# Species biology and demographic history determine species vulnerability to climate change in tropical island endemic birds

**DOI:** 10.1101/2024.11.14.623644

**Authors:** Ratnesh Karjee, Vikram Iyer, Durbadal Chatterjee, Rajasri Ray, Kritika M. Garg, Balaji Chattopadhyay

## Abstract

Climate change and associated habitat fluctuations can expedite the diversification of insular lineages or lead to isolation and extinction. Tropical island birds are a model system to assess responses to climate change owing to their insular nature and ability to diversify rapidly. While there is some understanding of the diversification of tropical island avian clades, we are yet to understand the vulnerability of these species to climate change. Long-term species genetic diversity and historical demography are critical predictors of this. Therefore, we investigated the sensitivity of tropical island endemic birds to climate change and analysed how species traits determine these responses by comparing species traits with demographic histories and paleohabitat fluctuations during the Last Glacial Period (LGP). From publicly available whole genome and paleoclimatic datasets, we reconstructed tropical island endemics’ past demographic histories (effective population size (Ne)) using Pairwise Sequential Markovian Coalescent (PSMC) (n = 23) and suitable habitat (n = 29) during the LGP and the Holocene. We observed that most species experienced an increase in suitable habitat between the Last Interglacial and the Last Glacial Maximum. However, a concomitant increase in Ne was only observed in the hyper-diverse passerine clade, attesting to their ability to rapidly diversify. Overall, diet specialists and large-bodied species showed a decrease in Ne during the LGP. Our results indicate that species traits dictate tropical island endemics’ demographic responses to climate, and a plastic response to habitat availability could be a consequence of clades’ abilities to rapidly occupy new niches and diversify. Further, our analyses revealed that most species entered the Holocene with low effective population sizes. Given that tropical island endemics have small geographic ranges and are groups vulnerable to climate change, special efforts are necessary to conserve them. We recommend that conservation management policies add components like historical demography and species traits while assessing extinction threats for island populations.

## INTRODUCTION

Tropical islands of the Indo-Australian Archipelago (IAA), the Indo-Pacific, and the Caribbean and Atlantic have had complex geological pasts that have affected their species distribution patterns. While several Pacific islands are true oceanic islands arising *de novo* from the seafloor, many islands of the Caribbean and the IAA were once connected with each other or to the continental landmasses during periods of global sea-level fall in the Quaternary (Lohman et al., 2011; Voris, 2000). These land bridges and the additional habitat they offered during periods of sea-level fall facilitated on one hand colonisations among islands and between islands and the mainland (Andersen et al., 2015; Cros et al., 2020; Irestedt et al., 2013; Moyle et al., 2009; Ng et al., 2017; Pujolar et al., 2022) and on the other, range expansions within islands in response to fluctuations in suitable habitat space. The habitat type and quality of these intermittently exposed land bridges varied, ranging from open savannah biomes to lowland forests (Cannon et al., 2009; Lohman et al., 2011). In the case of stratified elevational gradients, with tropical upland montane forests expanding onto lower slopes during periods of global cooling, upland montane species might have been able to disperse across otherwise persistent dual barriers of land and sea (Cannon et al., 2009).

Tropical islands’ geologic pasts make them an important natural system to study species responses to changing habitat at both ecological and evolutionary timescales. Birds are an ideal clade for this because they are taxonomically well-characterised, are ecologically well-studied, and charismatic. Avian lineages have arisen on tropical islands during the Pleistocene (Andersen et al., 2015; Garg et al., 2018; Irestedt et al., 2013; Moyle et al., 2009), concomitant with complex, clade-wise colonisation and recolonisation trajectories (Filardi and Moyle, 2005; Jønsson et al., 2008, 2011b, 2014) including colonisations of the mainland (Jønsson et al., 2011a) and repeated, independent colonisation events (Cibois et al., 2011). In birds, these responses to changing habitat are governed by species traits in several continental taxa and non-single-island endemics (Brüniche-Olsen et al., 2019, 2021), but responses in single-island endemics are poorly understood.

With such complex biogeographic pasts, tropical, single-island endemics represent either refugial populations, or *in-situ* diversifications as explained by taxon cycles (Ricklefs, 1970; Ricklefs and Cox, 1972, 1978). Being confined to single islands, these species are exceptionally vulnerable to ongoing anthropogenic climate change which is unprecedented and unlike the past Pleistocene and early Holocene climatic change (Crowley, 1990). This is exacerbated because we are currently experiencing a period of relatively high sea levels with reduced habitat availability for species resulting in population bottlenecks across taxa (Hewitt, 2000; Willis et al., 2004) including birds (Nadachowska-Brzyska et al., 2015; Smith et al., 2021). Information on tropical, single-island endemics’ demographic responses to past climate change can inform conservation efforts, owing to the genomic signatures that predispose a species to extinction (Mays et al., 2018; Spielman et al., 2004) and the fact that demographic history is directly correlated to extinction risk and resilience (Wilder et al., 2023).

Pairwise Sequential Markovian Coalescent (PSMC) is a powerful method to reconstruct the effective population size (Ne) for a species using a single diploid genome (Li and Durbin, 2011) and has been used to infer demographic history across taxa (Chattopadhyay et al., 2019; Kim et al., 2016; Kozma et al., 2016; Murray et al., 2017; Nadachowska-Brzyska et al., 2015). Using this along with paleo-ecological niche modelling allows us to directly correlate the demographic history of a species with its distributional range at different time points (Chattopadhyay et al., 2019). We perform these analyses on a global panel of tropical single-island endemics to understand the effects of past climatic changes on species.

## RESULTS

### Bird species panel

Our final species panel comprises 31 tropical single-island endemic bird species. Out of these, PSMC analyses were possible for 23 species and Ecological Niche Modelling (ENM) was possible for 29 species. Both PSMC and ENM analyses could be done successfully for 21 species (Table 1). These included eight Papuan species, five Philippine species, three Caribbean species, and three species from the Bismarck Archipelago. Of these 21 species, 14 species were passerines, with two from the white-eye family (Zosteropidae). Of the non-passerines, two species were parrots (family Psittacidae). All other families were represented by single species.

**Table 1:** Details of the taxa included in this study. PSMC = Pairwise Sequential Markovian Coalescent, ENM= Ecological Niche Modelling, y = yes, n = no.

| **Species** | **PSMC** | **ENM** | **Endemic**<br>**Region** | **Passerine** | **Family** |
| --- | --- | --- | --- | --- | --- |
| *Actenoides hombroni* | y | y | Philippines | n | Alcedinidae |
| *Aleadryas rufinucha* | y | y | Papua New<br>Guinea | y | Oreoicidae |
| *Alectura lathamii* | y | y | Australia | n | Megapodiidae |
| *Amazona guildingii* | y | y | St Vincent | n | Psittacidae |
| *Amazona vittata* | y | y | Puerto Rico | n | Psittacidae |
| *Amblyornis subalaris* | y | n | Papua New<br>Guinea | y | Ptilonorhynchi<br>dae |
| *Cacatua alba* | y | n | Indonesia | n | Cacatuidae |
| *Centropus unirufus* | y | y | Philippines | n | Cuculidae |
| *Chaetorhynchus*<br>*papuensis* | n | y | Papua New<br>Guinea | y | Rhipiduridae |
| *Cicinnurus regius* | y | y | Papua New<br>Guinea | y | Paradisaeidae |
| *Cnemophilus lorae* | y | y | Papua New<br>Guinea | y | Cnemophilida<br>e |
| *Dicaeum eximium* | y | y | Bismarck | y | Dicaeidae |
|  |  |  | Archipelago |  |  |
| *Diphyllodes magnificus* | y | y | Papua New Guinea | y | Paradisaeidae |
| *Eulacestoma nigropectus* | n | y | Papua New Guinea | y | Eulacestomati<br>dae |
| *Ifrita kowaldii* | n | y | Papua New Guinea | y | Ifritidae |
| *Irena cyanogastra* | y | y | Philippines | y | Irenidae |
| *Machaerirhynchus nigripectus* | y | y | Papua New Guinea | y | Machaerirhyn<br>chidae |
| *Melanocharis versteri* | y | y | Papua New Guinea | y | Melanochariti<br>dae |
| *Myiagra hebetior* | y | y | Bismarck Archipelago | y | Monarchidae |
| *Oreocharis arfaki* | n | y | Papua New Guinea | y | Paramythiidae |
| *Pachycephala philippinensis* | y | y | Philippines | y | Pachycephalid<br>ae |
| *Paradisaea raggiana* | y | y | Papua New Guinea | y | Paradisaeidae |
| *Paradisaea rubra* | n | y | Indonesia | y | Paradisaeidae |
| *Parotia lawesii* | n | y | Papua New Guinea | y | Paradisaeidae |
| *Pseudorectes*<br>*ferrugineus* | n | y | Papua New<br>Guinea | y | Pachycephalid<br>ae |
| *Ptilorrhoa leucosticta* | n | y | Papua New<br>Guinea | y | Cinclosomatid<br>ae |
| *Rhagologus*<br>*leucostigma* | y | y | Papua New<br>Guinea | y | Rhagologidae |
| *Rhynochetos jubatus* | y | y | New<br>Caledonia | n | Rhynochetida<br>e |
| *Sterrhoptilus*<br>*dennistouni* | y | y | Philippines | y | Zosteropidae |
| *Todus mexicanus* | y | y | Puerto Rico | n | Todidae |
| *Zosterops*<br>*hypoxanthus* | y | y | Bismarck<br>Archipelago | y | Zosteropidae |

### Paleo-habitat reconstruction

Among the 19 climatic variables used (Supplementary table S1), we observed precipitation of the warmest quarter (BIO18) to contribute the most for each island endemic bird species except for the Caribbean species *Amazona guildingii*. 25 out of the 29 bird species experienced an increase in suitable habitat from the Last Interglacial (LIG) to the Last Glacial Maximum (LGM) (Supplementary table S2). 12 out of 14 Papuan species showed an increase in habitat availability from the LIG to the LGM, with *Cicinnurus regius* and *Pseudorectus ferrugineus* as exceptions. All Philippine species showed an increase in habitat from the LIG to the LGM as well (Figure 1–2, Supplementary information S2). Two out of the three Caribbean species experienced a decline in suitable habitat during this period. Of the remaining species, all showed an increase in available habitat. Upland Papuan species remained confined to upland montane regions at all the time points we reconstructed available habitat, with *Rhagologus leucostigma* as a notable exception (Supplementary information S2). Papuan lowland species remained largely confined to regions which were not newly exposed land bridges at all times as well. From the LGM to the present day, habitat decreased for 20 out of the 29 bird species, and two species experienced no change (Supplementary table S2). For the present study, we primarily concentrate on habitat fluctuation between the LIG and the LGM as this is the period for which we have comparative evidence of fluctuations of both habitat as well as effective population size.

**Figure 1:**
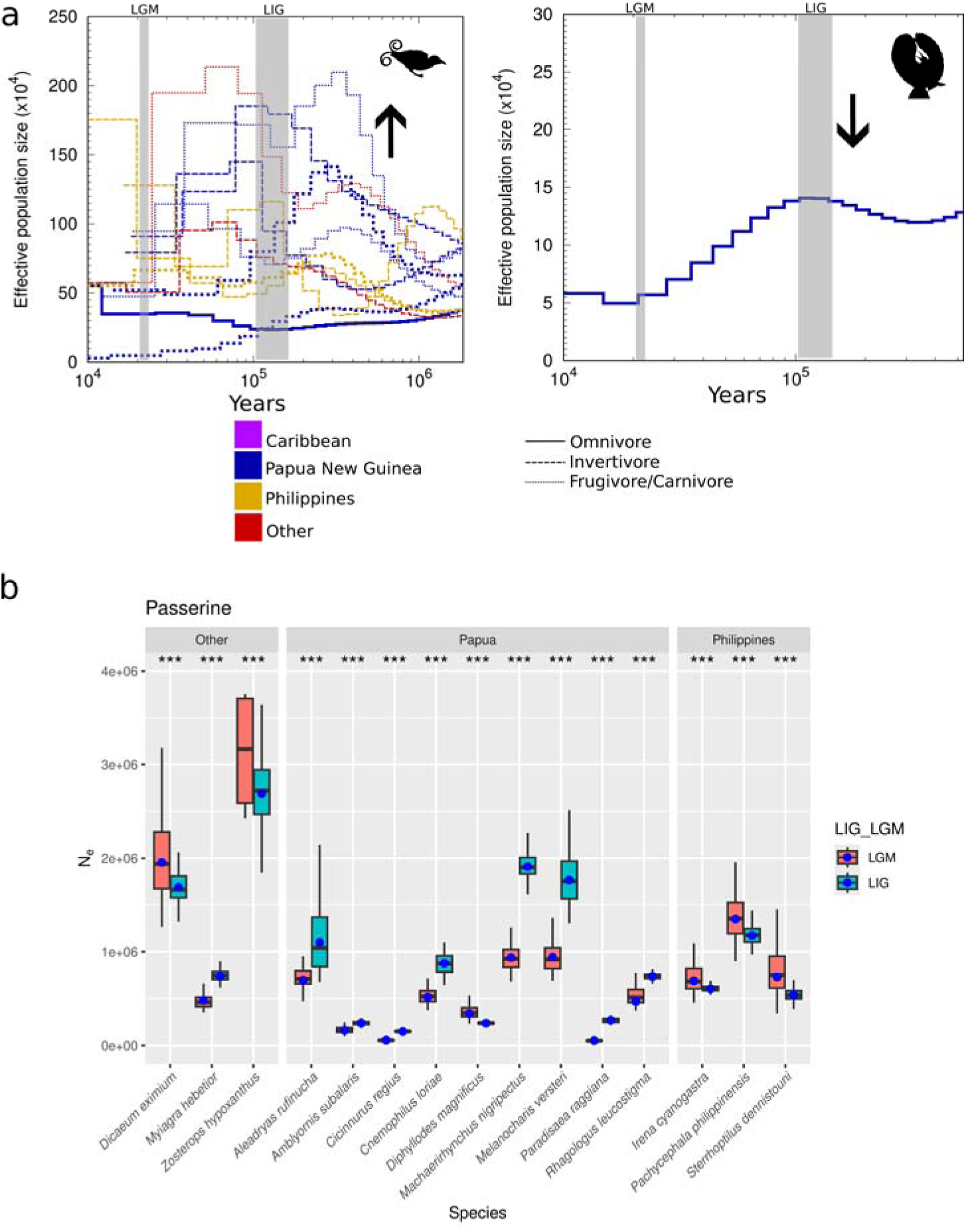
Effective population size for passerines. (a) Pairwise Sequential Markovian Coalescent (PSMC) plots using the settings –p “2 + 2 + 30 * 2 + 4 + 6 + 10” displaying reconstructed effective population size values with time for passerines based on whether habitat availability increased (left) or decreased (right) during the Last Glacial Period (LGP). Colours indicate the archipelago the bird belongs to, and the line style indicates the dietary habit of the bird species. Bold lines indicate large (> 50 g body mass) bird species. The grey bands indicate the approximate durations of the Last Interglacial (LIG) and the Last Glacial Maximum (LGM). Black arrows indicate if habitat availability increased or decreased during the LGP. A mutation rate of 1.4 x 10e–9 years/site and a generation time of 2 years for passerines, and a mutation rate of 1.91 x 10e–9 years/site and a generation time of 1 year for non-passerines were used to generate plots. Only species for which both Ecological Niche Modelling and PSMC analyses were possible are shown. *Zosterops hypoxanthus* is not displayed because its Ne values far exceed those of the other species. (b) Comparisons of Effective Population Size (Ne) at the Last Interglacial (LIG) and Last Glacial Maximum (LGM) incorporating bootstrapped Ne values generated using the same PSMC settings. Boxplots display bootstrapped Ne values, and blue points display the non-bootstrapped Ne value. Outlying bootstrapped and non-bootstrapped Ne values are not displayed. “***” indicates p < 0.001.

**Figure 2:**
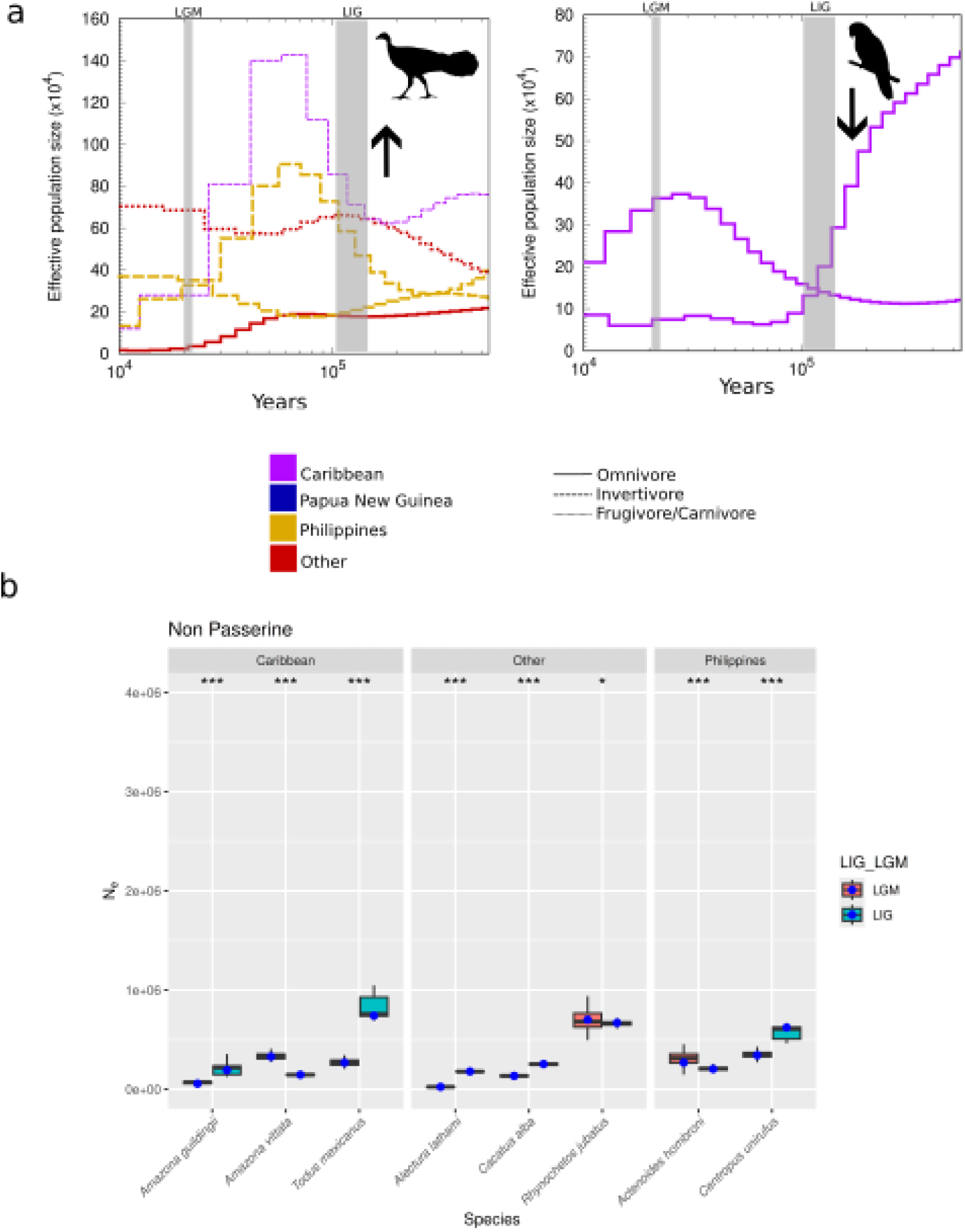
Effective population size for passerines. (a) Pairwise Sequential Markovian Coalescent (PSMC) plots using the settings –p “2 + 2 + 30 * 2 + 4 + 6 + 10” displaying reconstructed effective population size values with time for non passerines based on whether habitat availability increased (left) or decreased (right) during the Last Glacial Period (LGP). Colours indicate the archipelago the bird belongs to, and the line style indicates the dietary habit of the bird species. Bold lines indicate large (> 50 g body mass) bird species. The grey bands indicate the approximate durations of the Last Interglacial (LIG) and the Last Glacial Maximum (LGM). Black arrows indicate if habitat availability increased or decreased during the LGP. A mutation rate of 1.4 x 10e–9 years/site and a generation time of 2 years for passerines, and a mutation rate of 1.91 x 10e–9 years/site and a generation time of 1 year for non-passerines were used to generate plots. Only species for which both Ecological Niche Modelling and PSMC analyses were possible are shown. (b) Comparisons of Effective Population Size (Ne) at the Last Interglacial (LIG) and Last Glacial Maximum (LGM) incorporating bootstrapped Ne values generated using the same PSMC settings. Boxplots display bootstrapped Ne values, and blue points display the non-bootstrapped Ne value. Outlying bootstrapped and non-bootstrapped Ne values are not displayed. “***” indicates p < 0.001, and “*” indicates p < 0.05.

### Strong Quaternary fluctuations in effective population size

PSMC analyses could be successfully done for 23 species (Table 1). Results indicate large effective population size (Ne) variations in almost all bird species, corresponding to Quaternary climate shifts (Figure 3). The demographic history of these species reconstructed using PSMC analyses extends back over a million years (Supplementary information S1). Reconstructed Ne values were generally concordant across the three PSMC settings used, except for five species (*Centropus unirufus, Dicaeum eximium, Irena cyanogastra, Sterrhoptilus dennistouni, Zosterops hypoxanthus*) where Ne values had large differences across PSMC settings measured at the LIG and LGM (Supplementary table S3). However, in three of these species as well, trends of Ne increase or decrease from the LIG to LGM were robust across all three PSMC models considered (*Centropus unirufus, Irena cyanogastra, Sterrhoptilus dennistouni*) (Supplementary table S4). We found no significant difference between the different sets of PSMC settings used to estimate the change in Ne during the LGP (Kruskal-Wallis test, χ^2^= 0.11, DF = 2, p = 0.94).

**Figure 3:**
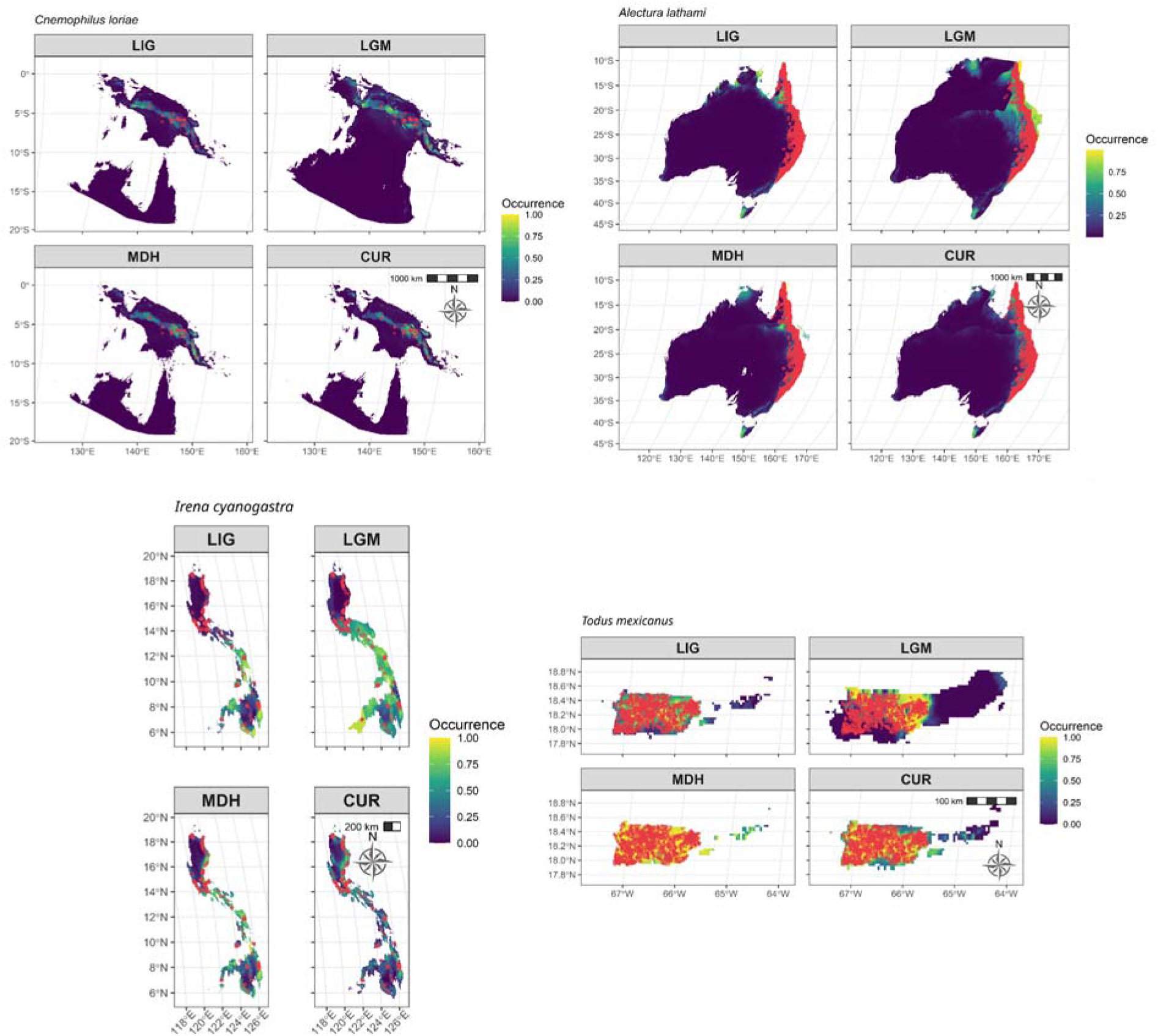
Example Ecological Niche Modelling plots for species from Papua (*Cnemophilus loriae)*, Australia (*Alectura lathami*), the Philippines (*Irena cyanogastra*), and the Caribbean (Puerto Rico, *Todus mexicanus*). LIG = Last Interglacial. LGM = Last Glacial Maximum. MDH = Mid Holocene. CUR = Current. The continuous heatmap represents the probability of occurrence of the species and red points are known occurrences from GBIF. For all the plots see Figure S2.

The 14 passerine species we analysed showed overall lower Ne values in the LGM as compared to the LIG (Figure 1–2). We obtained similar results for non-passerines as well, with *Rhynochetos jubatus* as a notable exception using the –p “4 + 30 * 2 + 4 + 6 + 10” setting, displaying a large peak followed by a crash in Ne (Supplementary information S1). This is likely a PSMC artefact, because plots using other parameter settings did not show a peak (Supplementary information S1). Along with overall lower values, we also see a smaller range of Ne values in the LGM as compared to the LIG (Figure 1–2).

### Correlation between historical fluctuations in Ne and distribution

Habitat change was poorly associated with change in Ne for the 21 species for which both PSMC and ENM analyses were possible (Cramer’s V = 0.11). However, when analysed separately, passerine species only showed a strong association (Cramer’s V = 0.98), while non-passerines showed a weak negative association (Cramer’s V = –0.15).

Bayesian multivariate regression models revealed that an overall fluctuation in Ne was associated with species biology and habitat change during the Last Glacial Period (LGP) even after controlling for geographical island group as a random effect (Figure 4). Most confidence intervals and beta coefficients did not overlap with zero (Figure 4) suggesting significant relationships. We found a positive association between change in habitat area (LIG to LGM) and Ne (β = 9.4, 95% CI: [1.95, 21.07]) and a negative relationship with body mass (β = −8.32, 95% CI: [–15.84, –2.59]). That is, large-bodied species showed decreases in Ne during the LGP. The change in Ne was positively correlated with the interaction term of habitat change and body mass (β = 10.28, 95% CI: [0.05, 25.18]). This suggests that habitat change and body mass act synergistically to predict the change in Ne. Diet specialists like frugivores (β = –11.89, 95% CI: [–22.57, –3.36]) and invertivores (β = –11.32, 95% CI: [–21.85, –2.85]) also showed significant negative associations with population change. However, omnivores (β = –1.28, 95% CI: [–12.77, 11.24]) did not show any association with population change. We found a marginally negative effect of clade identity as well (β = –3.64, 95% CI: [–7.91, –0.43]), that is, passerine status was associated with reducing Ne in the LGP in our dataset. Whether or not a species was a passerine was an important predictor of Ne in combination with the change in habitat from LIG to LGM (β = 8.01, 95% CI: [1.9, 17.11], suggesting that passerines respond positively to habitat change, as suggested by Cramer’s V as well. Finally, the random effects parameter for country showed that there exists country-to-country variation (β = 0.86, 95% CI: [0.04, 2.25]) (Figure 4, Supplementary table S5).

**Figure 4:**
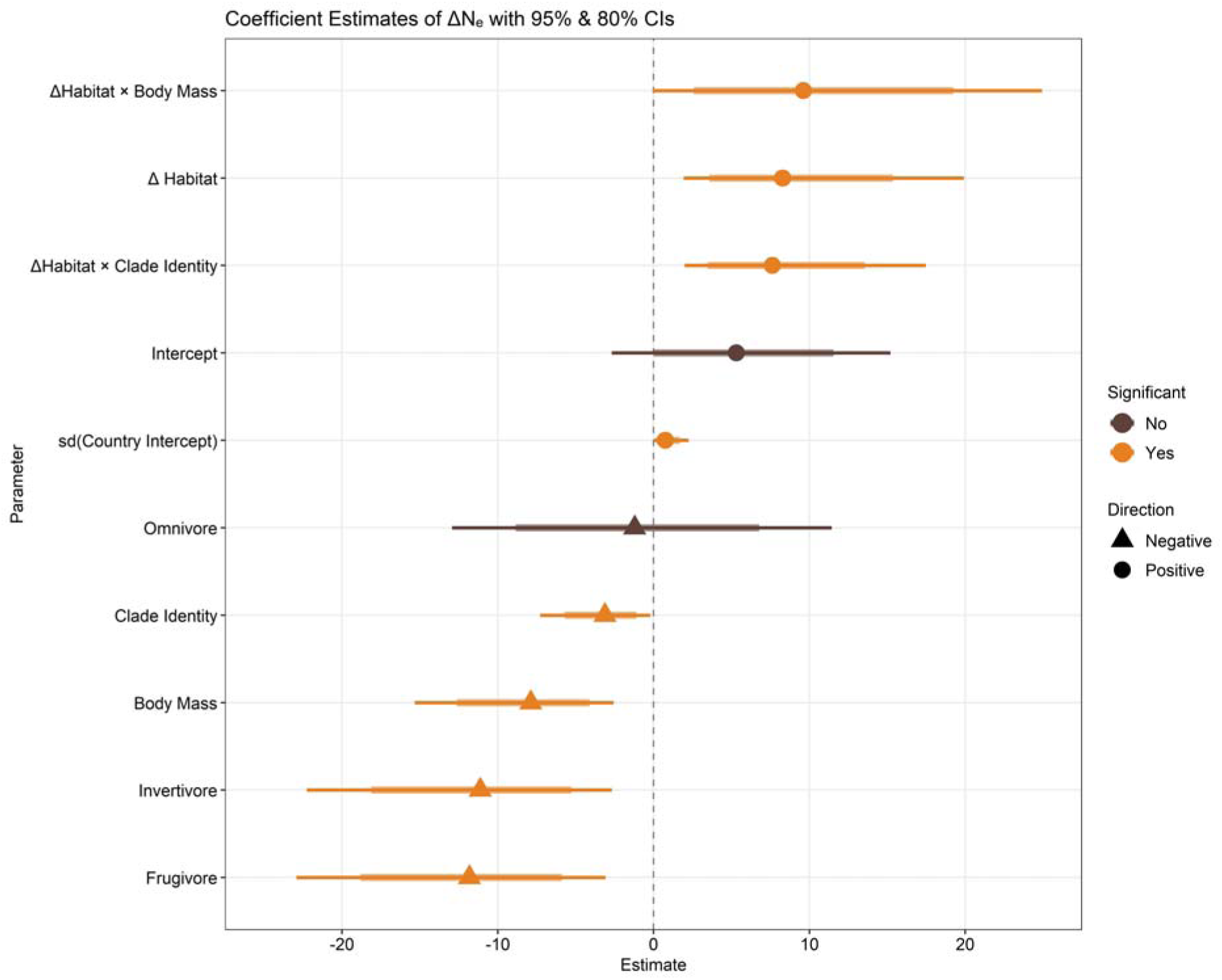
Results of the best Bayesian multivariate regression model performed. The response variable represents the changes in effective population size (increased (=1) or decreased (=0)) during the Last Glacial Period. Shapes represent mean values, thick lines represent the 80% confidence intervals, and thin lines represent 95% confidence intervals. Model parameters and coefficients are provided in table S6. ΔHabitat = the change in suitable habitat from the Last Interglacial to the Last Glacial Maximum.

## DISCUSSION

### Island birds are vulnerable to the effects of climate change

Our results on the fluctuations in paleohabitat and Ne during the LGP highlight the vulnerability of island birds to climatic fluctuations. Climate fluctuations change habitat availability and, in turn, Ne, making these species particularly sensitive to a changing environment. In response to landmass expansion, several species in our panel across islands showed an increase in their Ne during the LGP (Figure 1–2, Supplementary information S1), although the overall relationship was weak (Cramer’s V = 0.11). This could be because large amounts of intermittently exposed land bridge habitats remained uninhabited by the species in our panel (Supplementary figure S2). For example, we know that large amounts of exposed land bridge habitat available to our panel’s Papuan species ranged from lowland evergreen to seasonal rainforest, with an expansion of upland habitats of evergreen to seasonal rainforest. Transitional hill forest formed an ecotone between these two (Cannon et al., 2009). However, most of our species inhabit dense forest (Supplementary table), which is unlikely to comprise newly exposed landbridge habitat. Moreover, rainfall is overall lower in the LGM in the tropics including South and Southeast Asia, where most of our species are from (McGee, 2020), indicating a possible explanation of why we observed precipitation of the warmest quarter to be the largest contributing bioclimatic variable for all but one Caribbean species.

Our results also reveal that both passerine and non-passerine island endemics have entered the Holocene with low Ne, based on the PSMC plots (Figure 1–2). A loss of Ne predisposes species to extinction (Mays et al., 2018; Spielman et al., 2004) with a potentially causal relationship for birds (Evans and Sheldon, 2008). The current climate crisis has greatly imperilled biodiversity with the documented loss of many vertebrate species (Steadman and Martin, 2003; Tan et al., 2023; Willis et al., 2004). Fossil evidence has revealed the extinction of several species in the Caribbean since the LGM (Morgan and Woods, 1986; Orihuela et al., 2020; Steadman et al., 1984), and many of the species in our panel are Southeast Asian where rapid habitat loss is ongoing (Sodhi et al., 2004). Climate warming, sea level rise, and changes in vegetation are all associated with this loss, and flightless birds and endemics are particularly prone to extinction (Fromm and Meiri, 2021). Explicitly correlating past climate with demographic history—known to predict extinction risk (Wilder et al., 2023)—allows for a more robust validation of the latter.

Habitat loss, rise in sea level, and warming temperatures can rapidly accelerate species extinction, particularly in the tropics (Şekercioğlu et al., 2012). Effective population size, and in extension genetic diversity are associated with species survival and extinction risk (Frankham, 2005). Coalescent effective population size as used in PSMC is an important predictor of species vulnerability to extinction (Brüniche-Olsen et al., 2021; Wilder et al., 2023). Thus, an evolution-informed understanding of species vulnerability to climate change and their associations with paleohabitats can be an important tool to predict species vulnerability to the current climate crisis and future extinction risk (Brüniche-Olsen et al., 2021; Chattopadhyay et al., 2019; Gabrielli et al., 2024; Germain et al., 2023).

### A strong passerine association might be driven by their rapid diversification

Passerines are a hyperdiverse clade representing over 60% of extant avian diversity. The most recent, fossil-calibrated passerine phylogeny (Oliveros et al., 2019) shows that passerines began diversifying in the Middle (47 mya) to Late Eocene (38–39 mya) with increasing diversification rates suggested as we move towards the present. Crown passerines originated in the Australo-Pacific region (Oliveros et al., 2019), and this places the many Papuan and Australian species in our panel close to the passerine diversification centre which is known to have rapid diversification rates (McCullough et al., 2022). Because bird lineages are known to have arisen on tropical islands in the Pleistocene (Andersen et al., 2015; Irestedt et al., 2013; Moyle et al., 2009), it is likely that the Southeast Asian species in our panel represent newly arisen lineages over refugia, possibly belonging to currently rapidly diversifying clades. This is supported by the fact that several of these species belong to small, often oligotypic families such as Oreoicidae (*Aleadryas rufinucha*), Ptilonorhynchidae (*Amblyornis subalaris*), Cnemophilidae (*Cnemophilus loriae)*, Eulacestomatidae (*Eulacestoma nigropectus*), Ifritidae (*Ifrita kowaldi*), Machaerirhynchidae (*Machaerirhynchus nigripectus*), and Rhagologidae (*Rhagologus leucostigma*). Many of these species belong to poorly-studied tropical genera for which species-level divergence times are unavailable. The strong passerine association between change in Ne and available habitat area (Cramer’s V = 0.98; figure 4, β = 9.4) reveals that passerines are strongly plastic to the environment, most possibly as a consequence of their ability to rapidly diversify. This ability to diversify is reflected in their higher mutation rates as used in PSMC (Lanfear et al., 2010).

Non-passerines representing a paraphyletic outgroup do not show this trend (Cramer’s V = – 0.15). The seven non-passerine species for which both ENM and PSMC analyses were possible belong to six different families (Table 1), representing a diverse sample across the avian phylogeny and precluding non-passerine phylogenetic insight. Notably, we confirm (Cramer’s V = 0.11) that habitat area is positively correlated with genomic diversity in a clade non-specific manner for birds in general (Brüniche-Olsen et al., 2019, 2021).

### Species traits influenced historic fluctuations in Ne during the LGP

The interplay of species traits and habitat availability determine how Ne values fluctuate with time in non-endemic birds (Brüniche-Olsen et al., 2021), and our results confirm this for tropical island endemics as well. Overall, habitat change in the LGP was positively associated with Ne fluctuations (Figure 4, β = 9.4). While diet generalists like omnivores showed no significant association with Ne fluctuations, specialists like frugivores (Figure 4, β = –11.89) and invertivores (Figure 4, β = –11.32) showed significant negative associations with fluctuation in Ne, implying that specialist species tended to show decreases in Ne in the LGP. Previous work has shown that large-bodied species have lower values of standing genetic diversity at a given time-point (Brüniche-Olsen et al., 2019, 2021; Eo et al., 2011). We find large-bodied species to show a negative association with Ne fluctuations (Figure 4, β = –8.32) as well. Traits like body size and diet determine extinction risk (Ripple et al., 2017; Willis et al., 2004), and our results add to the understanding of how traits modulate Ne changes through time.

Assessing the relative contribution of species traits and habitat availability in determining Ne requires their quantitative measurements. Our methods generated values for available habitat at various time-points (Supplementary tableS2). However, Ne values can so far only be reconstructed using coalescent methods like PSMC. These values depend upon the parameter settings used. While we find these to be generally concordant across settings (Kruskal-Wallis, p = 0.94, Supplementary table S4), the precise, estimated values vary. We therefore chose to cautiously measure only the direction of change (increase or decrease) and not use numerical values.

### Caveats of PSMC analyses

Coalescent methods for inferring demographic history like PSMC assume no operant selection. Our bootstrapped PSMC estimates partially account for this by subsampling from across the genome and potentially breaking any linked regions under selection. Further, PSMC traces a local population’s demographic history rather than the entire species’ history (Gattepaille et al., 2013; Heller et al., 2013; Ptak and Przeworski, 2002). Population structure is thus a confounding factor. For the majority of the species in our panel, modelled habitat areas in the LGM were largely contiguous (Supplementary figure S2), making panmixia and reduced population structure a possibility, with a subsequent increase in isolation and population structure towards present day. Moreover, Papua is the only island in our panel large enough to likely support population structure.

Large peaks followed by apparent collapses in Ne have also recently been shown to be an artefact of PSMC analyses (Hilgers et al., 2024). We addressed this caveat by using three different parameter combinations and tested for consensus amongst the different parameters (increase/decrease/no change). Finally, hybridization may also increase the estimates of Ne, due to increase in heterozygosity. However, we currently lack species specific studies to address this issue. With additional population genomic data on island endemics, we can address this caveat in the future. Because of these potential caveats, we only infer the direction of population size change rather than actual estimates of Ne.

## METHODS

### Species selection

We queried Avibase (Lepage et al., 2014) for all species endemic to tropical islands and selected all species for which assembled whole genome sequences were available on GenBank using short-read data from the Illumina platform. We excluded genomes assembled from museum specimens. This resulted in a panel of 31 species (Table 1; Supplementary table S6–S7).

### Pairwise Sequential Markovian Coalescent (PSMC) analyses

For each assembled diploid genome, we identified the contigs corresponding to the sex chromosomes using Satsuma ver. 2.0 (Grabherr et al., 2010) by aligning contigs to a *Gallus gallus* (chicken; BioSample: SAMN02981218) genome as a reference. All contigs mapping to the sex chromosomes were removed following standard practice (Hawkins et al., 2018; Liu and Hansen, 2016). Next, we obtained all available Illumina short reads for each species from the Sequence Read Archive (SRA) database of the NCBI, and checked them for errors using FastQC (Andrews, 2010). We used Trimmomatic (Bolger et al., 2014) for preprocessing and trimming the reads using the parameter settings “ILLUMINACLIP:TruSeq3-PE.fa:2:30:10:2:True LEADING:3 TRAILING:3 SLIDINGWINDOW:4:20 MINLEN:36”.

The cleaned reads were aligned to the autosomes using BWA-MEM ver. 0.7.17 (Li, 2013) using default parameters. Next, we used SAMtools ver. 1.10 (Danecek et al., 2021) to merge all the resulting files to generate a single bam file for each species. We further sorted and removed duplicates using SAMtools and estimated the depth of coverage for each species. We excluded species with a depth of coverage < 18x. This is a sufficient value of coverage for PSMC analyses (Li and Durbin, 2011). Next, we implemented the SAMtools mpileup-bcftools pipeline to identify SNPs. The minimum and maximum depths for calling SNPs was set to one-third and double the mean depth respectively following Nadachowska-Brzyska et al. (Nadachowska-Brzyska et al., 2015).

However, because sex chromosome reads could potentially map to other regions of the genome, we also performed PSMC using an alternate method. For this, we directly mapped raw reads files onto the genome and then called SNPs on only autosomal regions using the SAMtools mpileup-bcftools pipeline, after which we performed PSMC as above (Supplementary Information S3–4). This was done for five species. Results between the two methods were not significantly different (Supplementary information S3–4).

We used three sets of parameters for PSMC analyses and looked for consensus trends amongst them because PSMC analyses can result in spurious peaks of effective population size (Hilgers et al., 2024). We used the following parameter sets for PSMC analyses: –t5 –b –r1 –p “4 + 30 * 2 + 4 + 6 + 10”, –t5 –b –r1 –p “2 + 2 + 30 * 2 + 4 + 6 + 10”, and –t5 –b –r1 –p “1 + 1 + 1 + 1 + 30 * 2 + 4 + 6 + 10”. These parameter options were chosen following Nadachowska-Brzyska et al. (2015). We performed 30 iterations for parameter optimisation and ran 100 bootstrap replicates using the “splitfa” PSMC utility on the psmcfa file to judge the uncertainty in our estimates. Bootstrapping was done by randomly sampling chromosome segments with replacement and running PSMC on them. To plot the results from the PSMC analyses, we used previously estimated values of mutation rates and generation times for passerines (1.4 x 10e–9 years/site and 2 years respectively; (Ellegren et al., 2012; Nadachowska-Brzyska et al., 2016) and non-passerines (1.91 x 10e–9 years/site and 1 year respectively) (Nam et al., 2010) (Supplementary material S1, supplementary table S3).

We extracted the precise values of the population scaled mutation rate (theta) from the bootstrapped and non-bootstrapped .psmc files and used these to calculate the precise values of Ne at the LIG and LGM (Supplementary information S5). We defined the LIG to be equal to 108,000–143,000 years ago, and the LGM to be equal to the atomic time interval closest to 20,000 years ago. We performed this for all three PSMC settings used and compared boxplots of bootstrapped PSMC values. We further performed the Kruskal-Wallis test for comparing across the three different PSMC settings using the “ggstatsplot” package in R (ver. 0.13.0; Patil, 2021).

### Species and climate records

We performed reconstructions of paleohabitats through ecological niche models and reconstructed species distributions from four time periods: the Last Interglacial (LIG, approx. 110,000–130,000 years ago), the Last Glacial Maximum (LGM, approx. 20,000 years ago), the Mid-Holocene (MDH, approx. 6000 years ago) and Current (CUR, present day).

We used R (ver. 4.2.1; R Core Team, 2021) for all paleo-habitat modelling analyses. We accessed the location records of our bird species from the Global Biodiversity Information Facility (GBIF) in September, 2023 using the “rbgif” package (ver. 3.7.9) (Chamberlain and Boettiger, 2017) in R. We discarded duplicates and retained only human observed records using the “tidyverse” (v-2.0.0) package (Wickham and RStudio, 2023) followed by the ‘CoordinateCleaner’ package (v-3.0.1) (Zizka et al., 2023). We used the “spThin” package (Aiello-Lammens et al., 2014) to account for spatial autocorrelation and finally discarded spurious distribution records like records overlaying water bodies, buildings, roadways, and railways using QGIS (v-3.34+; https://www.qgis.org/). GBIF dataset keys used to access all the data points are in Supplementary table S8.

We used 19 climate variables (labelled BIO1 – BIO19; Supplementary table S4) for the four different time periods considered. We downloaded CHELSA (https://chelsa-climate.org/; Karger et al., 2021) climate predictors at a resolution of 2.5 arc-minutes (∼5 km) for all regions.

We followed Chattopadhyay et al., (2019) and used a global dataset approach to account for idiosyncratic biases due to smaller datasets to extract the location points of each endemic-island group against each climatic variable. We tested for multicollinearity using variance inflation factor (VIF) using the ‘usdm’ R package (Naimi, 2014) and considered variables for further analyses if their VIF value was ≤ 5 (Shrestha, 2020).

### Habitat Suitability Modelling and Area Calculation

We used R to reconstruct paleo-climatic suitable habitat for 29 endemic island birds through ecological niche models for the four time-periods considered. For this, we accessed the Global Biodiversity Information Facility (GBIF) to extract our species’ occurrence records. We could not perform ENM analyses for two species due to a paucity in the number of GBIF occurrence points for them (Table 1).

### Data partitioning and model evaluation

For each species, we used available occurrence records from the present day to generate pseudo-absence data points. We generated 500–10,000 pseudo-absence/background points for each species depending upon the area of the island (Supplementary table S6), except for *Amazona guildingii* endemic to Saint Vincent where we generated 15 points. Saint Vincent’s small land area could not accommodate more points than this. We used a subset of bioclimatic variables to generate a weighted average ensemble species distribution model (eSDM) for each island group. eSDMs were implemented in the R package “sdm” (v-1.2.37; Naimi and Araujo, 2016) applying the ‘MaxEnt’, ‘GLM’, and ‘BRT’ algorithms. eSDMs account for the limitations of different models by generating a weighted average of multiple models (Araújo and New, 2007; Dormann et al., 2018; Naimi and Araujo, 2016). We used k-fold cross-validation (CV) with replication for each method for training and test datasets across each endemic island group (Supplementary table S9). Model accuracy was measured using the Area Under the Curve (AUC) and True Skill Statistic (TSS) metrics. We first performed the above analyses for the present-day (CUR) distribution using the weighted-average of AUC and TSS. AUC and TSS values greater than 0.9 and 0.75 respectively have been shown to be indicators of superior model performance (Ahmad et al., 2019) and we chose a similar threshold (AUC ≥ 0.9 & TSS ≥ 0.8) to get the best ensemble model for most of the species except for 10 species for which data quality was poor (Supplementary table S9). Models for the other time points: the LIG, LGM, and MDH, were generated using the eSDM model generated based on CUR data.

### Suitable area analysis

All spatial analyses were carried out using the R package ‘terra’ (ver. 1.7.13) (Hijmans et al., 2024). Equal area projections were used to calculate the absolute suitable area of each species across the four time-periods. To account for a spherical Earth and the limitations of landmass depiction of the original eSDM raster (World Geodetic System 1984) we reprojected it into a pseudocylindrical projection (Eckert IV). Further, we used the average quantile threshold pixels to measure the availability of absolute suitable areas for the LIG, LGM, MDH, and CUR periods (Supplementary table S2).

### Statistical Analyses

We checked for associations between Ne and habitat change during the Last Glacial Period (from LIG to LGM) (LGP) using Cramer’s V implemented in the“polycor” package in R (Fox, 2022). Cramver’s V varies between –1 to 1, with values closer to 0 suggesting no-correlation. This was done for the entire dataset, and then for passerines and non-passerines separately.

We further explored the effects of species biology, habitat, and phylogenetic constraint on the fluctuation in Ne during the LGP using Bayesian Multilevel Models (MLMs), using the “brms” package in R (Bürkner, 2018). Our fixed effect predictors were habitat change from the LIG to LGM, diet (invertivore, frugivore, or omnivore; Supplementary table S6), body mass, and clade identity (passerine and non-passerine). The latter allows us to check if belonging to the rapidly diversifying passerine clade results in a signal. We also included country (the island/archipelago a species is endemic to) as a random effect variable for the analysis to account for country-specific effects. For the response variable i.e., the change in Ne, a Bernoulli distribution with a logit link was used because it is a binary response variable. Thus, species traits, clade identity, and change in habitat were three sets of independent predictor variables, with the change in Ne as the response variable.

We ran a series of models (n=12) to find the best-fit model (brms11, Supplementary table S10) based on leave-one-out cross-validation. We generated a total of 16,000 posterior samples by using the No-U-Turn Sampler (Hamiltonian Monte Carlo) algorithm with 4 chains with 5000 iterations each (with 1000 warm-up iterations; supplementary Information S6). Further, we estimated the posterior parameters with 95% confidence intervals to find negative or positive associations between the Ne and its predictors. In addition, we assessed the posterior convergence of the sample through the R-hat statistic (*R*^ = 1) where values close to 1 suggest an ideal convergence. We also reported the effective sample size for bulk and tail distributions (Supplementary table S9).

## ACKNOWLEDGEMENTS

BC acknowledges the support of the Trivedi School of Biosciences. KMG acknowledges the support from the DBT-Ramalingaswami Fellowship (No. BT/HRD/35/02/2006). VI is supported by DBT grant (BT/PR42830/BRB/10/1995/2021) to KMG.

## DATA AVAILABILITY

The raw data used for this study are available on NCBI and details of the accession IDs are provided in Supplementary table S7. All codes used in this study are provided on the Zenodo URL https://doi.org/10.5281/zenodo.14603965.

## REFERENCES

1. Ahmad, R., Khuroo, A.A., Charles, B., Hamid, M., Rashid, I., Aravind, N.A., 2019. Global distribution modelling, invasion risk assessment and niche dynamics of Leucanthemum vulgare (Ox-eye Daisy) under climate change. Sci. Rep. 9, 11395. 10.1038/s41598-019-47859-1

2. Aiello-Lammens, M.E., Boria, R.A., Radosavljevic, A., Vilela, B., Anderson, R.P., 2014. spThin: Functions for Spatial Thinning of Species Occurrence Records for Use in Ecological Models. 10.32614/CRAN.package.spThin

3. Andersen, M.J., Shult, H.T., Cibois, A., Thibault, J.-C., Filardi, C.E., Moyle, R.G., 2015. Rapid diversification and secondary sympatry in Australo-Pacific kingfishers (Aves: Alcedinidae: *Todiramphus*). R. Soc. Open Sci. 2, 140375. 10.1098/rsos.140375

4. Andrews, S., 2010. FastQC: A Quality Control Tool for High Throughput Sequence Data.

5. Araújo, M.B., New, M., 2007. Ensemble forecasting of species distributions. Trends Ecol. Evol. 22, 42–47. 10.1016/j.tree.2006.09.010

6. Bolger, A.M., Lohse, M., Usadel, B., 2014. Trimmomatic: a flexible trimmer for Illumina sequence data. Bioinformatics 30, 2114–2120. 10.1093/bioinformatics/btu170

7. Brüniche-Olsen, A., Kellner, K.F., DeWoody, J.A., 2019. Island area, body size and demographic history shape genomic diversity in Darwin’s finches and related tanagers. Mol. Ecol. 28, 4914–4925. 10.1111/mec.15266

8. Brüniche-Olsen, A., Kellner, K.F., Belant, J.L., DeWoody, J.A., 2021. Life-history traits and habitat availability shape genomic diversity in birds: implications for conservation. Proc. R. Soc. B Biol. Sci. 288, 20211441. 10.1098/rspb.2021.1441

9. Bürkner, P.-C., 2018. Advanced Bayesian Multilevel Modeling with the R Package brms. R J. 10, 395–411.

10. Cannon, C.H., Morley, R.J., Bush, A.B.G., 2009. The current refugial rainforests of Sundaland are unrepresentative of their biogeographic past and highly vulnerable to disturbance. Proc. Natl. Acad. Sci. 106, 11188–11193. 10.1073/pnas.0809865106

11. Chamberlain, S., Boettiger, C., 2017. R Python, and Ruby clients for GBIF species occurrence data. PeerJ Prepr.

12. Chattopadhyay, B., Garg, K.M., Ray, R., Rheindt, F.E., 2019. Fluctuating fortunes: genomes and habitat reconstructions reveal global climate-mediated changes in bats’ genetic diversity. Proc. R. Soc. B Biol. Sci. 286, 20190304. 10.1098/rspb.2019.0304

13. Cibois, A., Beadell, J.S., Graves, G.R., Pasquet, E., Slikas, B., Sonsthagen, S.A., Thibault, J.-C., Fleischer, R.C., 2011. Charting the course of reed-warblers across the Pacific islands. J. Biogeogr. 38, 1963–1975. 10.1111/j.1365-2699.2011.02542.x

14. Cros, E., Chattopadhyay, B., Garg, K.M., Ng, N.S.R., Tomassi, S., Benedick, S., Edwards, D.P., Rheindt, F.E., 2020. Quaternary land bridges have not been universal conduits of gene flow. Mol. Ecol. 29, 2692–2706. 10.1111/mec.15509

15. Crowley, T.J., 1990. Are There Any Satisfactory Geologic Analogs for a Future Greenhouse Warming?

16. Danecek, P., Bonfield, J.K., Liddle, J., Marshall, J., Ohan, V., Pollard, M.O., Whitwham, A., Keane, T., McCarthy, S.A., Davies, R.M., Li, H., 2021. Twelve years of SAMtools and BCFtools. GigaScience 10, giab008. 10.1093/gigascience/giab008

17. Dormann, C.F., Calabrese, J.M., Guillera-Arroita, G., Matechou, E., Bahn, V., Bartoń, K., Beale, C.M., Ciuti, S., Elith, J., Gerstner, K., Guelat, J., Keil, P., Lahoz-Monfort, J.J., Pollock, L.J., Reineking, B., Roberts, D.R., Schröder, B., Thuiller, W., Warton, D.I., Wintle, B.A., Wood, S.N., Wüest, R.O., Hartig, F., 2018. Model averaging in ecology: a review of Bayesian, information-theoretic, and tactical approaches for predictive inference. Ecol. Monogr. 88, 485–504. 10.1002/ecm.1309

18. Ellegren, H., Smeds, L., Burri, R., Olason, P.I., Backström, N., Kawakami, T., Künstner, A., Mäkinen, H., Nadachowska-Brzyska, K., Qvarnström, A., Uebbing, S., Wolf, J.B.W., 2012. The genomic landscape of species divergence in *Ficedula* flycatchers. Nature 491, 756–760. 10.1038/nature11584

19. Eo, S.H., Doyle, J.M., DeWoody, J.A., 2011. Genetic diversity in birds is associated with body mass and habitat type. J. Zool. 283, 220–226. 10.1111/j.1469-7998.2010.00773.x

20. Evans, S.R., Sheldon, B.C., 2008. Interspecific Patterns of Genetic Diversity in Birds: Correlations with Extinction Risk. Conserv. Biol. 22, 1016–1025. 10.1111/j.1523-1739.2008.00972.x

21. Filardi, C.E., Moyle, R.G., 2005. Single origin of a pan-Pacific bird group and upstream colonization of Australasia. Nature 438, 216–219. 10.1038/nature04057

22. Fox, J., 2022. polycor: Polychoric and Polyserial Correlations.

23. Frankham, R., 2005. Genetics and extinction. Biol. Conserv. 126, 131–140. 10.1016/j.biocon.2005.05.002

24. Frankham, R., 1995. Effective population size/adult population size ratios in wildlife: a review. Genet. Res. 66, 95–107. 10.1017/S0016672300034455

25. Fromm, A., Meiri, S., 2021. Big, flightless, insular and dead: Characterising the extinct birds of the Quaternary. J. Biogeogr. 48, 2350–2359. 10.1111/jbi.14206

26. Gabrielli, M., Leroy, T., Salmona, J., Nabholz, B., Milá, B., Thébaud, C., 2024. Demographic responses of oceanic island birds to local and regional ecological disruptions revealed by whole-genome sequencing. Mol. Ecol. 33, e17243. 10.1111/mec.17243

27. Garg, K.M., Chattopadhyay, B., Wilton, P.R., Malia Prawiradilaga, D., Rheindt, F.E., 2018. Pleistocene land bridges act as semipermeable agents of avian gene flow in Wallacea. Mol. Phylogenet. Evol. 125, 196–203. 10.1016/j.ympev.2018.03.032

28. Gattepaille, L.M., Jakobsson, M., Blum, M.G., 2013. Inferring population size changes with sequence and SNP data: lessons from human bottlenecks. Heredity 110, 409–419. 10.1038/hdy.2012.120

29. Germain, R.R., Feng, S., Chen, G., Graves, G.R., Tobias, J.A., Rahbek, C., Lei, F., Fjeldså, J., Hosner, P.A., Gilbert, M.T.P., Zhang, G., Nogués-Bravo, D., 2023. Species-specific traits mediate avian demographic responses under past climate change. Nat. Ecol. Evol. 7, 862–872. 10.1038/s41559-023-02055-3

30. Grabherr, M.G., Russell, P., Meyer, M., Mauceli, E., Alföldi, J., Di Palma, F., Lindblad-Toh, K., 2010. Genome-wide synteny through highly sensitive sequence alignment: Satsuma. Bioinformatics 26, 1145–1151. 10.1093/bioinformatics/btq102

31. Hawks, J., 2017. Introgression Makes Waves in Inferred Histories of Effective Population Size. Human Biology 89, 67–80.

32. Hawkins, M.T.R., Culligan, R.R., Frasier, C.L., Dikow, R.B., Hagenson, R., Lei, R., Louis, E.E., 2018. Genome sequence and population declines in the critically endangered greater bamboo lemur (*Prolemur simus*) and implications for conservation. BMC Genomics 19, 445. 10.1186/s12864-018-4841-4

33. Heller, R., Chikhi, L., Siegismund, H.R., 2013. The Confounding Effect of Population Structure on Bayesian Skyline Plot Inferences of Demographic History. PLOS ONE 8, e62992. 10.1371/journal.pone.0062992

34. Hewitt, G., 2000. The genetic legacy of the Quaternary ice ages. Nature 405, 907–913. 10.1038/35016000

35. Hijmans, R.J., Bivand, R., Dyba, K., Pebesma, E., Sumner, M.D., 2024. terra: Spatial Data Analysis.

36. Hilgers, L., Liu, S., Jensen, A., Brown, T., Cousins, T., Schweiger, R., Guschanski, K., Hiller, M., 2024. Avoidable false PSMC population size peaks occur across numerous studies. 10.1101/2024.06.17.599025

37. Irestedt, M., Fabre, P.-H., Batalha-Filho, H., Jønsson, K.A., Roselaar, C.S., Sangster, G., Ericson, P.G.P., 2013. The spatio-temporal colonization and diversification across the Indo-Pacific by a ‘great speciator’ (Aves, *Erythropitta erythrogaster*). Proc. R. Soc. B Biol. Sci. 280, 20130309. 10.1098/rspb.2013.0309

38. Jønsson, K.A., Fabre, P.-H., Ricklefs, R.E., Fjeldså, J., 2011a. Major global radiation of corvoid birds originated in the proto-Papuan archipelago. Proc. Natl. Acad. Sci. 108, 2328–2333. 10.1073/pnas.1018956108

39. Jønsson, K.A., Irestedt, M., Bowie, R.C.K., Christidis, L., Fjeldså, J., 2011b. Systematics and biogeography of Indo-Pacific ground-doves. Mol. Phylogenet. Evol. 59, 538–543. 10.1016/j.ympev.2011.01.007

40. Jønsson, K.A., Irestedt, M., Christidis, L., Clegg, S.M., Holt, B.G., Fjeldså, J., 2014. Evidence of taxon cycles in an Indo-Pacific passerine bird radiation (Aves: *Pachycephala*). Proc. R. Soc. B Biol. Sci. 281, 20131727. 10.1098/rspb.2013.1727

41. Jønsson, K.A., Irestedt, M., Fuchs, J., Ericson, P.G.P., Christidis, L., Bowie, R.C.K., Norman, J.A., Pasquet, E., Fjeldså, J., 2008. Explosive avian radiations and multi-directional dispersal across Wallacea: Evidence from the Campephagidae and other Crown Corvida (Aves). Mol. Phylogenet. Evol. 47, 221–236. 10.1016/j.ympev.2008.01.017

42. Karger, D.N., Wilson, A.M., Mahony, C., Zimmermann, N.E., Jetz, W., 2021. Global daily 1 km land surface precipitation based on cloud cover-informed downscaling. Sci. Data 8, 307. 10.1038/s41597-021-01084-6

43. Kim, Soonok, Cho, Y.S., Kim, H.-M., Chung, O., Kim, H., Jho, S., Seomun, H., Kim, J., Bang, W.Y., Kim, C., An, J., Bae, C.H., Bhak, Y., Jeon, S., Yoon, H., Kim, Y., Jun, J., Lee, HyeJin, Cho, S., Uphyrkina, O., Kostyria, A., Goodrich, J., Miquelle, D., Roelke, M., Lewis, J., Yurchenko, A., Bankevich, A., Cho, J., Lee, S., Edwards, J.S., Weber, J.A., Cook, J., Kim, Sangsoo, Lee, Hang, Manica, A., Lee, I., O’Brien, S.J., Bhak, J., Yeo, J.-H., 2016. Comparison of carnivore, omnivore, and herbivore mammalian genomes with a new leopard assembly. Genome Biol. 17, 211. 10.1186/s13059-016-1071-4

44. Kozma, R., Melsted, P., Magnússon, K.P., Höglund, J., 2016. Looking into the past – the reaction of three grouse species to climate change over the last million years using whole genome sequences. Mol. Ecol. 25, 570–580. 10.1111/mec.13496

45. Lanfear, R., Ho, S.Y.W., Love, D., Bromham, L., 2010. Mutation rate is linked to diversification in birds. Proceedings of the National Academy of Sciences 107, 20423–20428. 10.1073/pnas.1007888107

46. Lepage, D., Vaidya, G., Guralnick, R., 2014. Avibase – a database system for managing and organizing taxonomic concepts. ZooKeys 420, 117–135. 10.3897/zookeys.420.7089

47. Li, H., 2013. Aligning sequence reads, clone sequences and assembly contigs with BWA-MEM. 10.48550/arXiv.1303.3997

48. Li, H., Durbin, R., 2011. Inference of human population history from individual whole-genome sequences. Nature 475, 493–496. 10.1038/nature10231

49. Liu, S., Hansen, M.M., 2017. PSMC (pairwise sequentially Markovian coalescent) analysis of RAD (restriction site associated DNA) sequencing data. Molecular Ecology Resources 17, 631–641. 10.1111/1755-0998.12606

50. Lohman, D.J., Bruyn, M. de, Page, T., Rintelen, K. von, Hall, R., Ng, P.K.L., Shih, H.-T., Carvalho, G.R., Rintelen, T. von, 2011. Biogeography of the Indo-Australian Archipelago. Annu. Rev. Ecol. Evol. Syst. 42, 205–226. 10.1146/annurev-ecolsys-102710-145001

51. Mays, H.L., Hung, C.-M., Shaner, P.-J., Denvir, J., Justice, M., Yang, S.-F., Roth, T.L., Oehler, D.A., Fan, J., Rekulapally, S., Primerano, D.A., 2018. Genomic Analysis of Demographic History and Ecological Niche Modeling in the Endangered Sumatran Rhinoceros *Dicerorhinus sumatrensis*. Curr. Biol. 28, 70–76.e4. 10.1016/j.cub.2017.11.021

52. McCullough, J.M., Oliveros, C.H., Benz, B.W., Zenil-Ferguson, R., Cracraft, J., Moyle, R.G., Andersen, M.J., 2022. Wallacean and Melanesian Islands Promote Higher Rates of Diversification within the Global Passerine Radiation Corvides. Syst. Biol. 71, 1423–1439. 10.1093/sysbio/syac044

53. McGee, D., 2020. Glacial–Interglacial Precipitation Changes. Annual Review of Marine Science 12, 525–557. 10.1146/annurev-marine-010419-010859

54. Morgan, G.S., Woods, C.A., 1986. Extinction and the zoogeography of West Indian land mammals. Biol. J. Linn. Soc. 28, 167–203. 10.1111/j.1095-8312.1986.tb01753.x

55. Moyle, R.G., Filardi, C.E., Smith, C.E., Diamond, J., 2009. Explosive Pleistocene diversification and hemispheric expansion of a “great speciator.” Proc. Natl. Acad. Sci. 106, 1863–1868. 10.1073/pnas.0809861105

56. Murray, G.G.R., Soares, A.E.R., Novak, B.J., Schaefer, N.K., Cahill, J.A., Baker, A.J., Demboski, J.R., Doll, A., Da Fonseca, R.R., Fulton, T.L., Gilbert, M.T.P., Heintzman, P.D., Letts, B., McIntosh, G., O’Connell, B.L., Peck, M., Pipes, M.-L., Rice, E.S., Santos, K.M., Sohrweide, A.G., Vohr, S.H., Corbett-Detig, R.B., Green, R.E., Shapiro, B., 2017. Natural selection shaped the rise and fall of passenger pigeon genomic diversity. Science 358, 951–954. 10.1126/science.aao0960

57. Nadachowska-Brzyska, K., Burri, R., Smeds, L., Ellegren, H., 2016. PSMC analysis of effective population sizes in molecular ecology and its application to black-and-white *Ficedula* flycatchers. Mol. Ecol. 25, 1058–1072. 10.1111/mec.13540

58. Nadachowska-Brzyska, K., Li, C., Smeds, L., Zhang, G., Ellegren, H., 2015. Temporal Dynamics of Avian Populations during Pleistocene Revealed by Whole-Genome Sequences. Curr. Biol. 25, 1375–1380. 10.1016/j.cub.2015.03.047

59. Naimi B, Hamm Na, Groen TA, Skidmore AK, Toxopeus AG (2014). Where is positional uncertainty a problem for species distribution modelling. Ecography, 37, 191–203. doi:10.1111/j.1600-0587.2013.00205.x

60. Naimi, B., Araujo, M.B., 2016. sdm: a reproducible and extensible R platform for species distribution modelling. Ecography 39, 368–375. 10.1111/ecog.01881

61. Nam, K., Mugal, C., Nabholz, B., Schielzeth, H., Wolf, J.B., Backström, N., Künstner, A., Balakrishnan, C.N., Heger, A., Ponting, C.P., Clayton, D.F., Ellegren, H., 2010. Molecular evolution of genes in avian genomes. Genome Biol. 11, R68. 10.1186/gb-2010-11-6-r68

62. Ng, N.S.R., Wilton, P.R., Prawiradilaga, D.M., Tay, Y.C., Indrawan, M., Garg, K.M., Rheindt, F.E., 2017. The effects of Pleistocene climate change on biotic differentiation in a montane songbird clade from Wallacea. Mol. Phylogenet. Evol. 114, 353–366. 10.1016/j.ympev.2017.05.007

63. Oliveros, C.H., Field, D.J., Ksepka, D.T., Barker, F.K., Aleixo, A., Andersen, M.J., Alström, P., Benz, B.W., Braun, E.L., Braun, M.J., Bravo, G.A., Brumfield, R.T., Chesser, R.T., Claramunt, S., Cracraft, J., Cuervo, A.M., Derryberry, E.P., Glenn, T.C., Harvey, M.G., Hosner, P.A., Joseph, L., Kimball, R.T., Mack, A.L., Miskelly, C.M., Peterson, A.T., Robbins, M.B., Sheldon, F.H., Silveira, L.F., Smith, B.T., White, N.D., Moyle, R.G., Faircloth, B.C., 2019. Earth history and the passerine superradiation. Proc. Natl. Acad. Sci. 116, 7916–7925. 10.1073/pnas.1813206116

64. Orihuela, J., Viñola, L.W., Jiménez Vázquez, O., Mychajliw, A.M., Hernández de Lara, O., Lorenzo, L., Soto-Centeno, J.A., 2020. Assessing the role of humans in Greater Antillean land vertebrate extinctions: New insights from Cuba. Quat. Sci. Rev. 249, 106597. 10.1016/j.quascirev.2020.106597

65. Patil, I., 2021. statsExpressions: R Package for Tidy Dataframes and Expressions with Statistical Details. Journal of Open Source Software 6, 3236. 10.21105/joss.03236

66. Ptak, S.E., Przeworski, M., 2002. Evidence for population growth in humans is confounded by fine-scale population structure. Trends Genet. 18, 559–563. 10.1016/S0168-9525(02)02781-6

67. Pujolar, J.M., Blom, M.P.K., Reeve, A.H., Kennedy, J.D., Marki, P.Z., Korneliussen, T.S., Freeman, B.G., Sam, K., Linck, E., Haryoko, T., Iova, B., Koane, B., Maiah, G., Paul, L., Irestedt, M., Jønsson, K.A., 2022. The formation of avian montane diversity across barriers and along elevational gradients. Nat. Commun. 13, 268. 10.1038/s41467-021-27858-5

68. R Core Team, 2021. R: A Language and Environment for Statistical Computing. R Foundation for Statistical Computing, Vienna, Austria.

69. Ricklefs, R.E., 1970. Stage of Taxon Cycle and Distribution of Birds on Jamaica, Greater Antilles. Evolution 24, 475–477. 10.2307/2406820

70. Ricklefs, R.E., Cox, G.W., 1972. Taxon Cycles in the West Indian Avifauna. Am. Nat. 106, 195–219. 10.1086/282762

71. Ricklefs, R.E., Cox, G.W., 1978. Stage of Taxon Cycle, Habitat Distribution, and Population Density in the Avifauna of the West Indies. Am. Nat. 112, 875–895. 10.1086/283329

72. Ripple, W.J., Wolf, C., Newsome, T.M., Hoffmann, M., Wirsing, A.J., McCauley, D.J., 2017. Extinction risk is most acute for the world’s largest and smallest vertebrates. Proc. Natl. Acad. Sci. 114, 10678–10683. 10.1073/pnas.1702078114

73. Şekercioğlu, Ç.H., Primack, R.B., Wormworth, J., 2012. The effects of climate change on tropical birds. Biol. Conserv. 148, 1–18. 10.1016/j.biocon.2011.10.019

74. Smith, B.T., Gehara, M., Harvey, M.G., 2021. The demography of extinction in eastern North American birds. Proc. R. Soc. B Biol. Sci. 288, 20201945. 10.1098/rspb.2020.1945

75. Sodhi, N.S., Koh, L.P., Brook, B.W., Ng, P.K.L., 2004. Southeast Asian biodiversity: an impending disaster. Trends Ecol. Evol. 19, 654–660. 10.1016/j.tree.2004.09.006

76. Spielman, D., Brook, B.W., Frankham, R., 2004. Most species are not driven to extinction before genetic factors impact them. Proc. Natl. Acad. Sci. 101, 15261–15264. 10.1073/pnas.0403809101

77. Shrestha, N., 2020. Detecting Multicollinearity in Regression Analysis. American Journal of Applied Mathematics and Statistics 8, 39–42. 10.12691/ajams-8-2-1

78. Steadman, D.W., Martin, P.S., 2003. The late Quaternary extinction and future resurrection of birds on Pacific islands. Earth-Sci. Rev. 61, 133–147. 10.1016/S0012-8252(02)00116-2

79. Steadman, D.W., Pregill, G.K., Olson, S.L., 1984. Fossil vertebrates from Antigua, Lesser Antilles: Evidence for late Holocene human-caused extinctions in the West Indies. Proc. Natl. Acad. Sci. 81, 4448–4451. 10.1073/pnas.81.14.4448

80. Tan, H.Z., Jansen, J.J., Allport, G.A., Garg, K.M., Chattopadhyay, B., Irestedt, M., Pang, S.E., Chilton, G., Gwee, C.Y., Rheindt, F.E., 2023. Megafaunal extinctions, not climate change, may explain Holocene genetic diversity declines in Numenius shorebirds. eLife 12, e85422. 10.7554/eLife.85422

81. Voris, H.K., 2000. Maps of Pleistocene sea levels in Southeast Asia: shorelines, river systems and time durations. J. Biogeogr. 27, 1153–1167. 10.1046/j.1365-2699.2000.00489.x

82. Wang, J., Santiago, E., Caballero, A., 2016. Prediction and estimation of effective population size. Heredity 117, 193–206. 10.1038/hdy.2016.43

83. Wickham, H., RStudio, 2023. tidyverse: Easily Install and Load the “Tidyverse.”

84. Wilder, A.P., Supple, M.A., Subramanian, A., Mudide, A., Swofford, R., Serres-Armero, A., Steiner, C., Koepfli, K.-P., Genereux, D.P., Karlsson, E.K., Lindblad-Toh, K., Marques-Bonet, T., Munoz Fuentes, V., Foley, K., Meyer, W.K., Zoonomia Consortium, Ryder, O.A., Shapiro, B., 2023. The contribution of historical processes to contemporary extinction risk in placental mammals. Science 380, eabn5856. 10.1126/science.abn5856

85. Willis, K.J., Bennett, K.D., Walker, D., Hewitt, G.M., 2004. Genetic consequences of climatic oscillations in the Quaternary. Philos. Trans. R. Soc. Lond. B. Biol. Sci. 359, 183–195. 10.1098/rstb.2003.1388

86. Zizka, A., Silvestro, D., Andermann, T., Azevedo, J., Ritter, C.D., Edler, D., Farooq, H., Herdean, A., Ariza, M., Scharn, R., Svanteson, S., Wengstrom, N., Zizka, V., Antonelli, A., sp, B.V. (Bruno updated the package to remove dependencies on, raster, rgdal, maptools, packages), and rgeos, ropensci, I.S. (Irene reviewed the package for, see <https://github.com/ropensci/onboarding/issues/210>), ropensci, F.R.-S. (Francisco reviewed the package for, see <https://github.com/ropensci/onboarding/issues/210>), 2023. CoordinateCleaner: Automated Cleaning of Occurrence Records from Biological Collections.

