## Supplementary figures for "Species biology and demographic history determine species vulnerability to climate change in tropical island endemic birds"

### **SUPPLEMENTARY INFORMATION**

**SUPPLEMENTARY FIGURES**

### Figure S1: PSMC plots with bootstrapped replicates of all species included in this study. The plots for all three settings of atomic time intervals used in the current study are provided. For each species, the top panel: The first population-size parameter spans the first four time intervals. The next 30 parameters span two intervals each, and the last three parameters span four, six and 10 intervals respectively (–p “4 + 30 * 2 + 4 + 6 + 10”); middle panel: –p “2 + 2 + 30 * 2 + 4 + 6 + 10”; bottom panel: –p “1 + 1 + 1 + 1 + 30 * 2 + 4 + 6 + 10”.

#### *Actenoides hombroni*

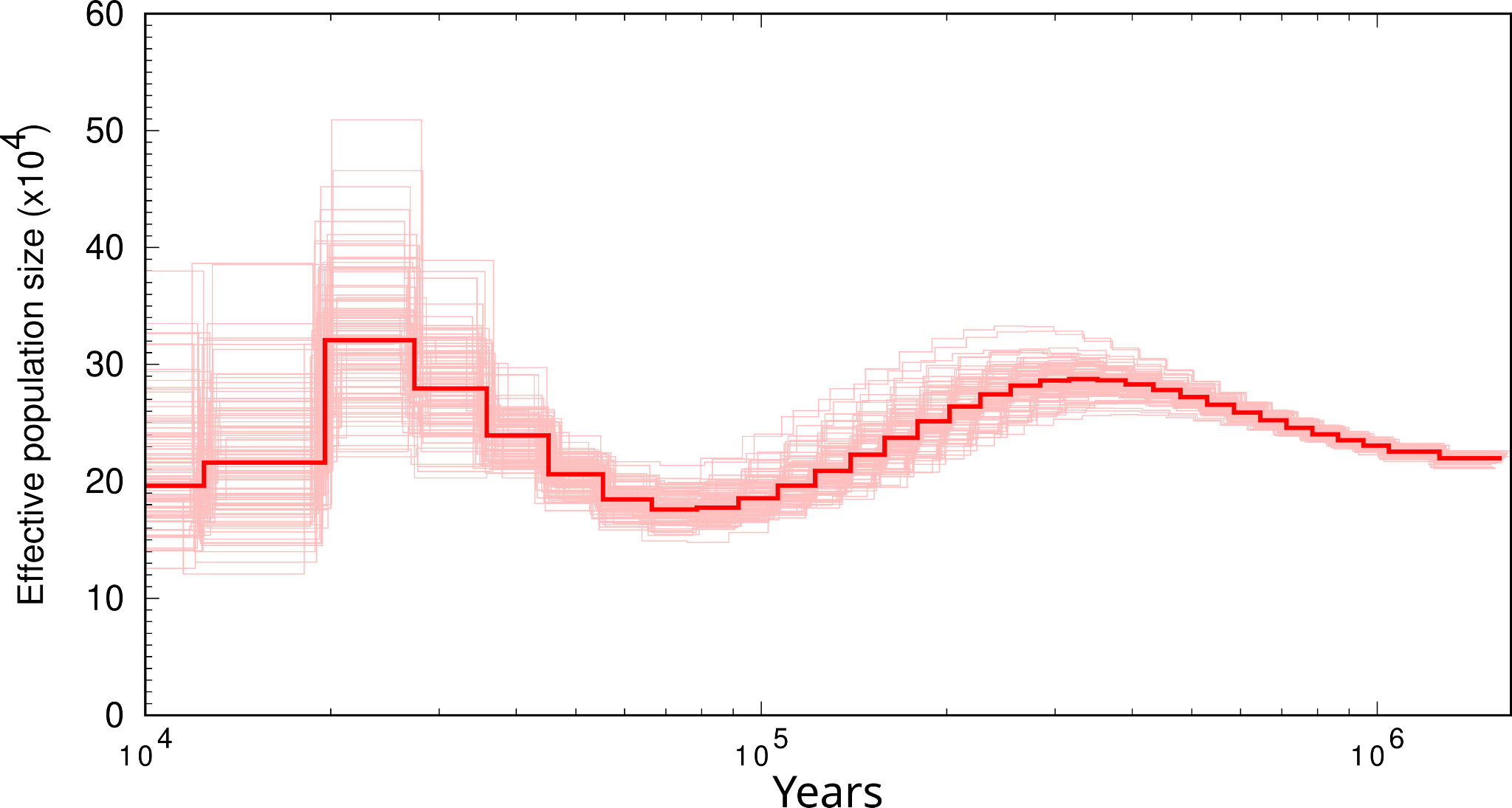

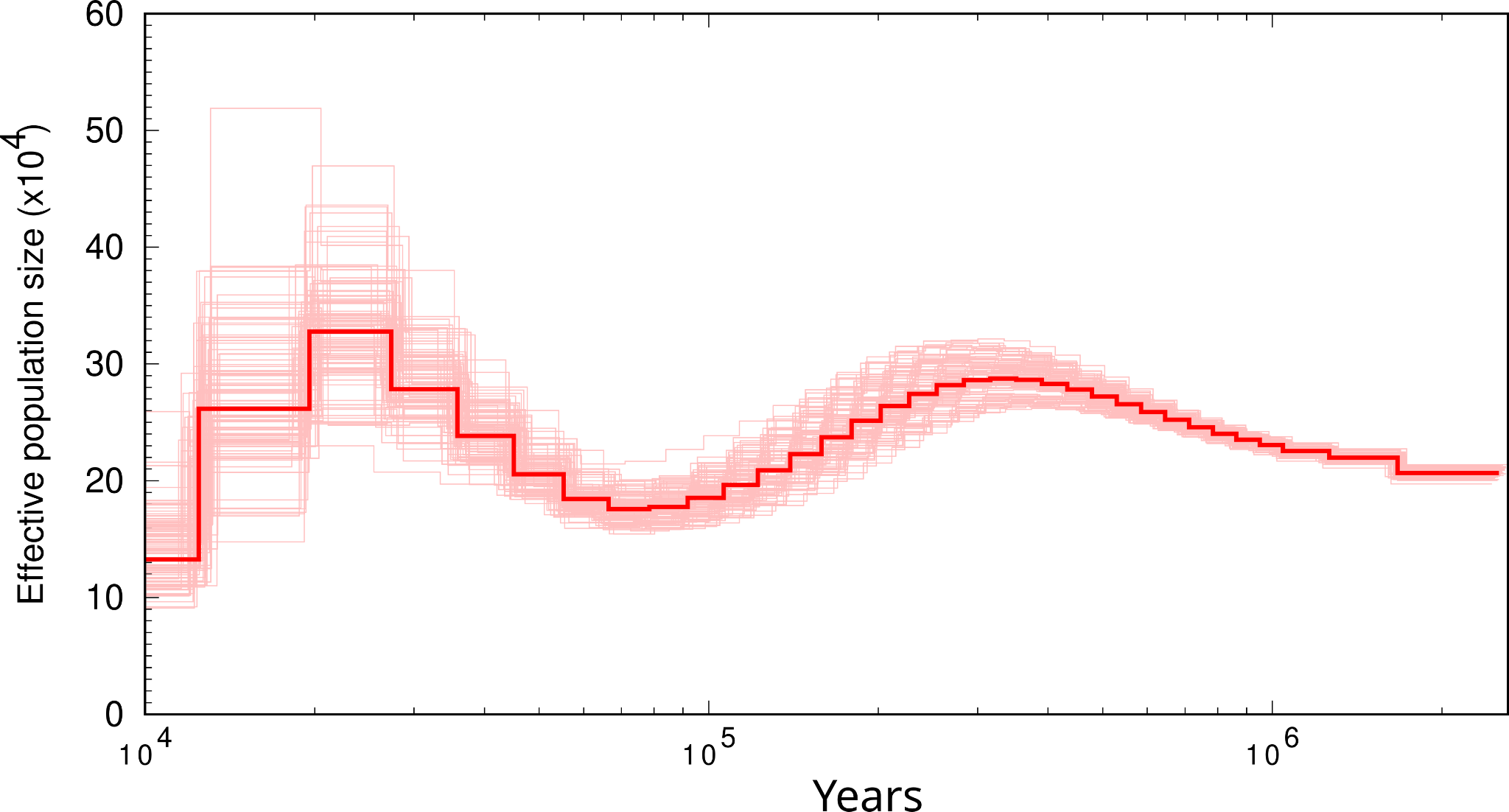

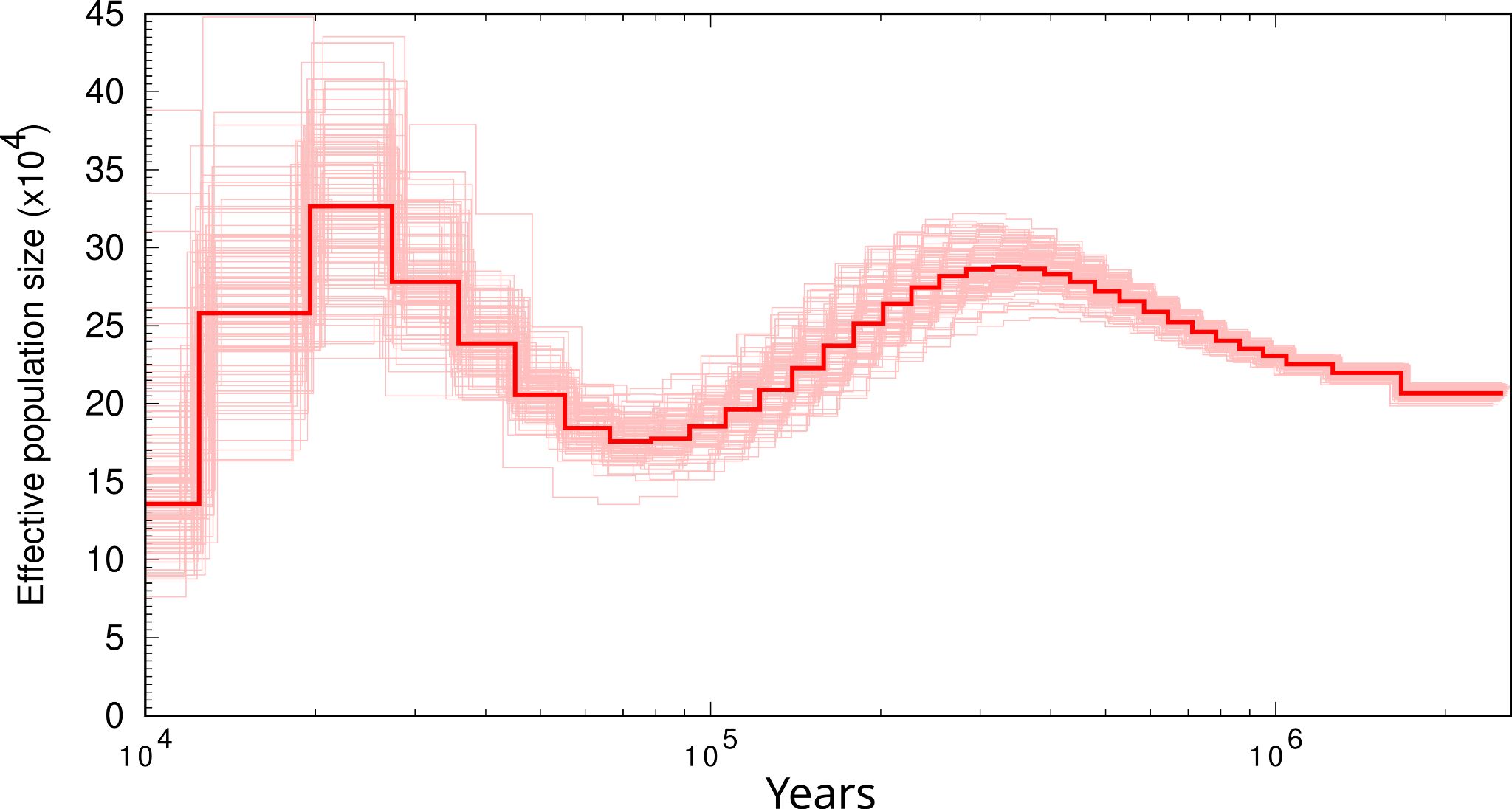

#### *Aleadryas rufinucha*

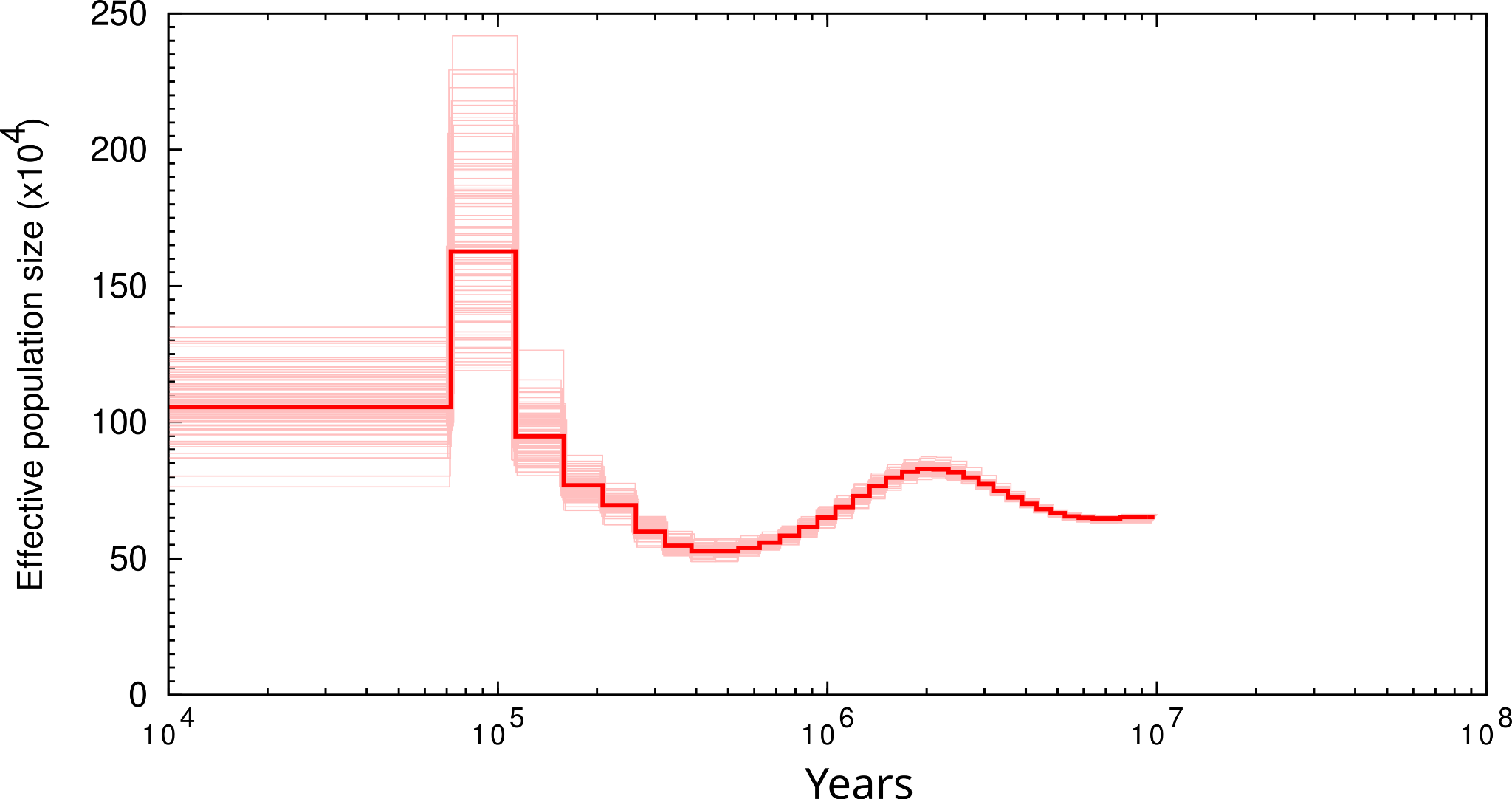

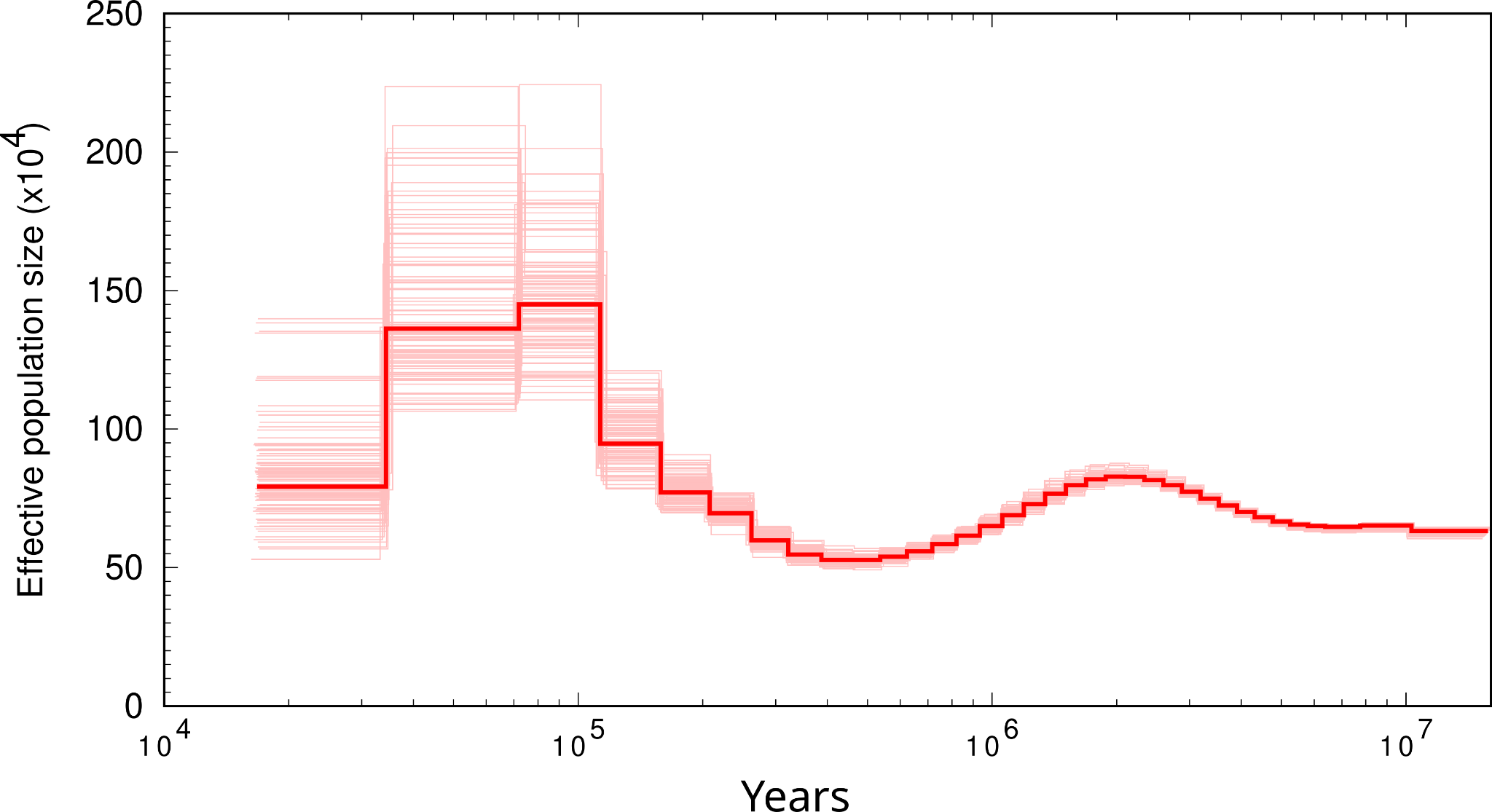

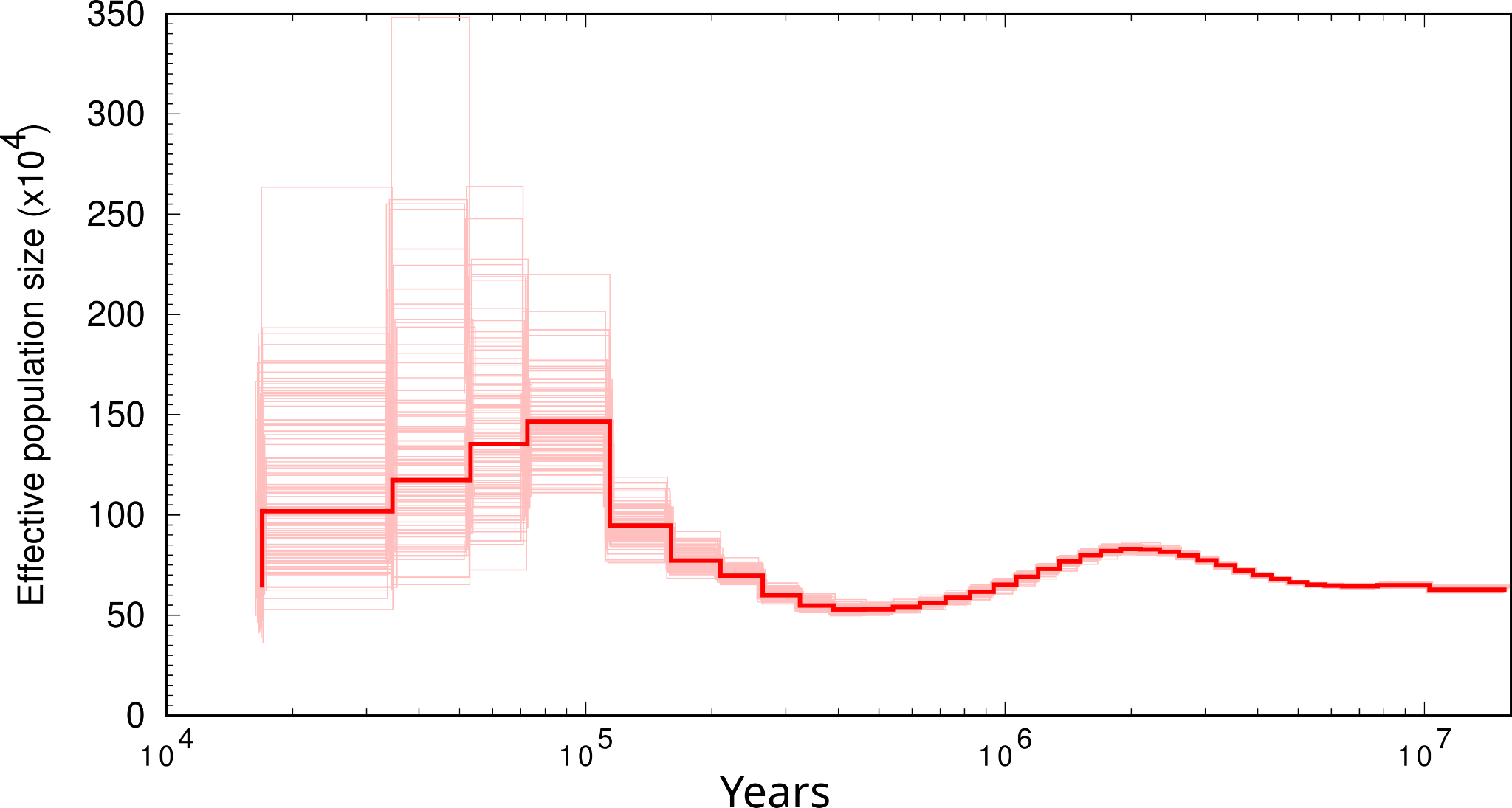

#### *Alectura lathami*

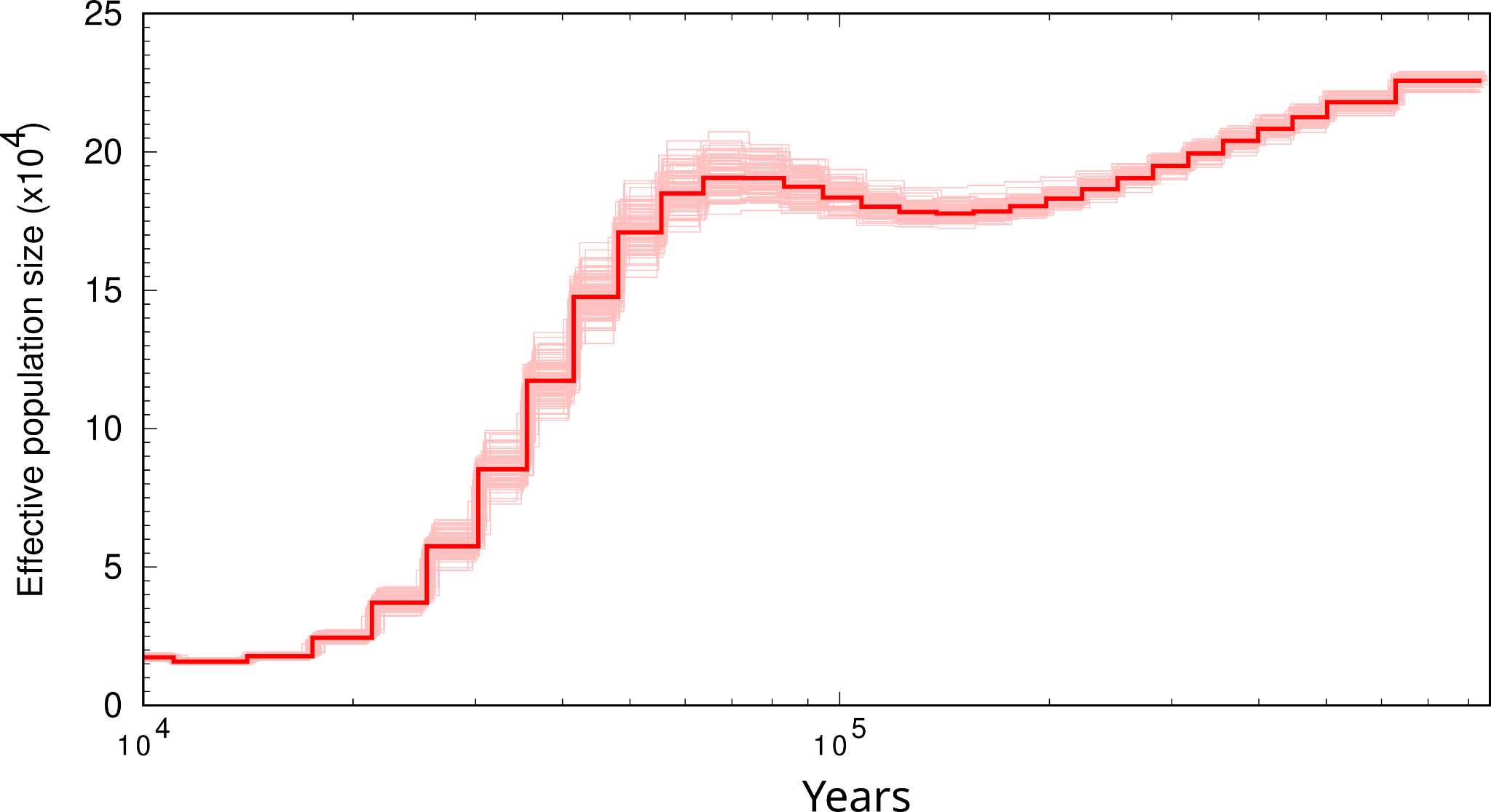

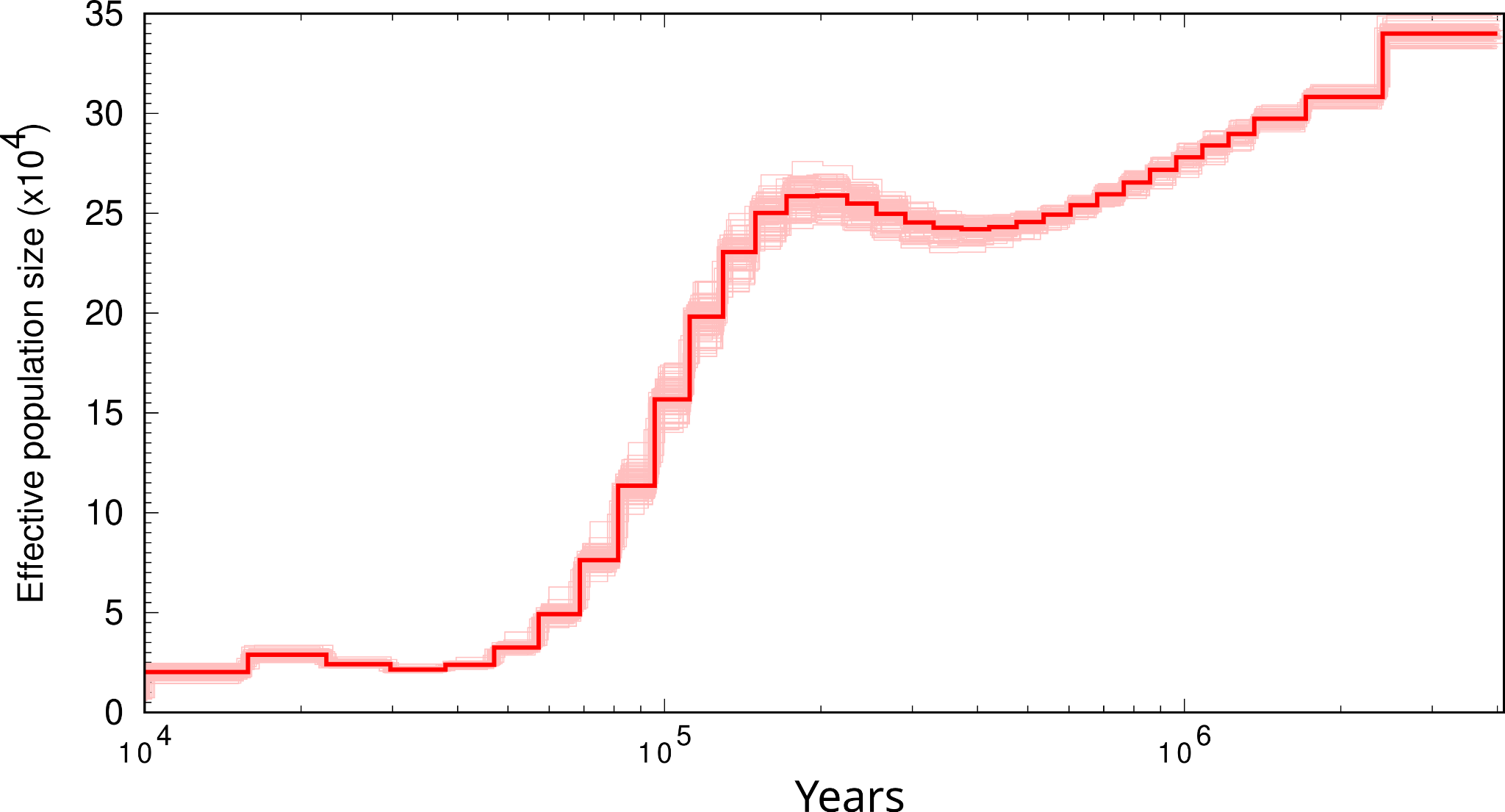

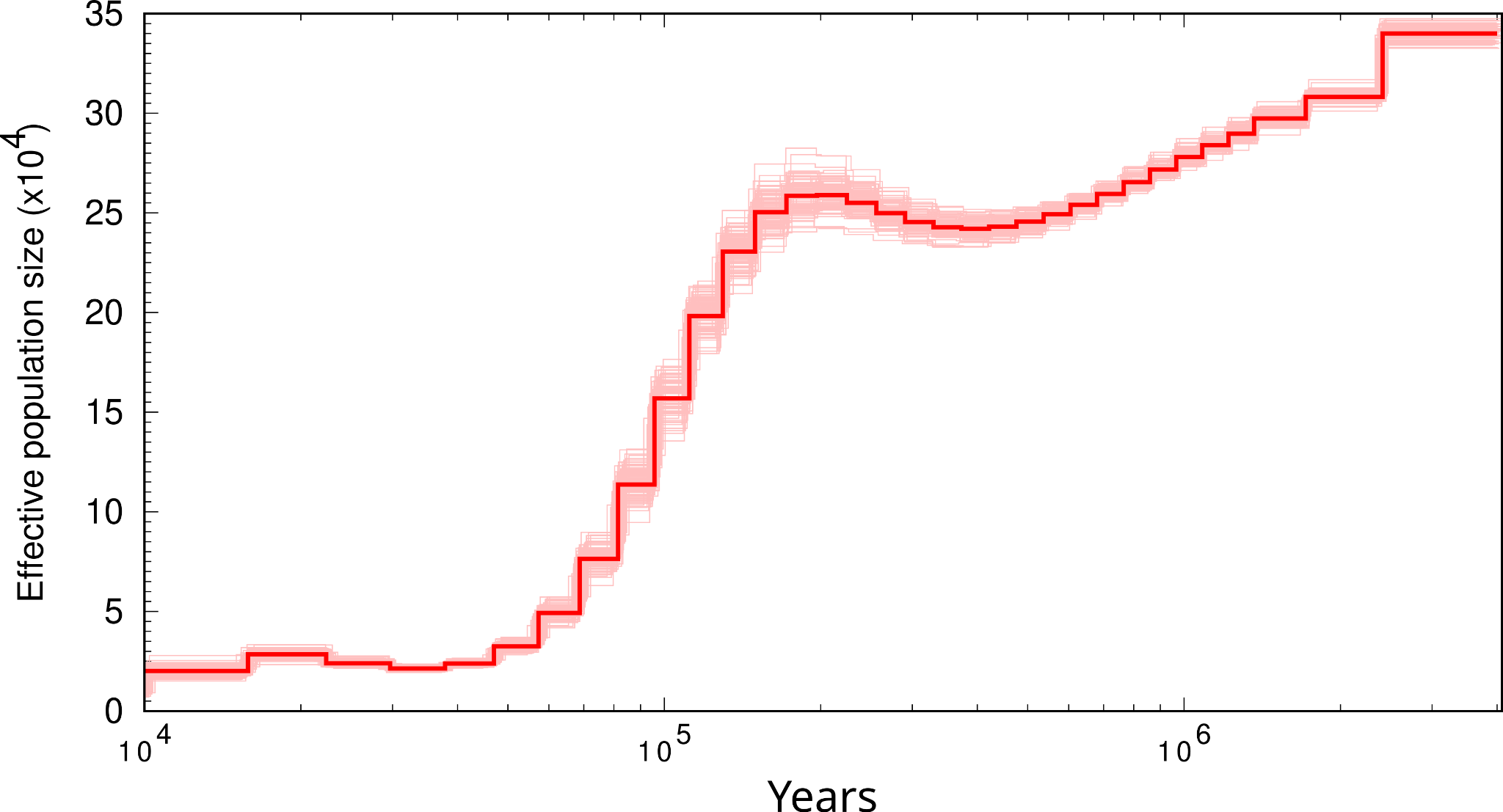

#### *Amazona guildingii*

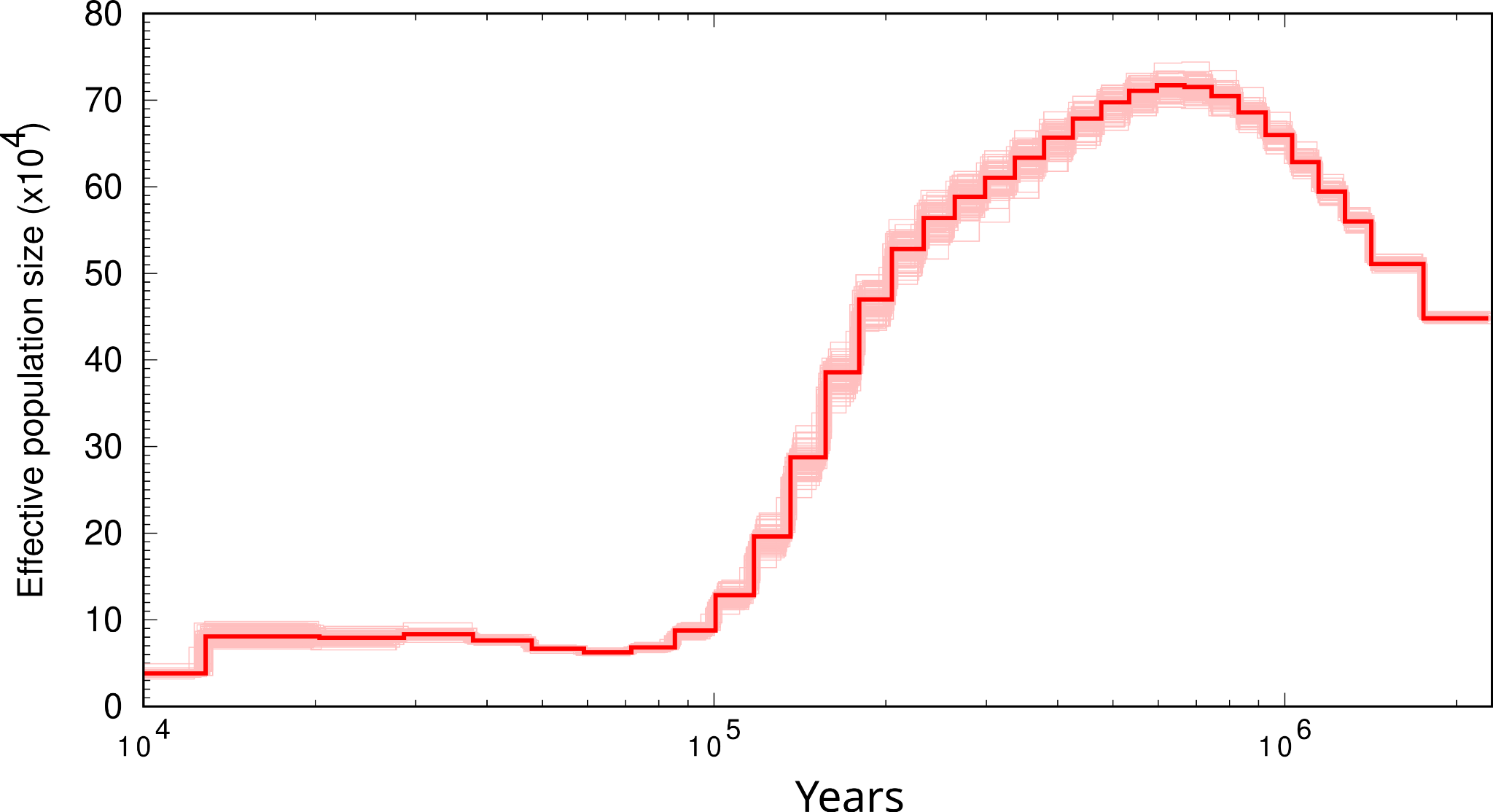

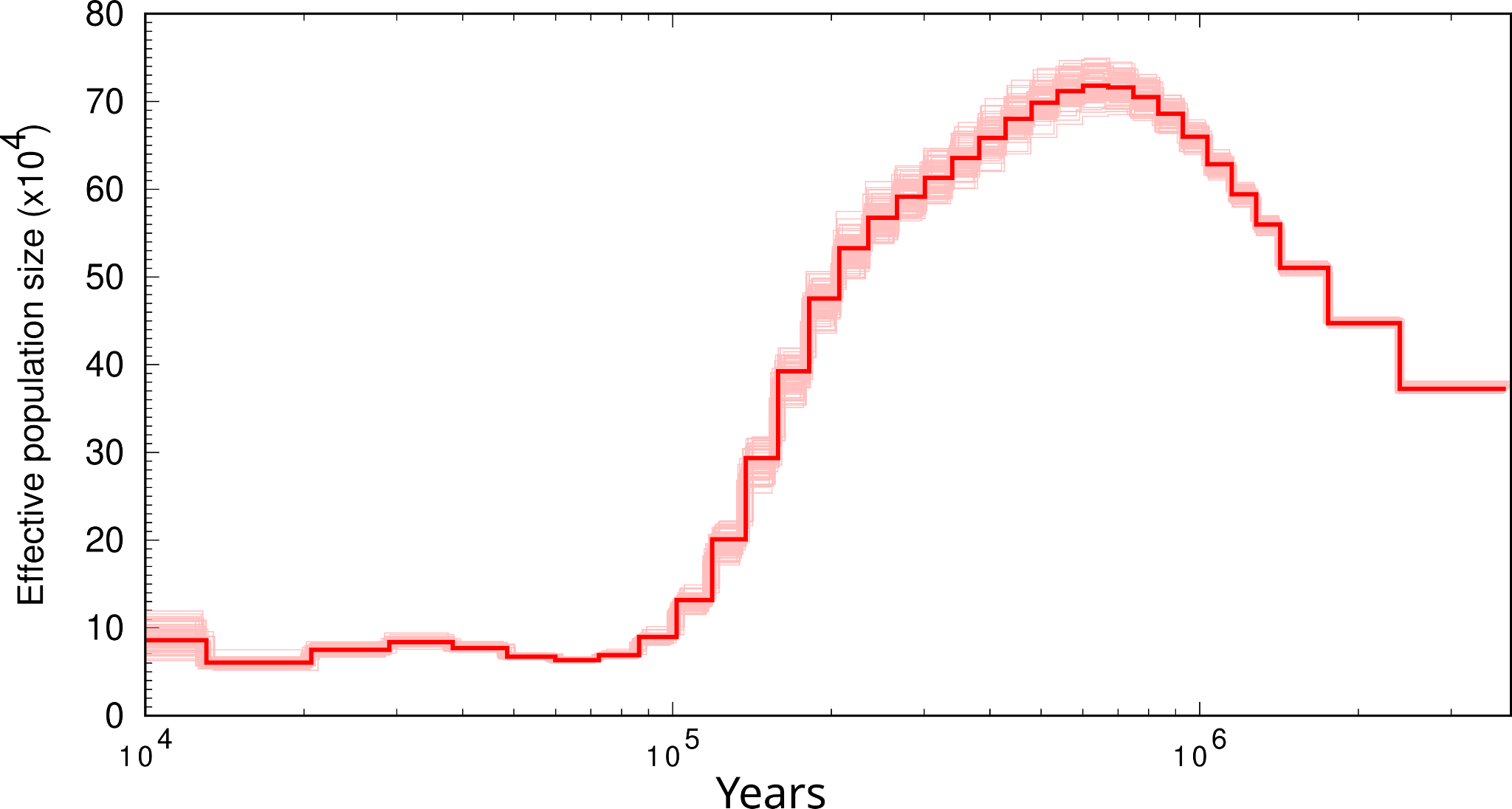

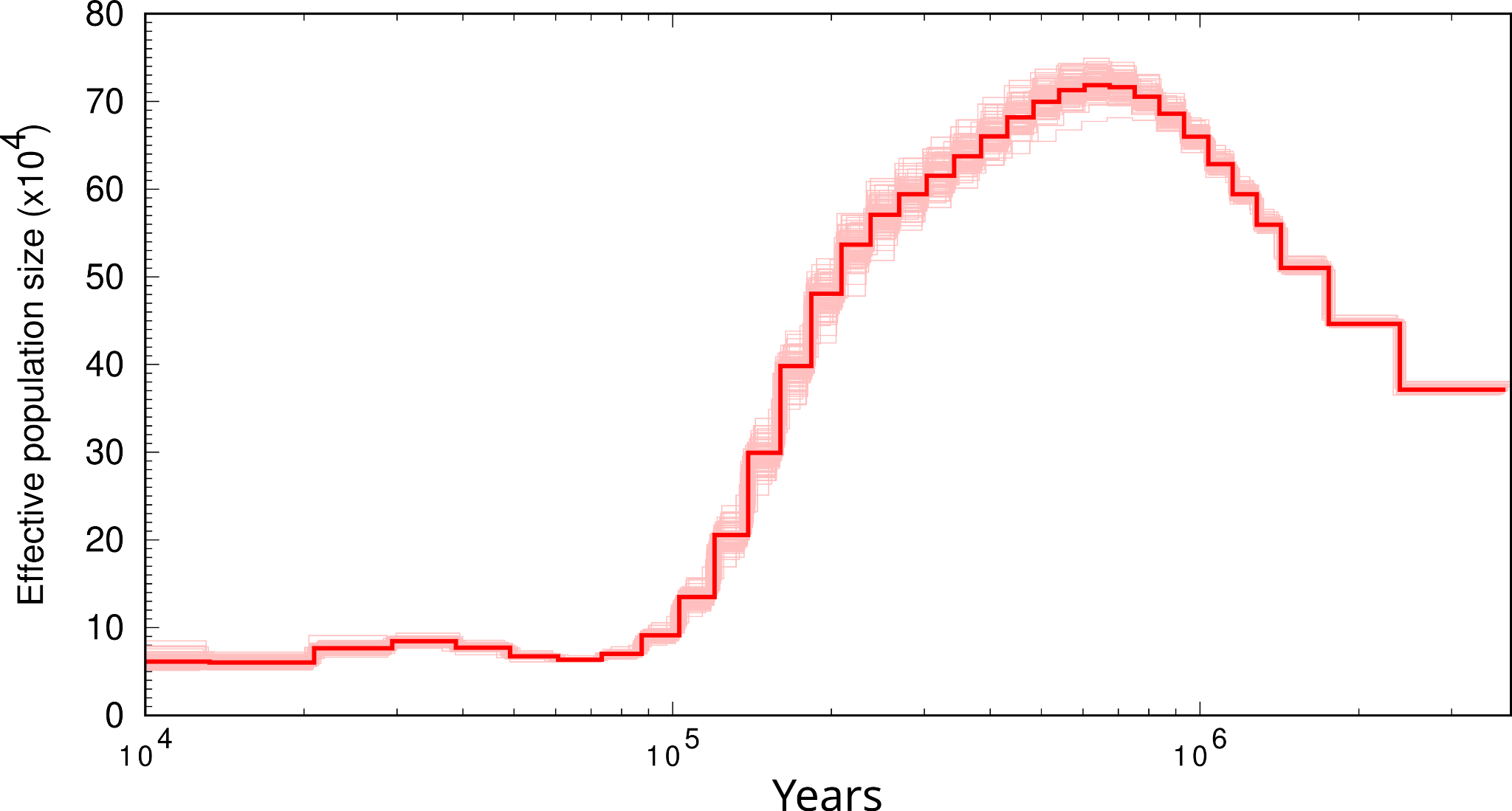

#### *Amazona vittata*

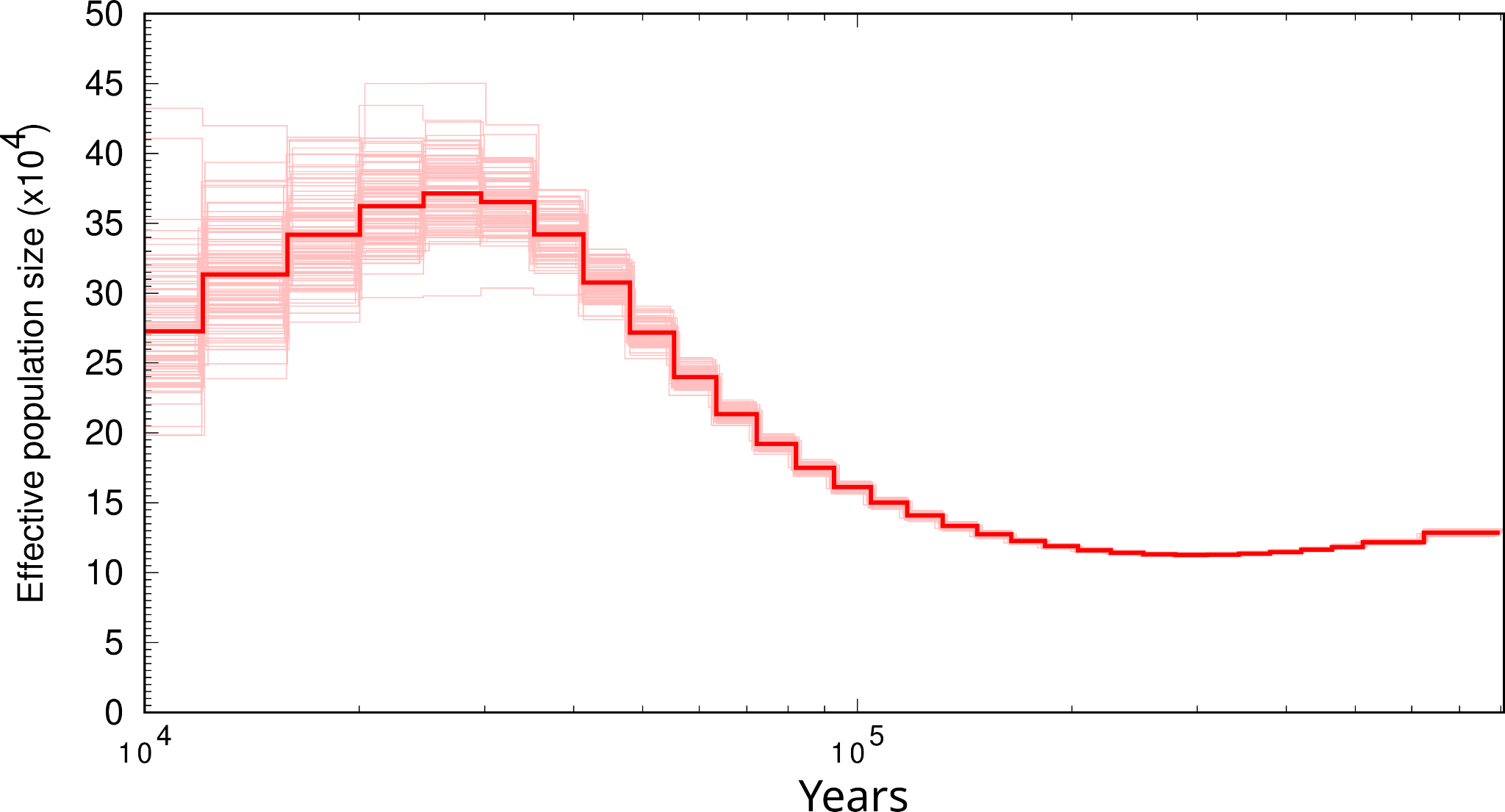

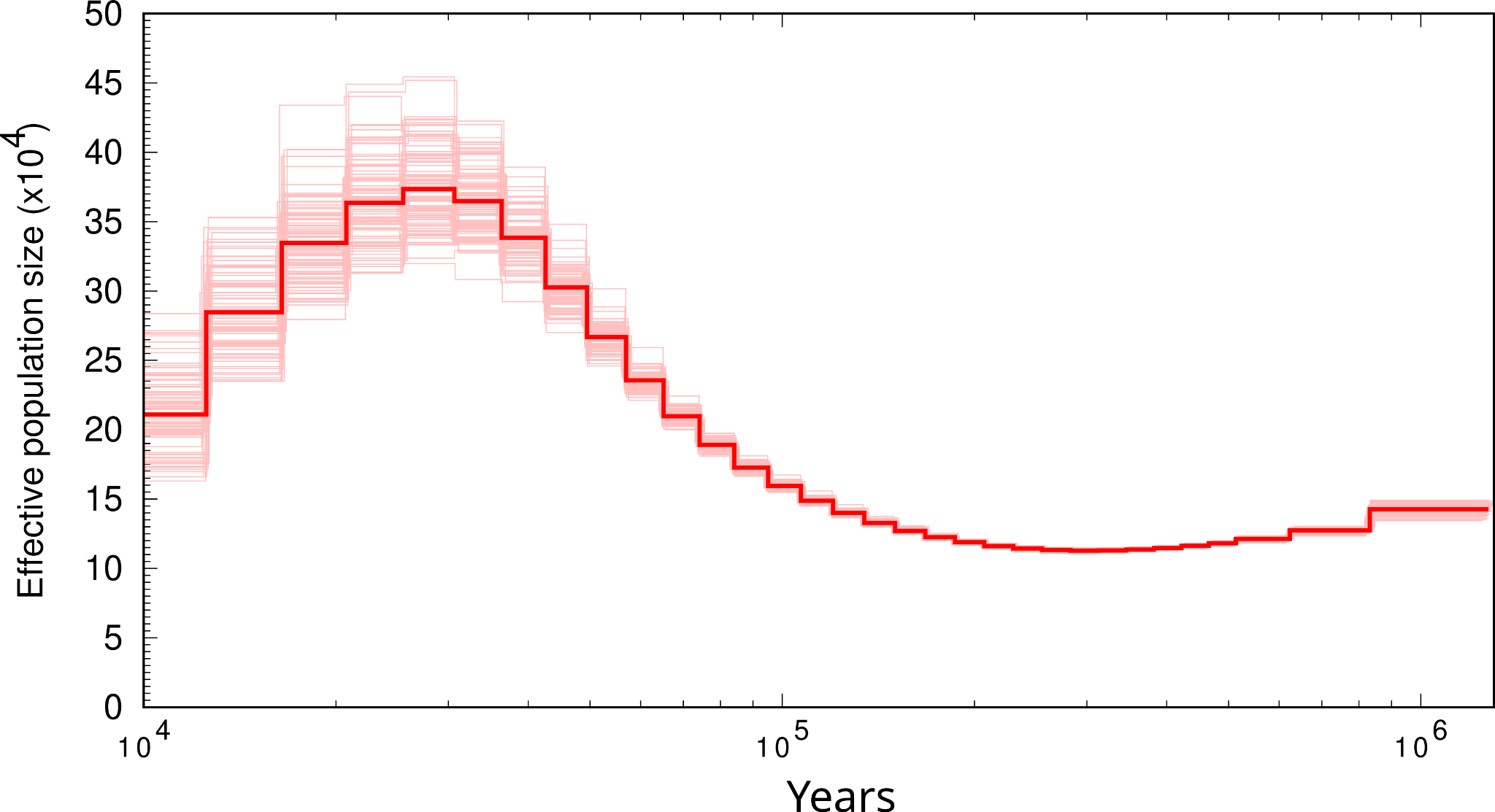

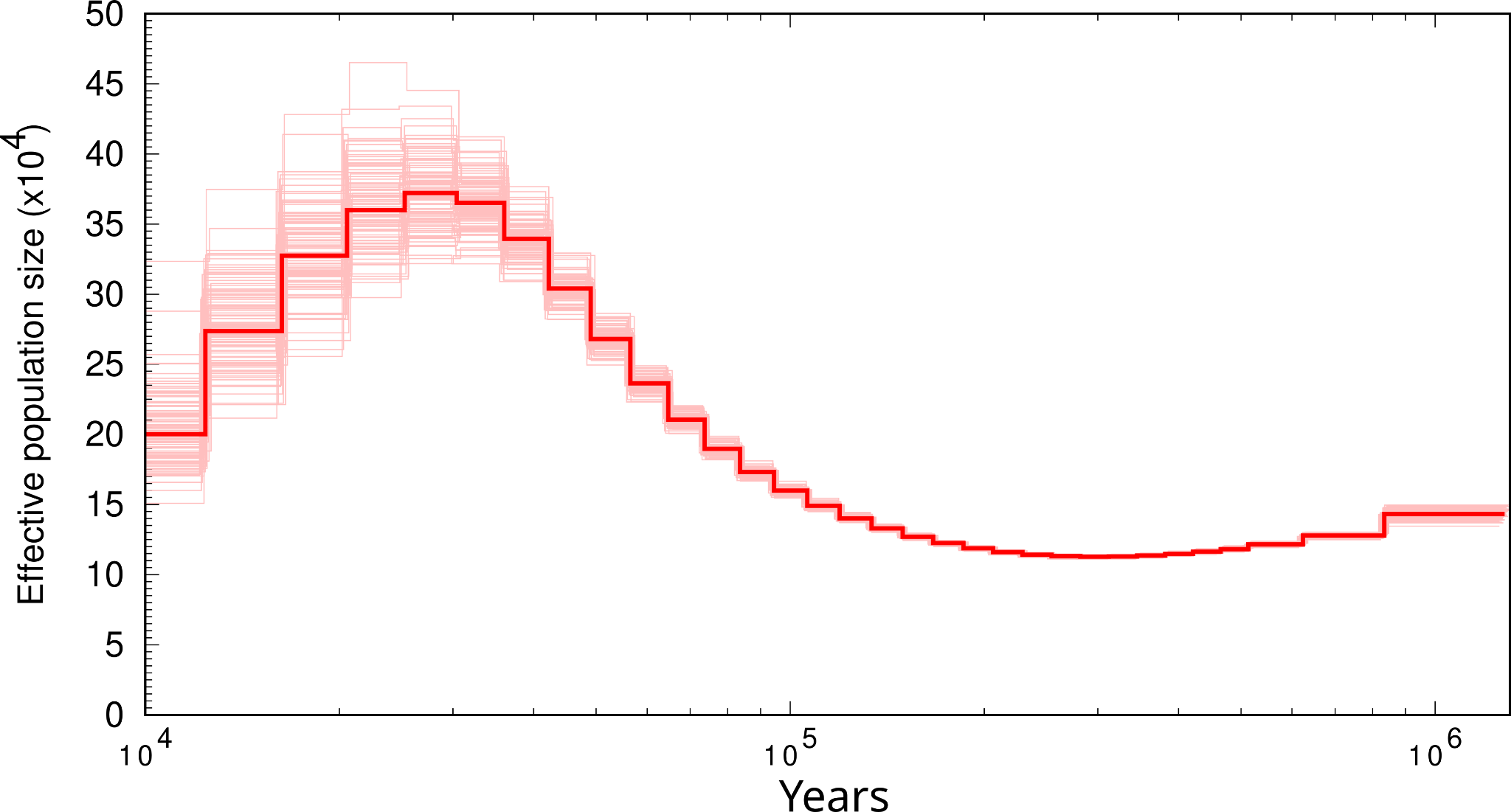

#### *Amblyornis subalaris*

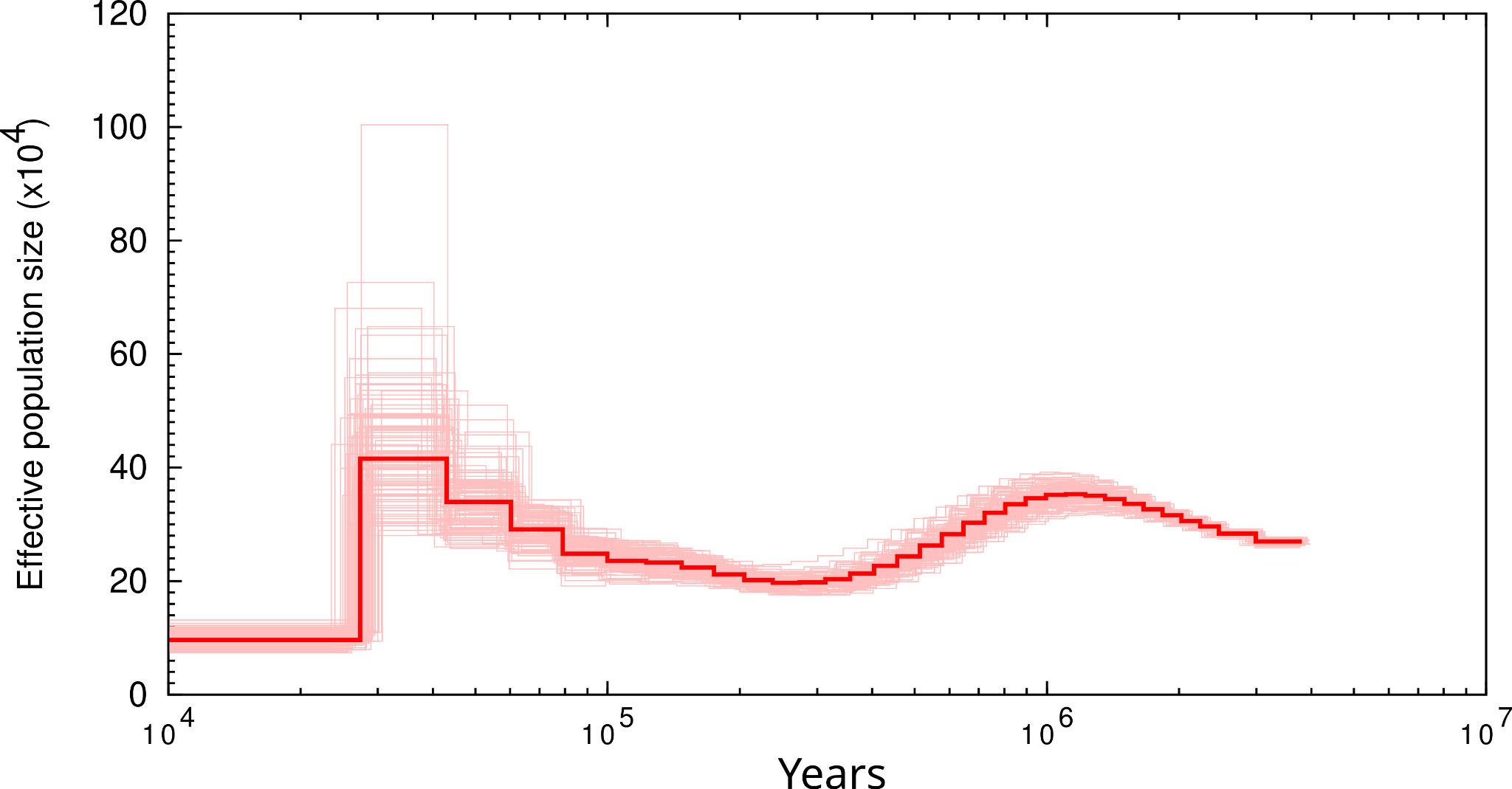

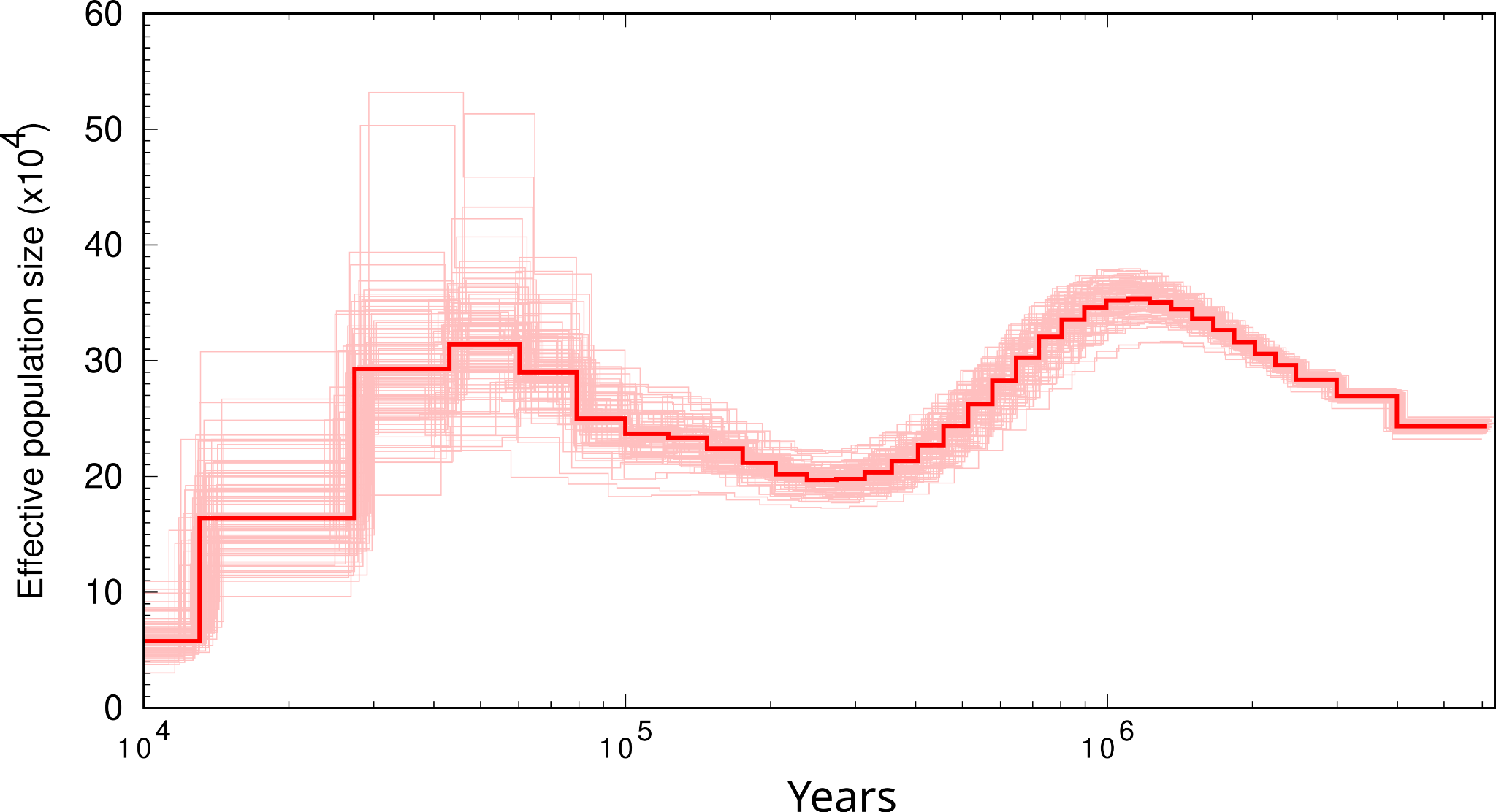

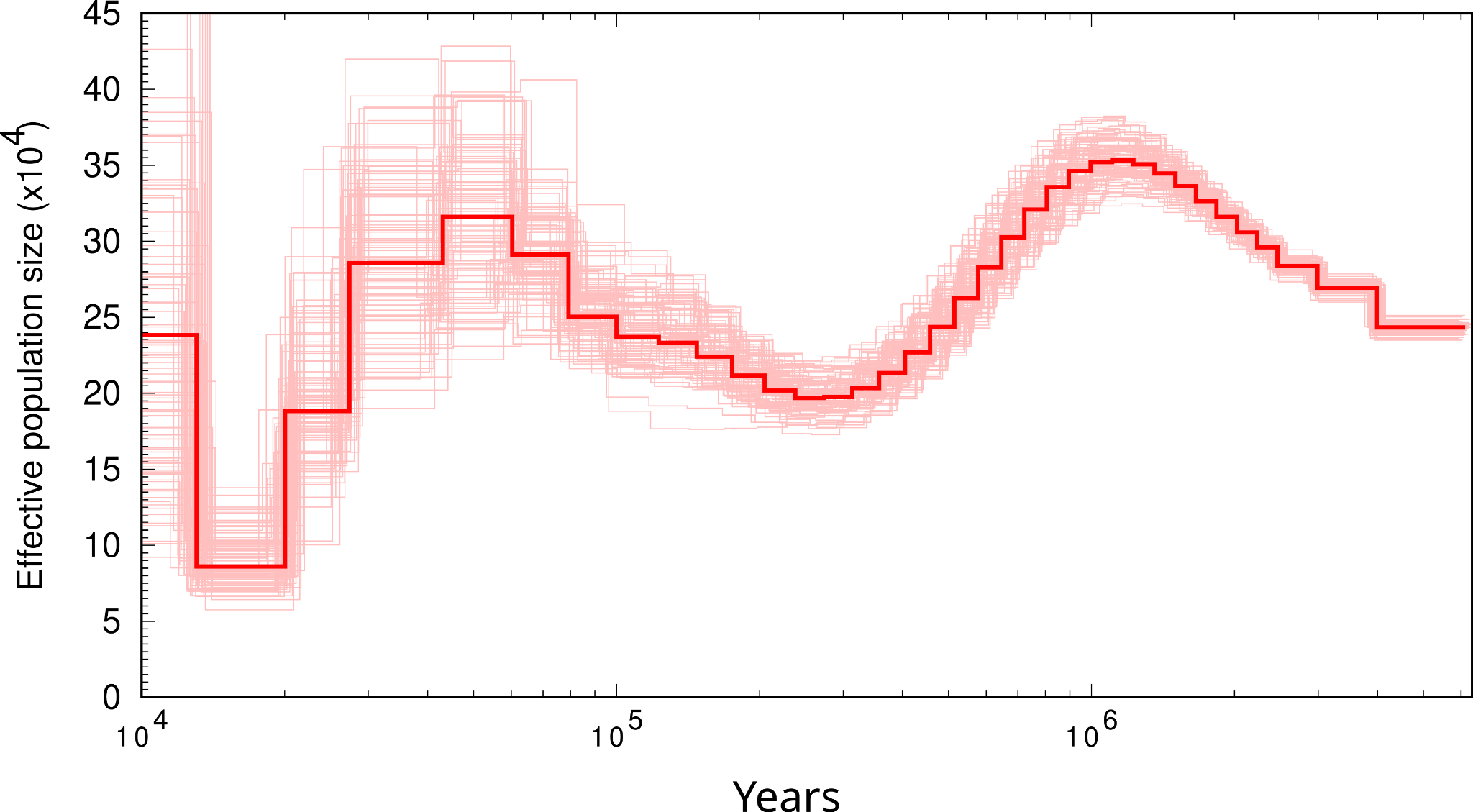

### *Cacatua alba*

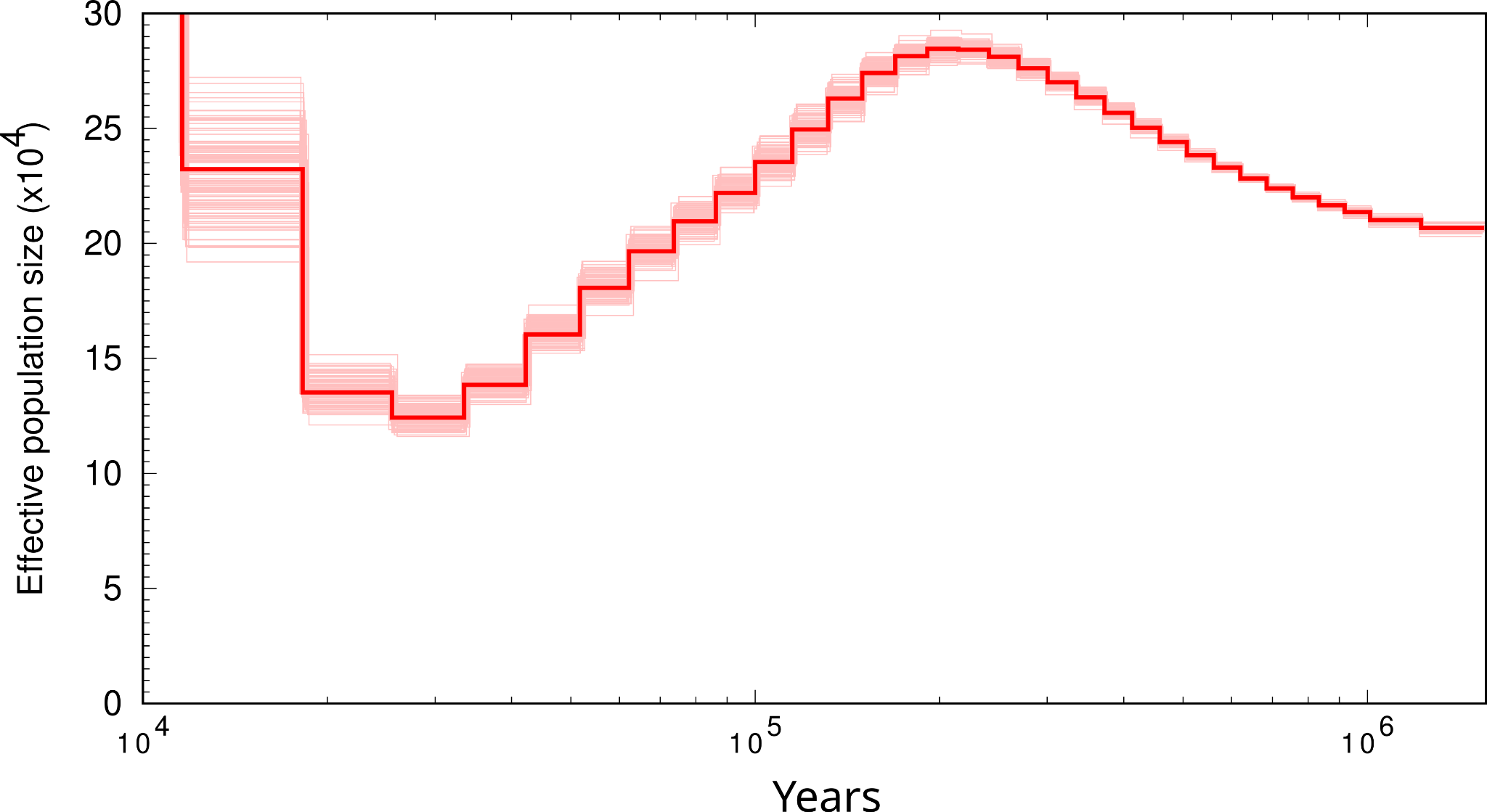

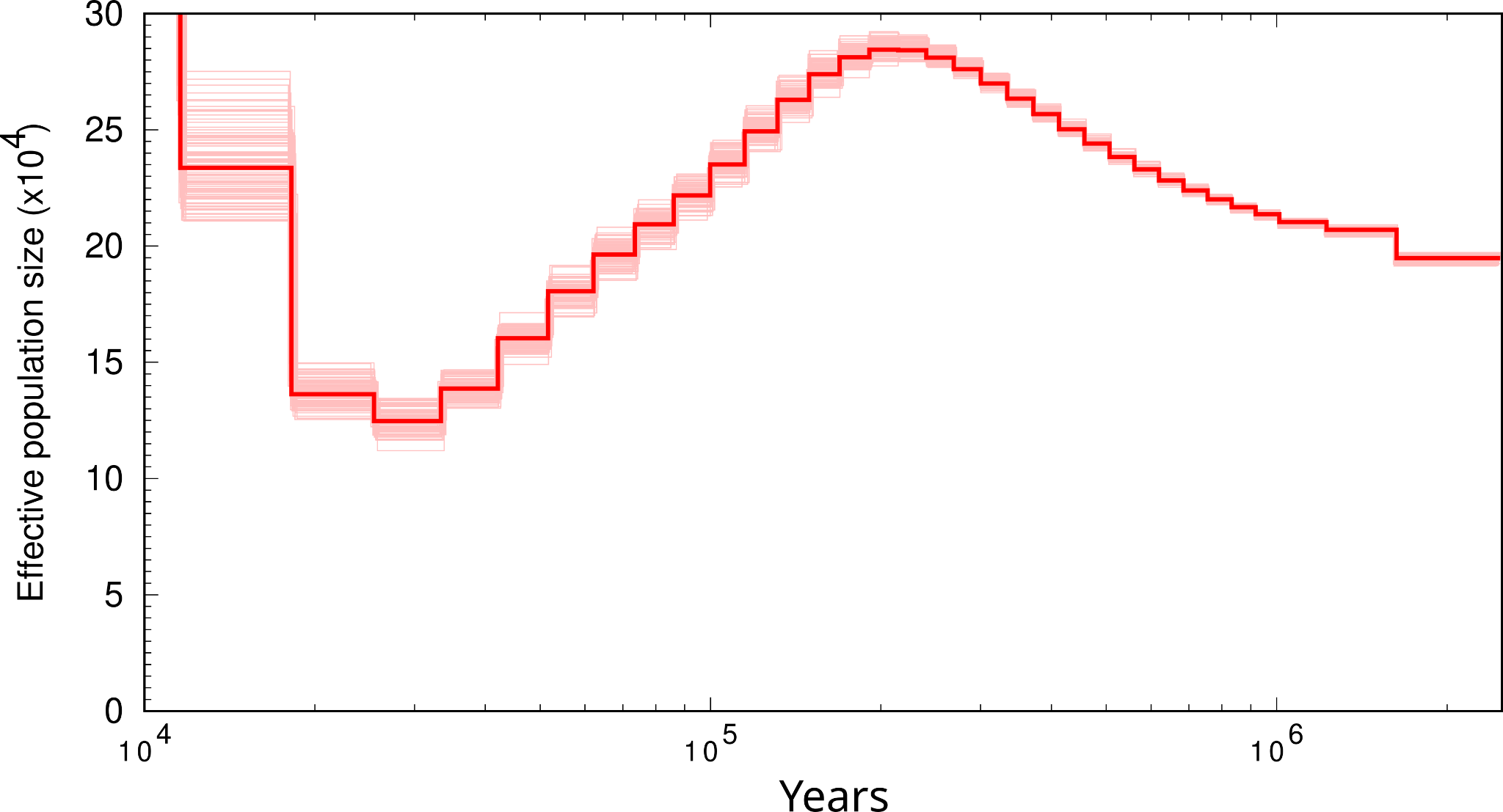

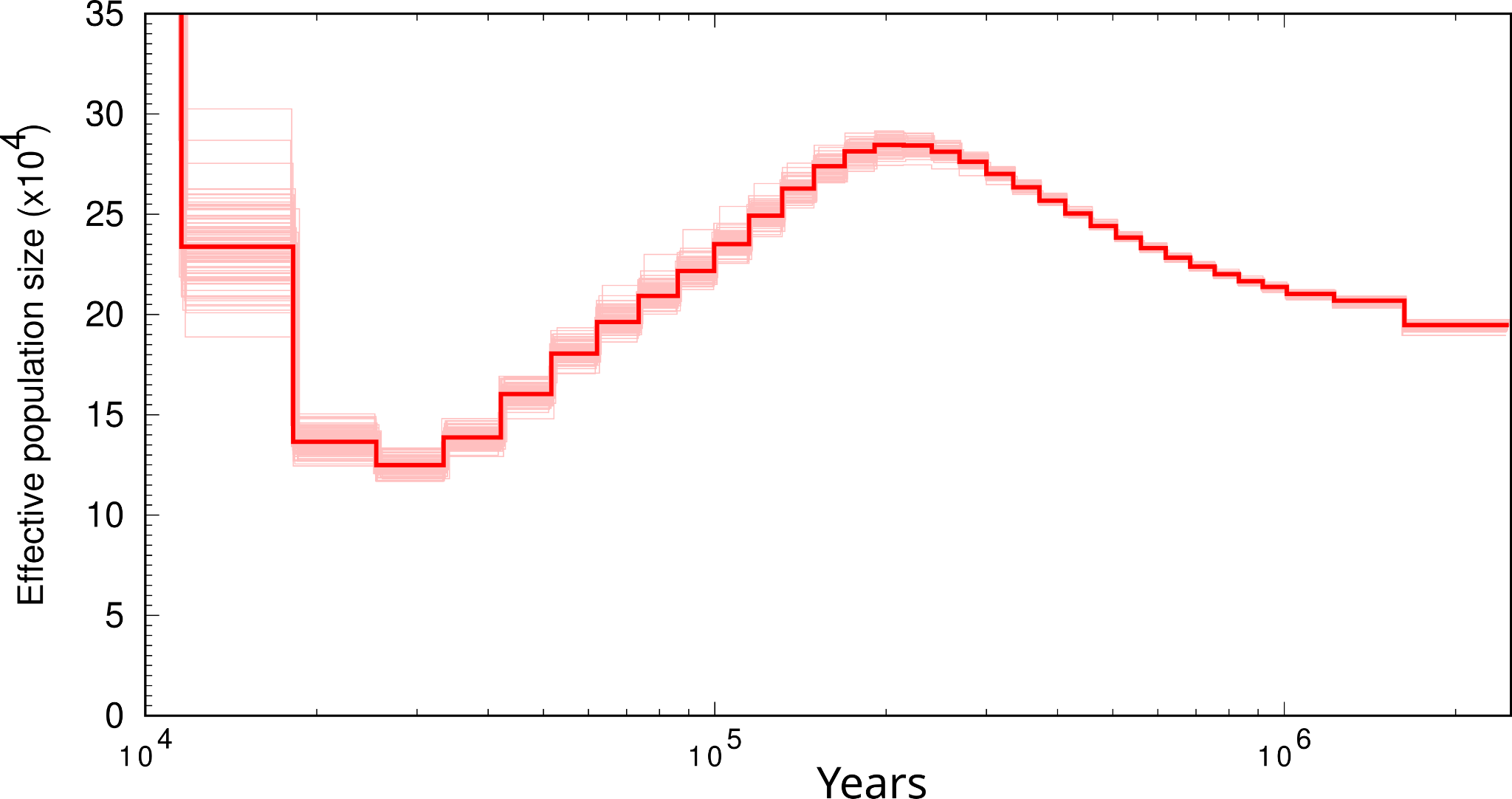

#### *Centropus unirufus*

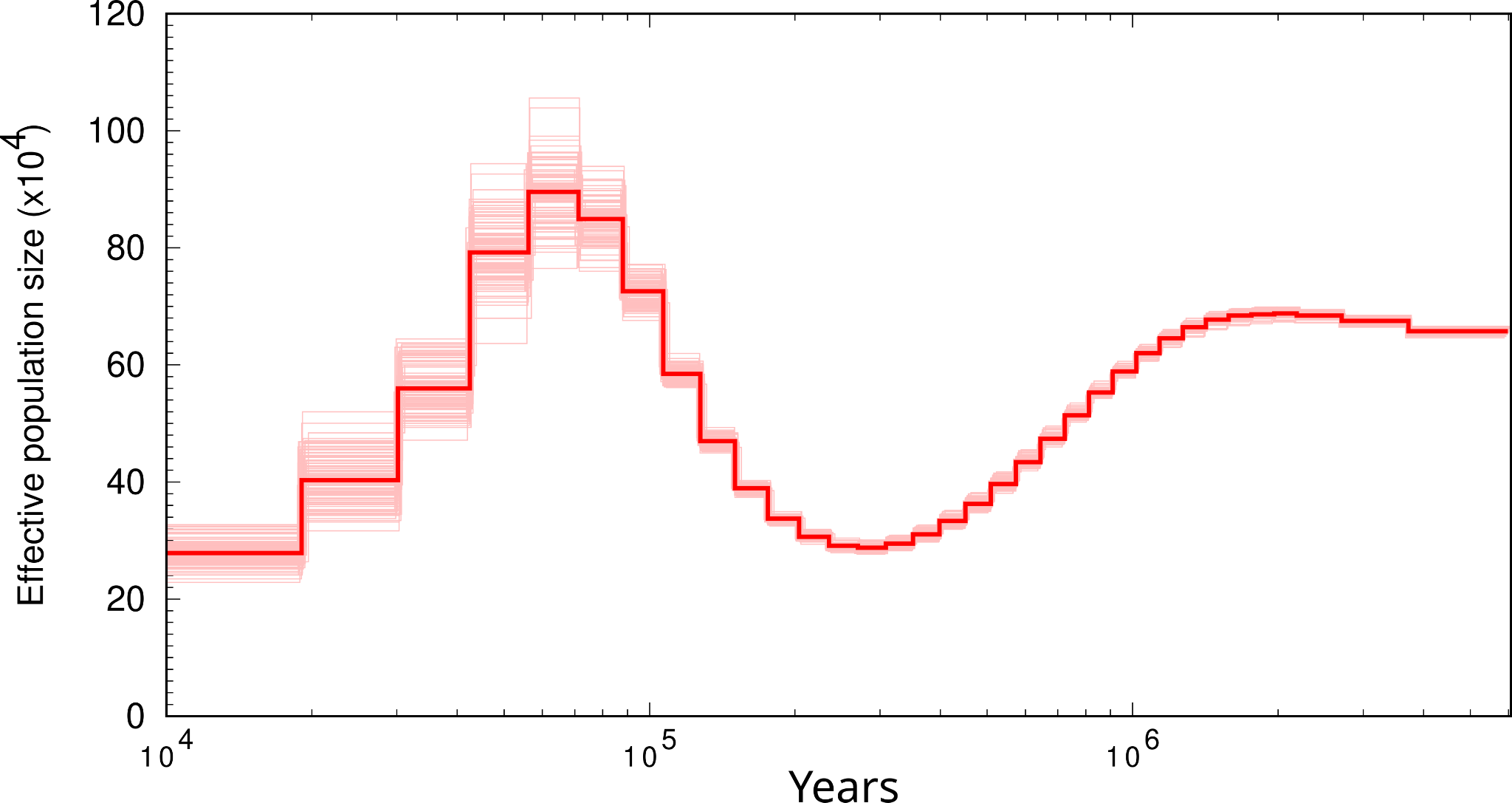

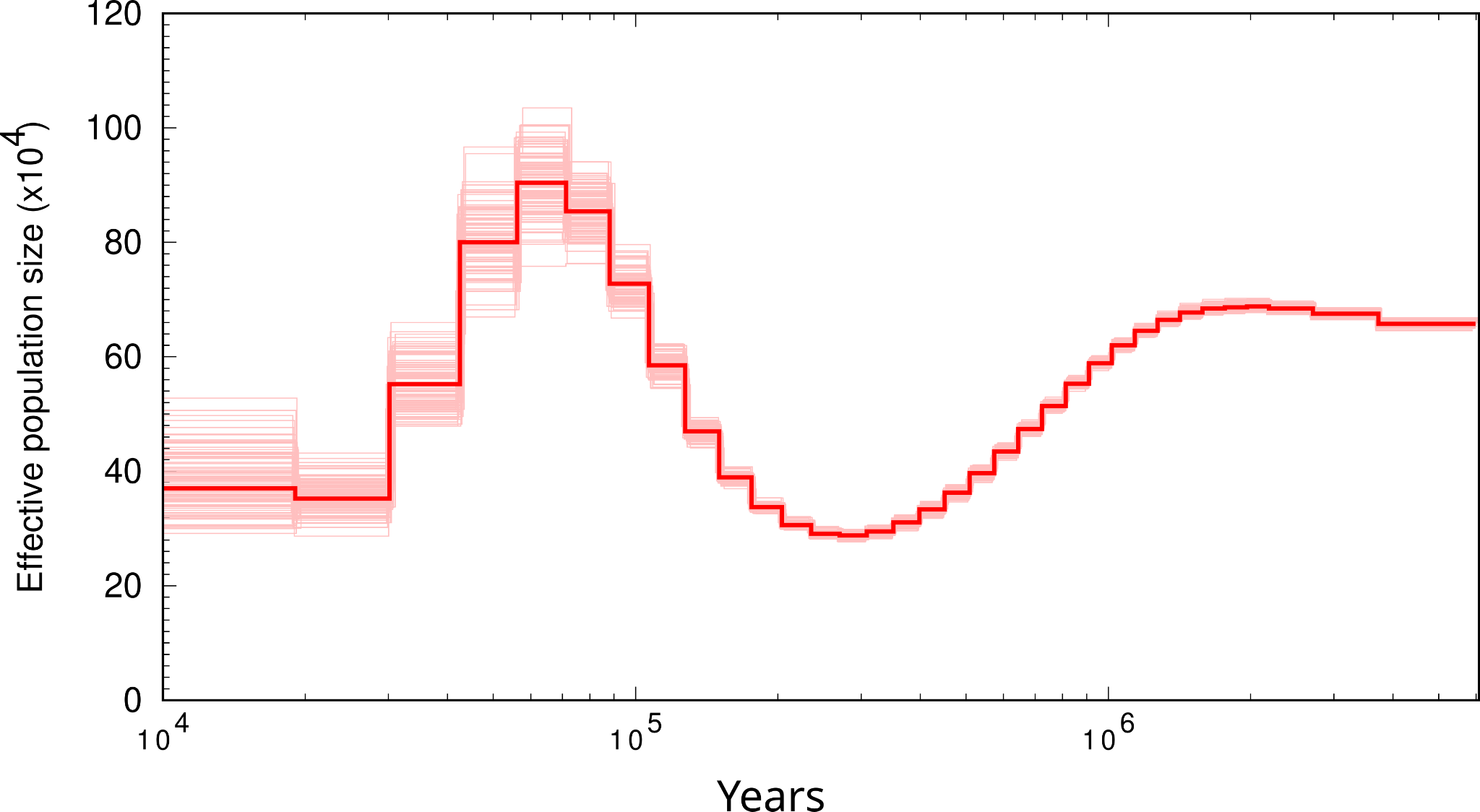

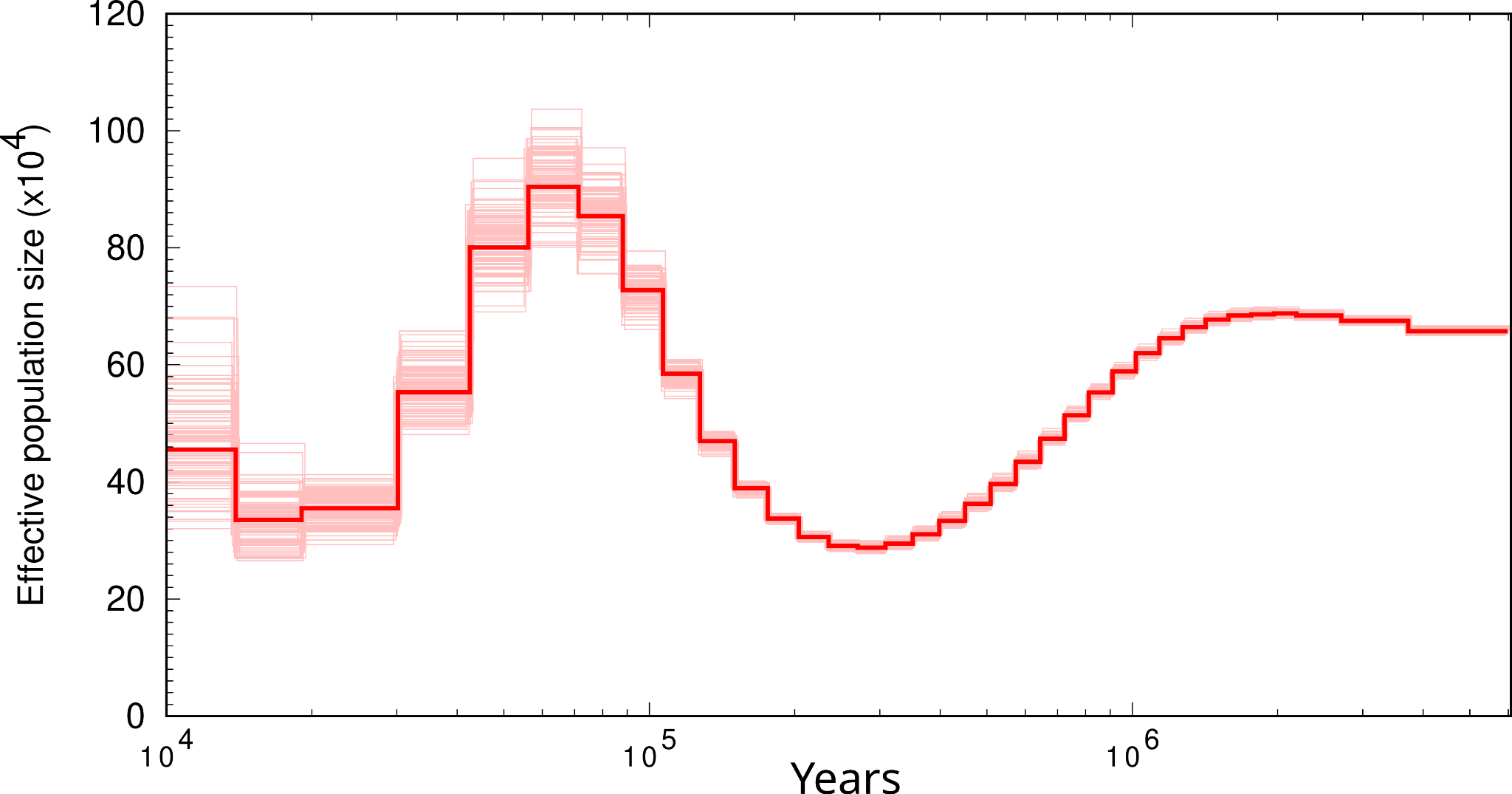

#### *Cicinnurus regius*

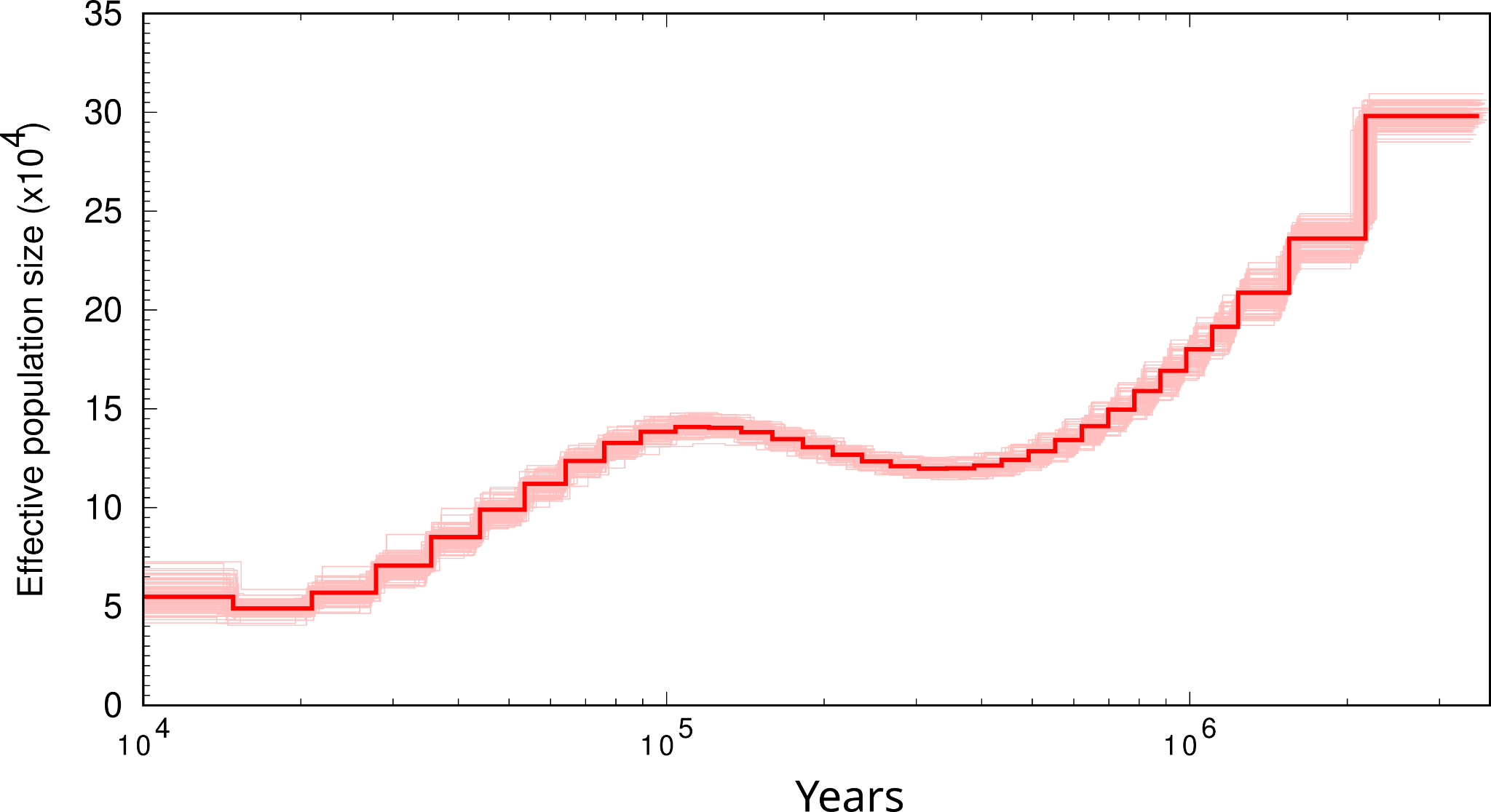

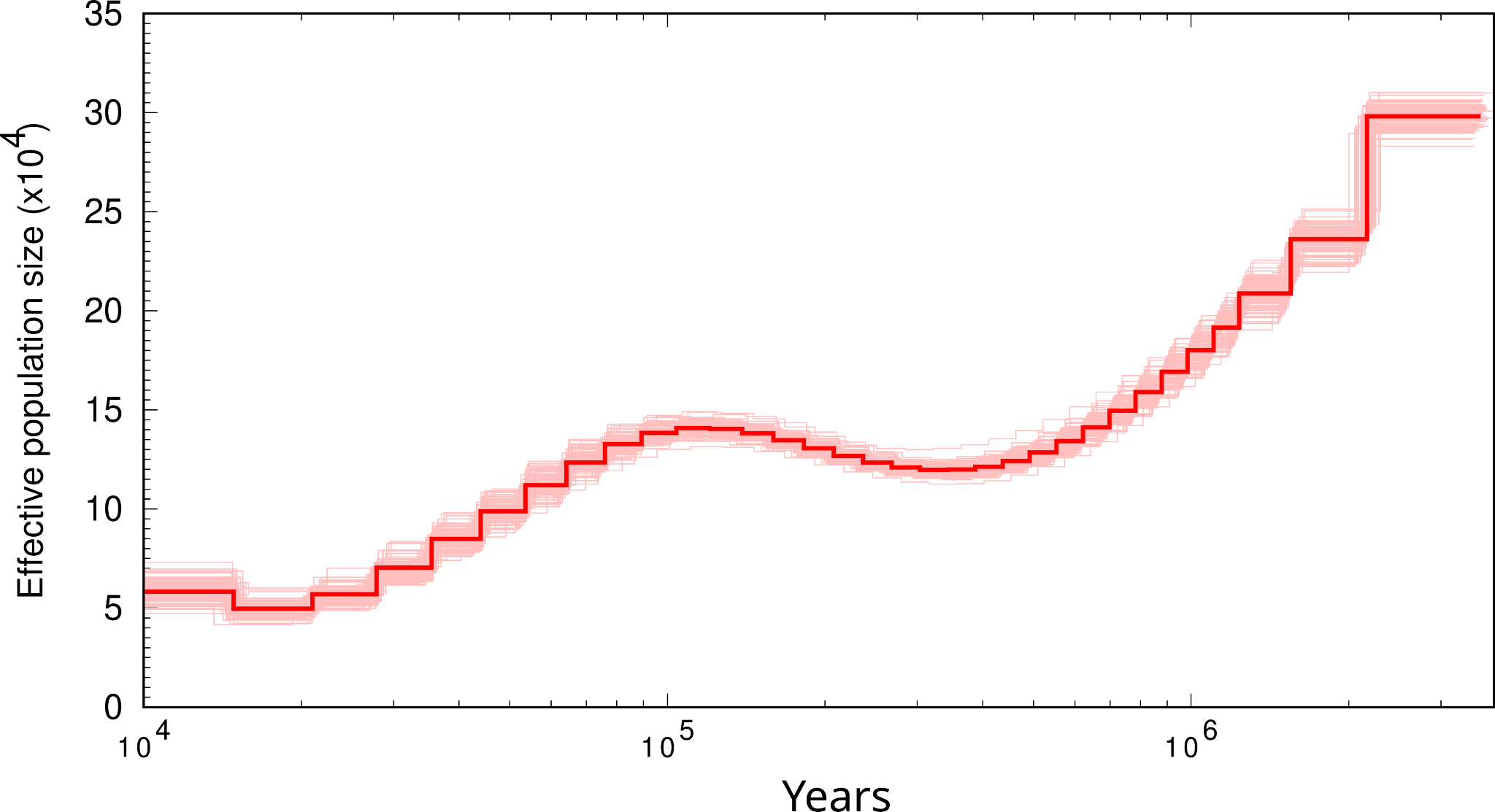

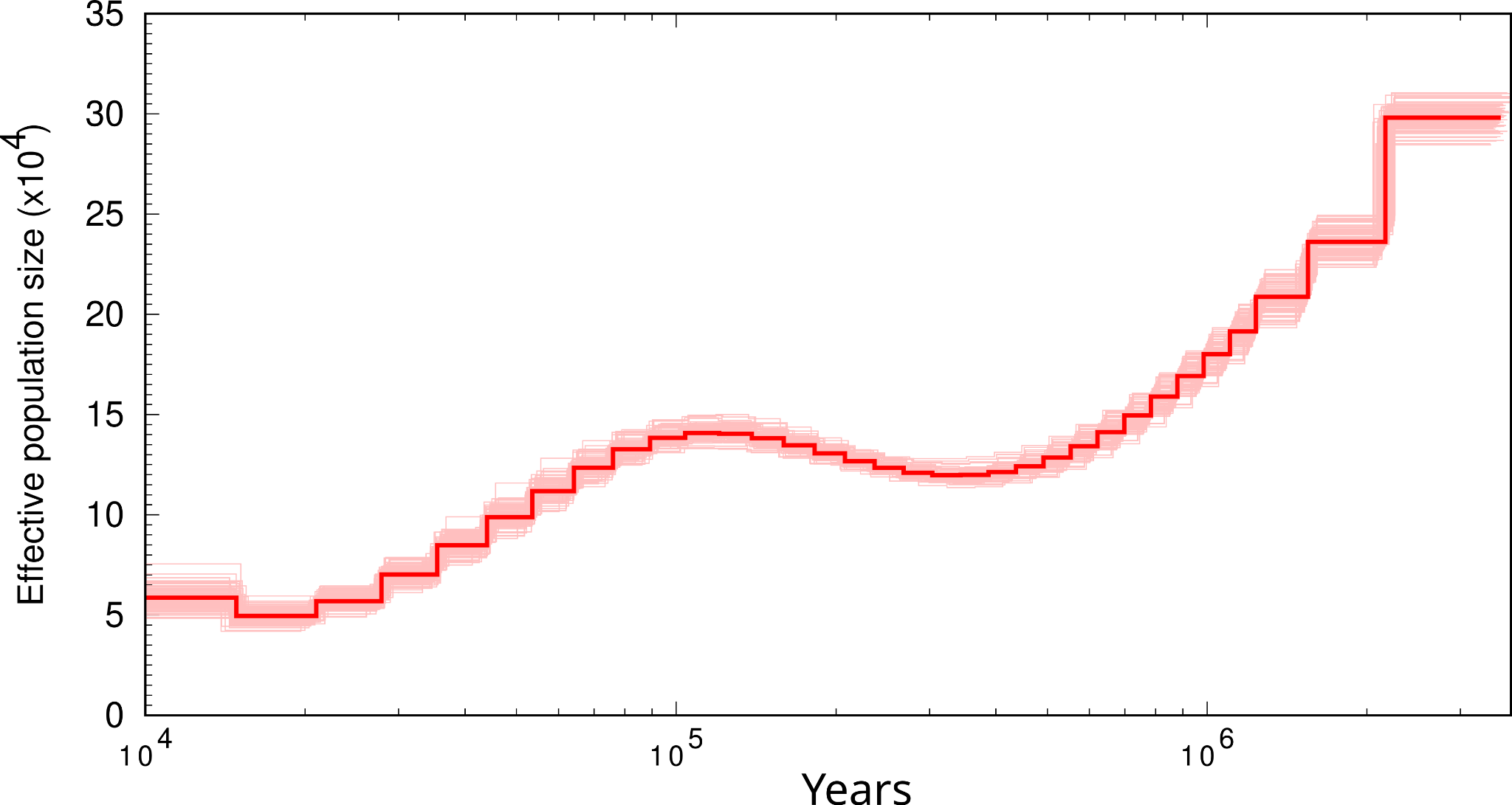

#### *Cnemophilus loriae*

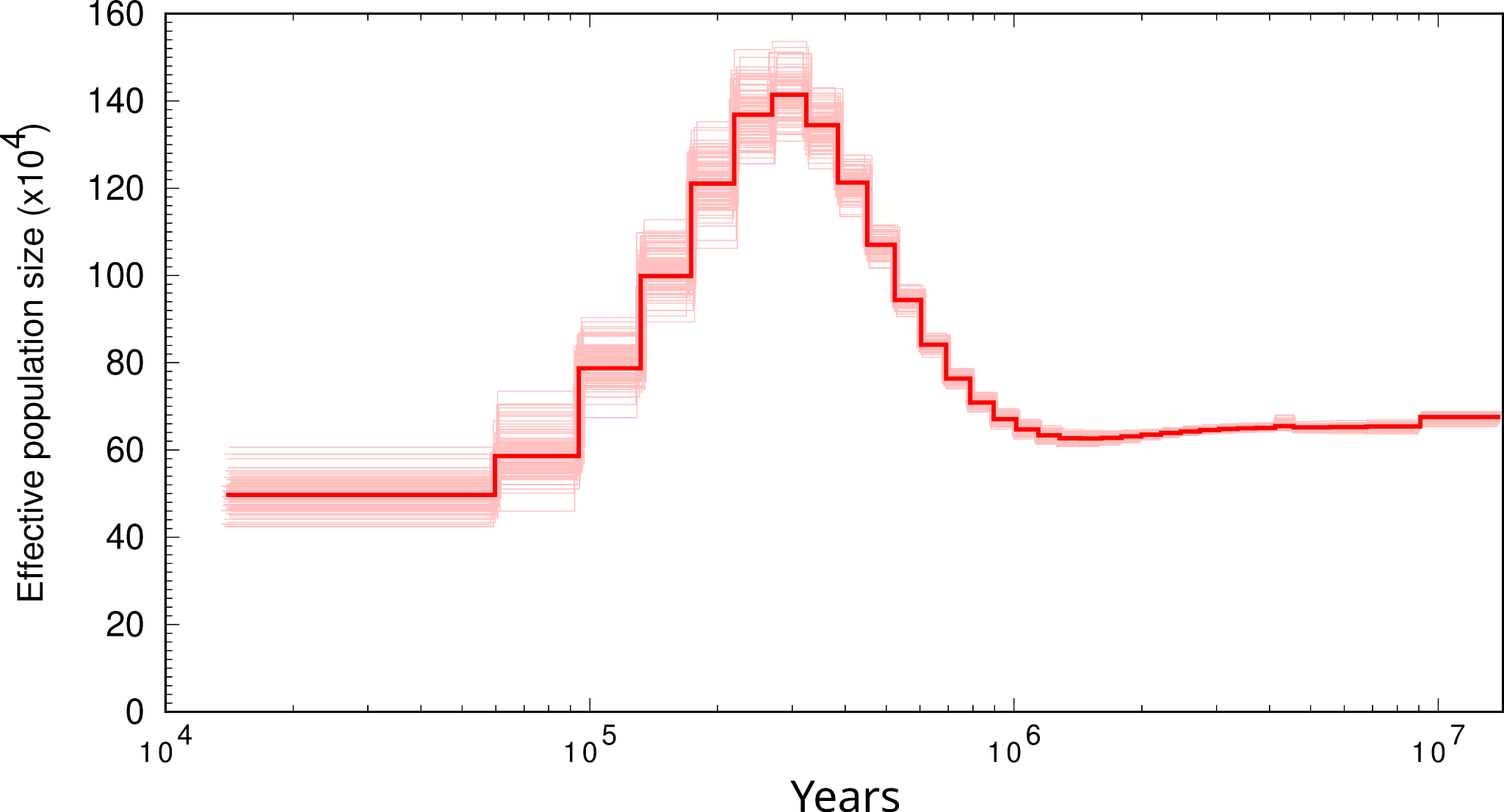

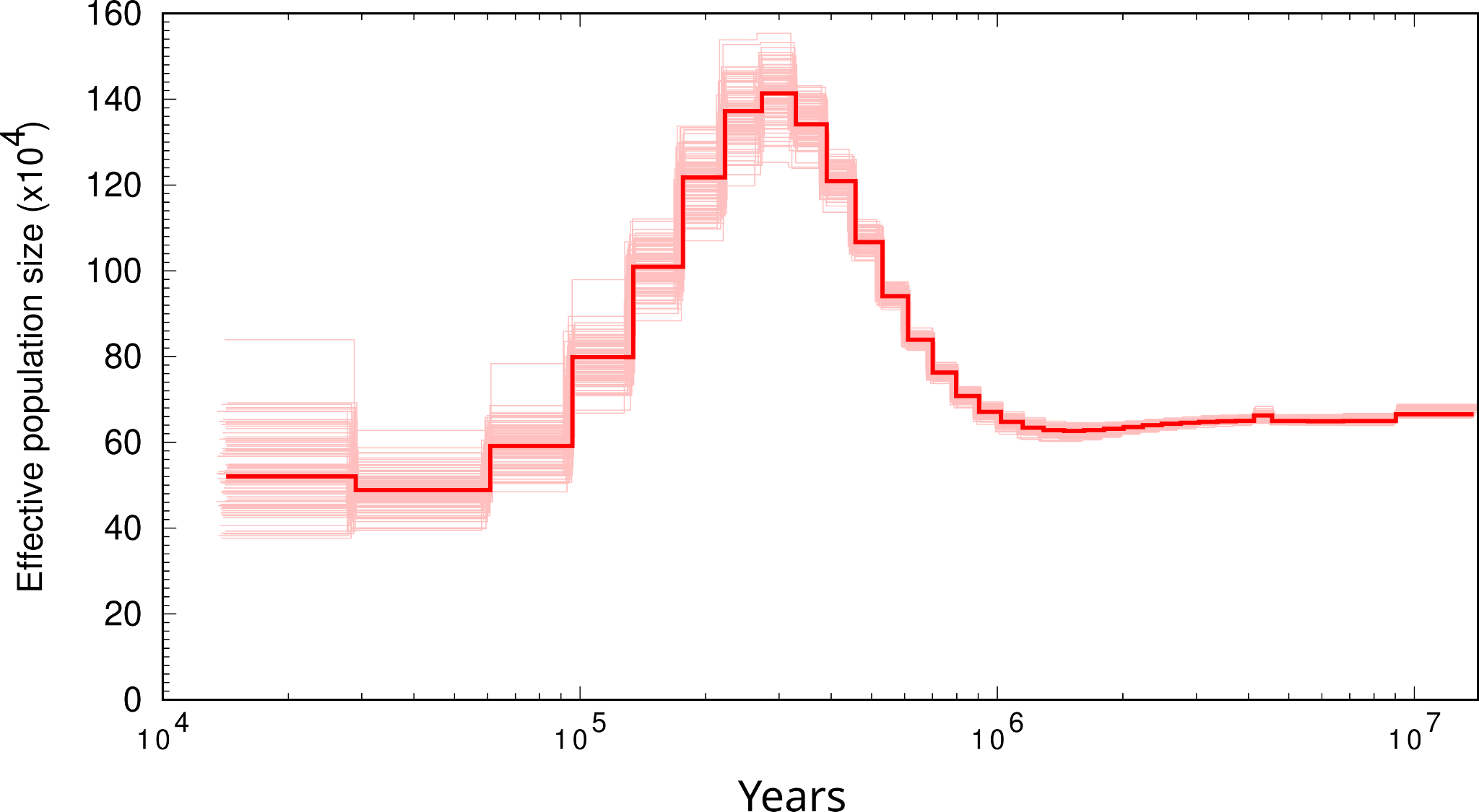

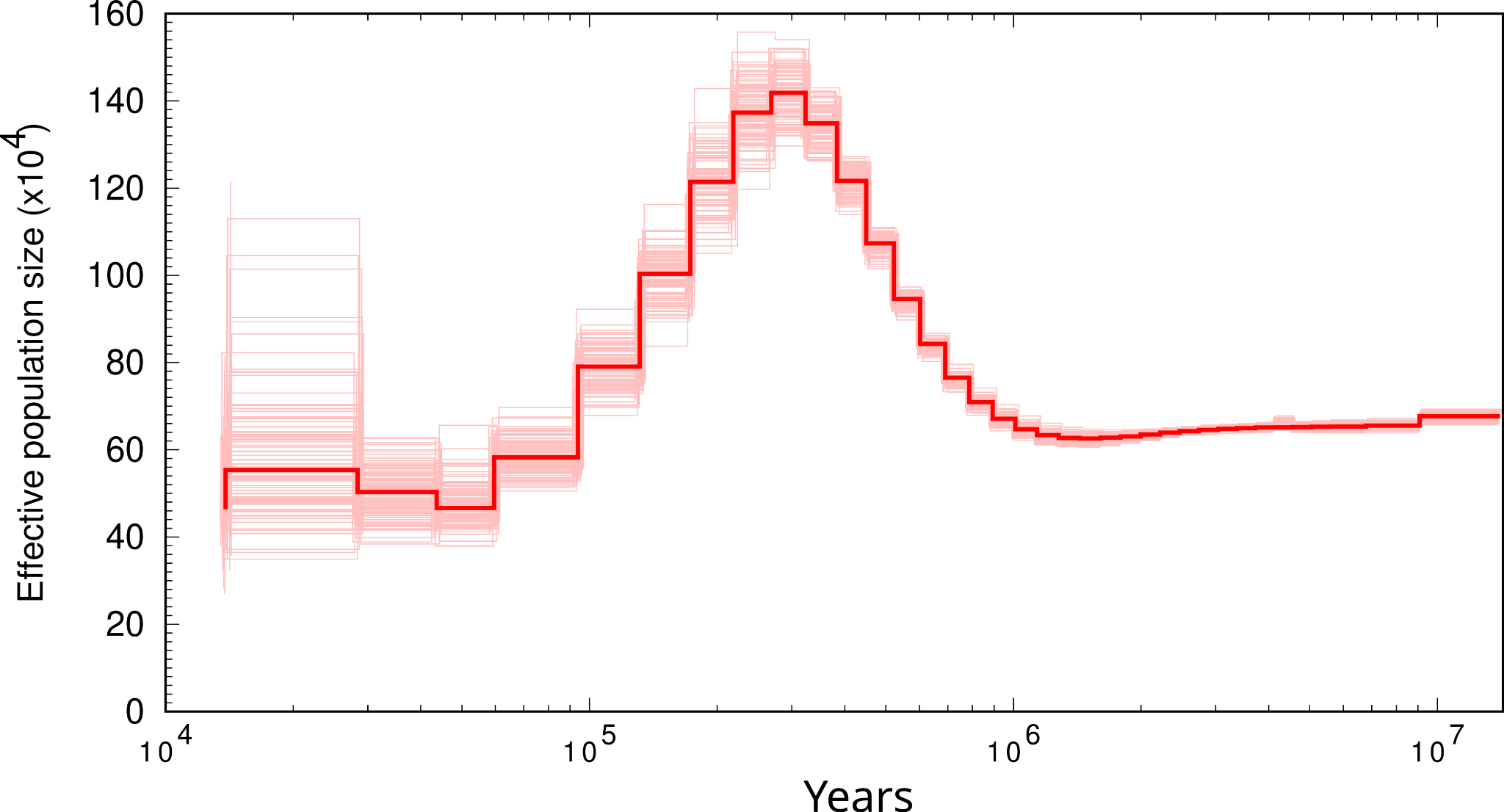

#### *Dicaeum eximium*

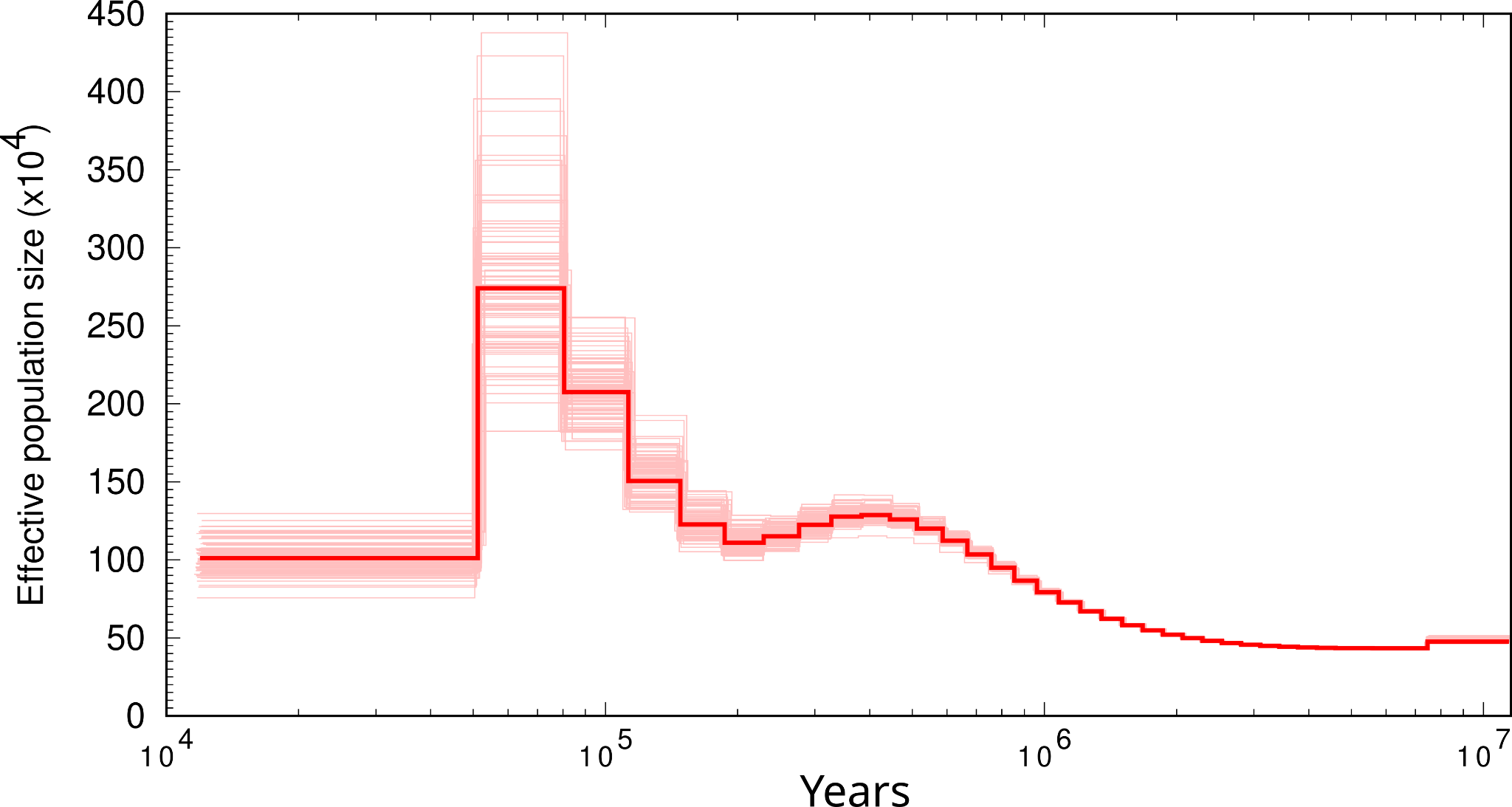

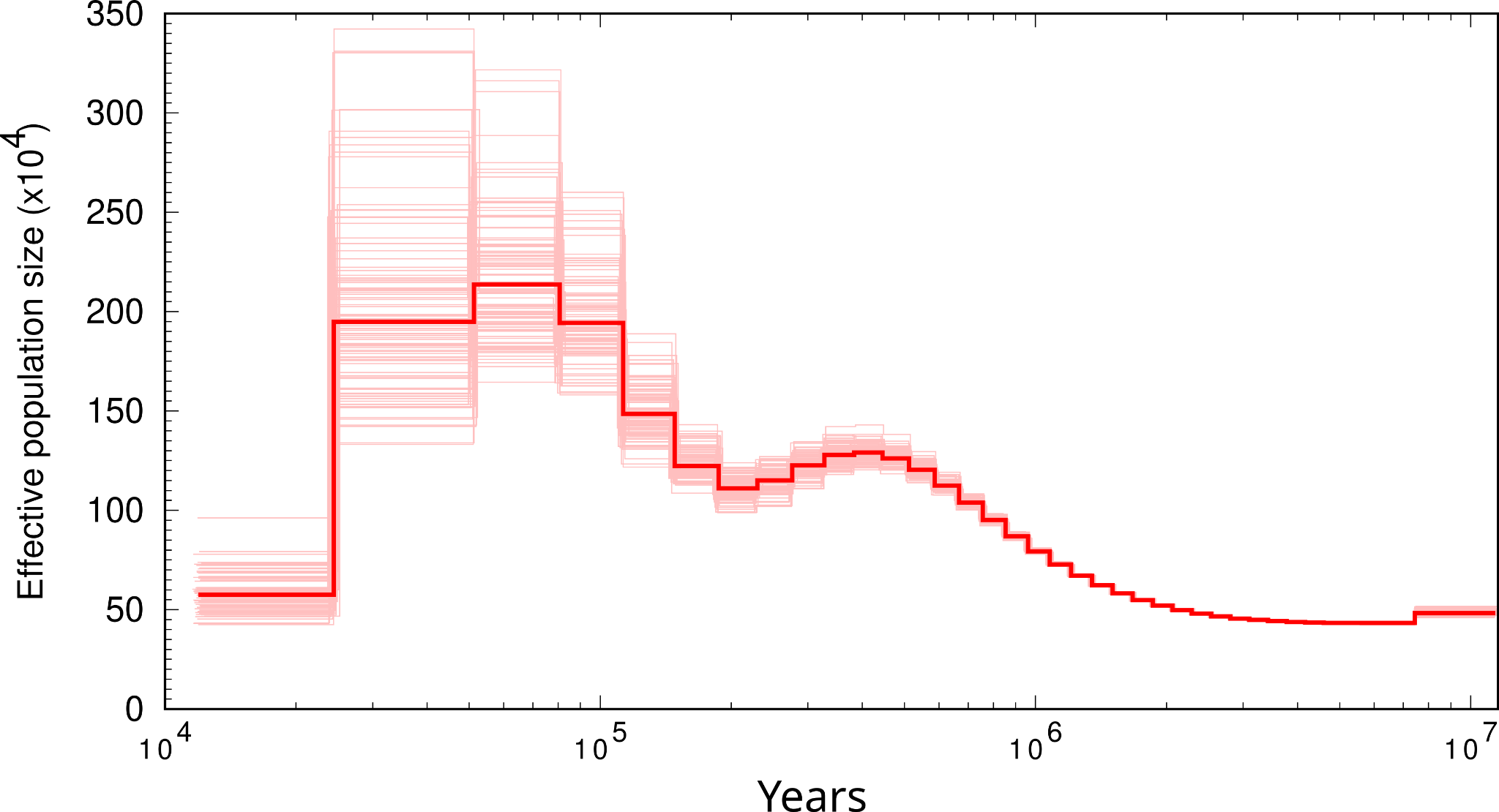

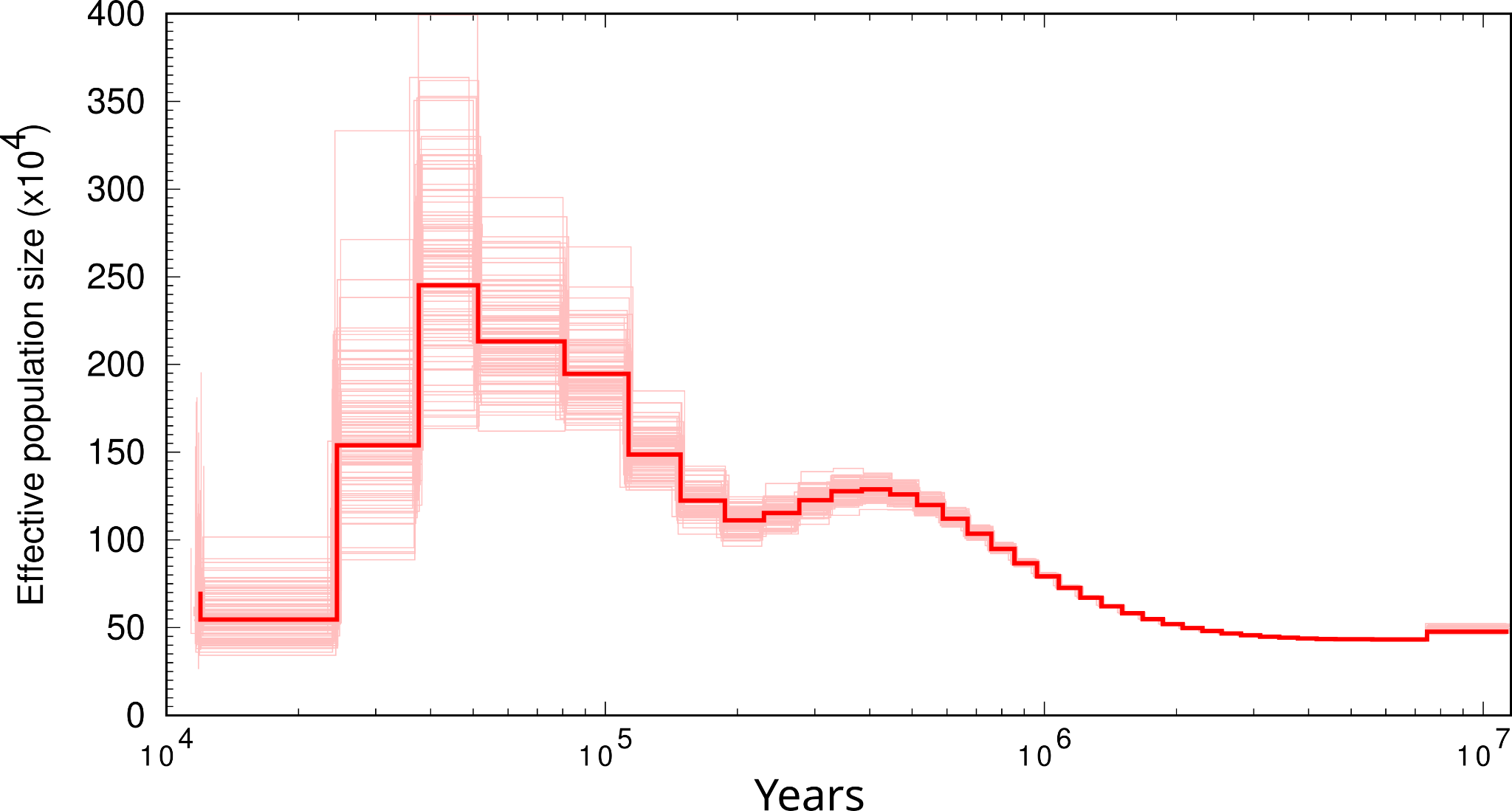

#### *Diphyllodes magnificus*

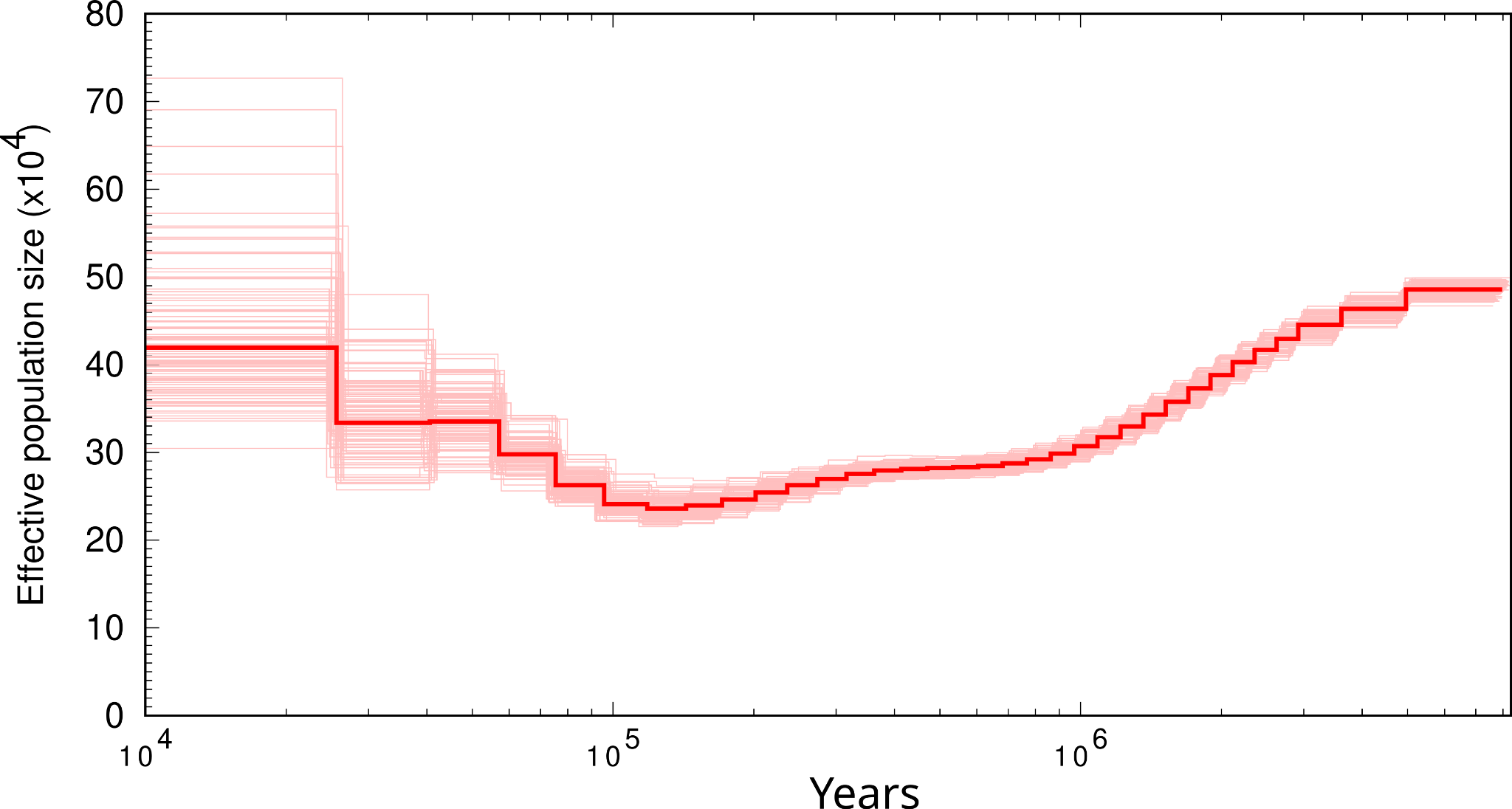

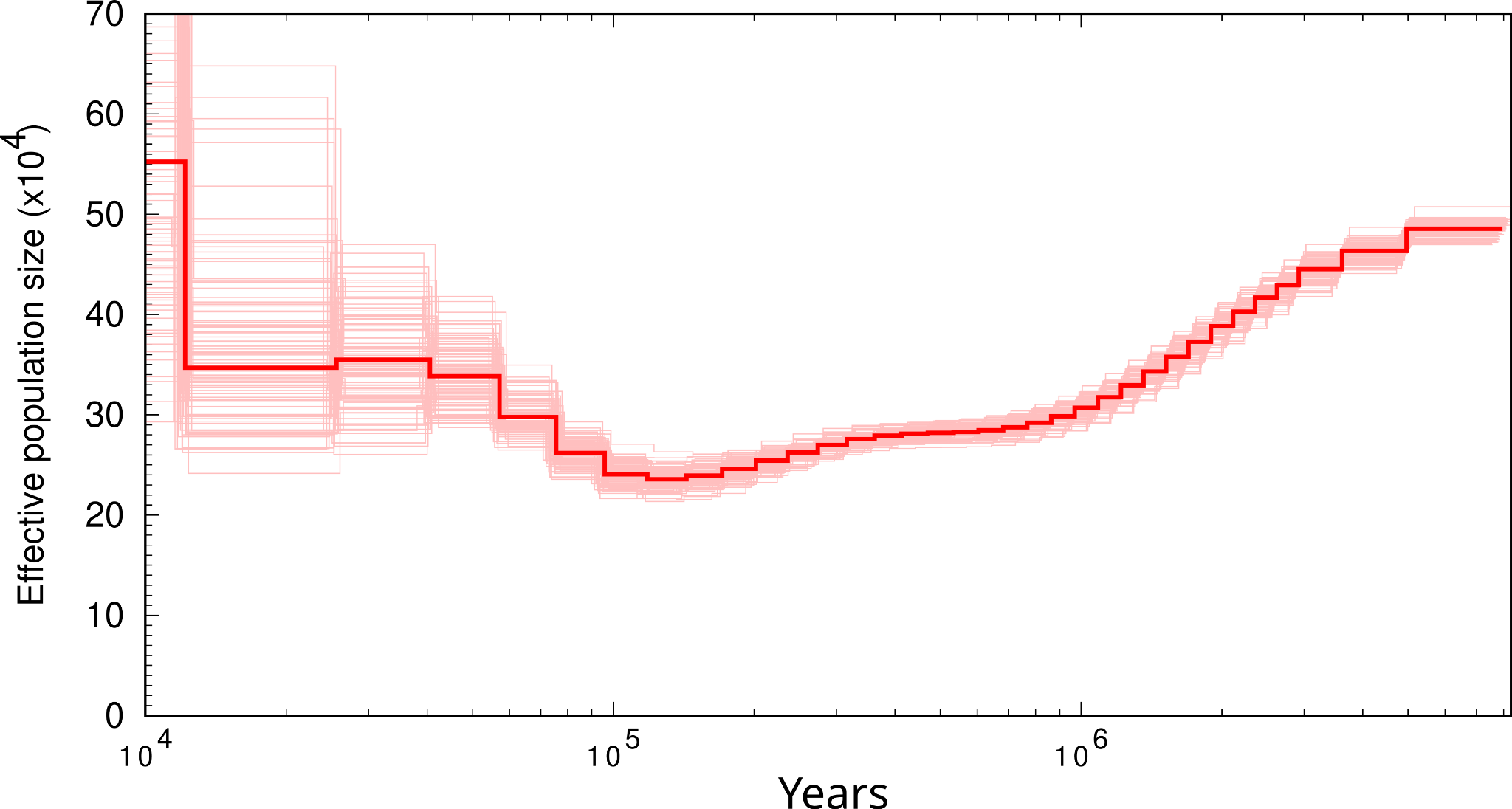

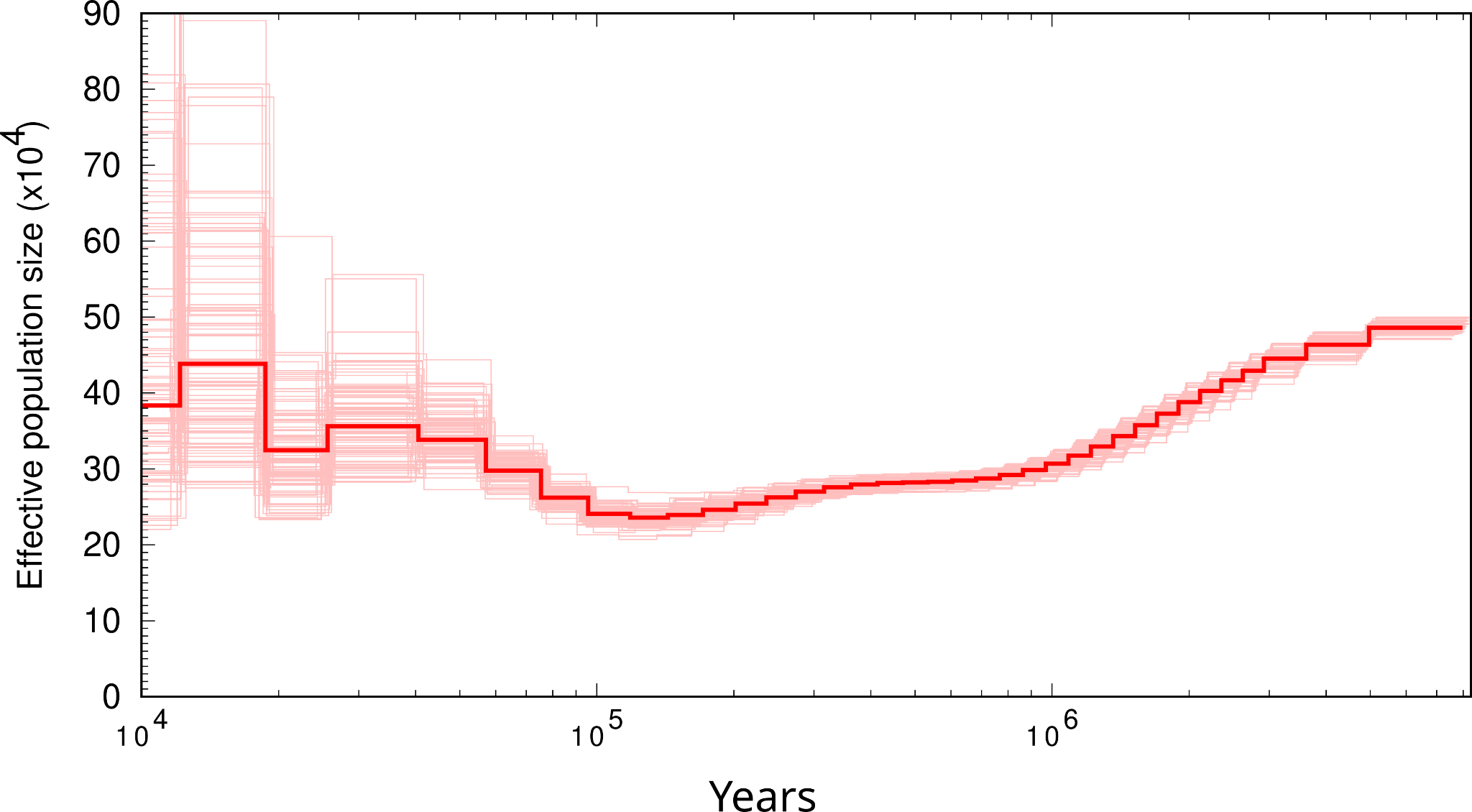

#### *Irena cyanogastra*

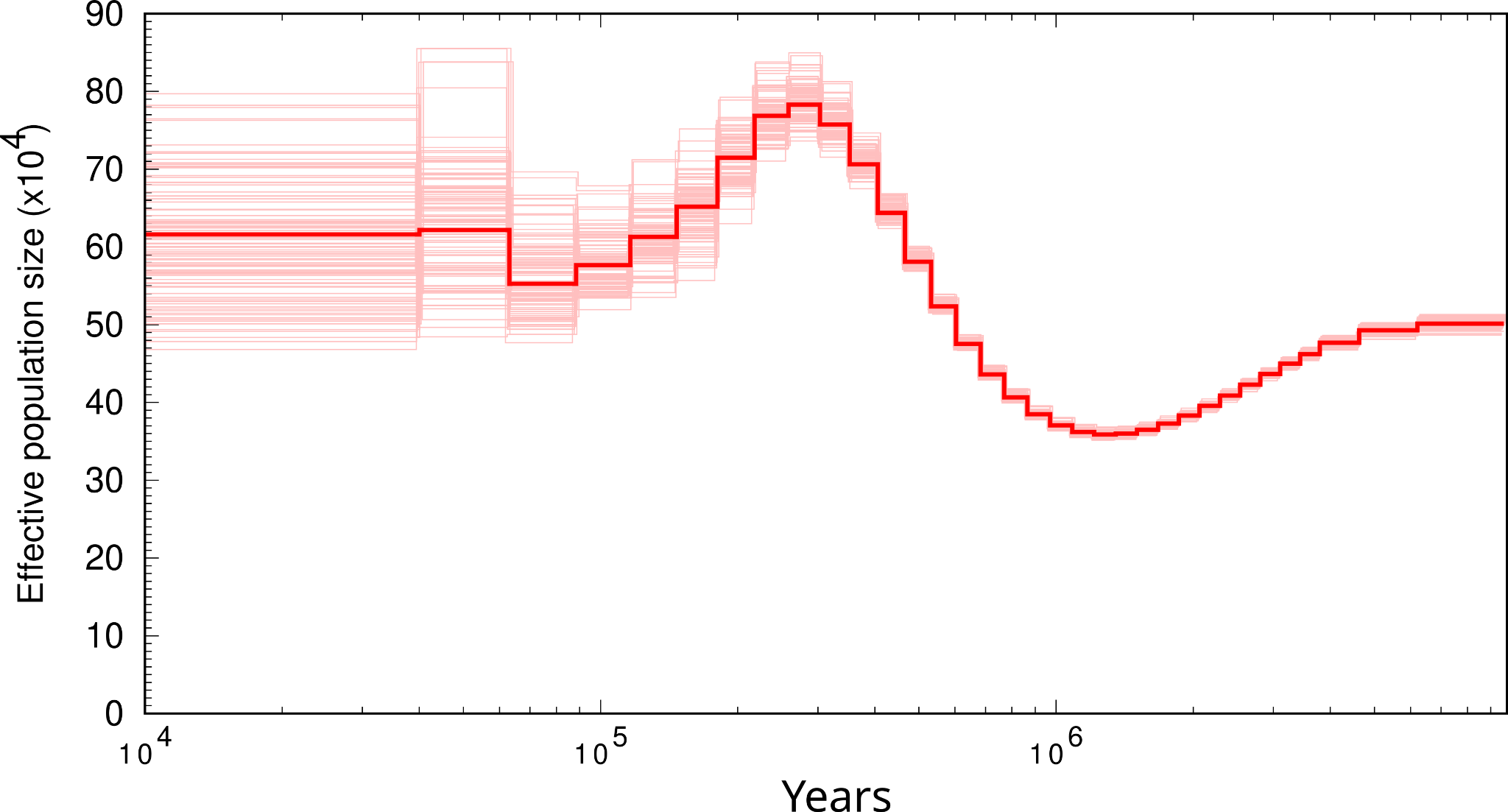

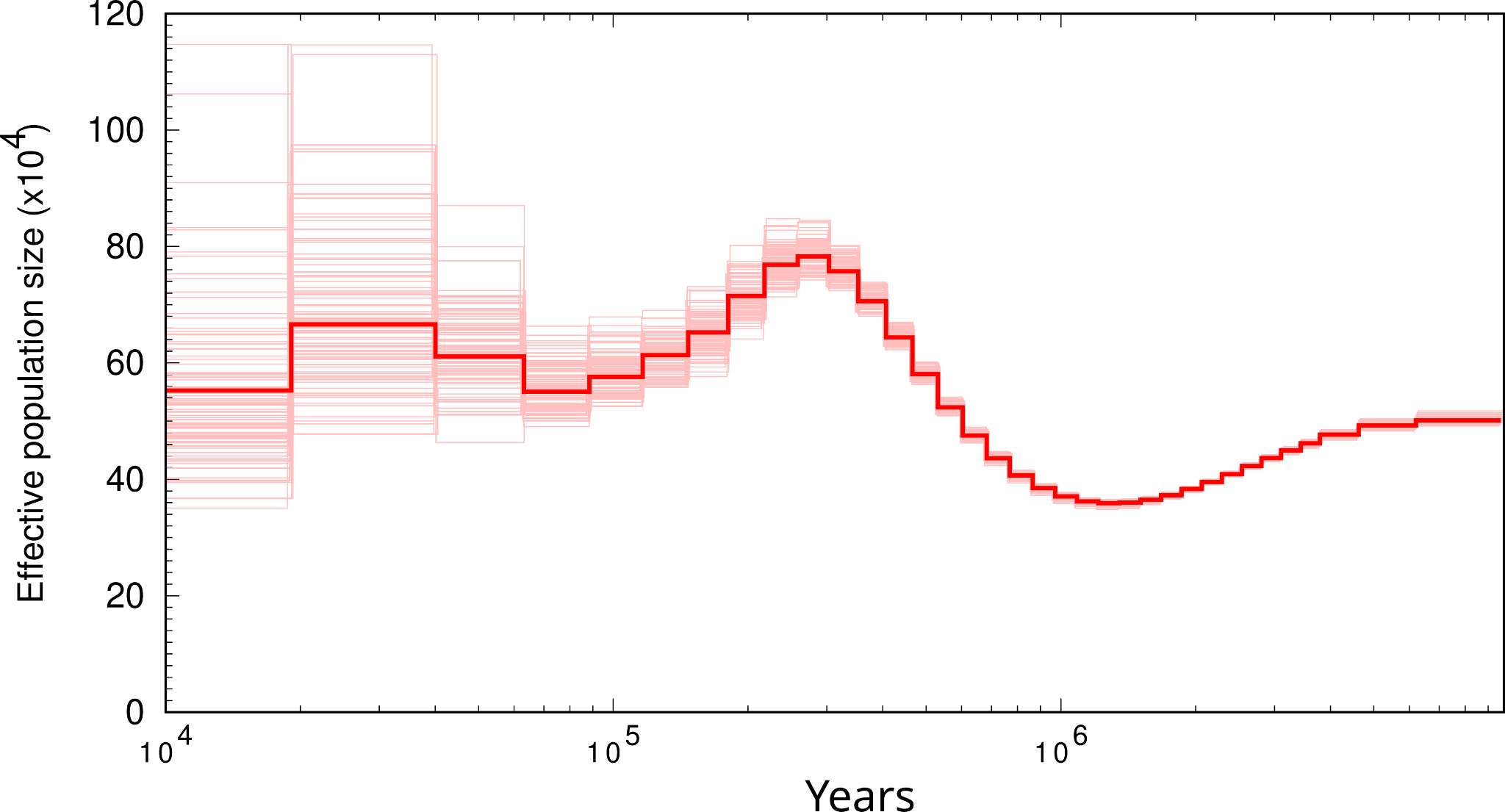

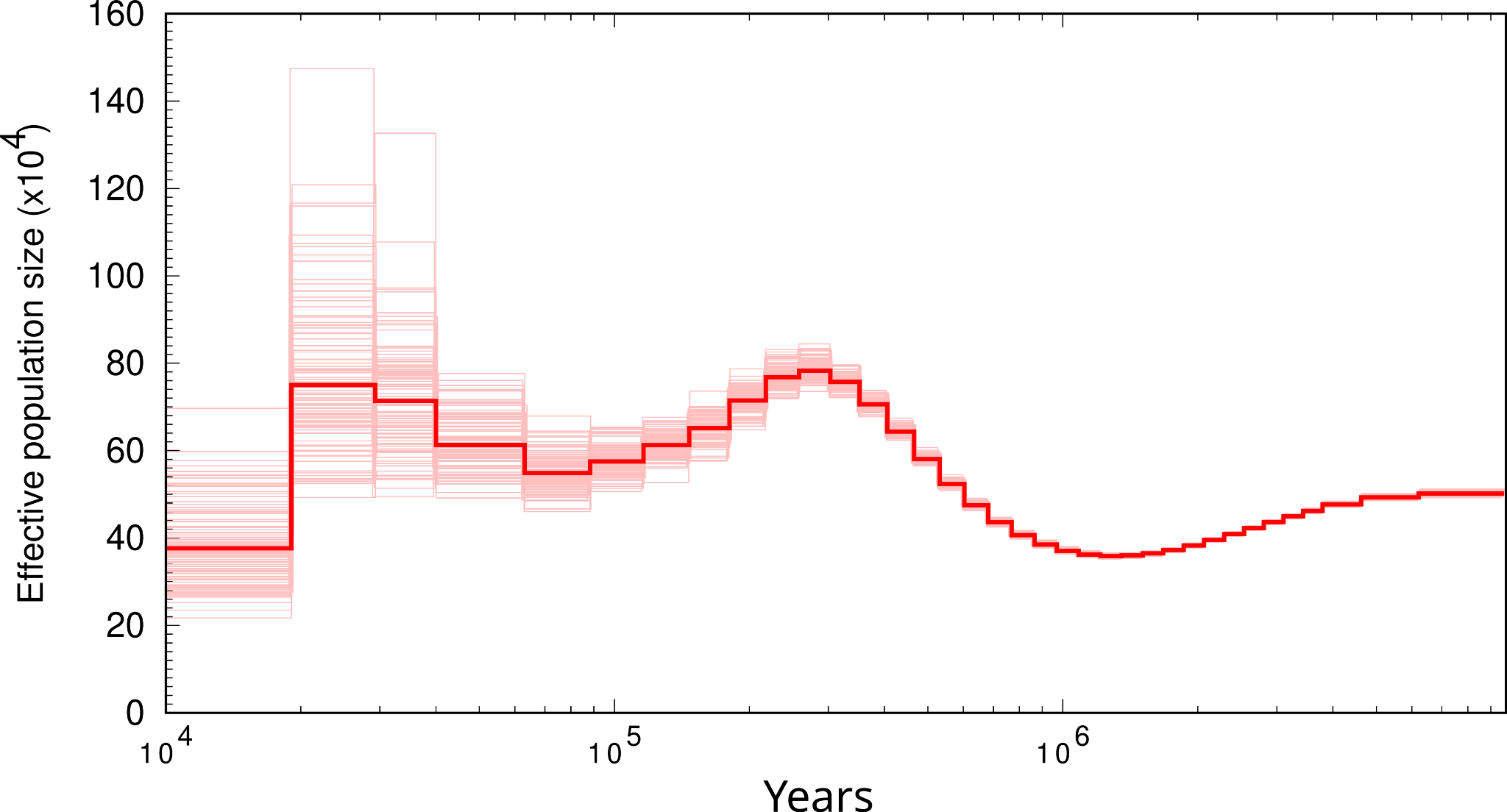

*Machaerirhynchus nigripectus*

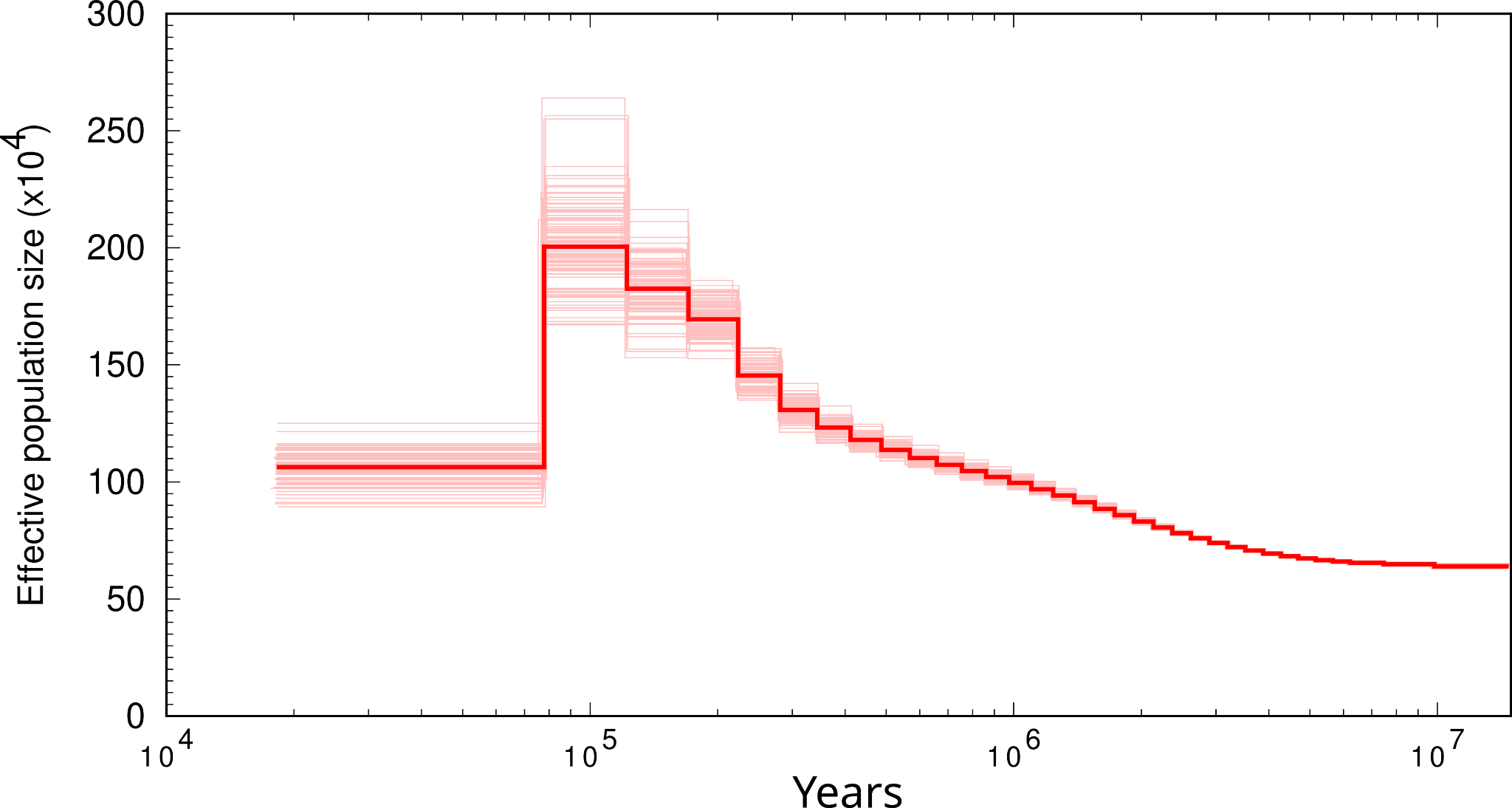

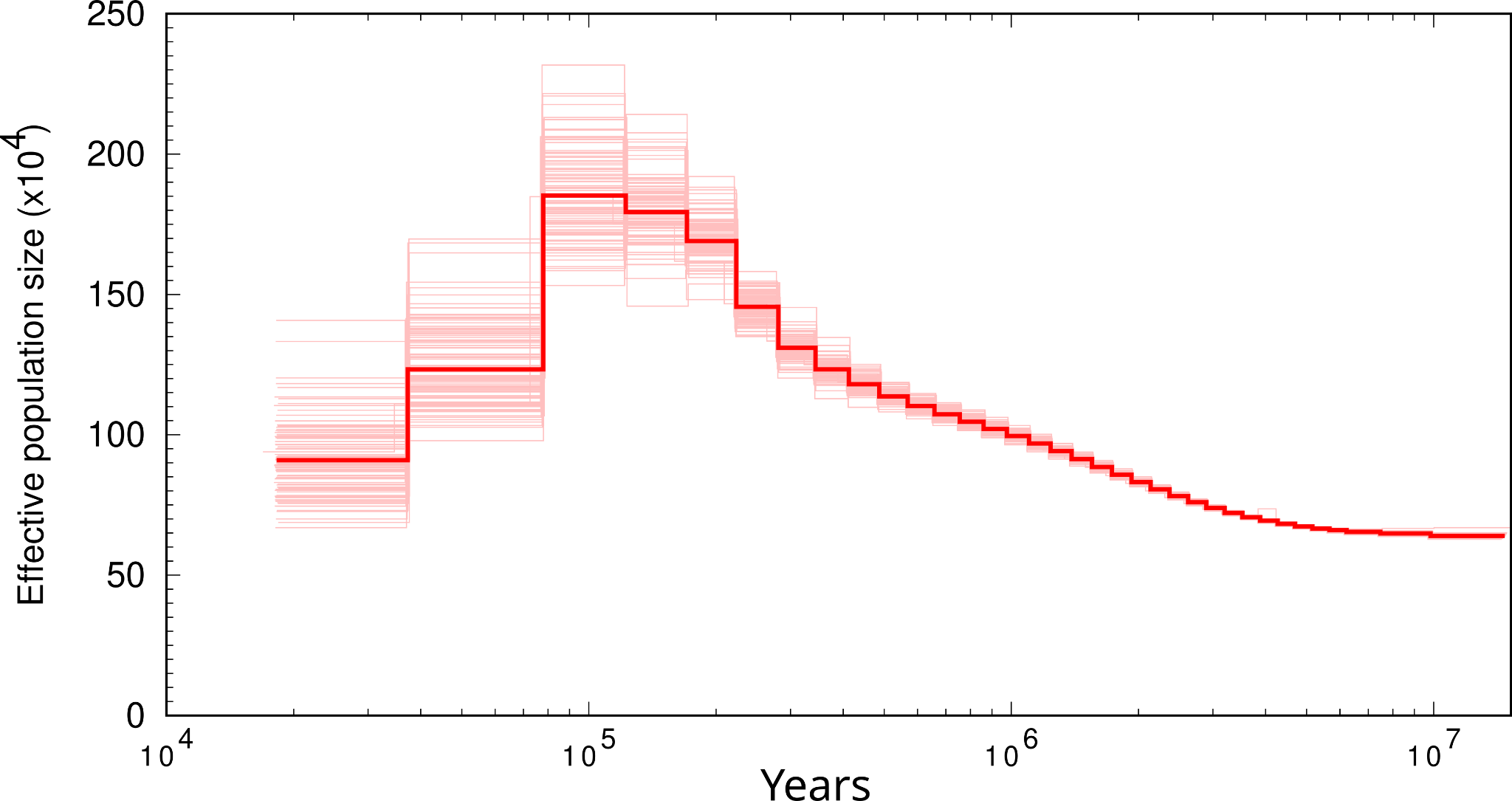

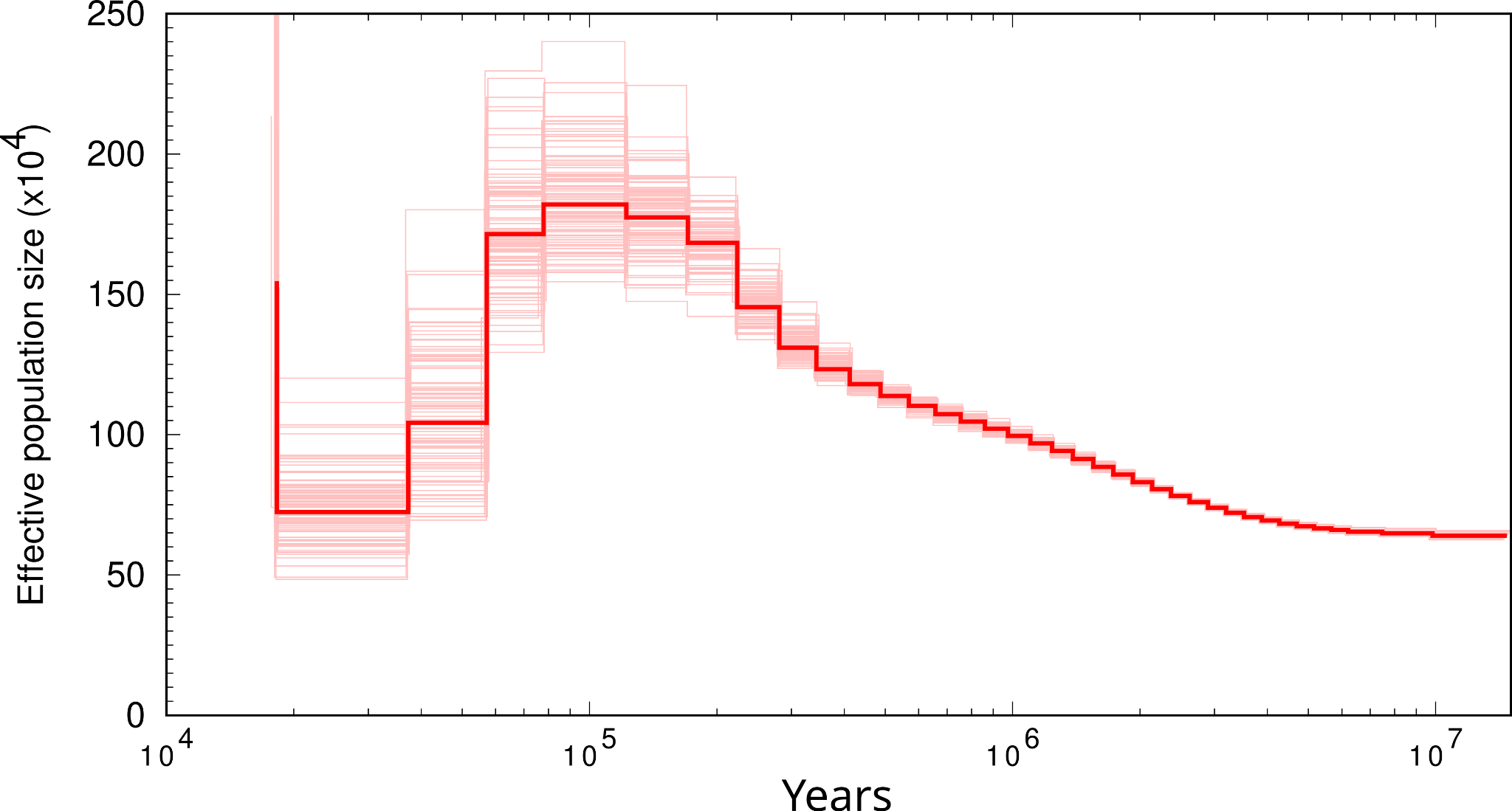

#### *Melanocharis versteri*

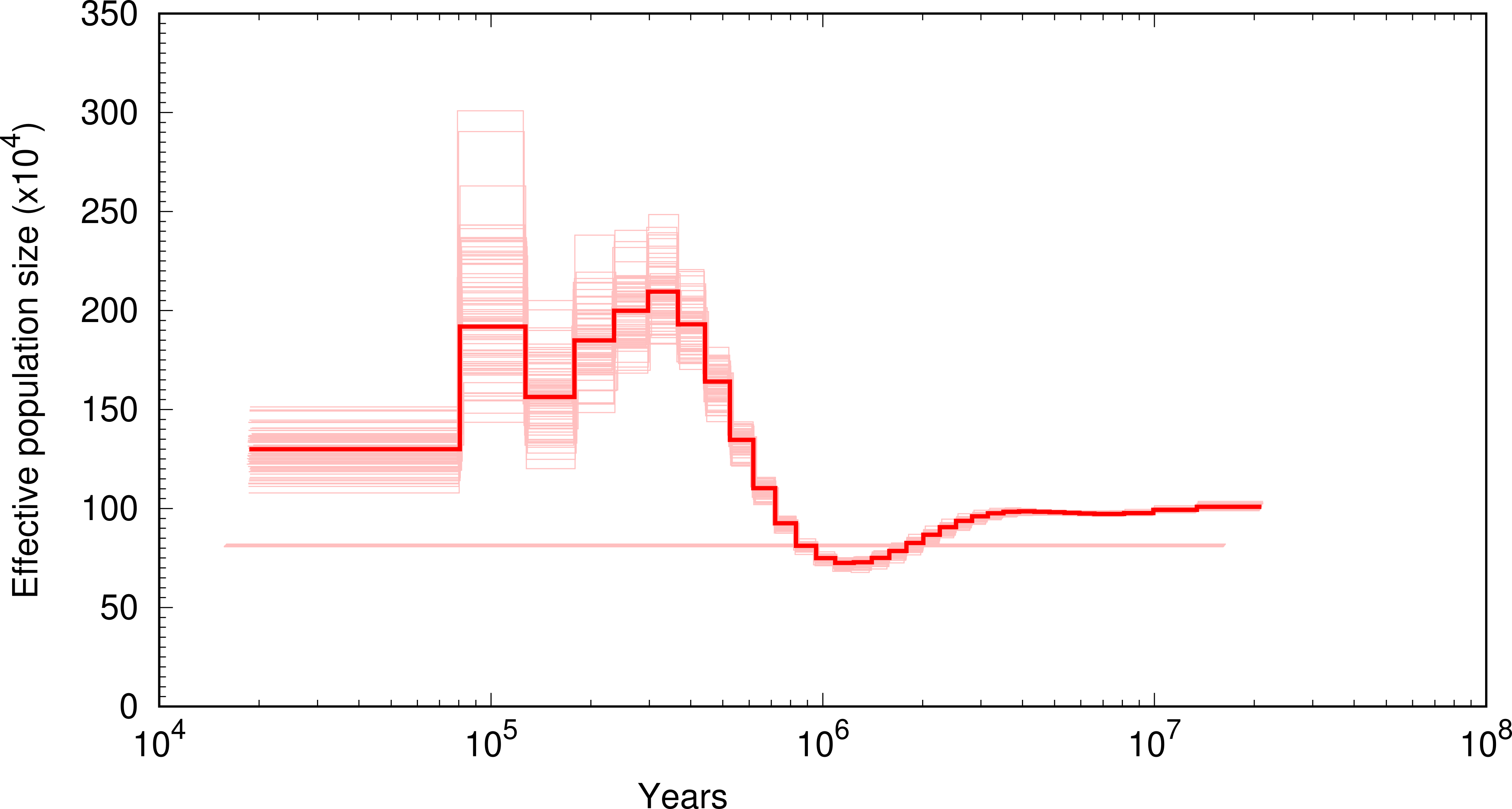

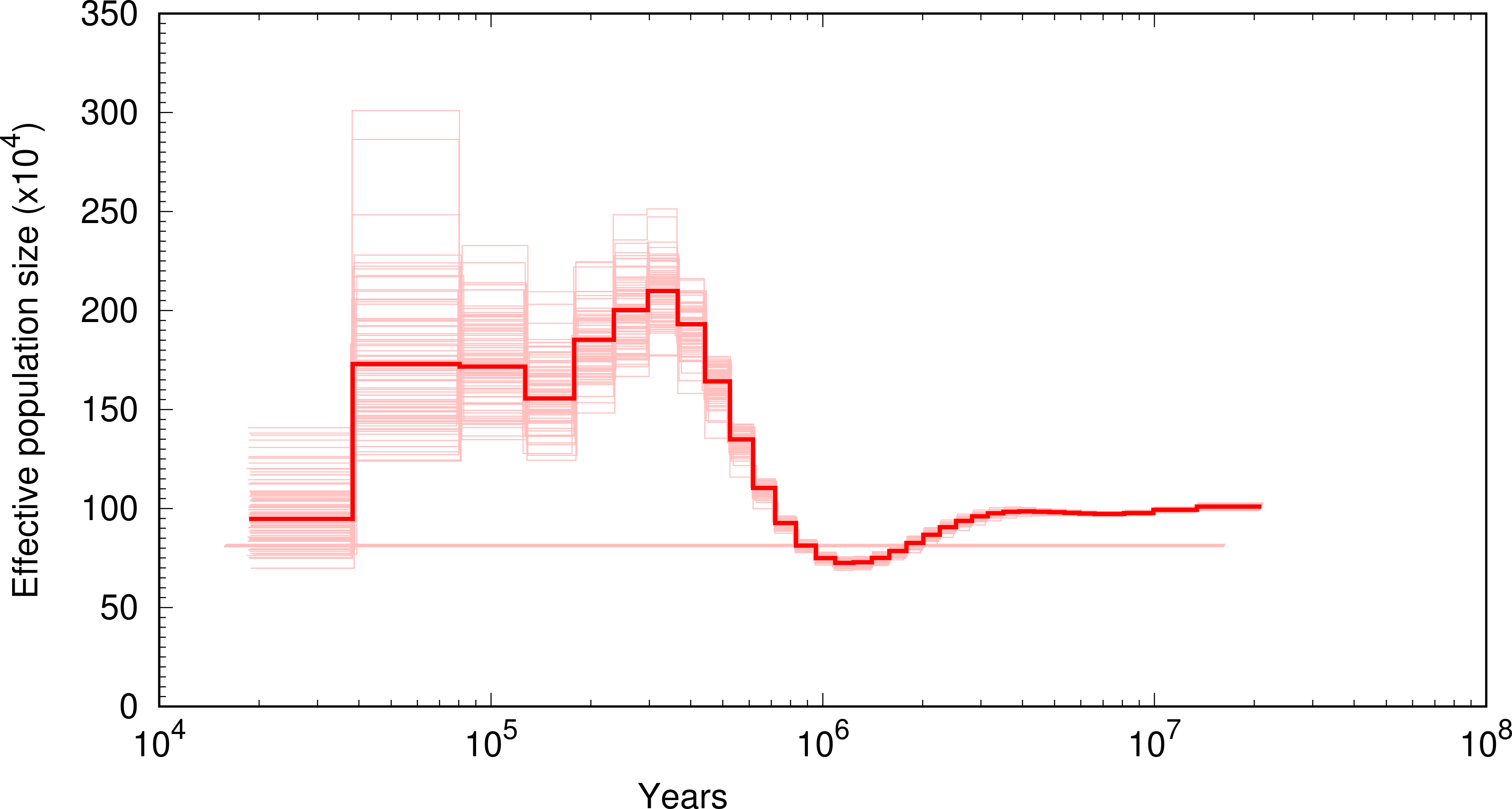

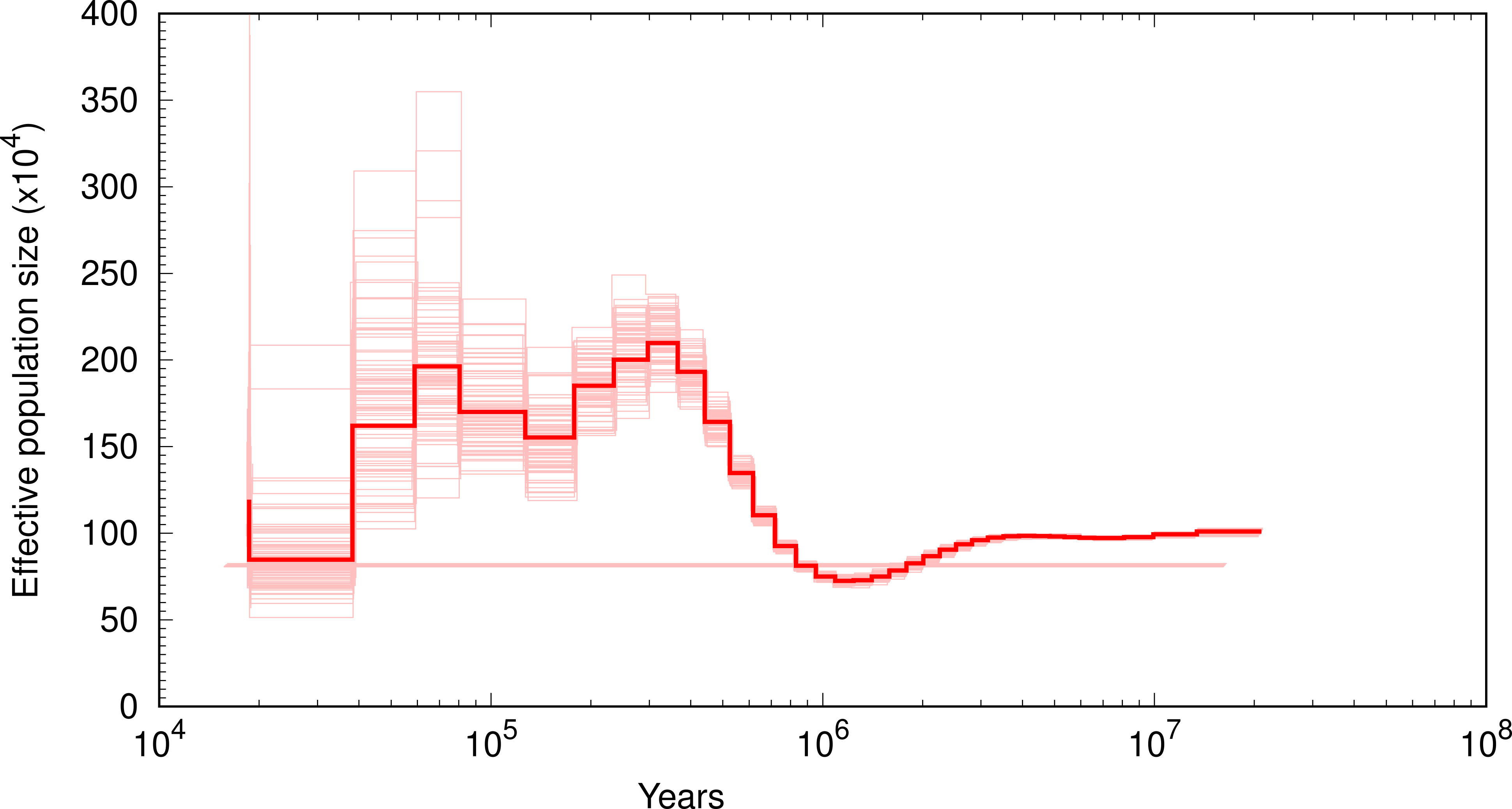

#### *Myiagra hebetior*

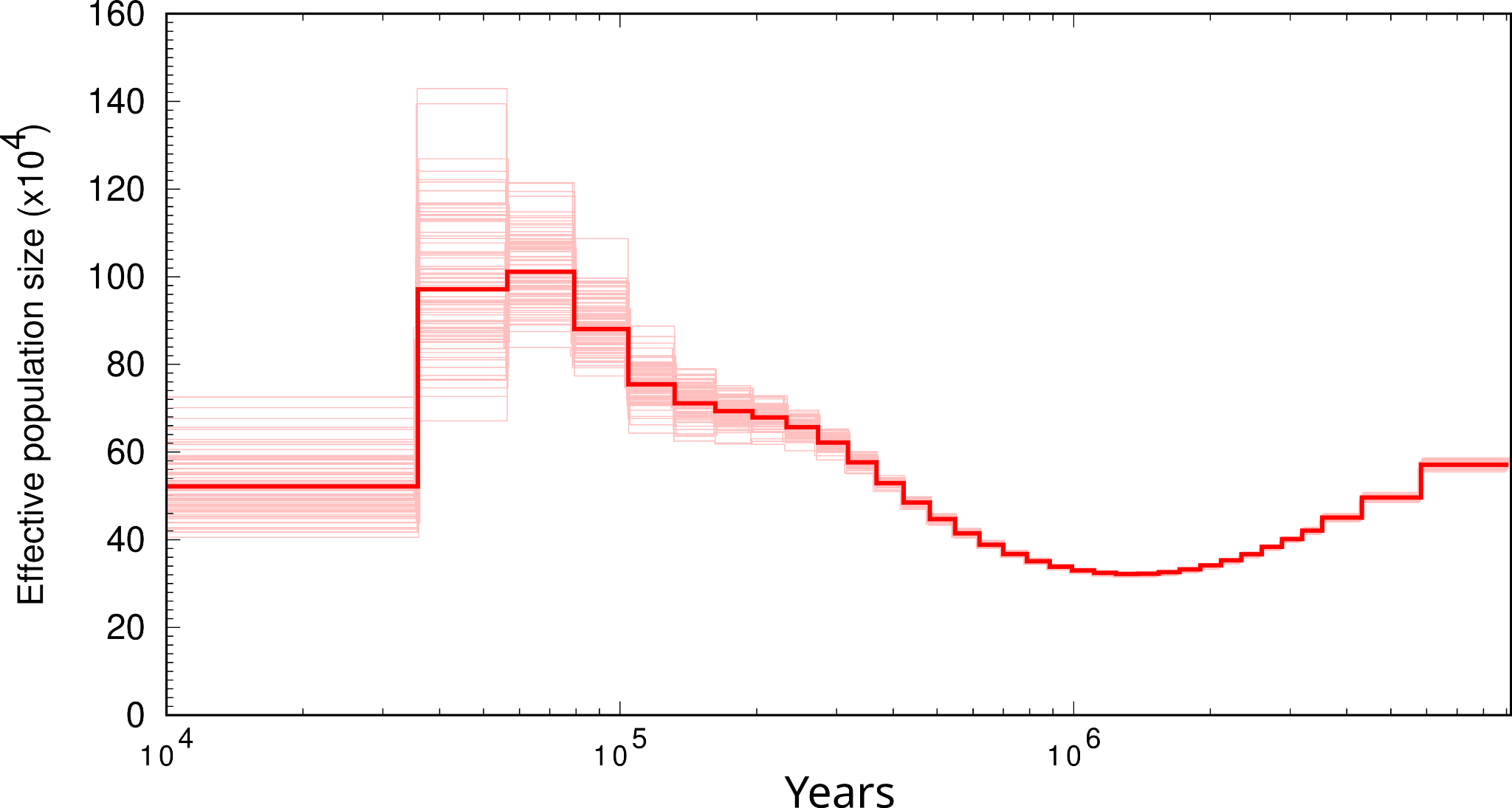

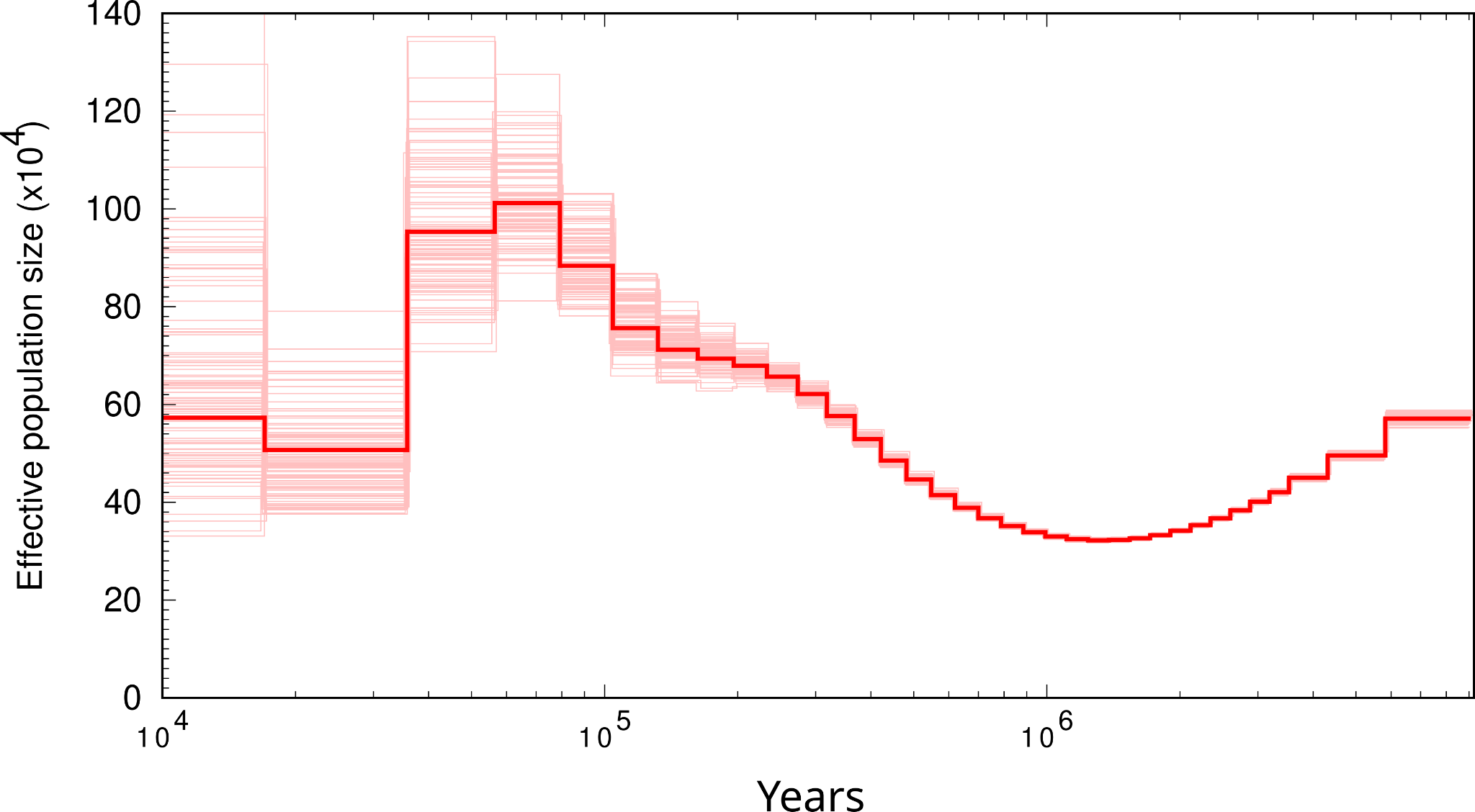

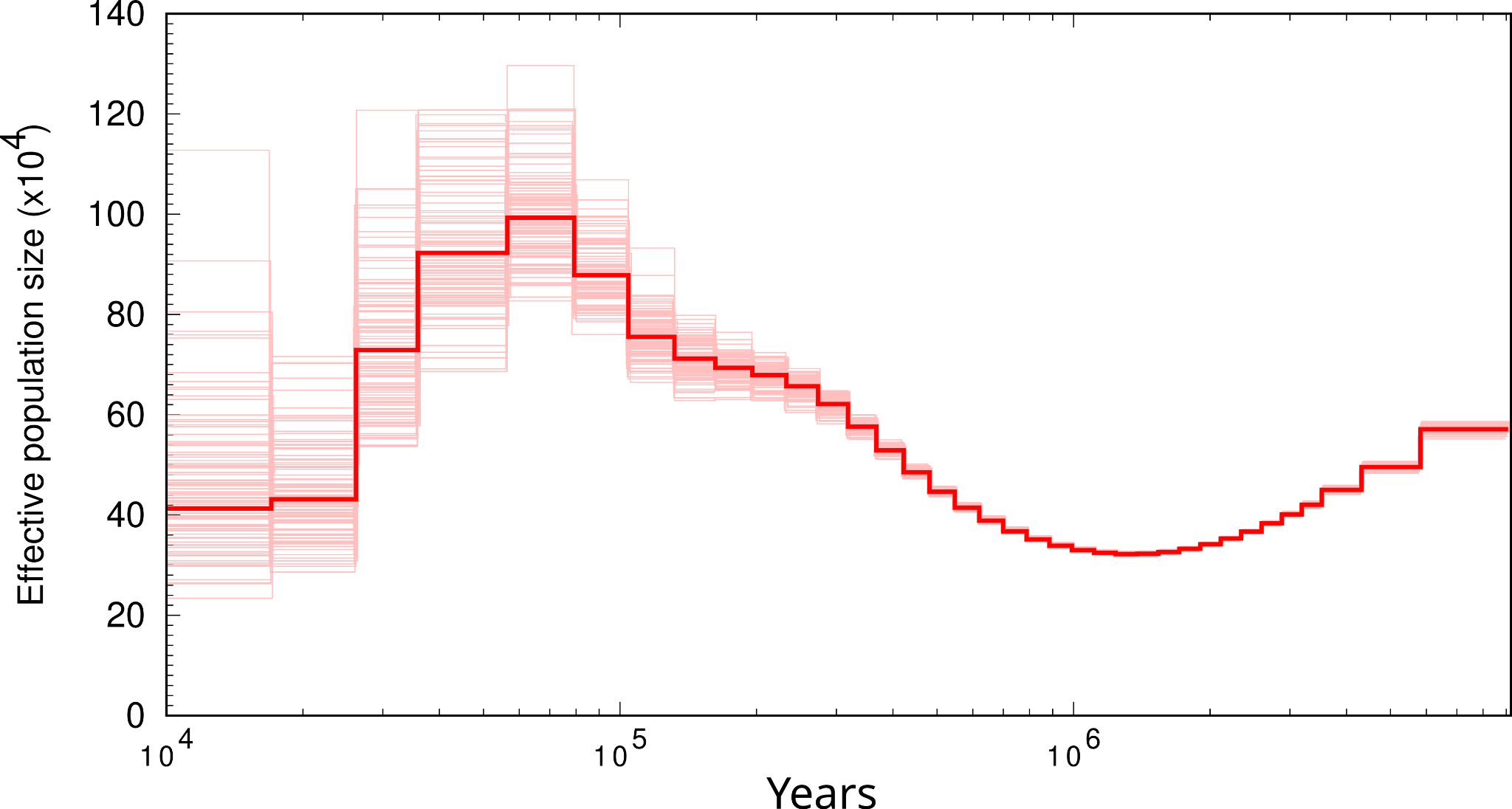

#### *Pachycephala philippinensis*

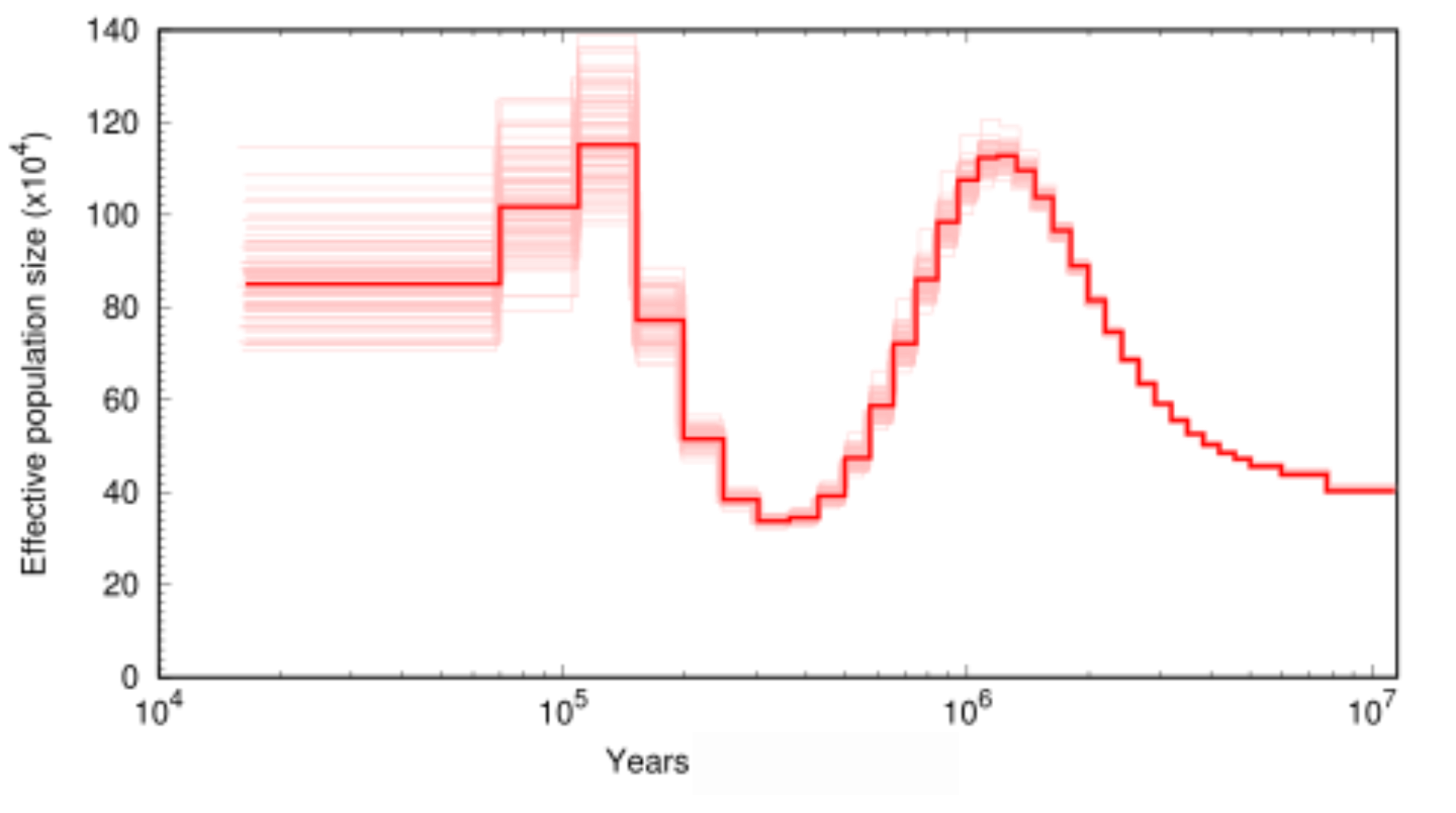

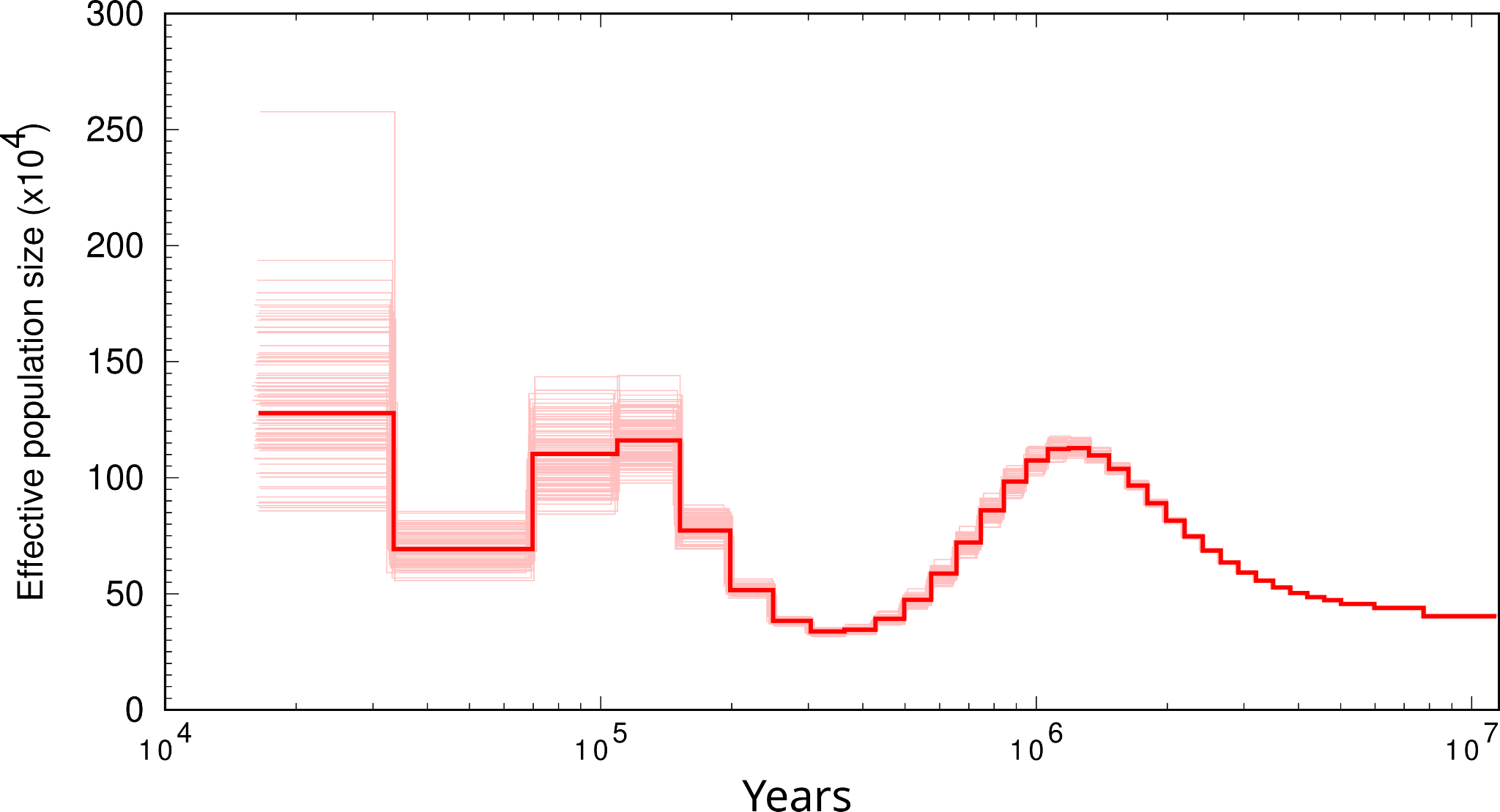

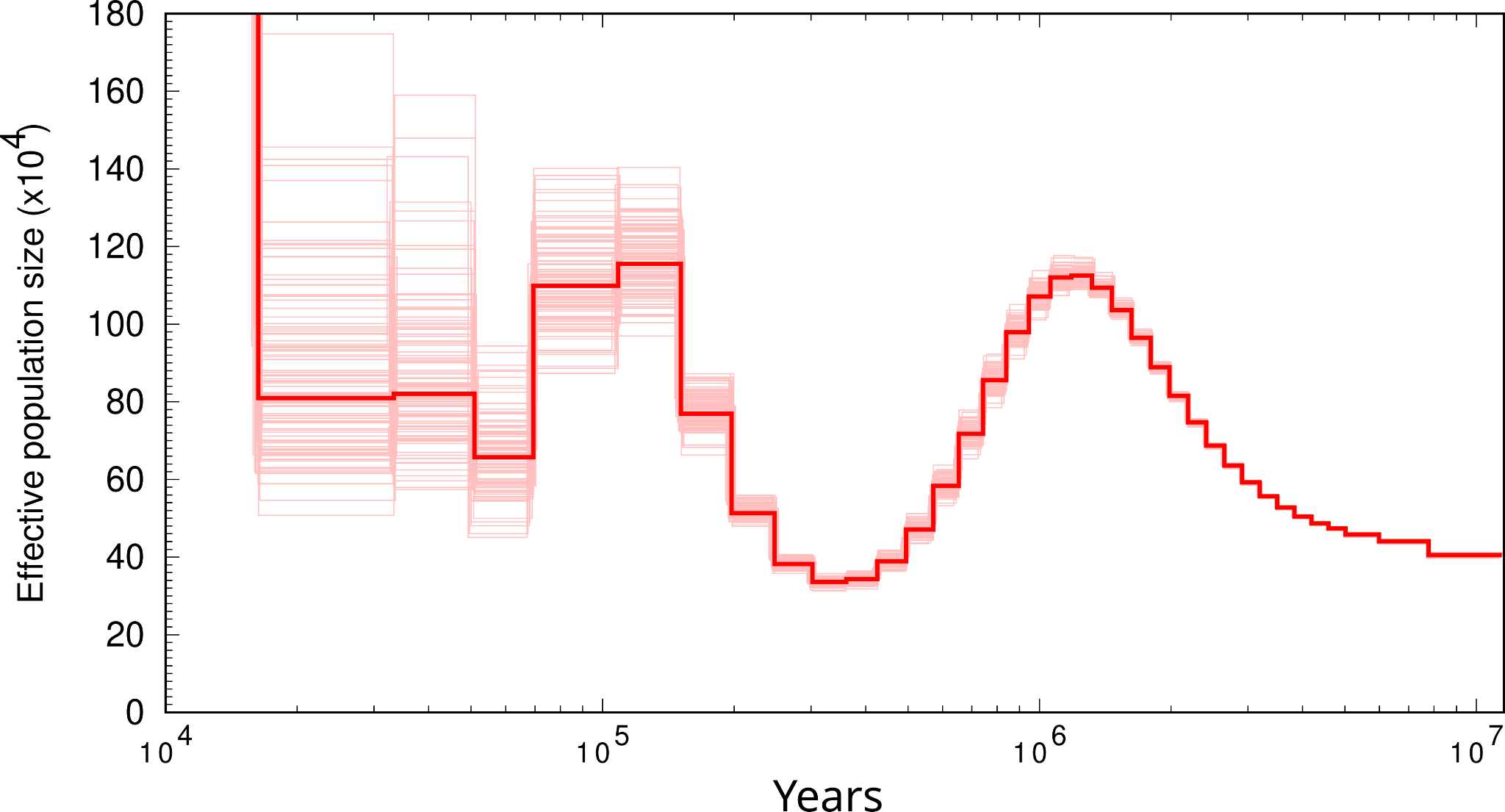

*Paradisaea raggiana*

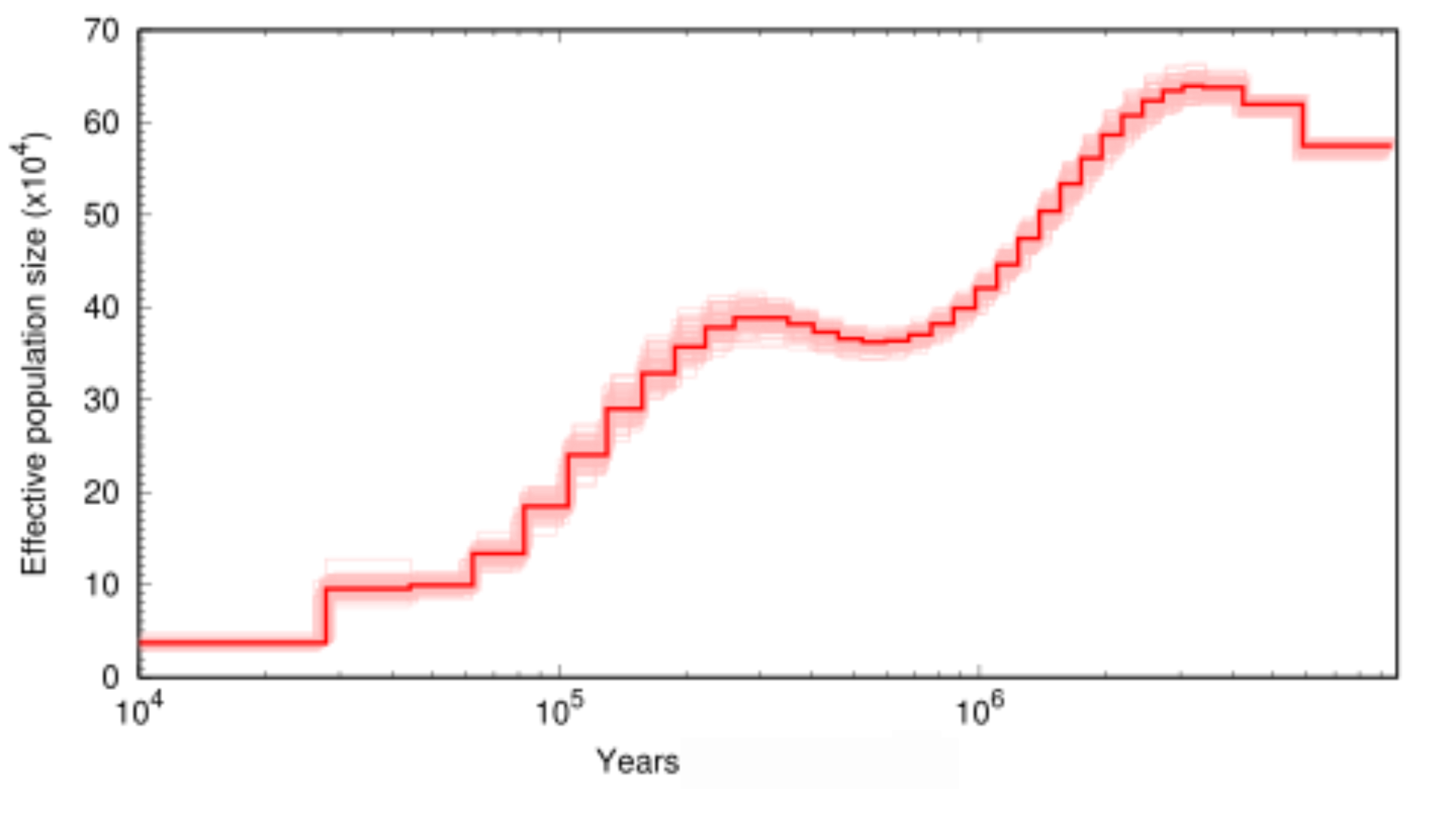

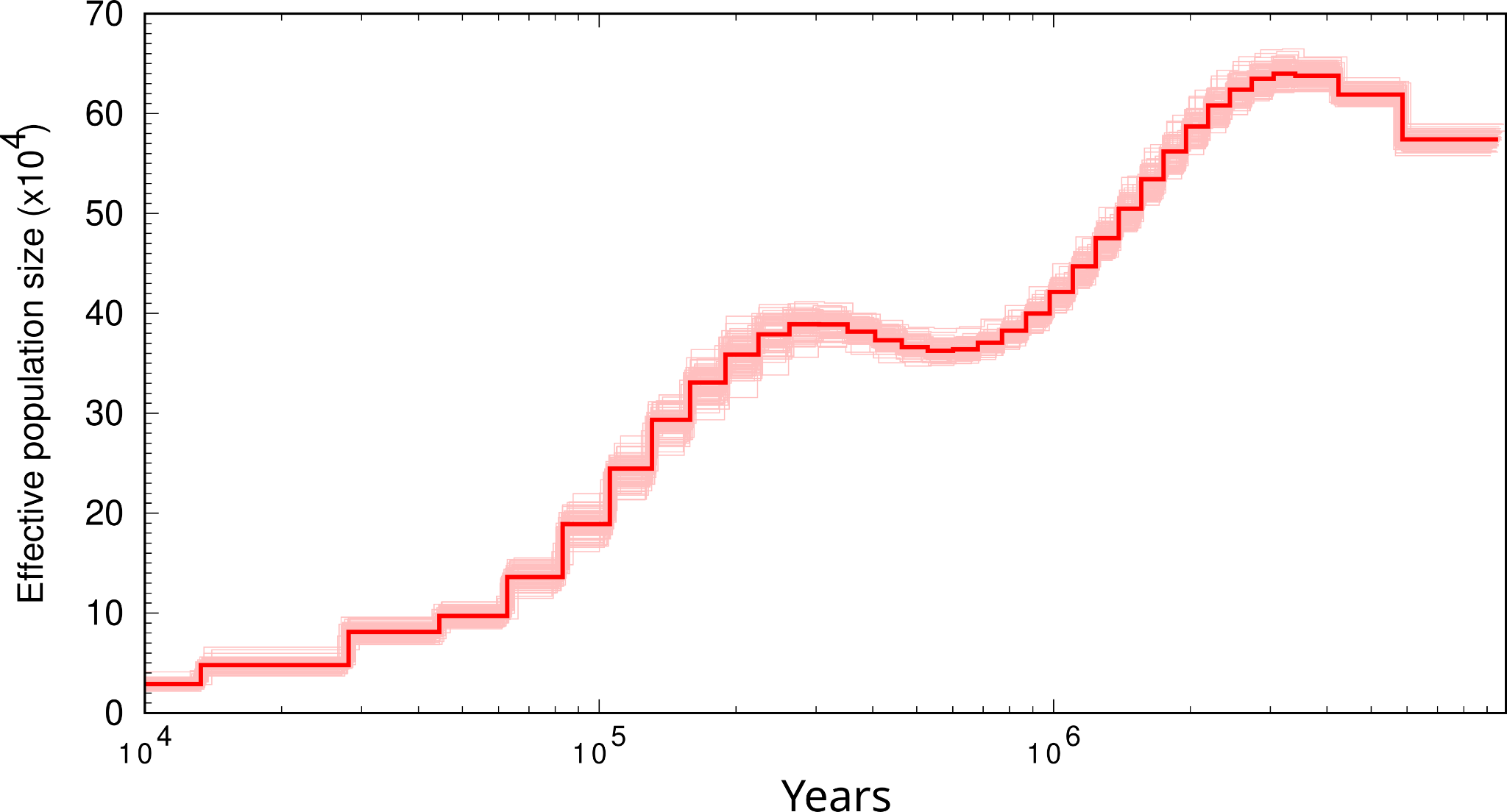

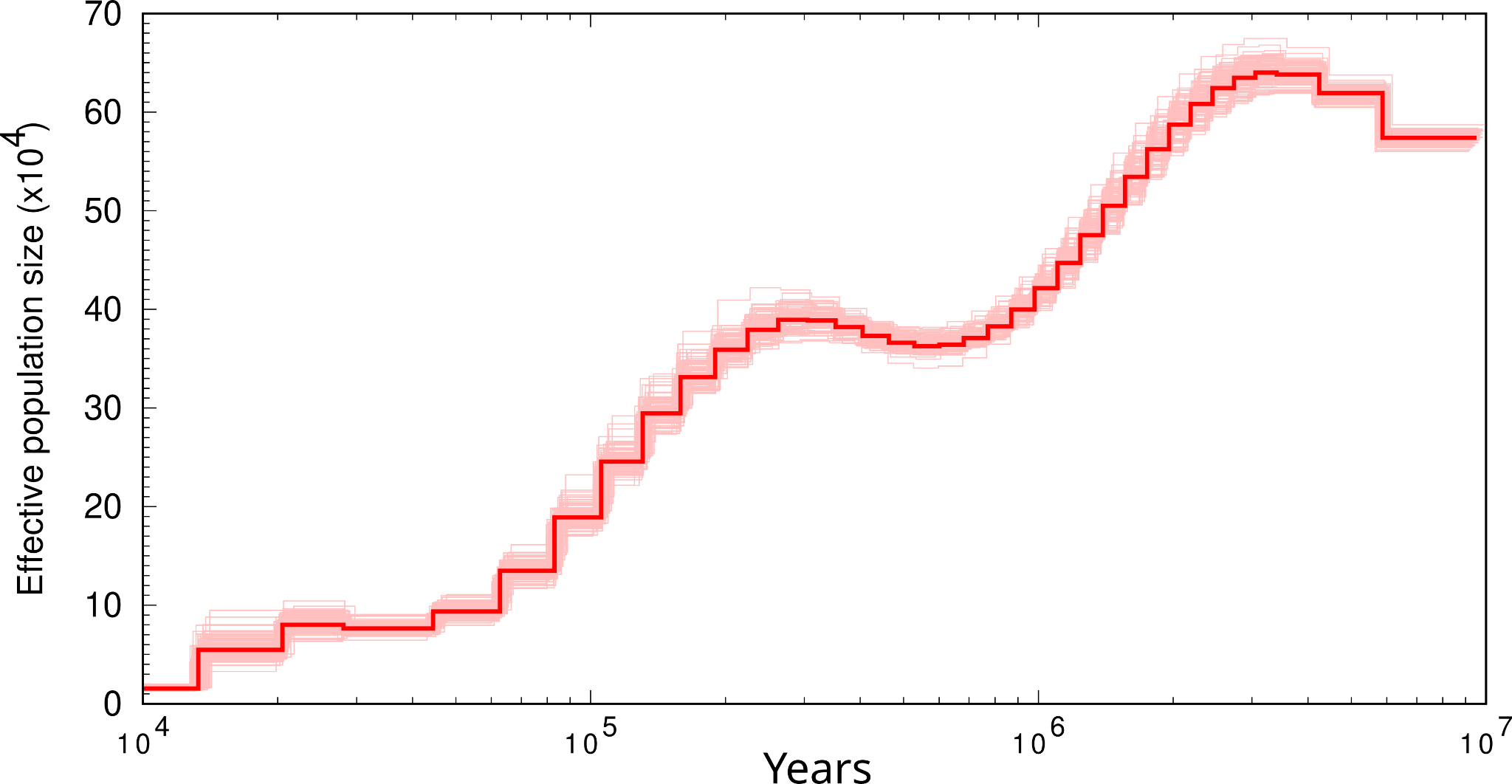

#### *Rhagologus leucostigma*

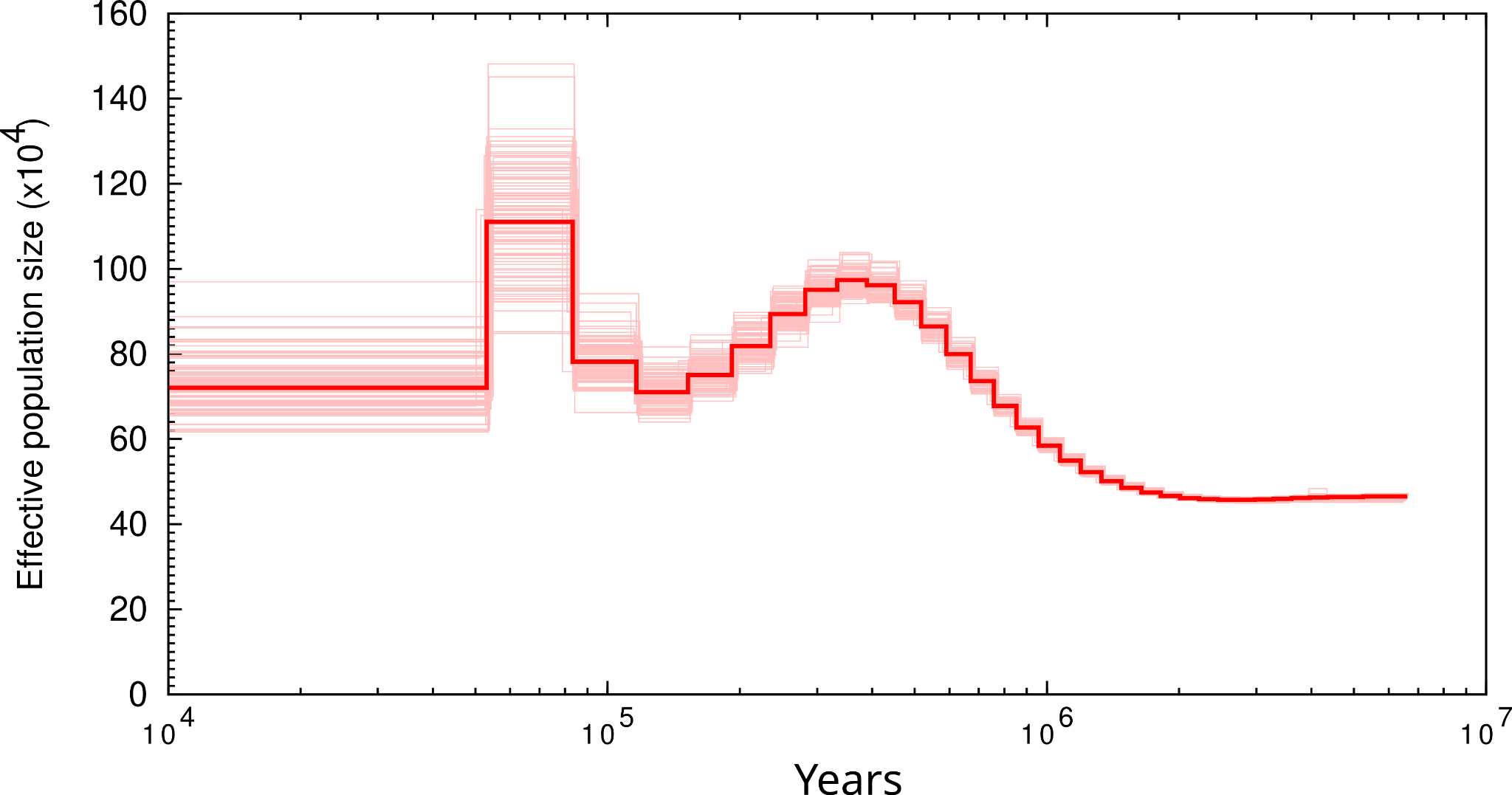

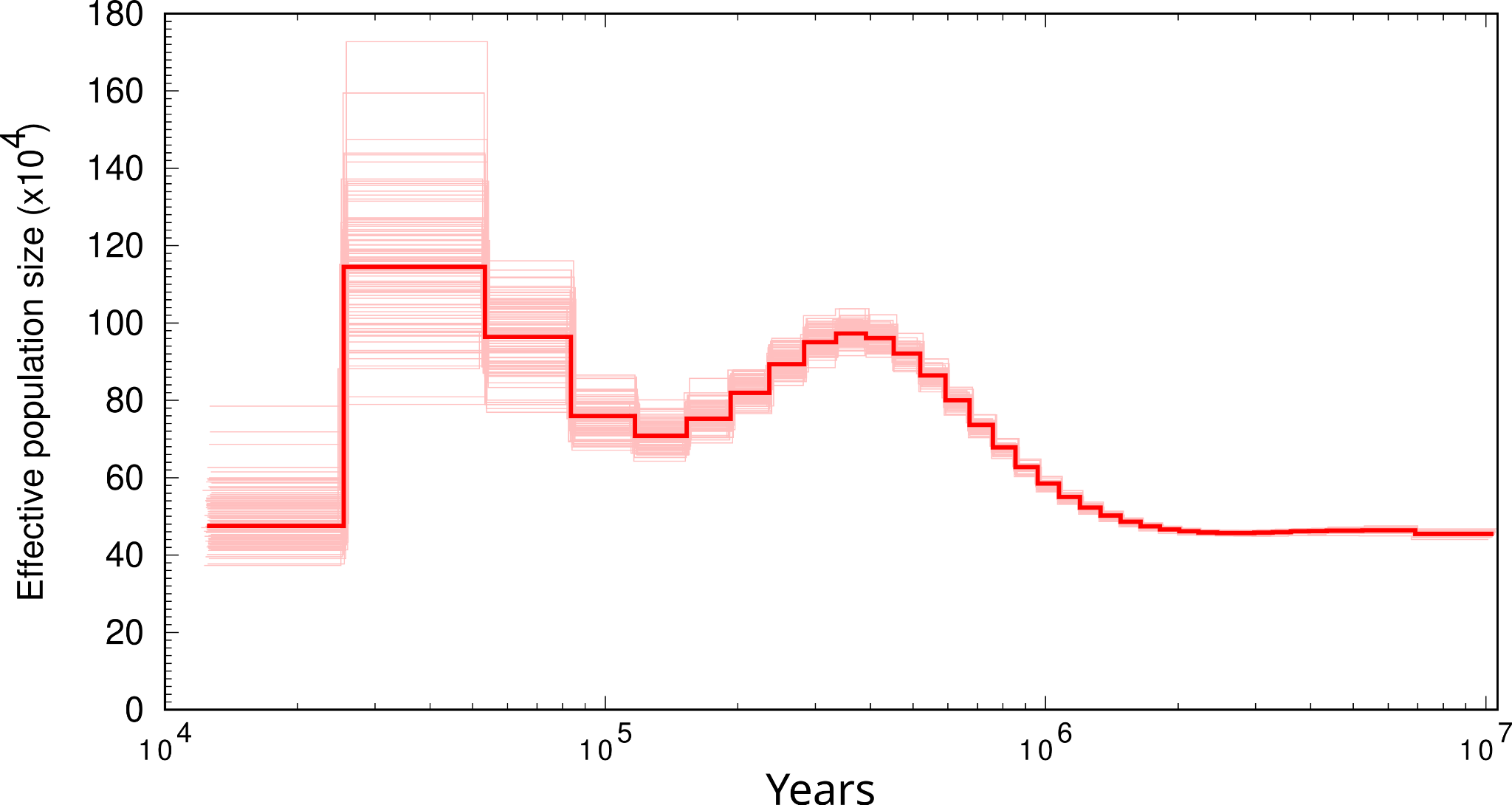

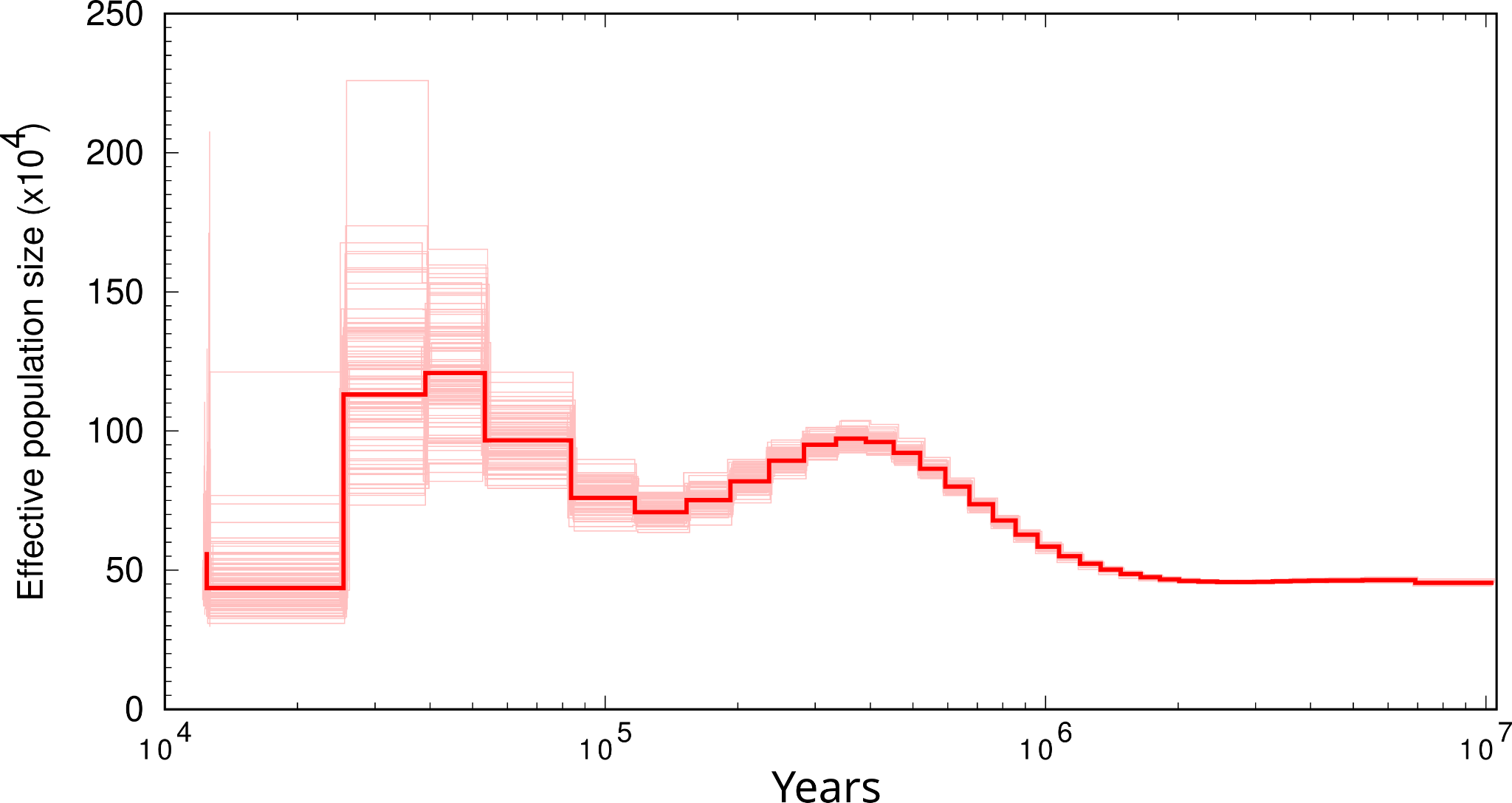

#### *Rhynochetos jubatus*

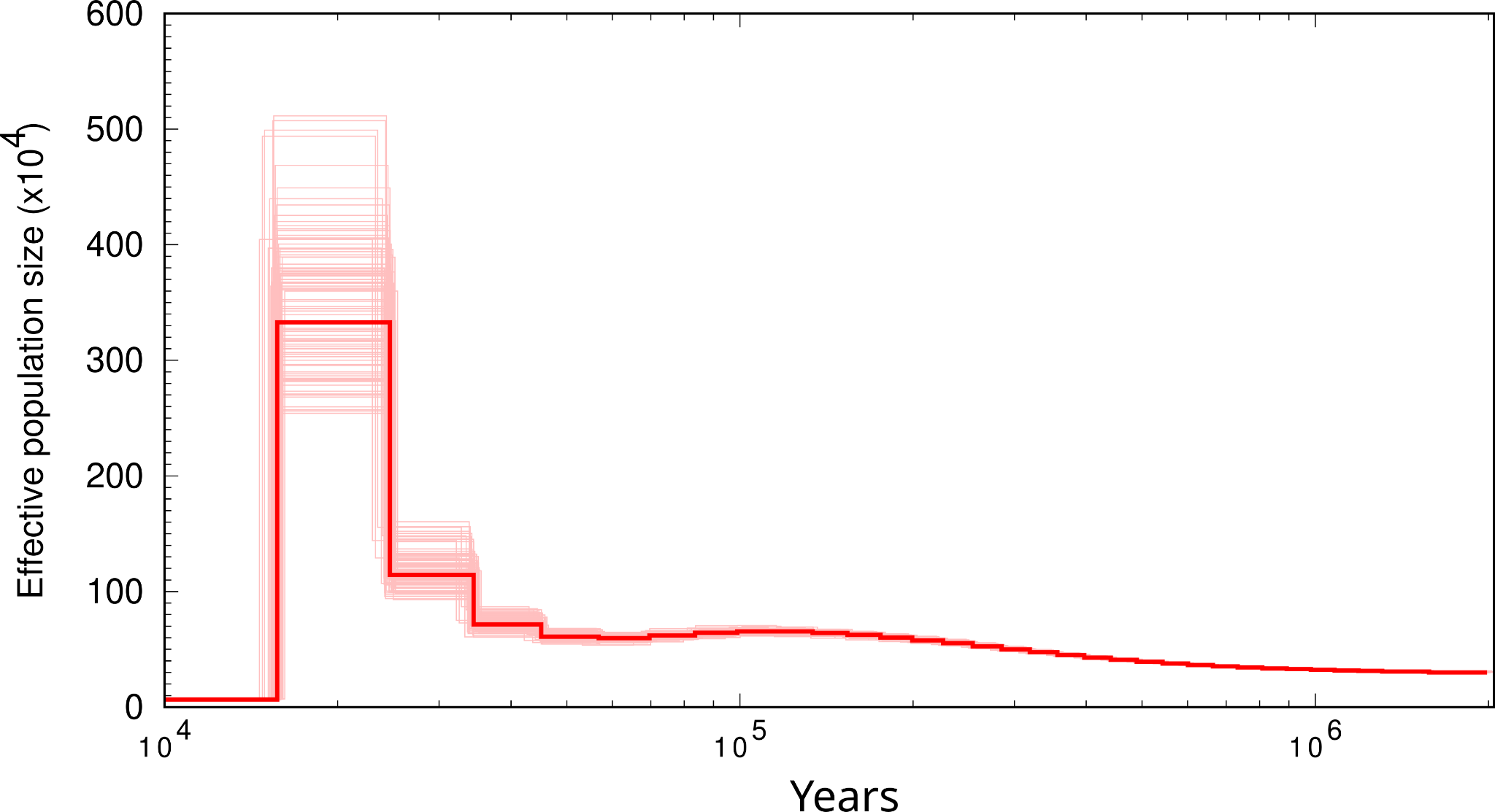

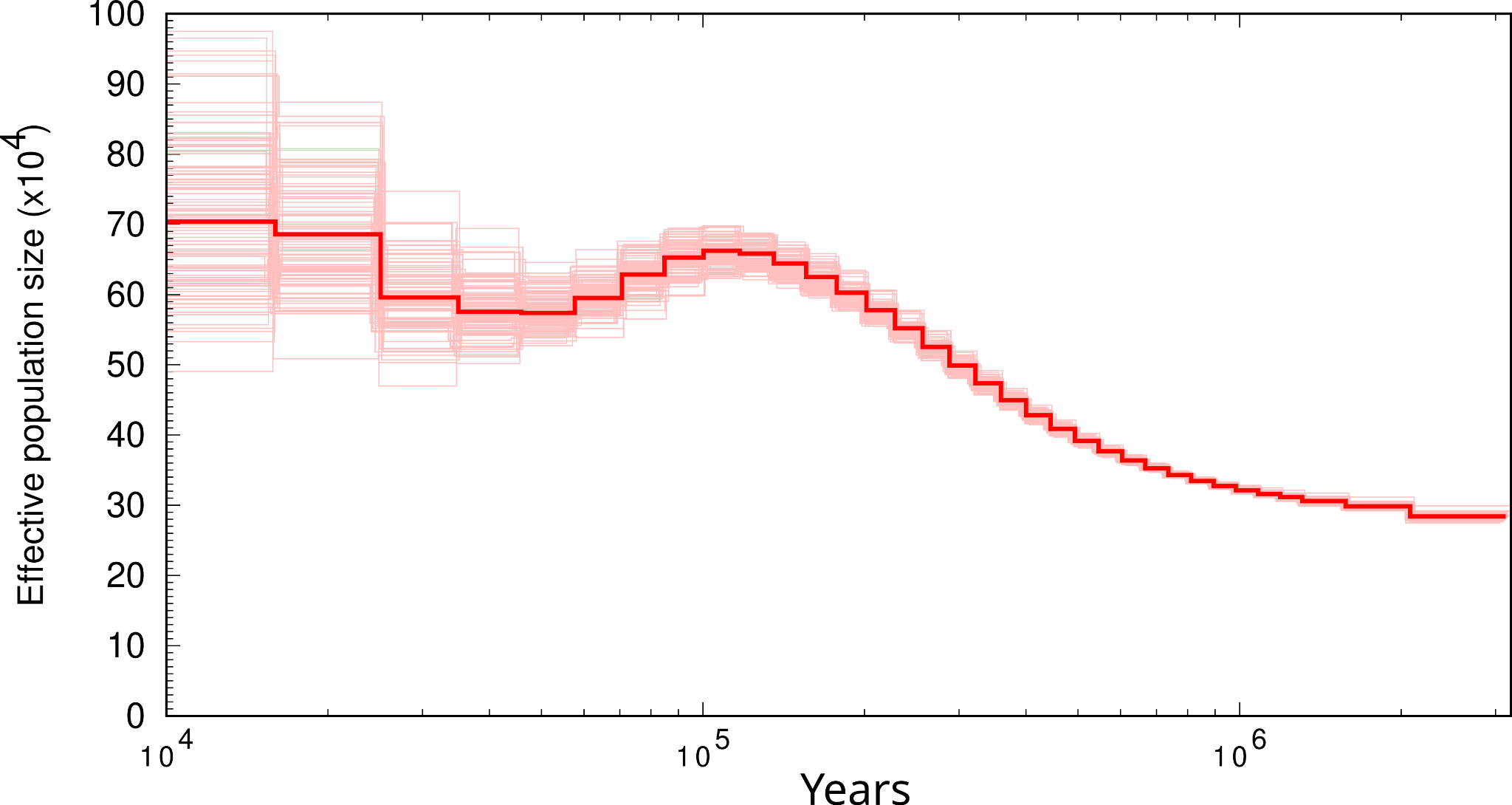

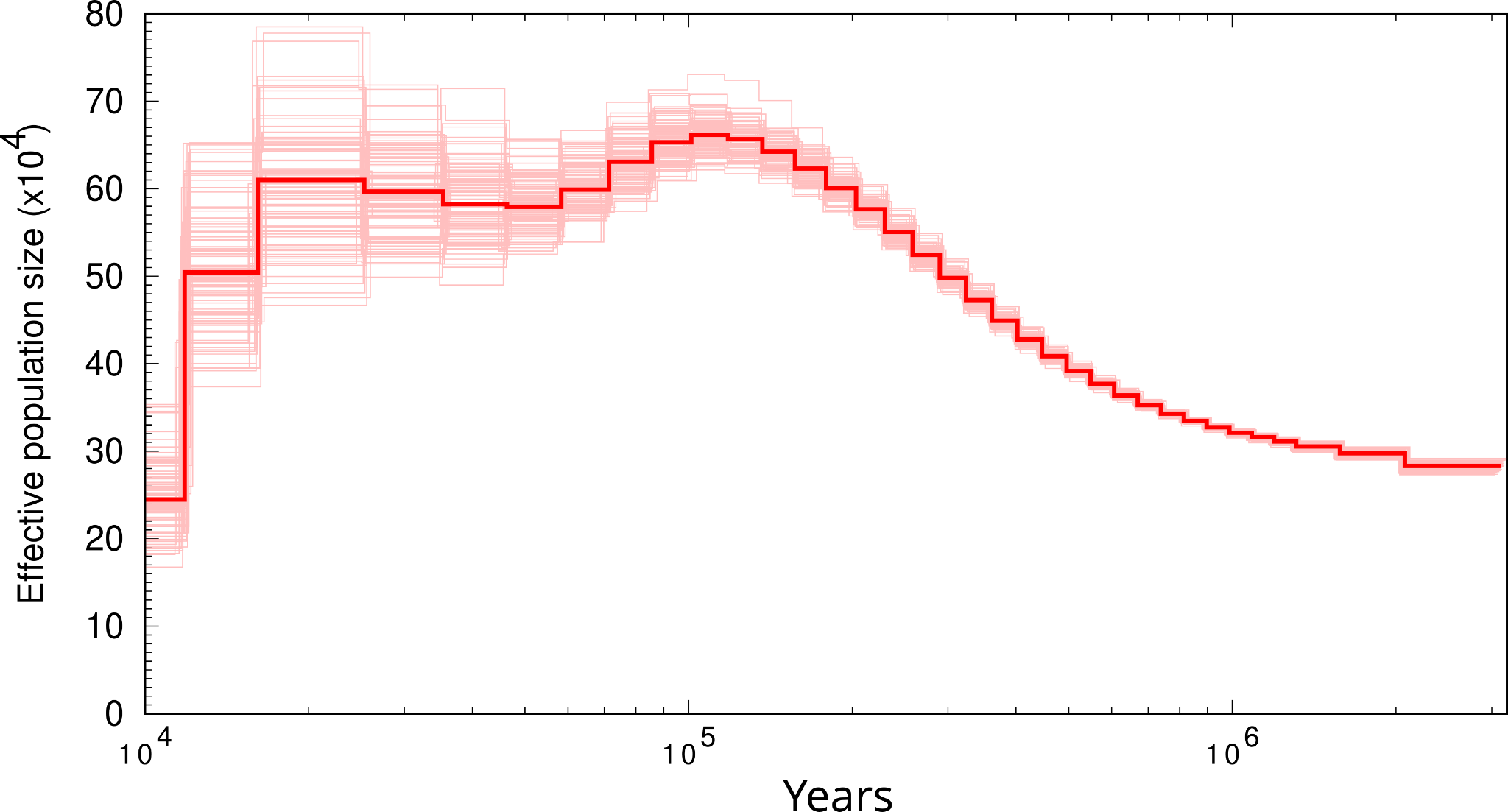

#### *Sterrhoptilus dennistouni*

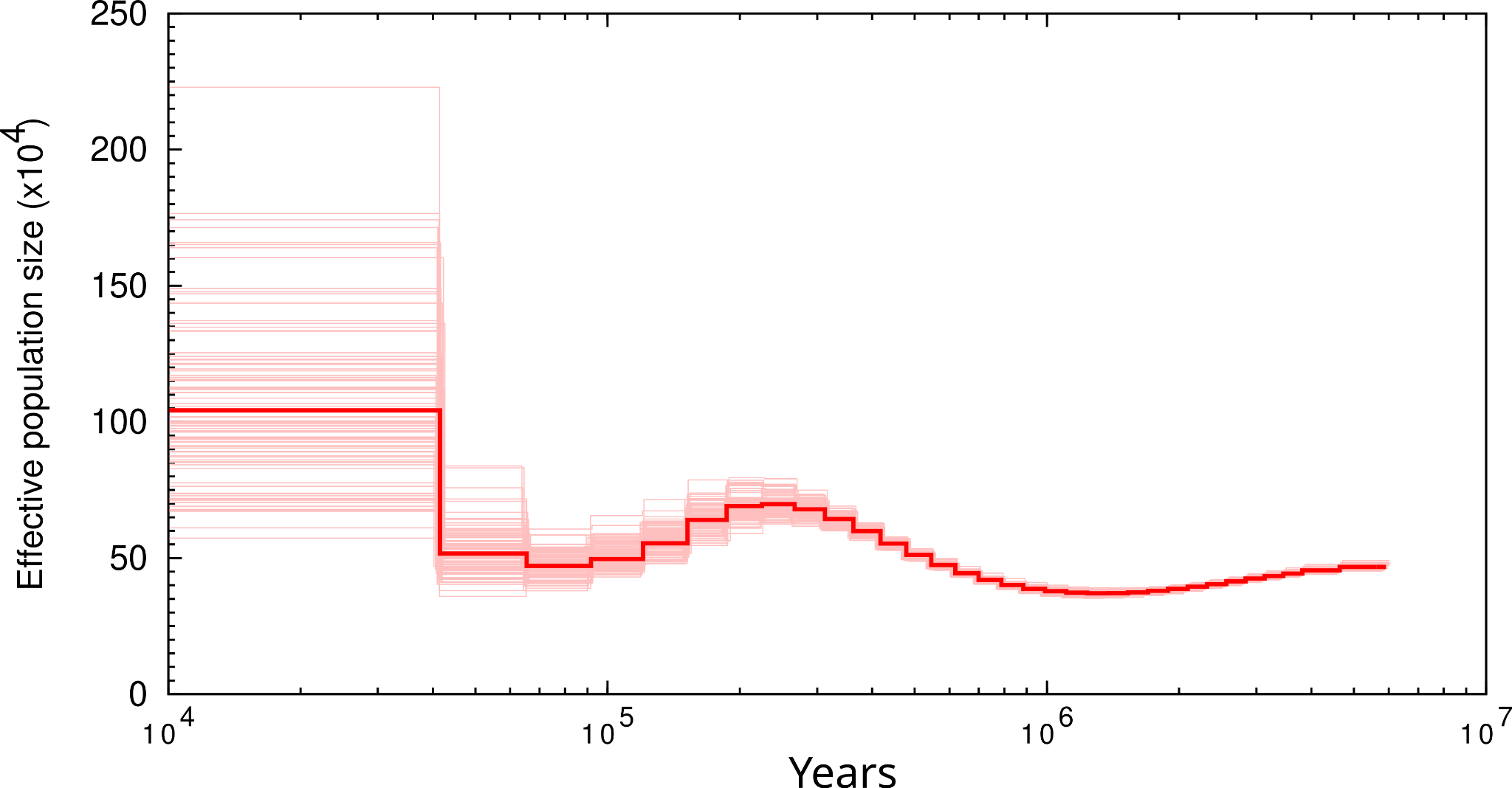

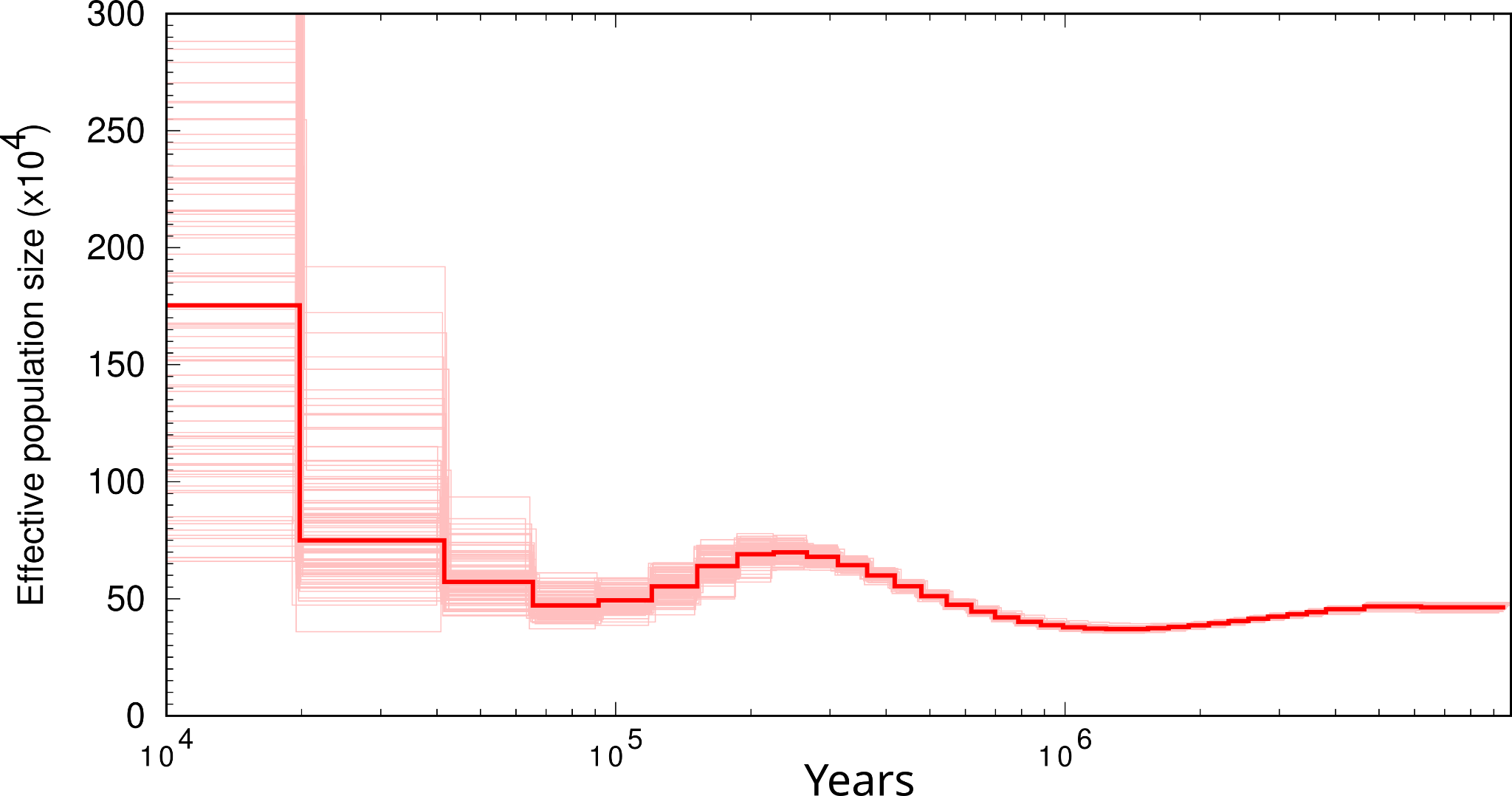

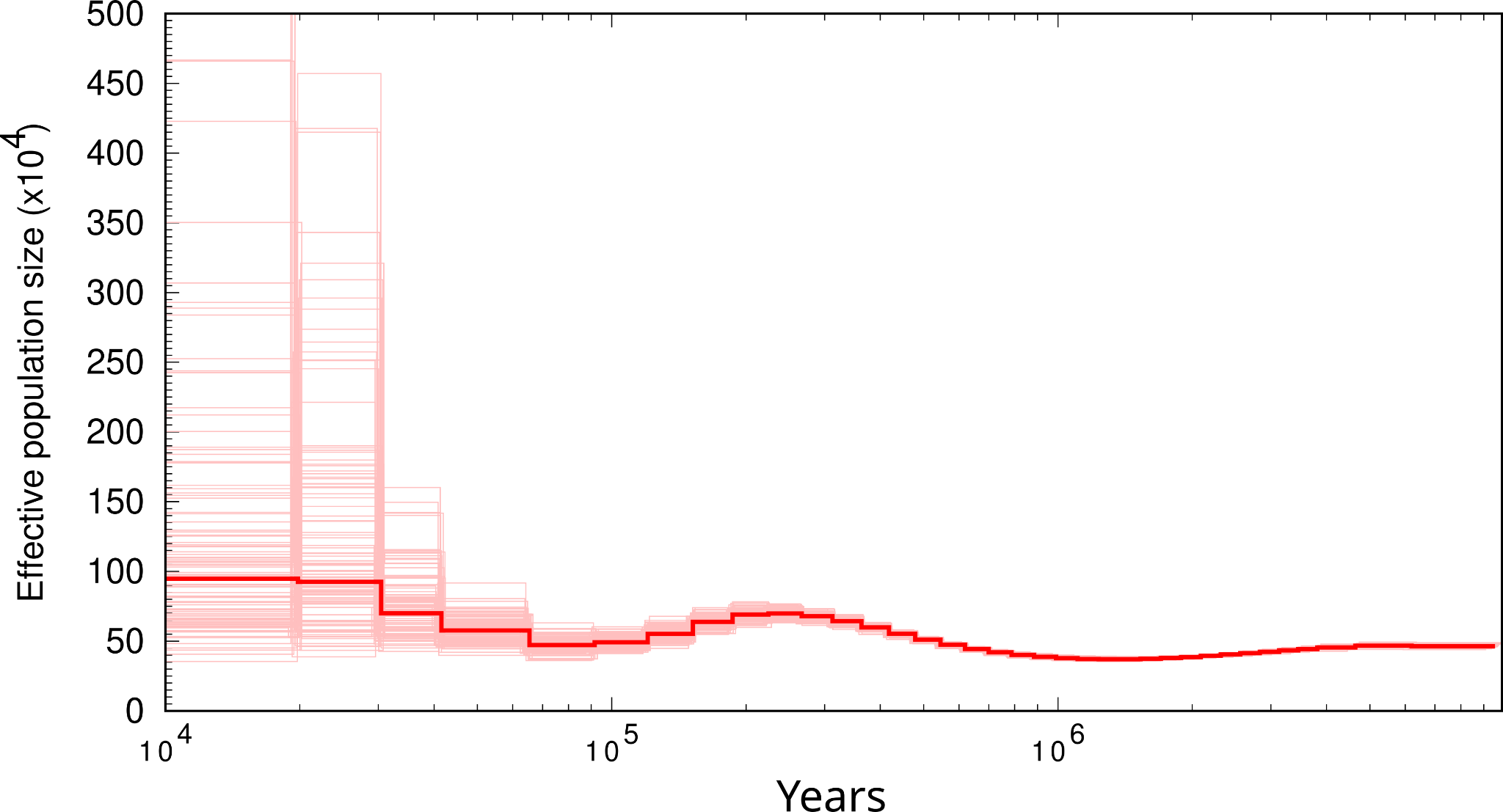

#### *Todus mexicanus*

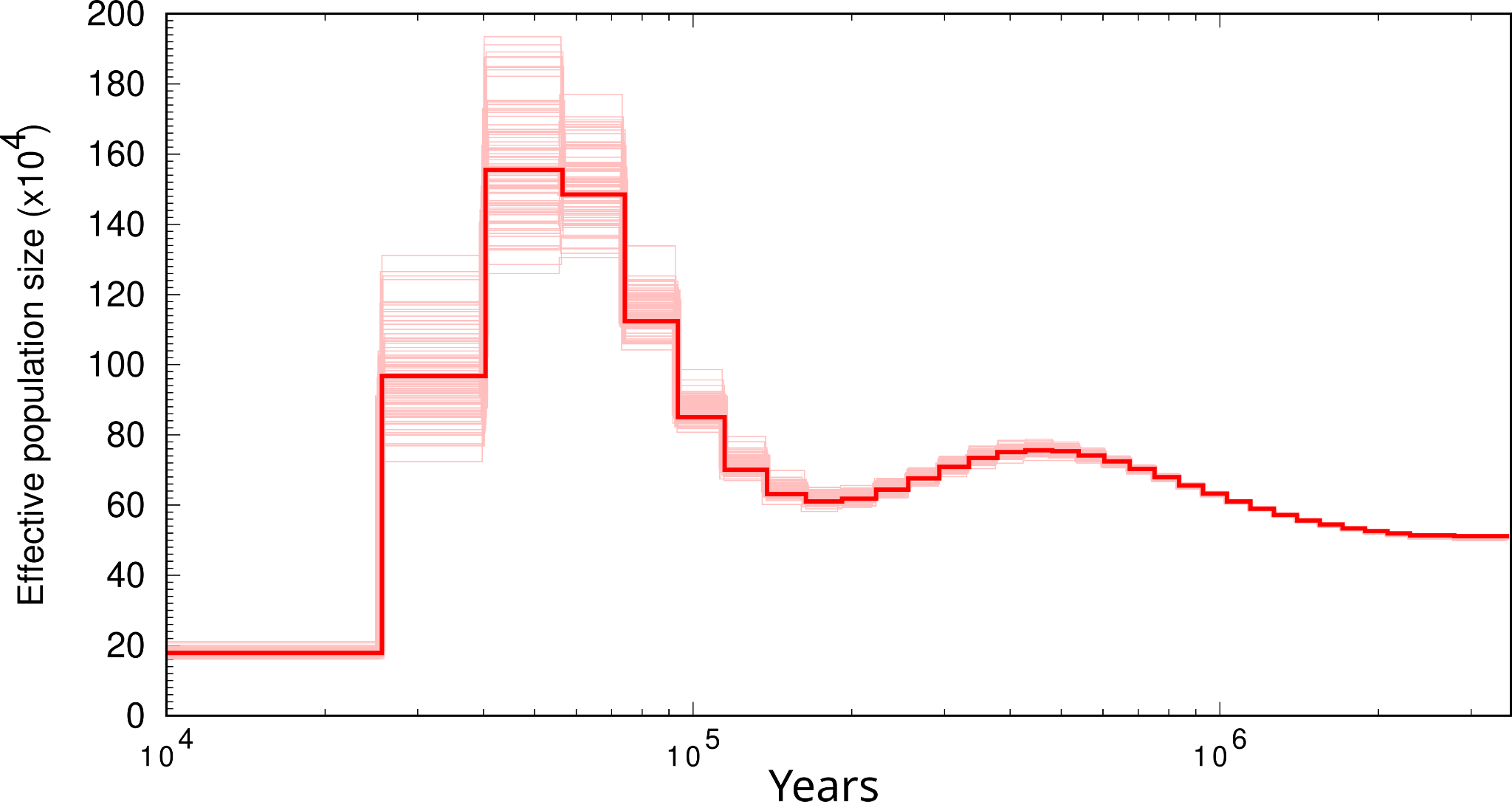

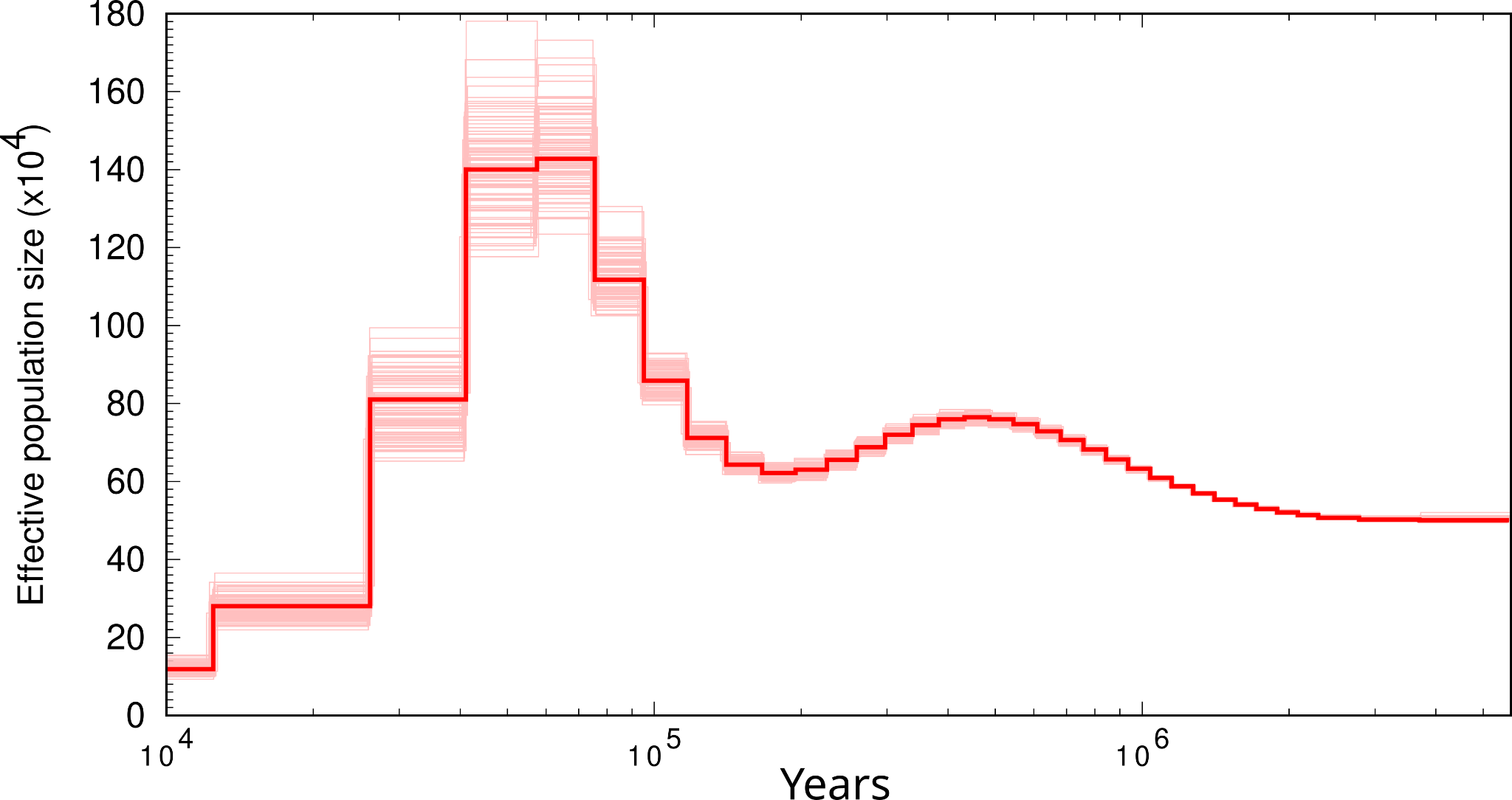

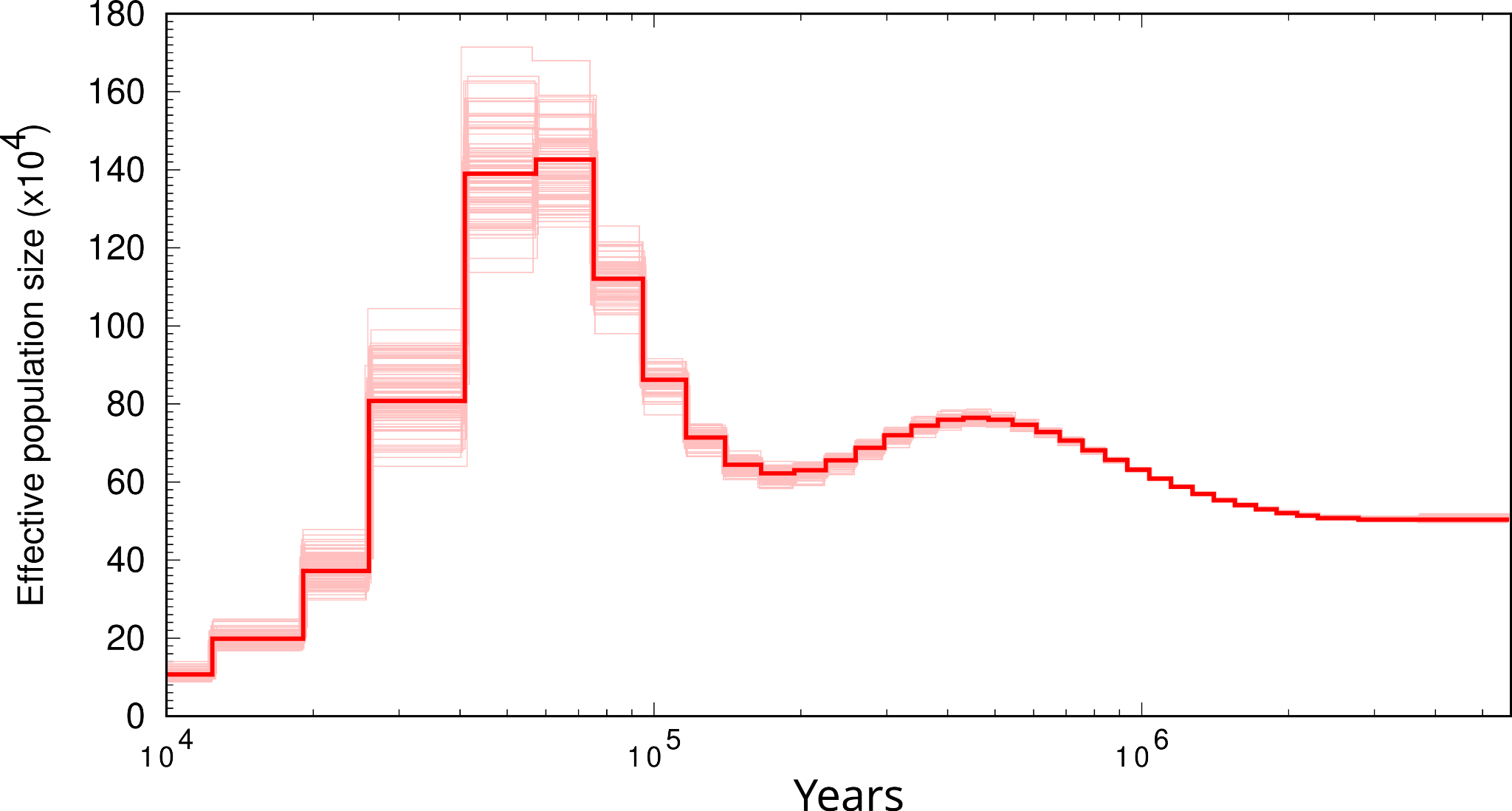

#### *Zosterops hypoxanthus*

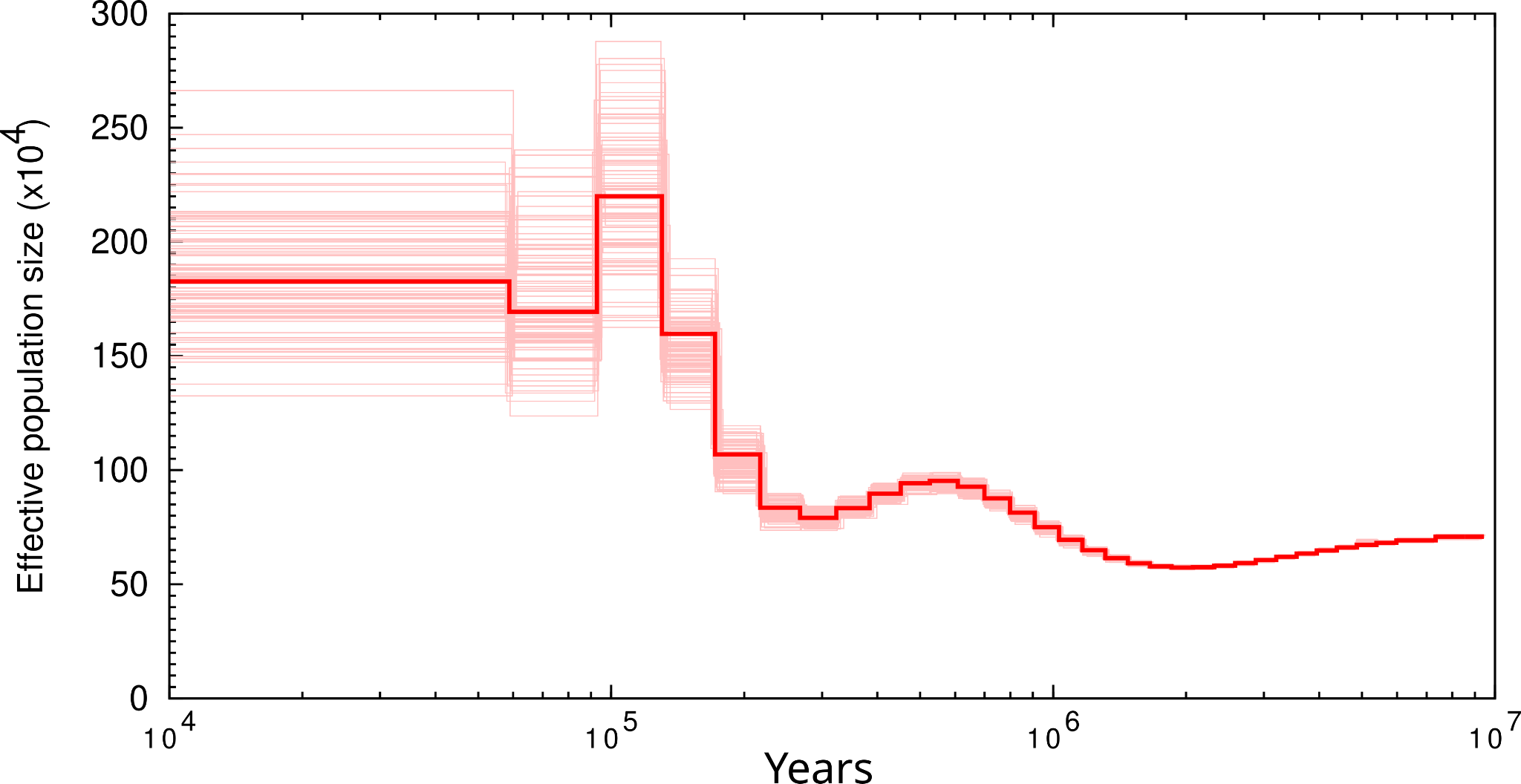

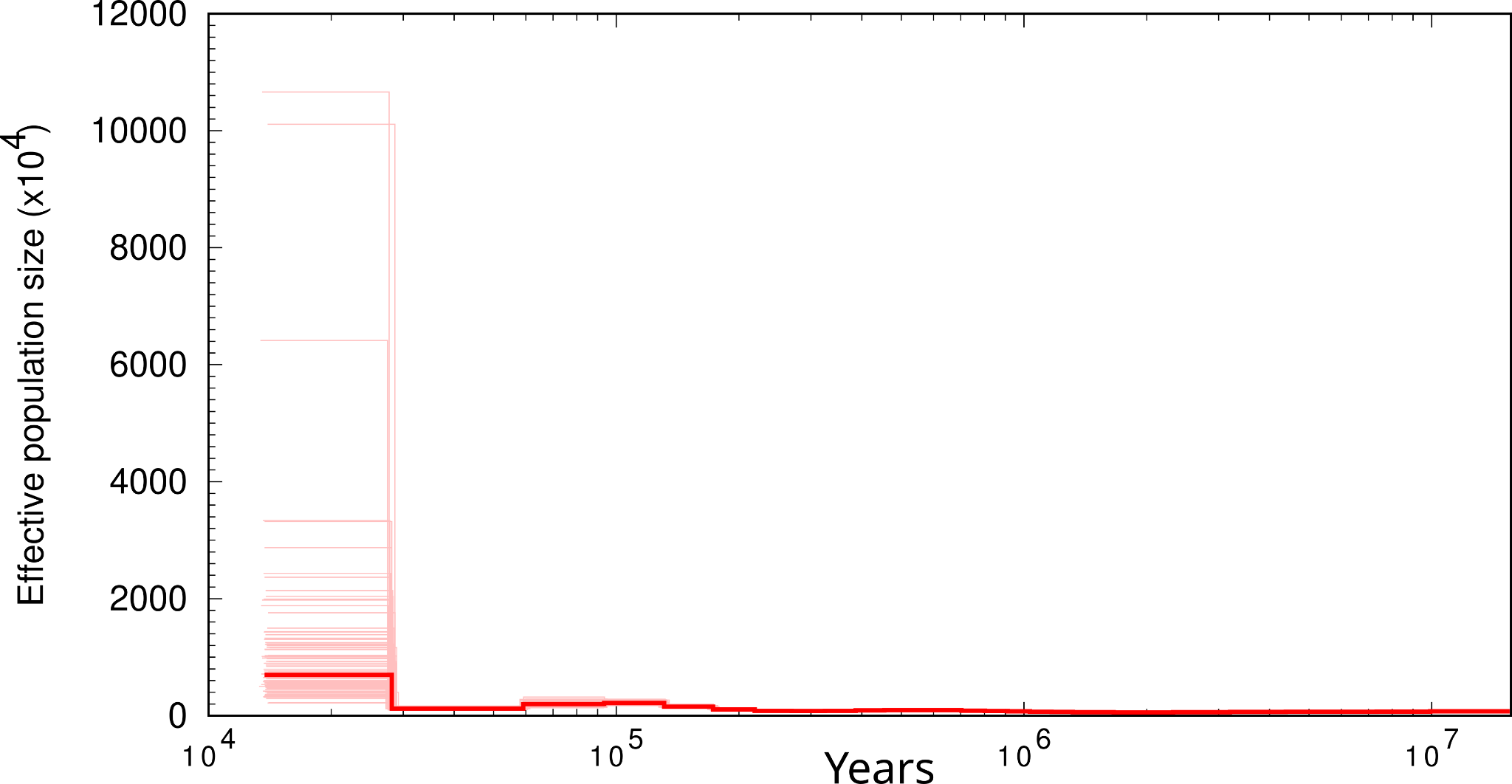

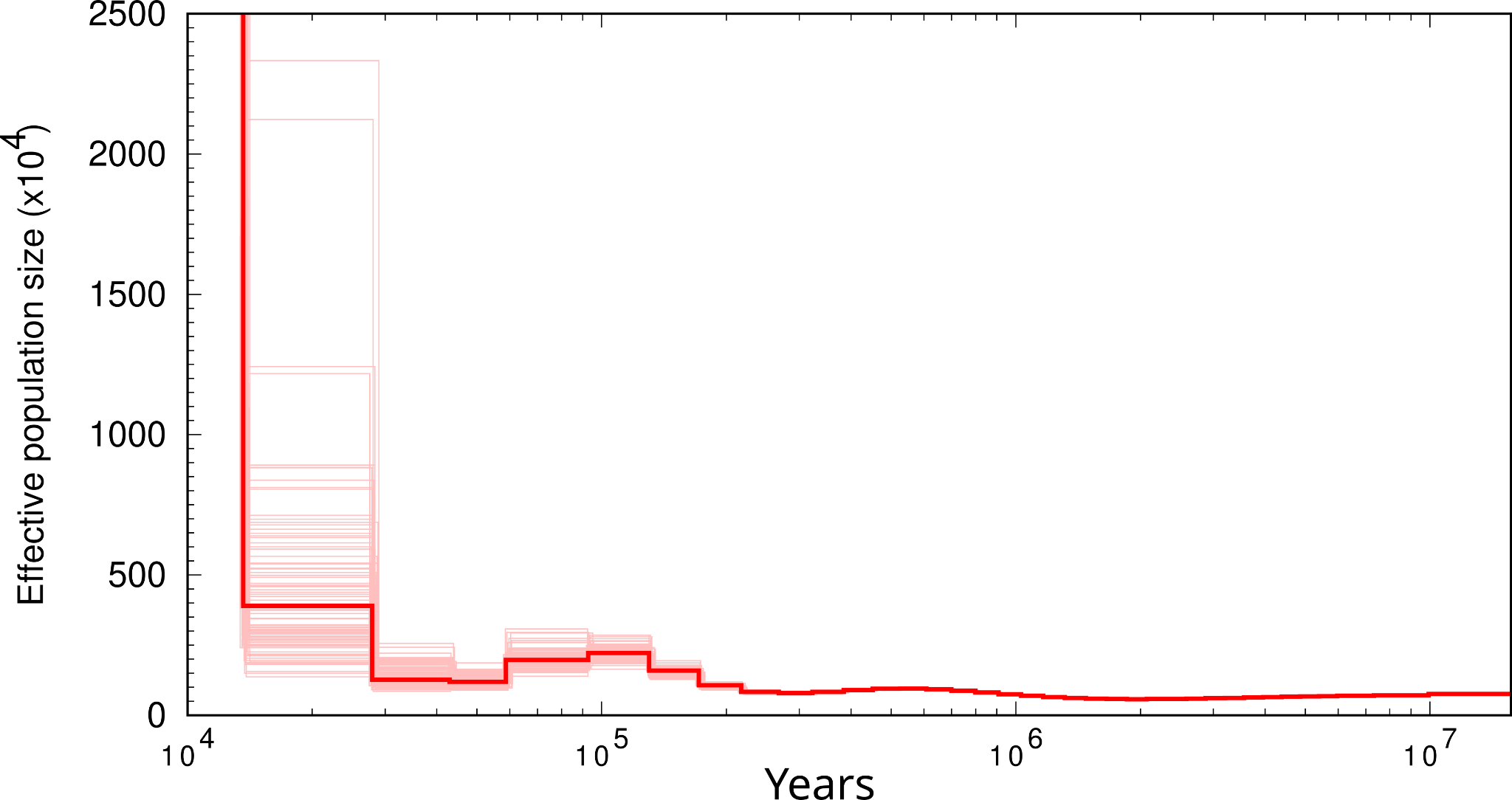

### Figure S2: Ecological Niche Modelling (ENM) plots of all the reconstructed habitat ranges. LIG = Last Inter Glacial, LGM = Last Glacial Maximum, MDH = Mid Holocene, CUR = Current. The continuous heatmap represents the probability of occurrence of the species and red points are known occurrences from the Global Biodiversity Information System (GBIF). Stable = Unchanged habitat range. Increase = Habitat gained. Decrease = Habitat lost.

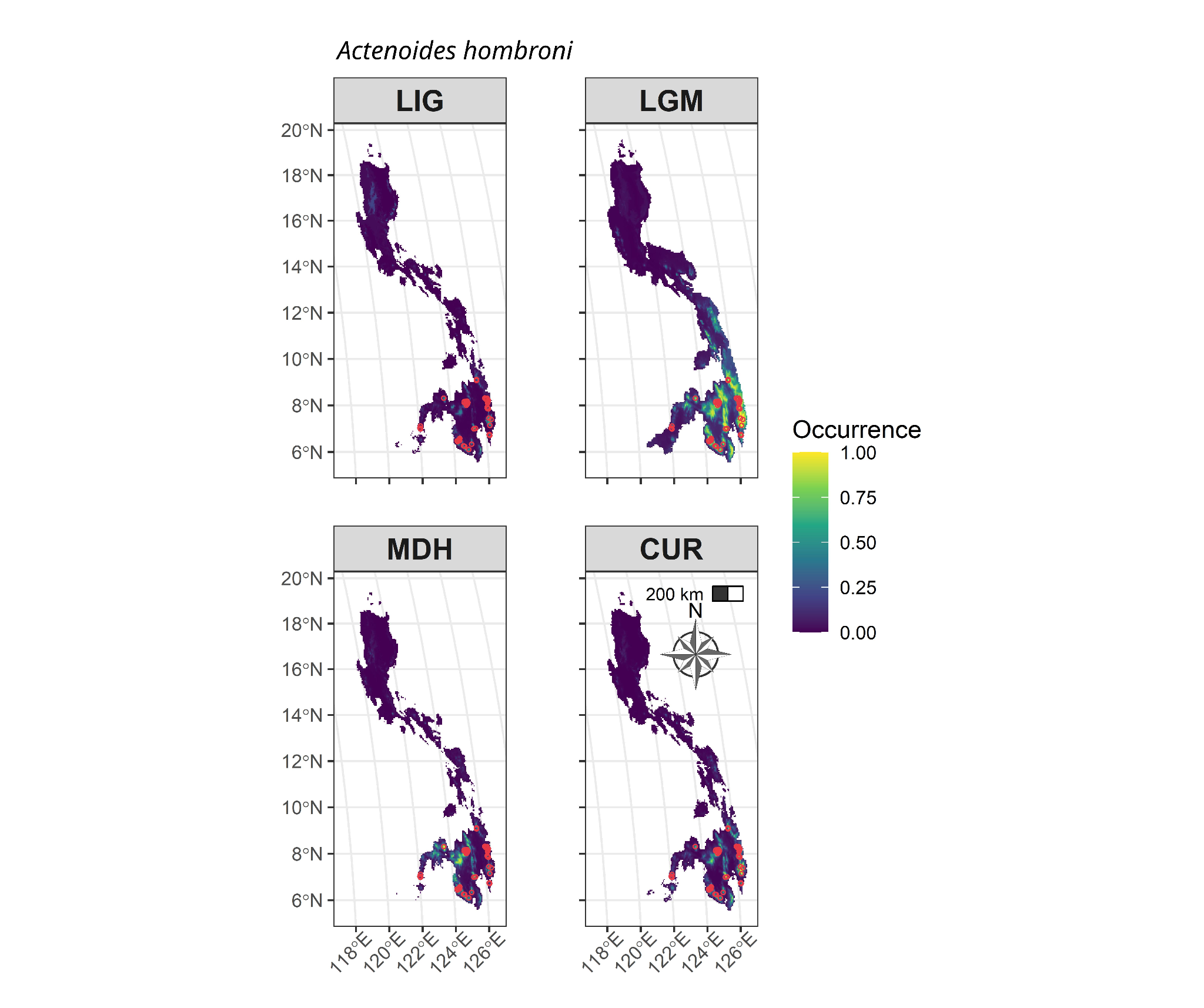

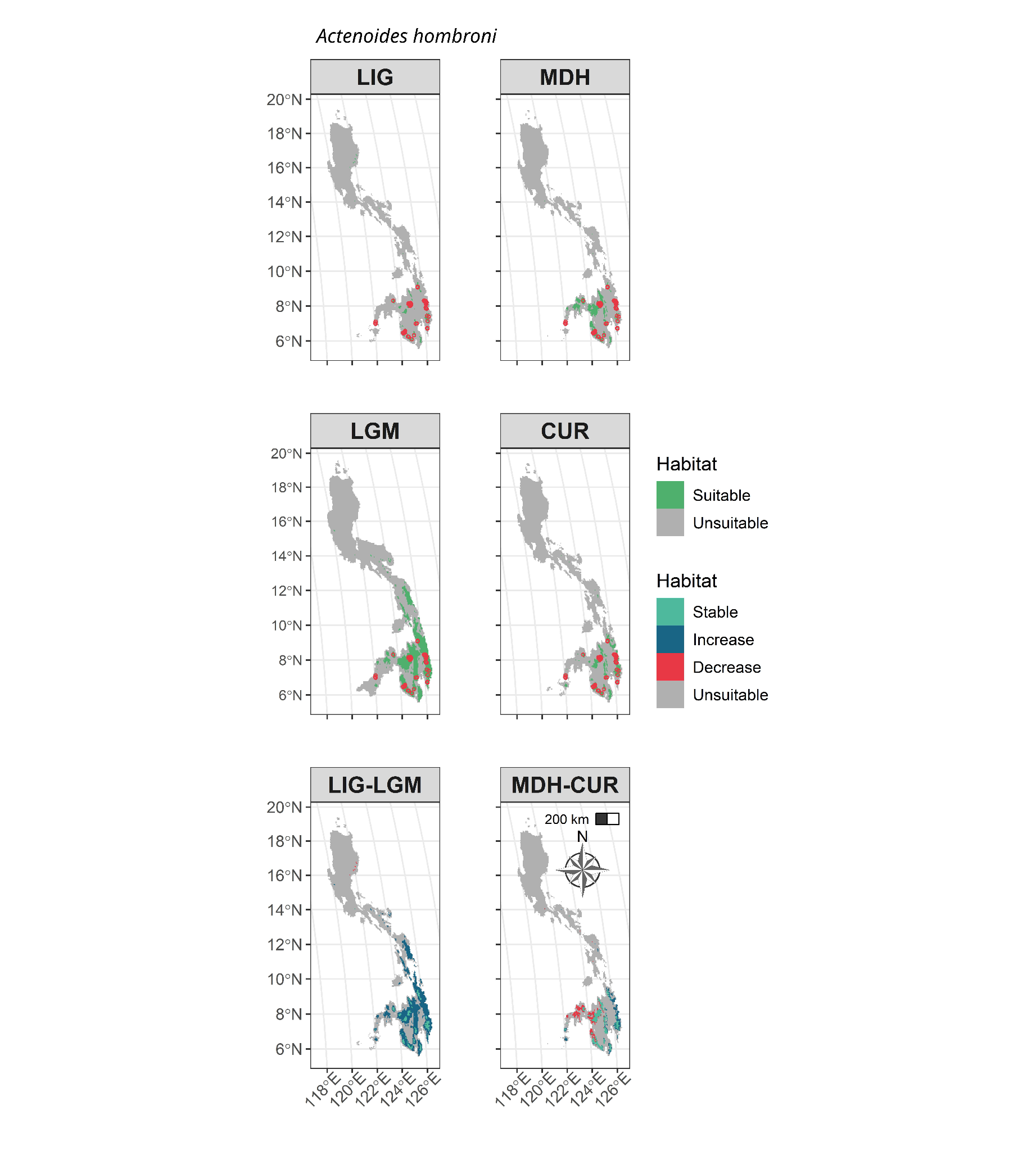

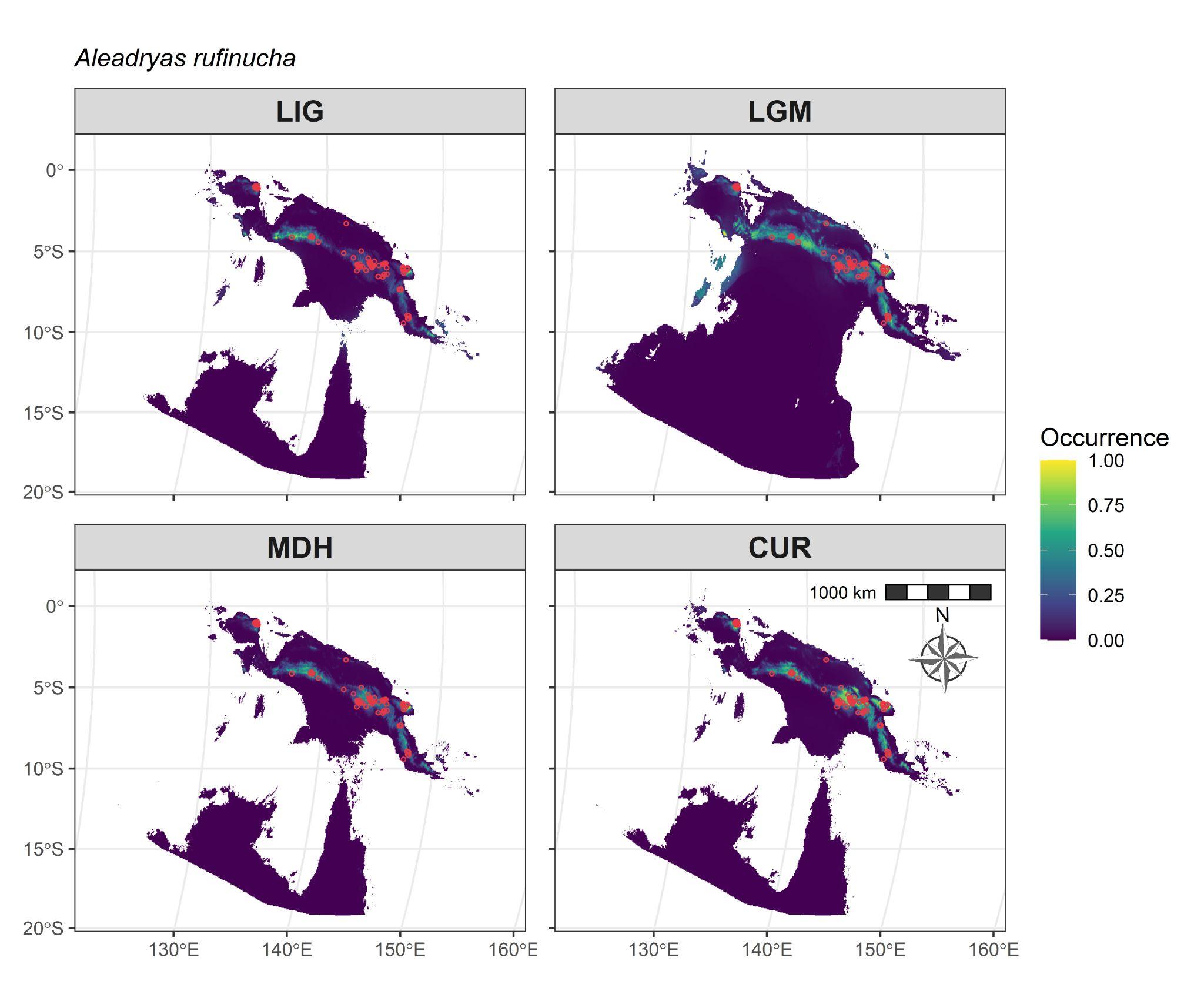

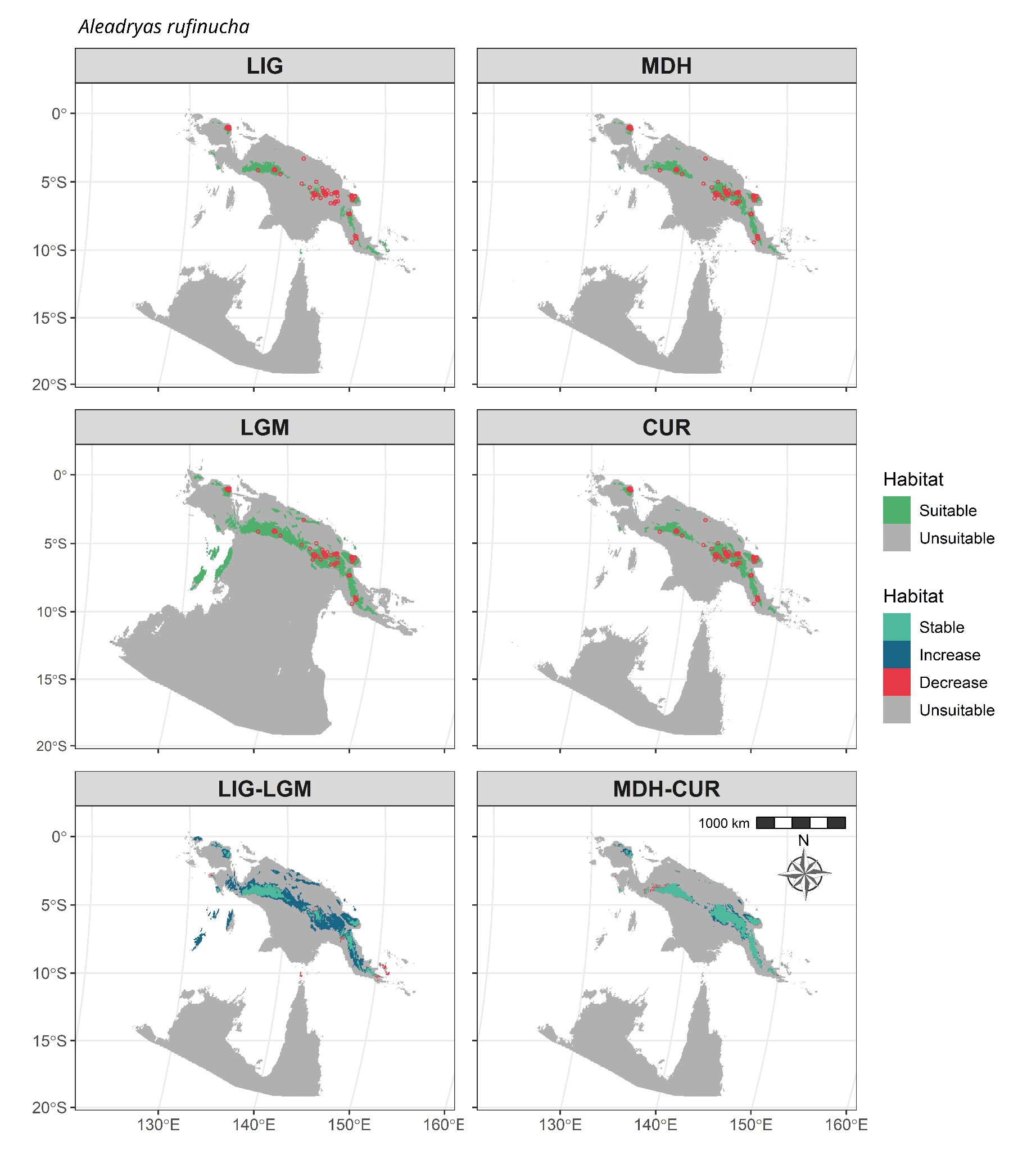

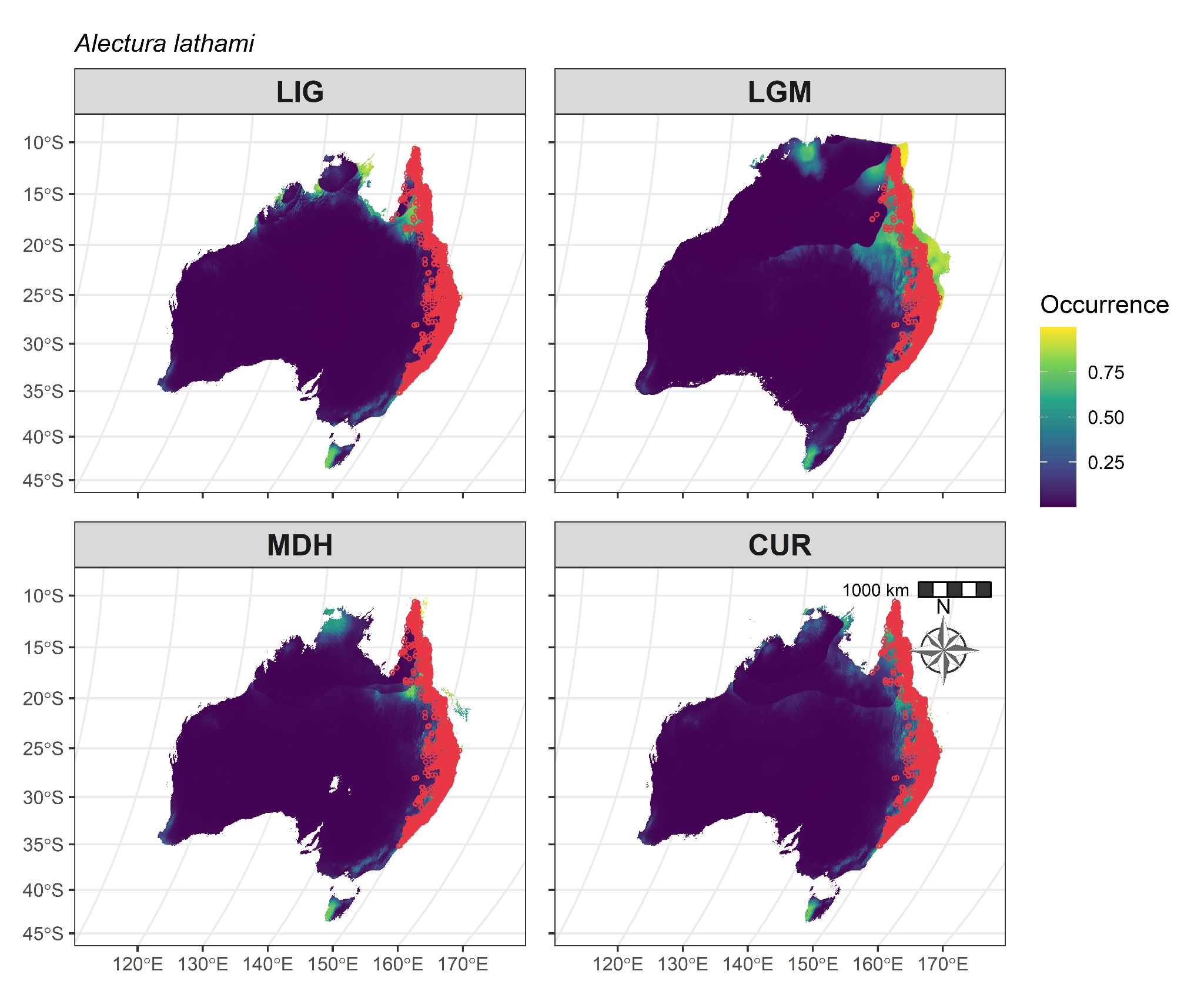

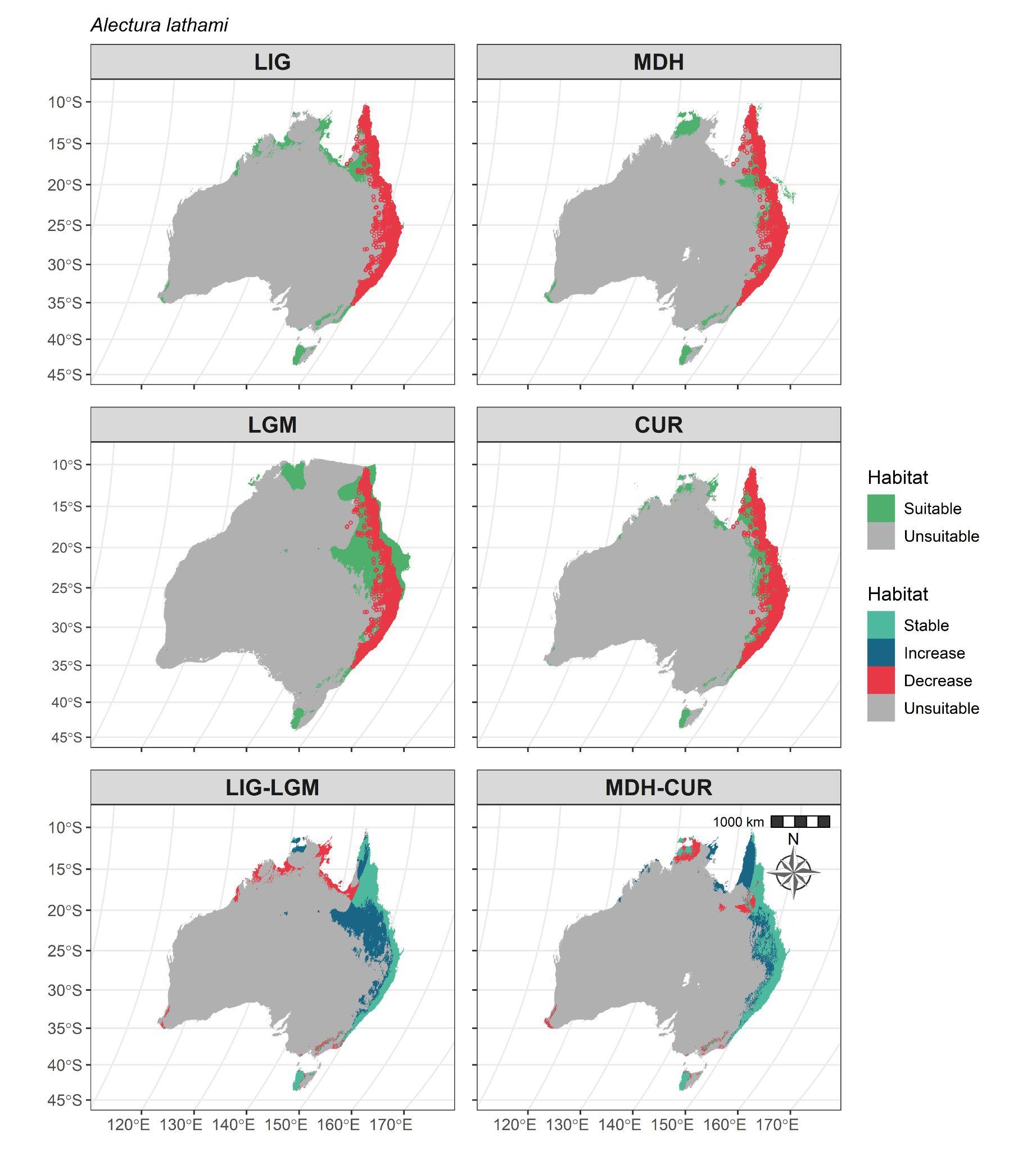

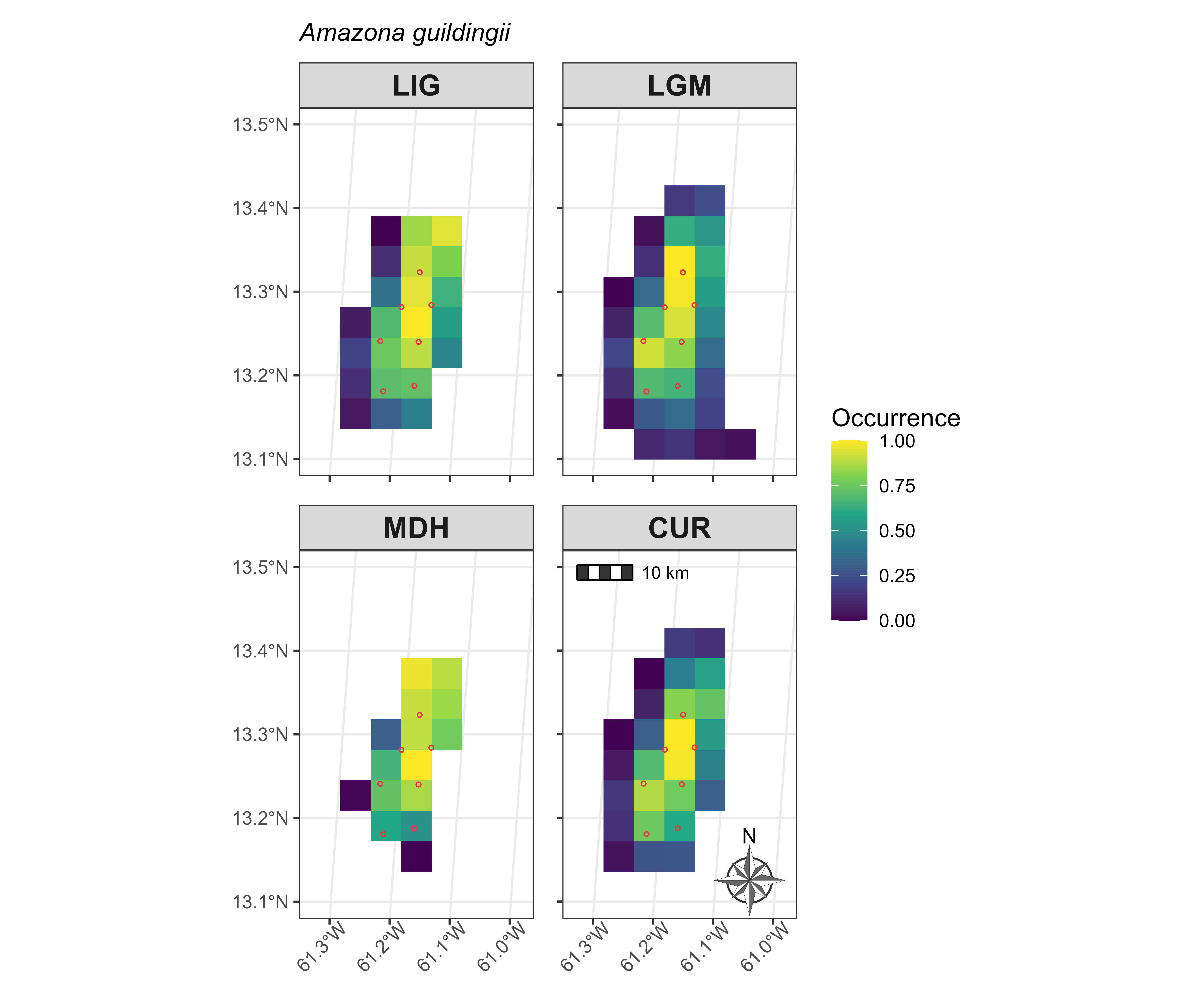

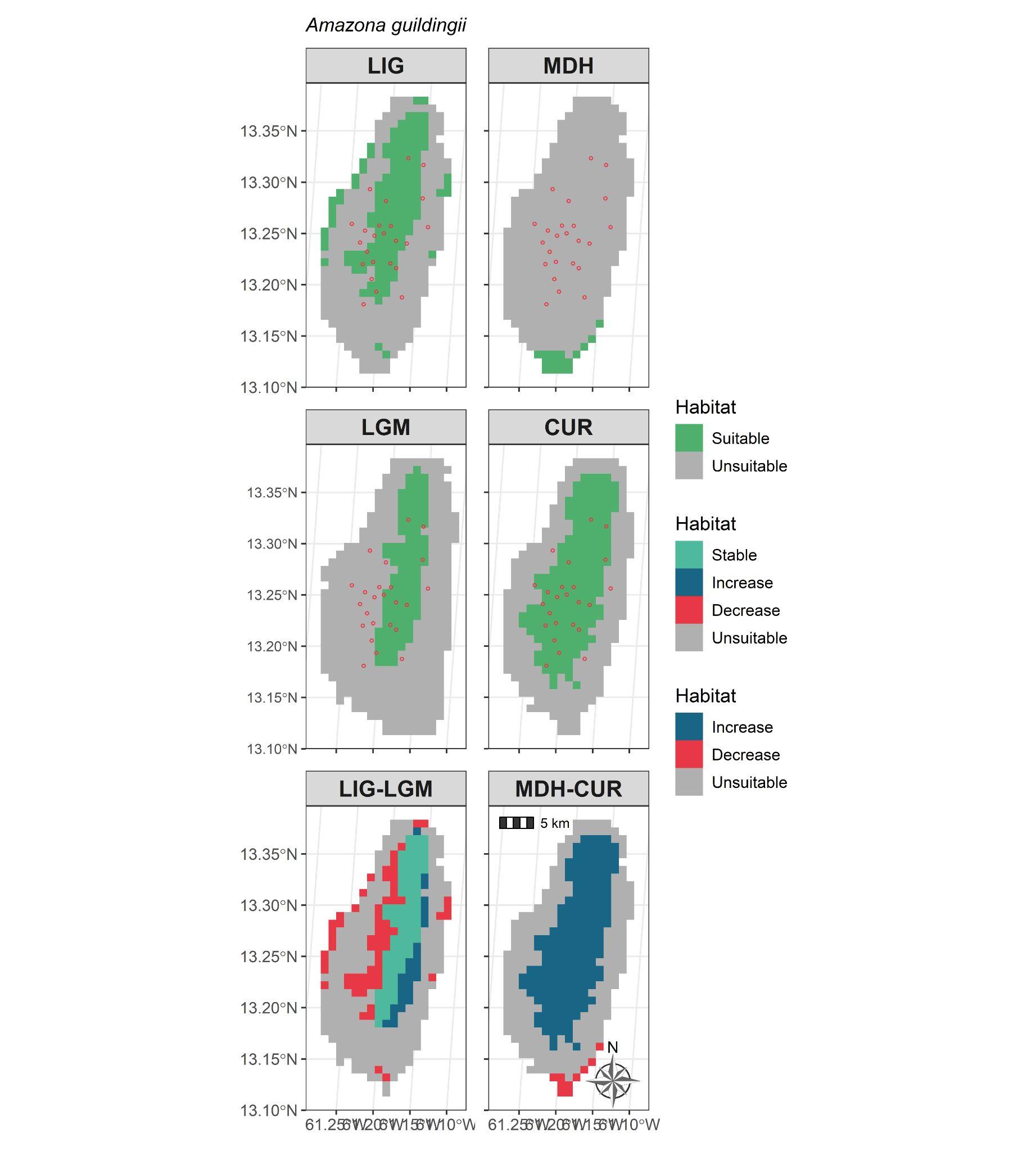

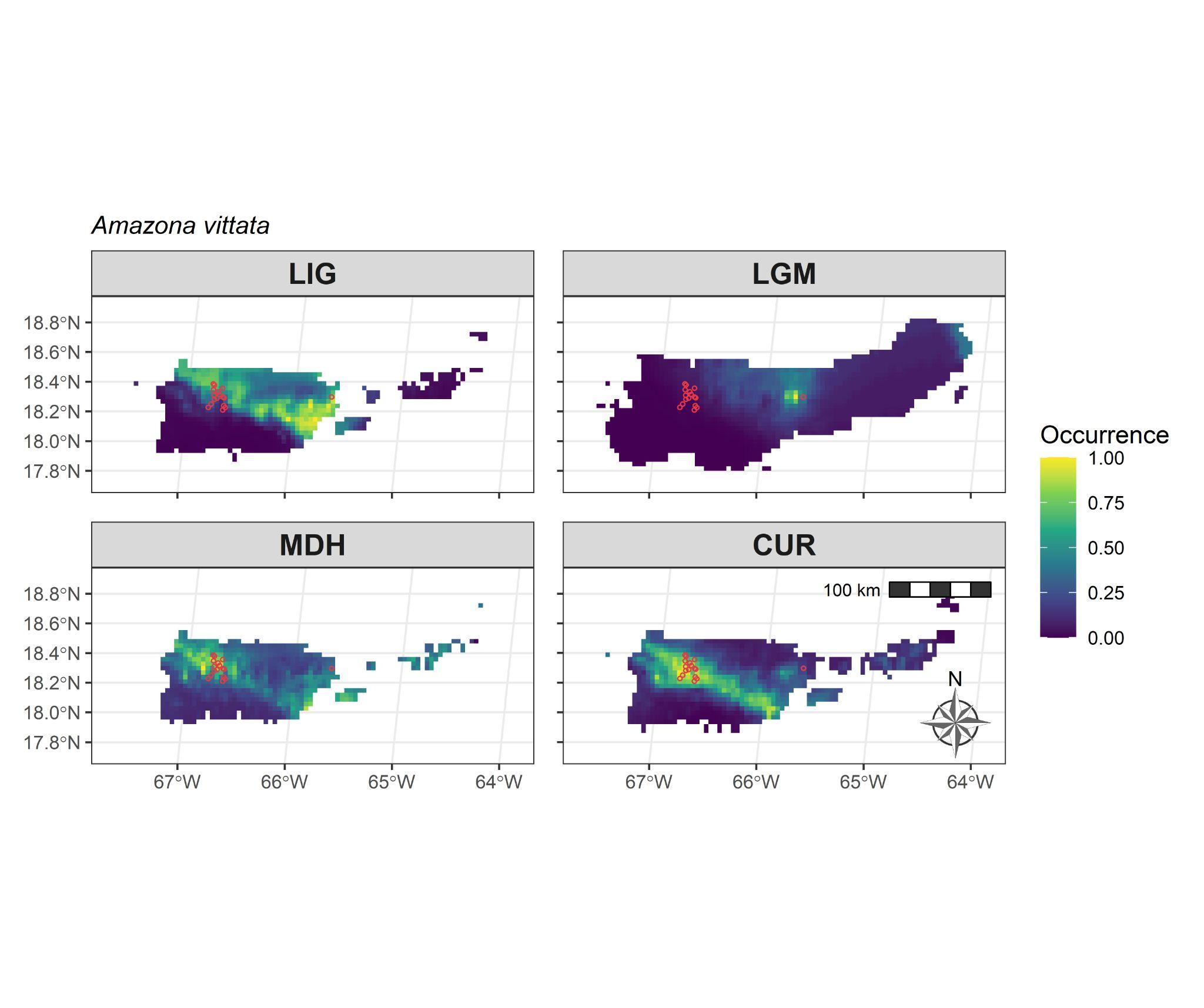

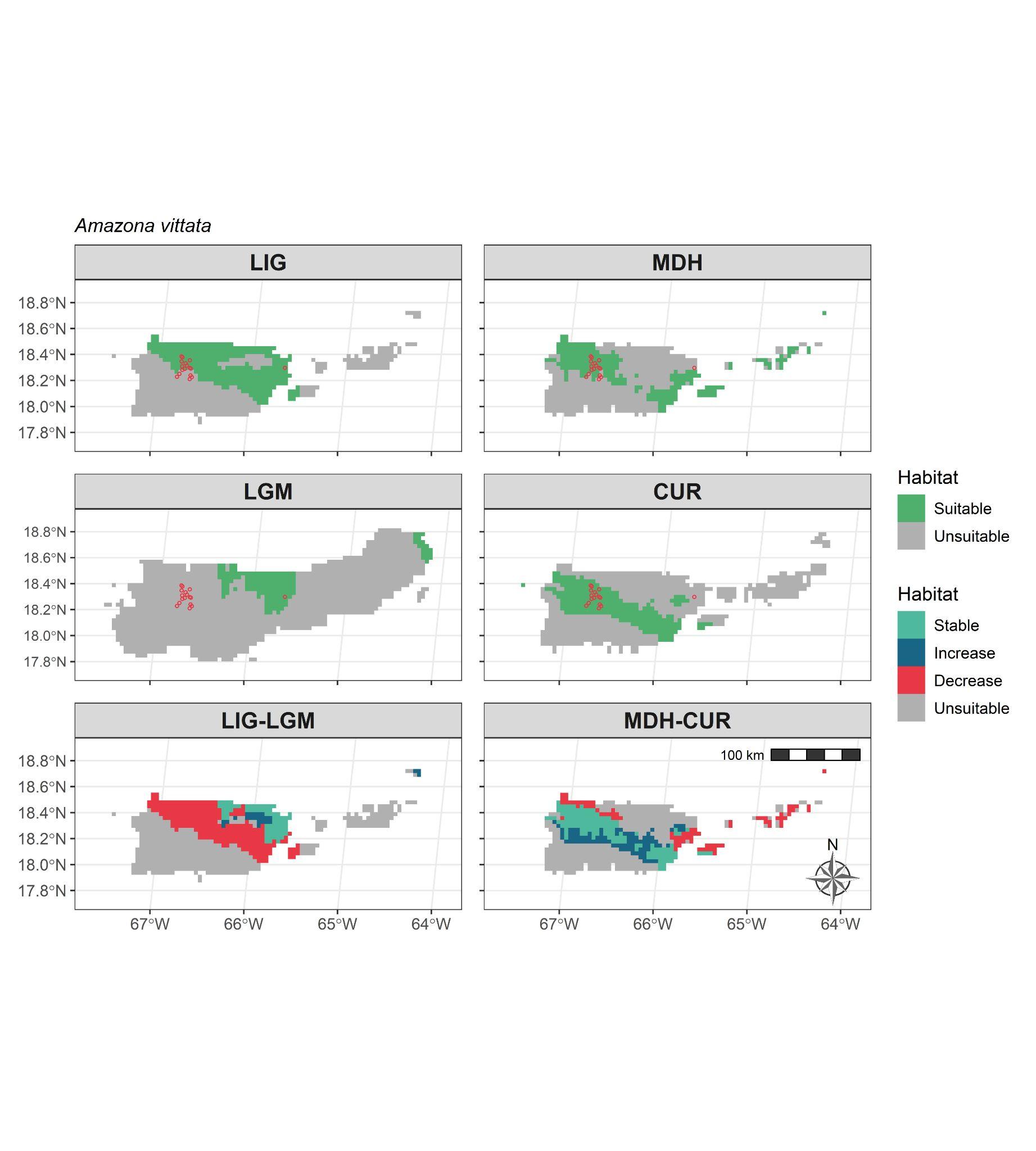

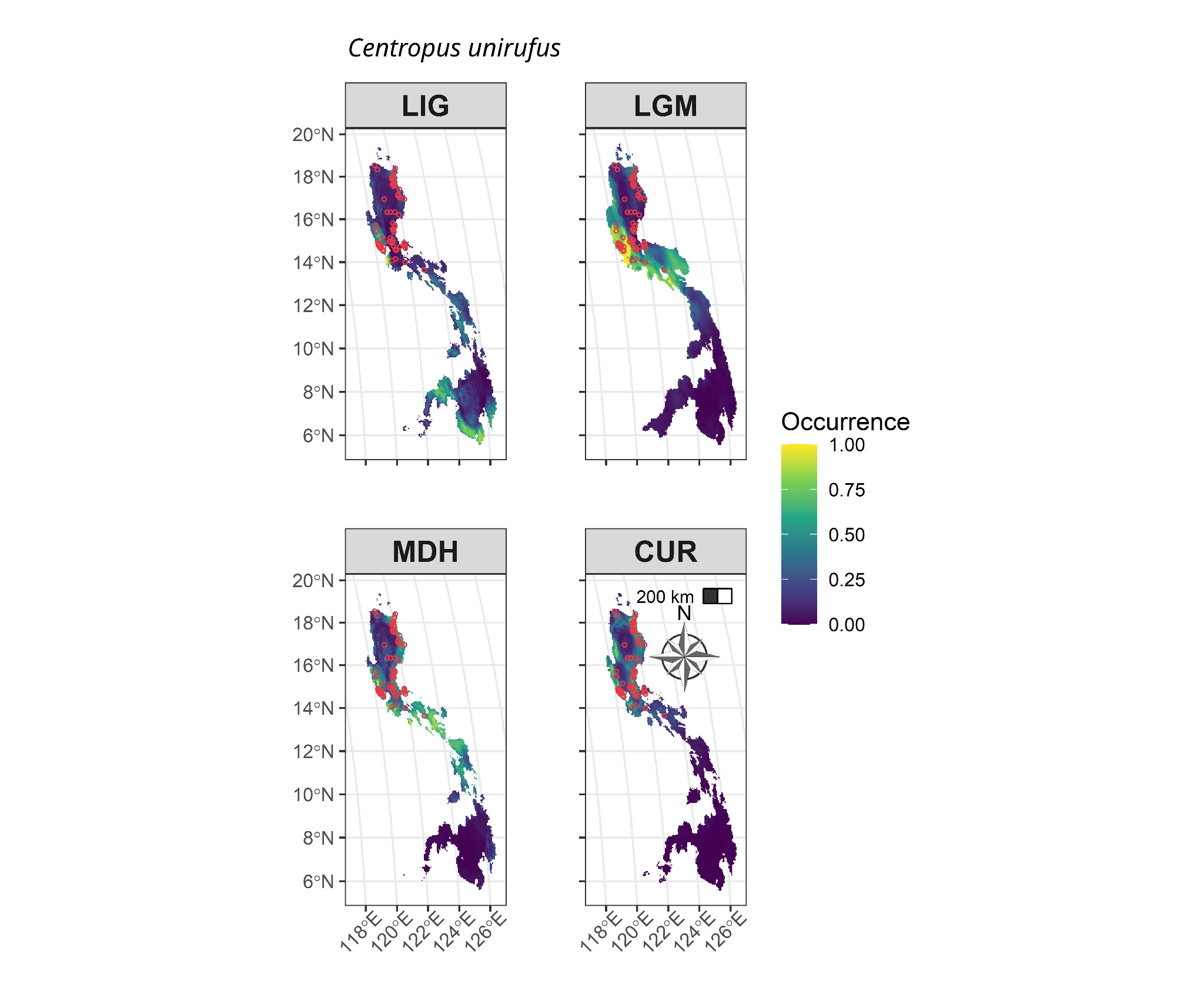

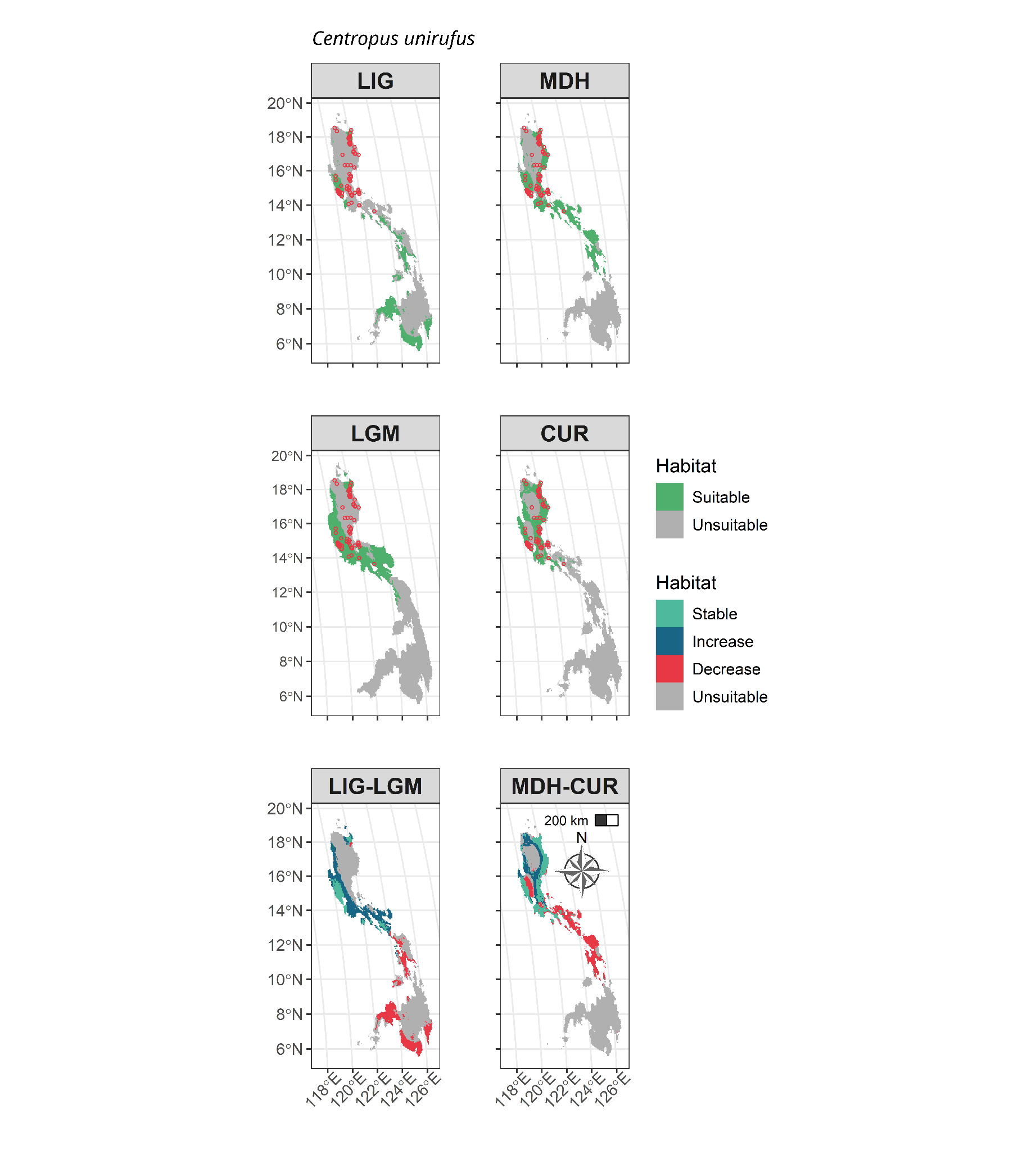

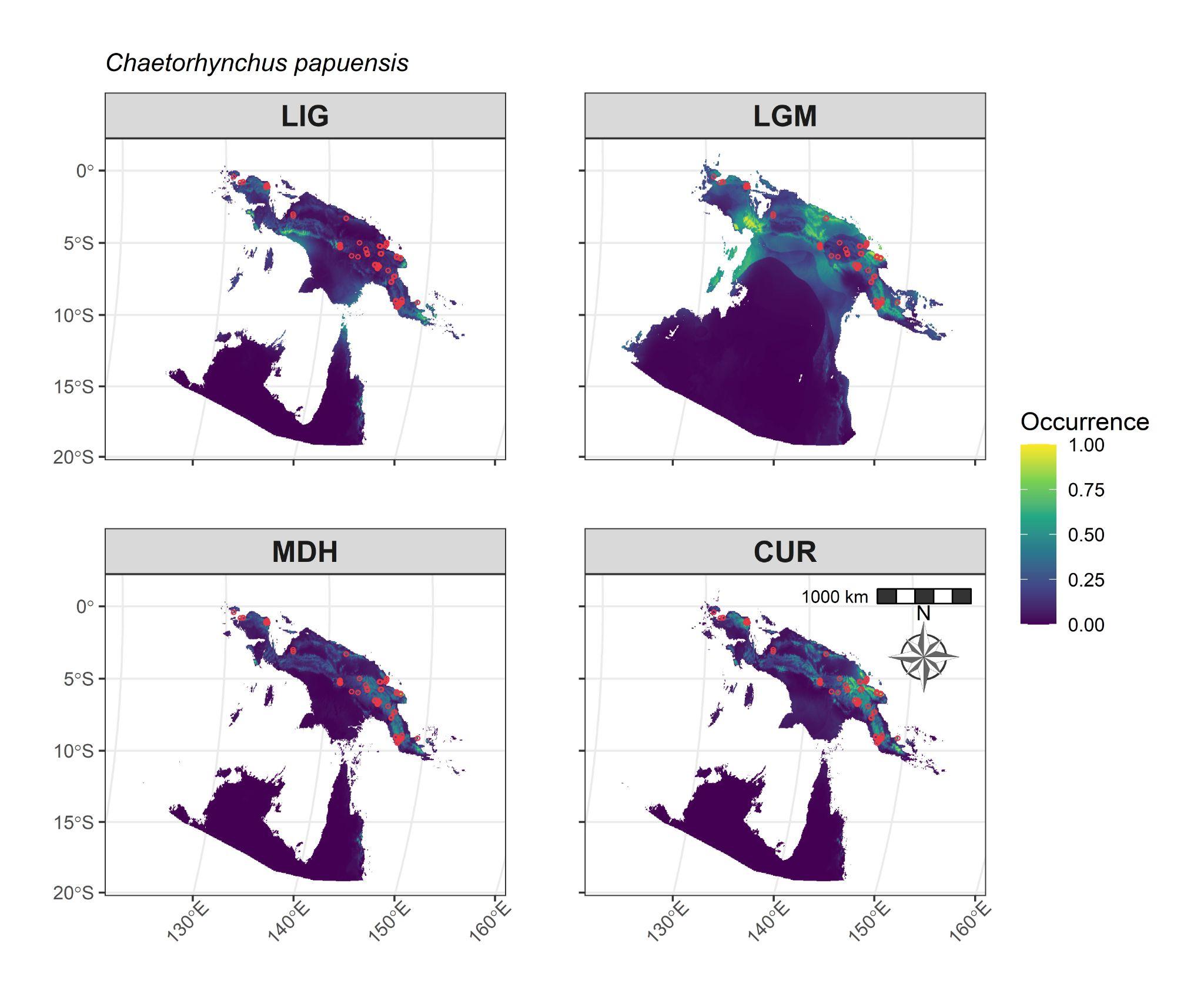

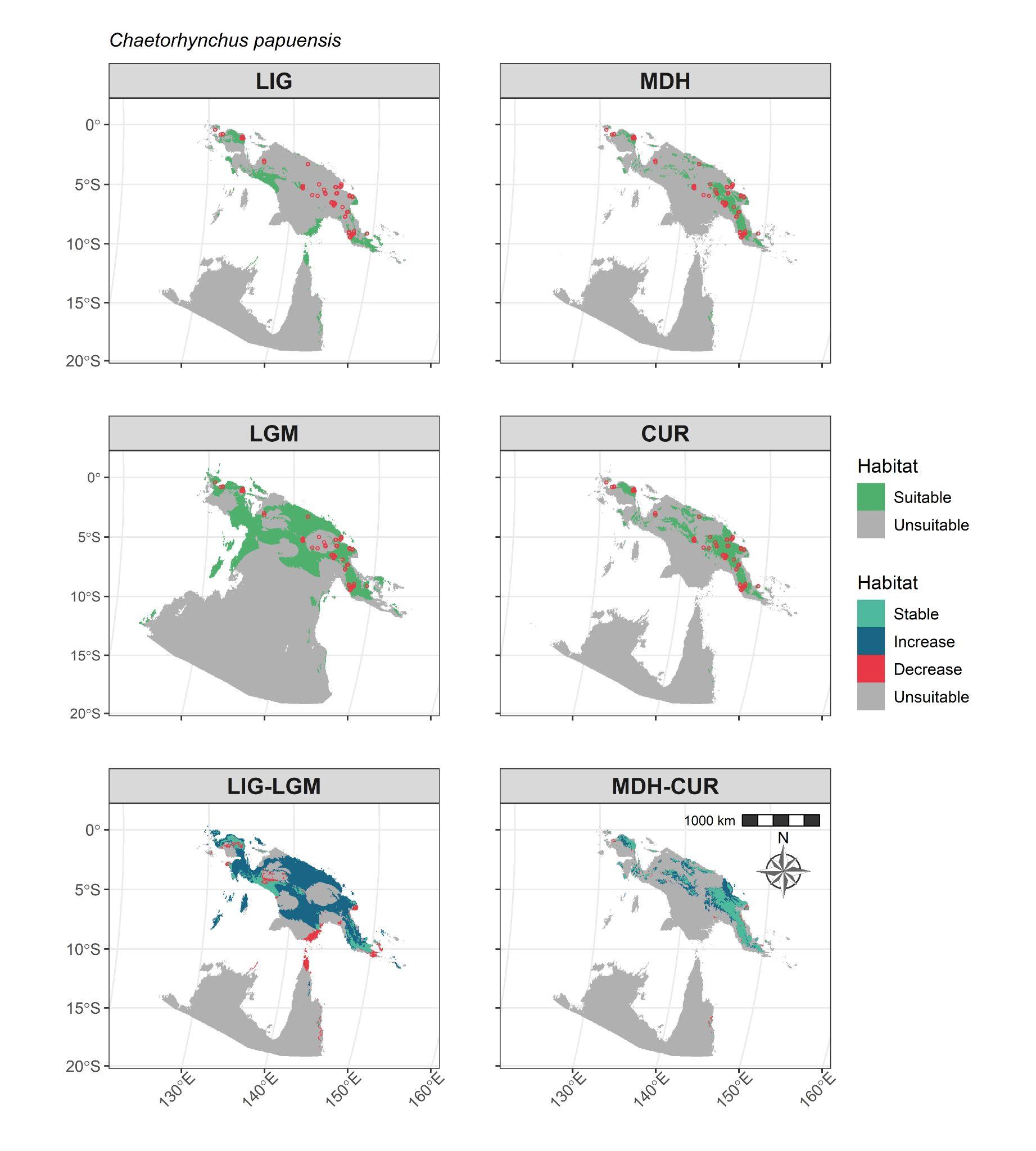

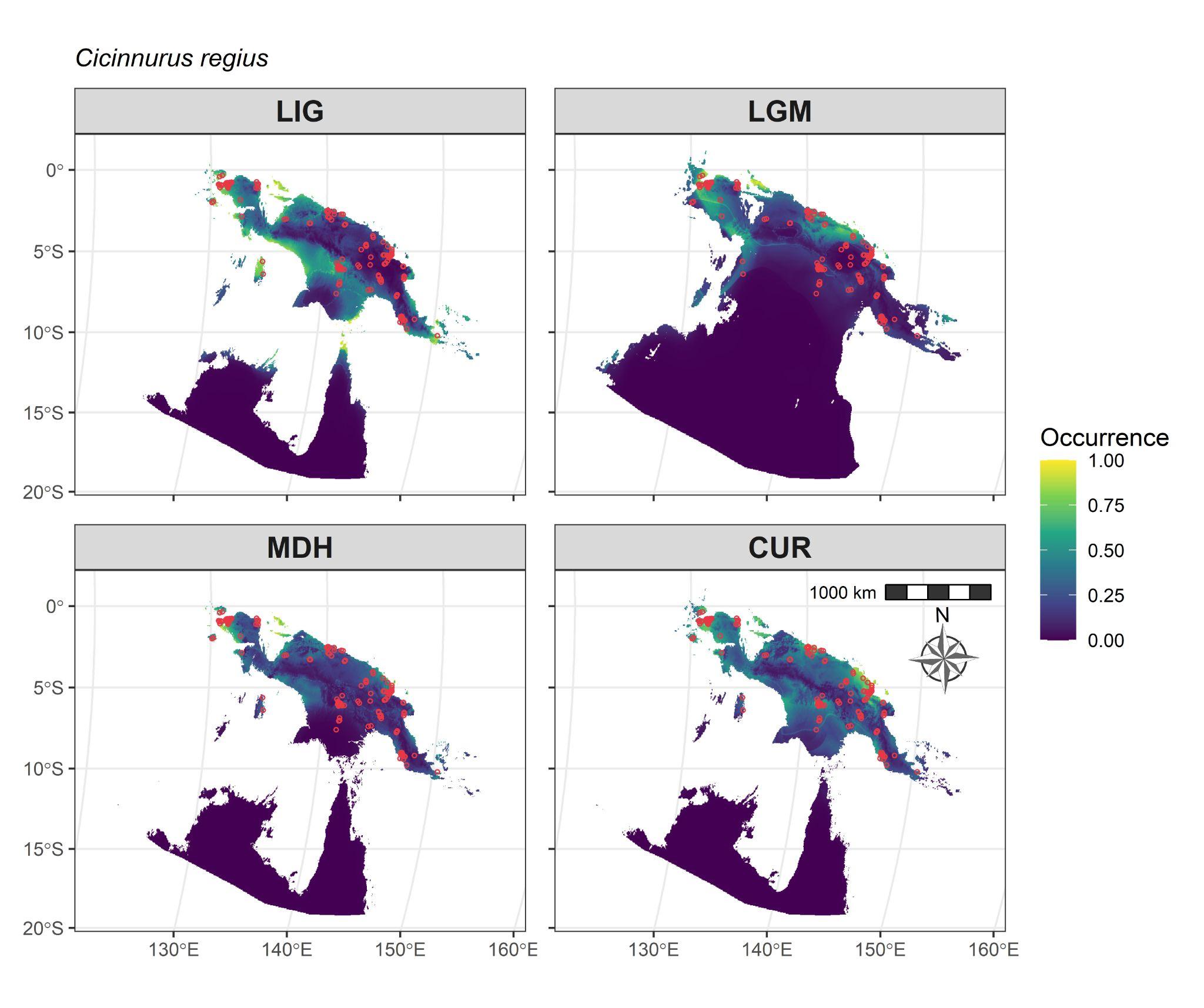

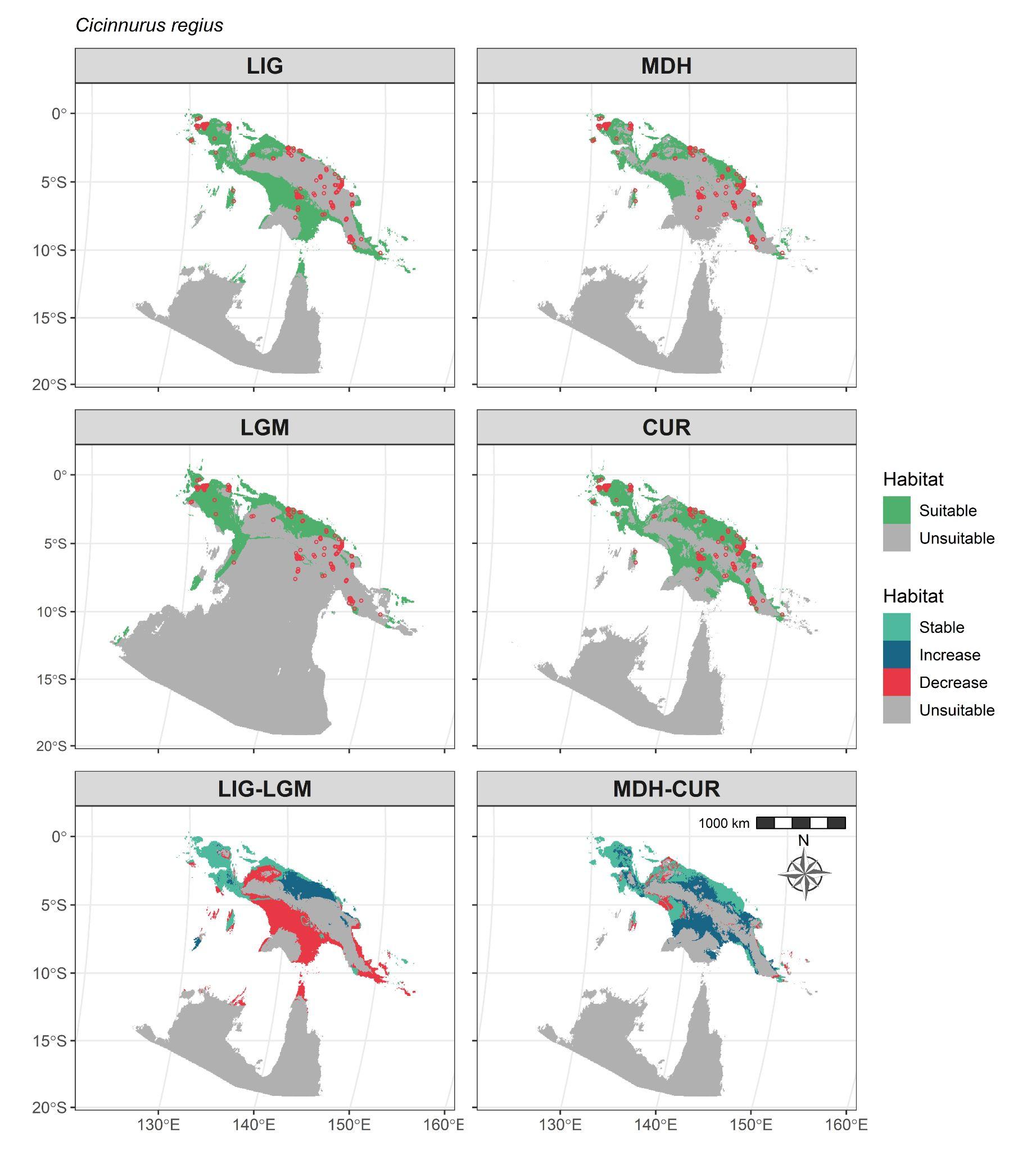

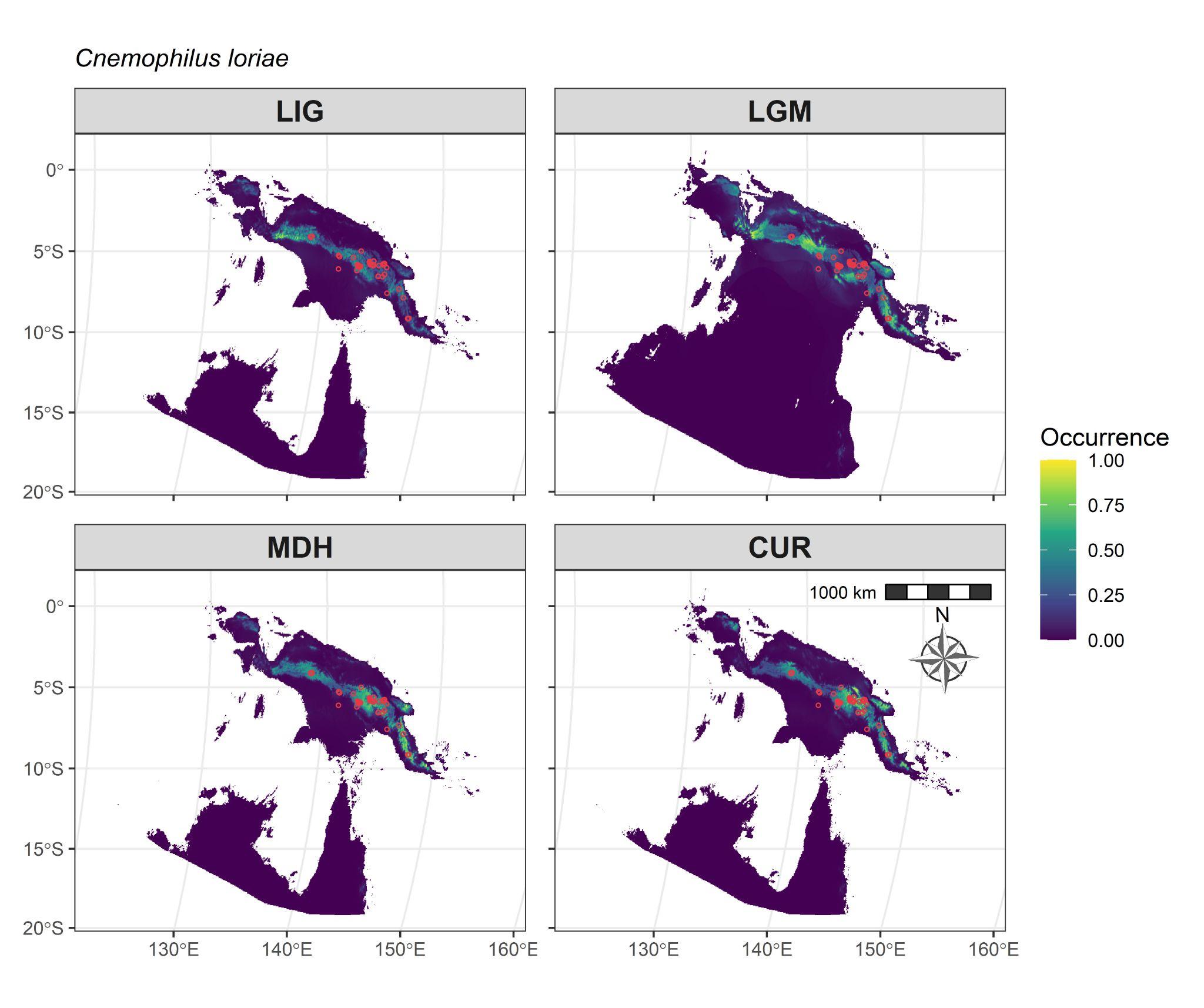

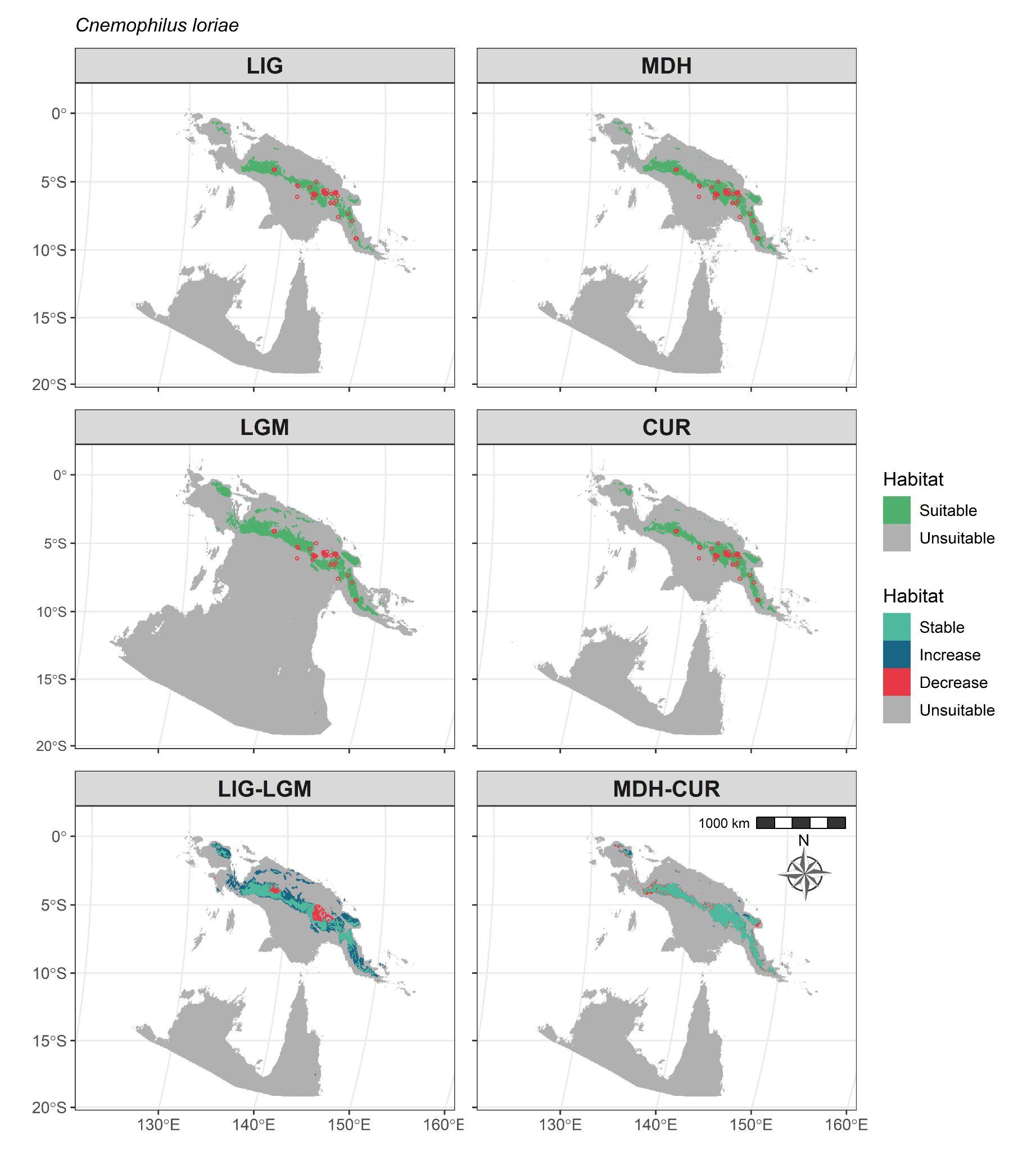

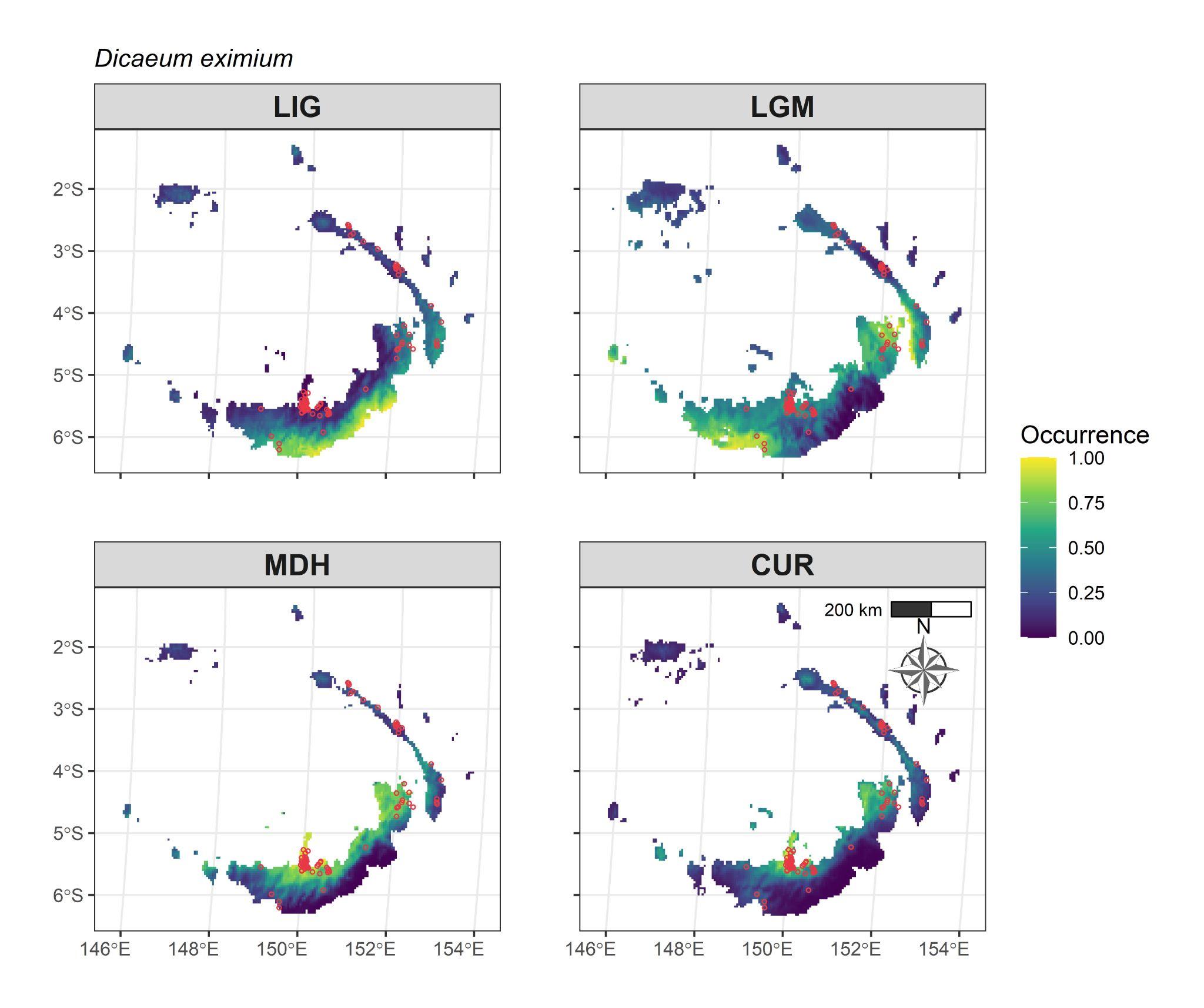

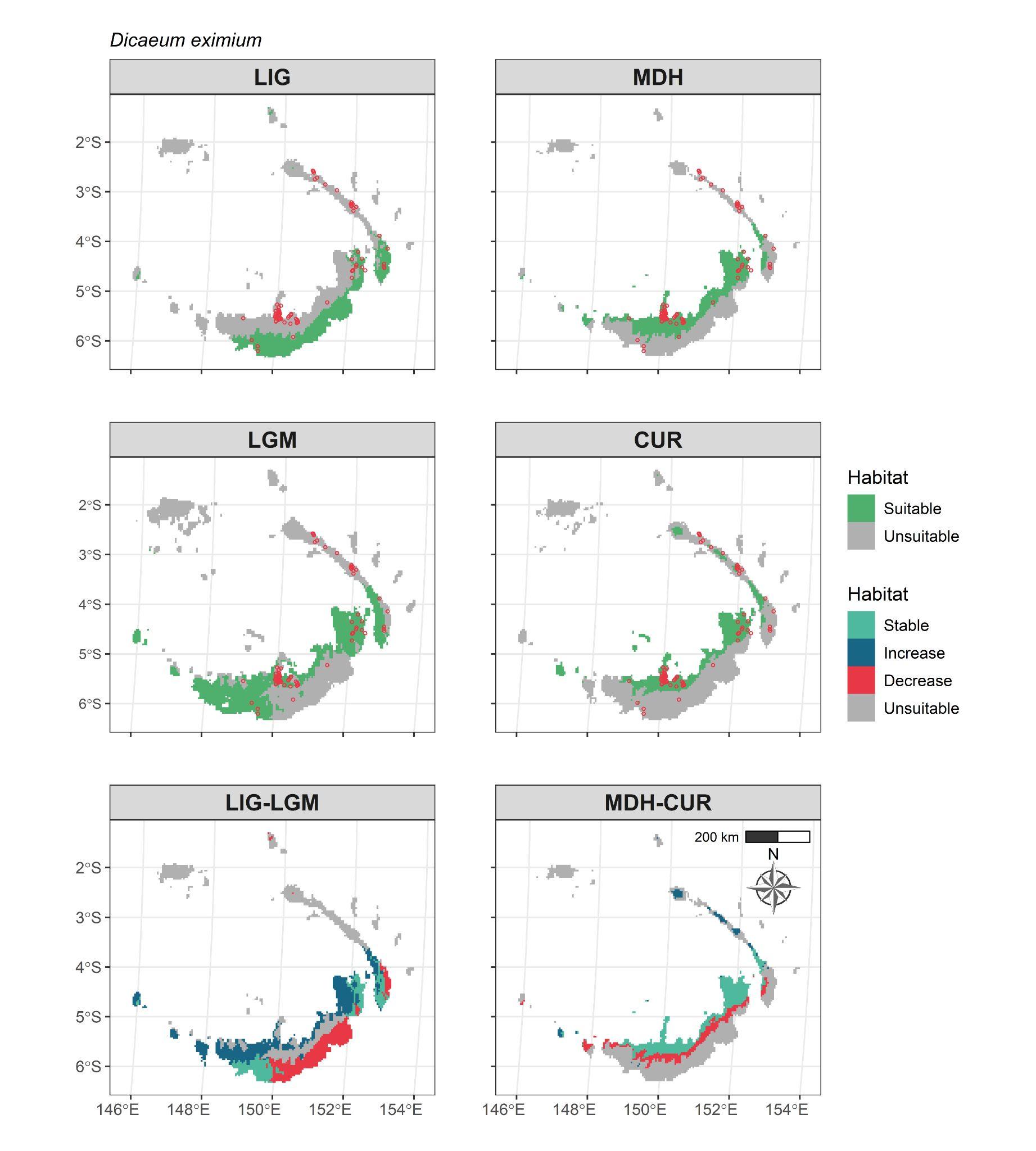

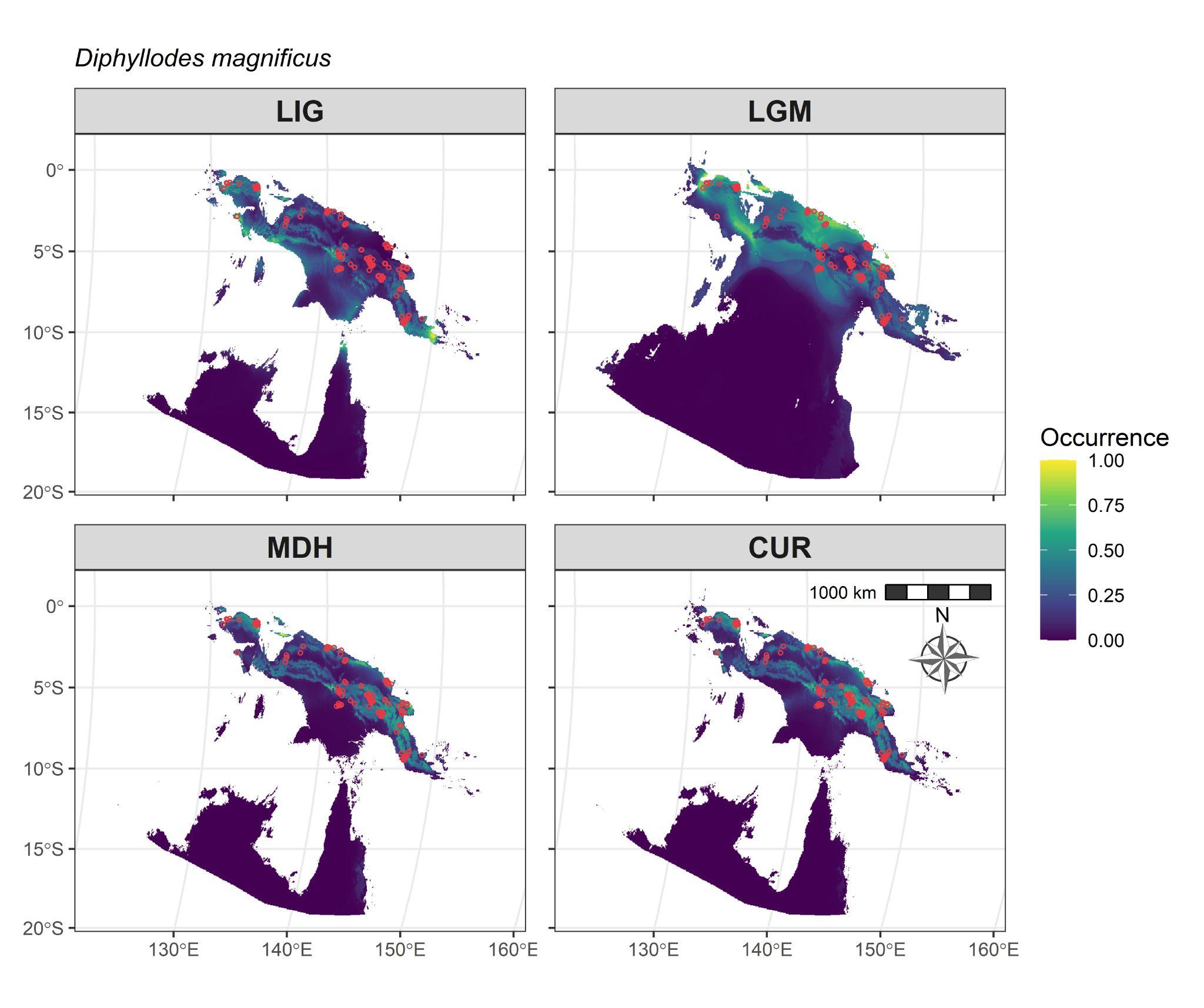

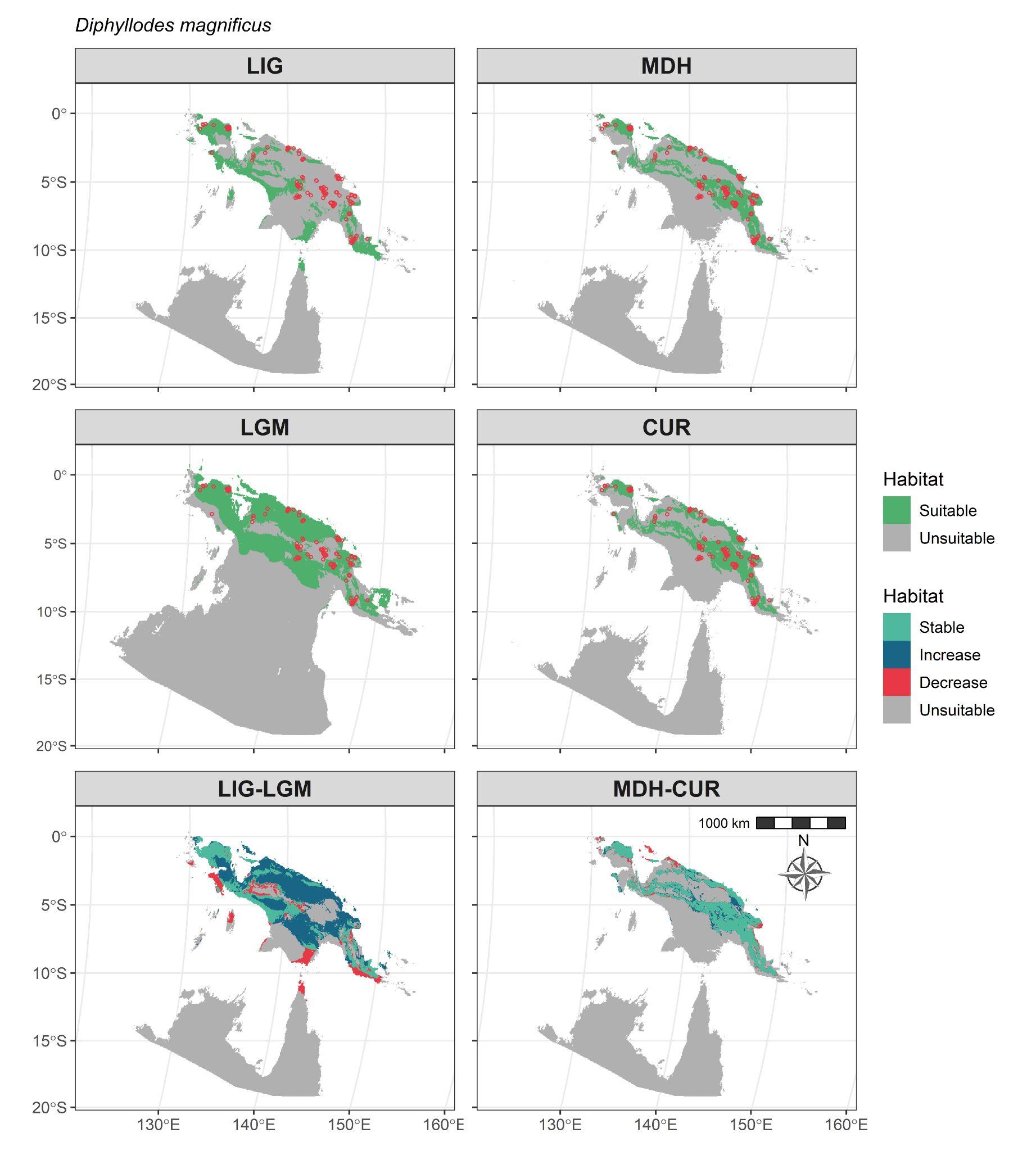

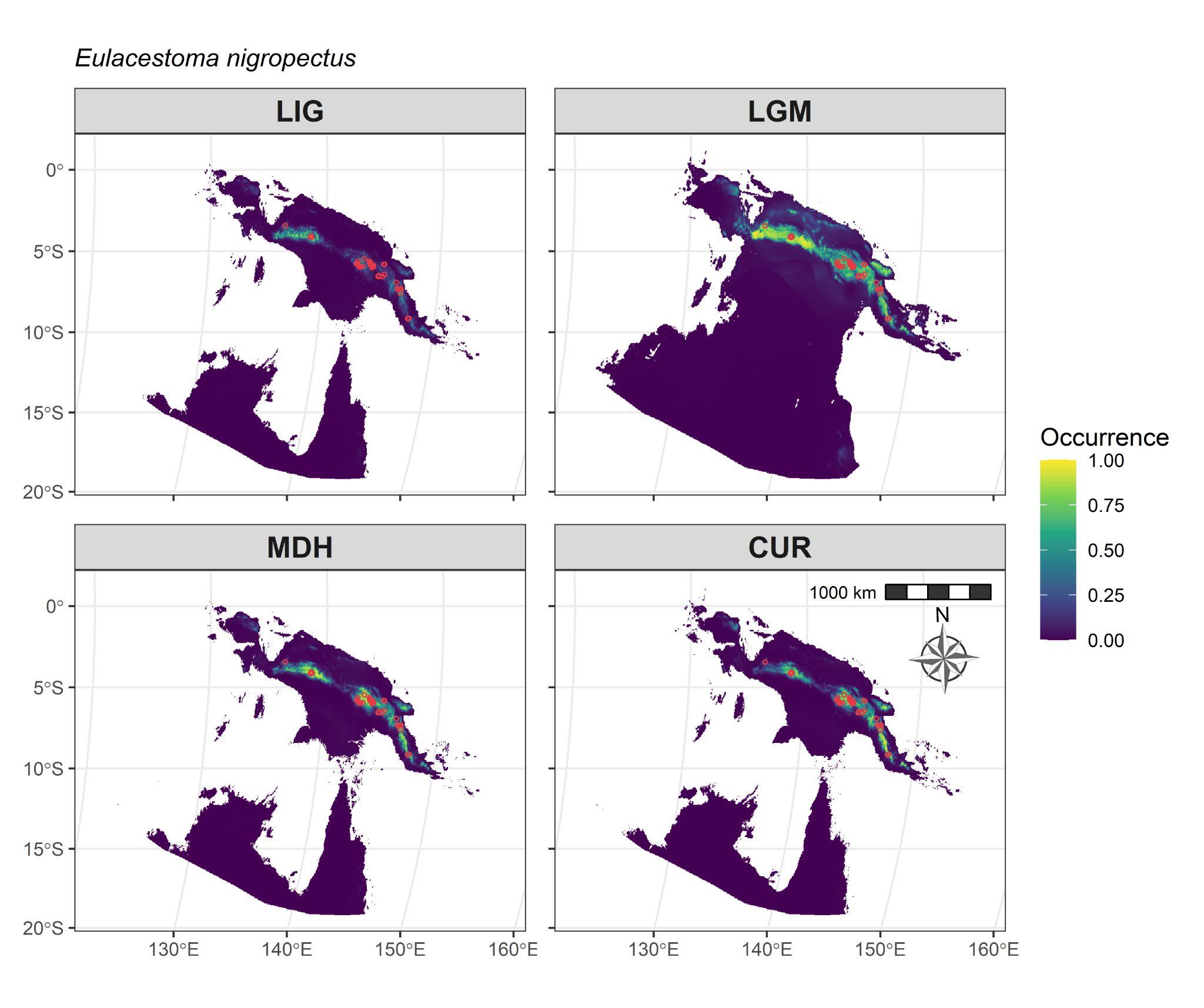

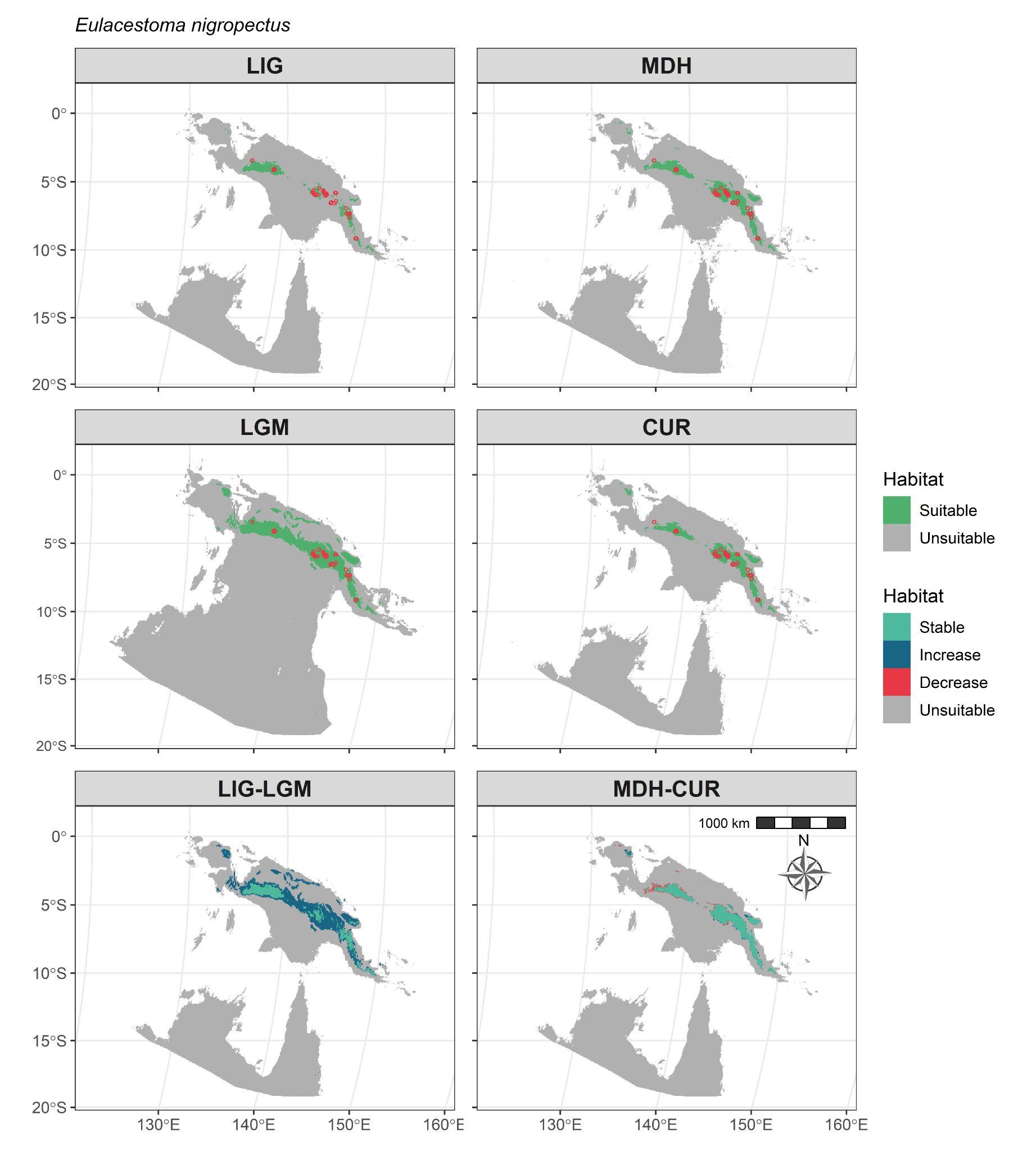

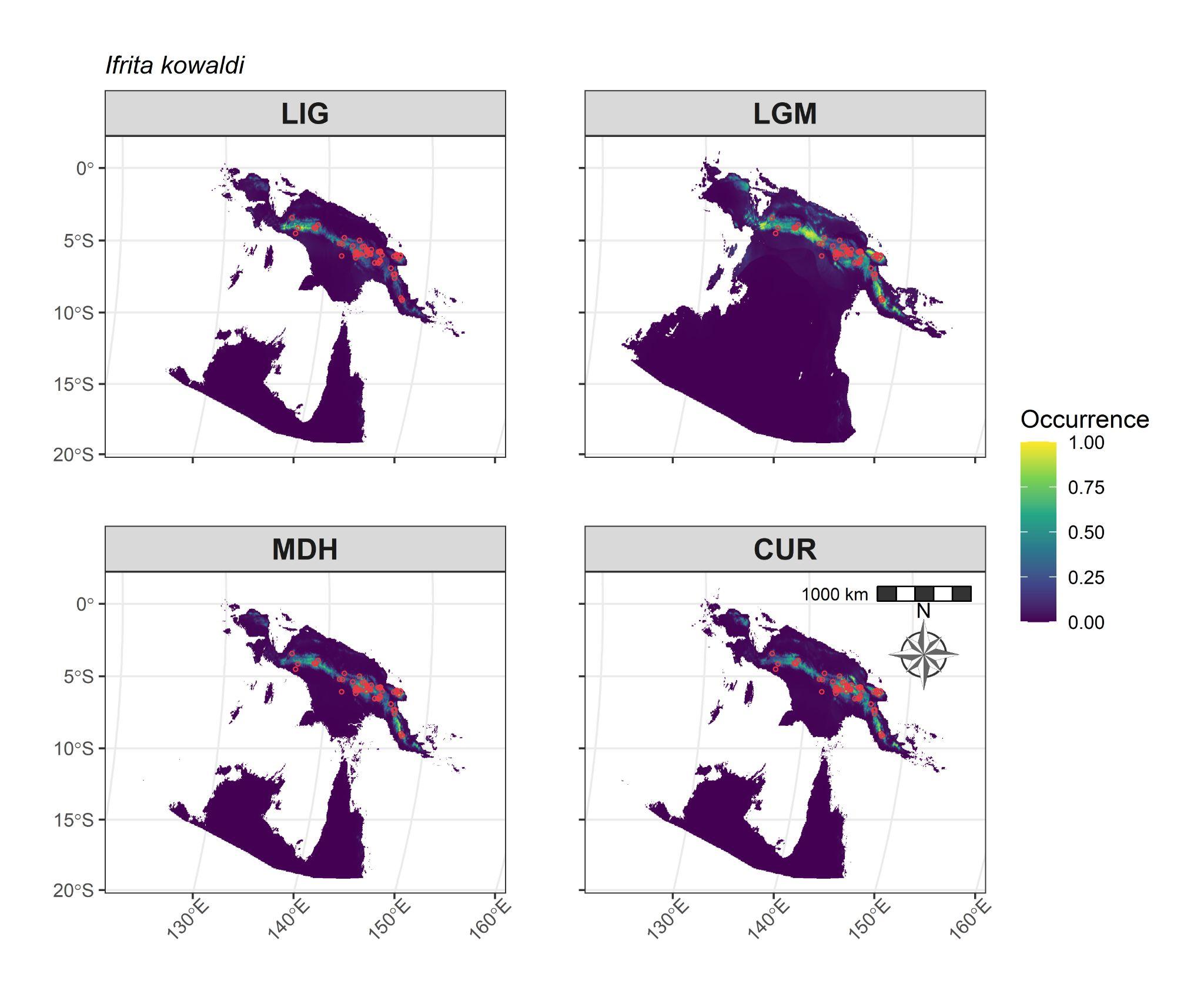

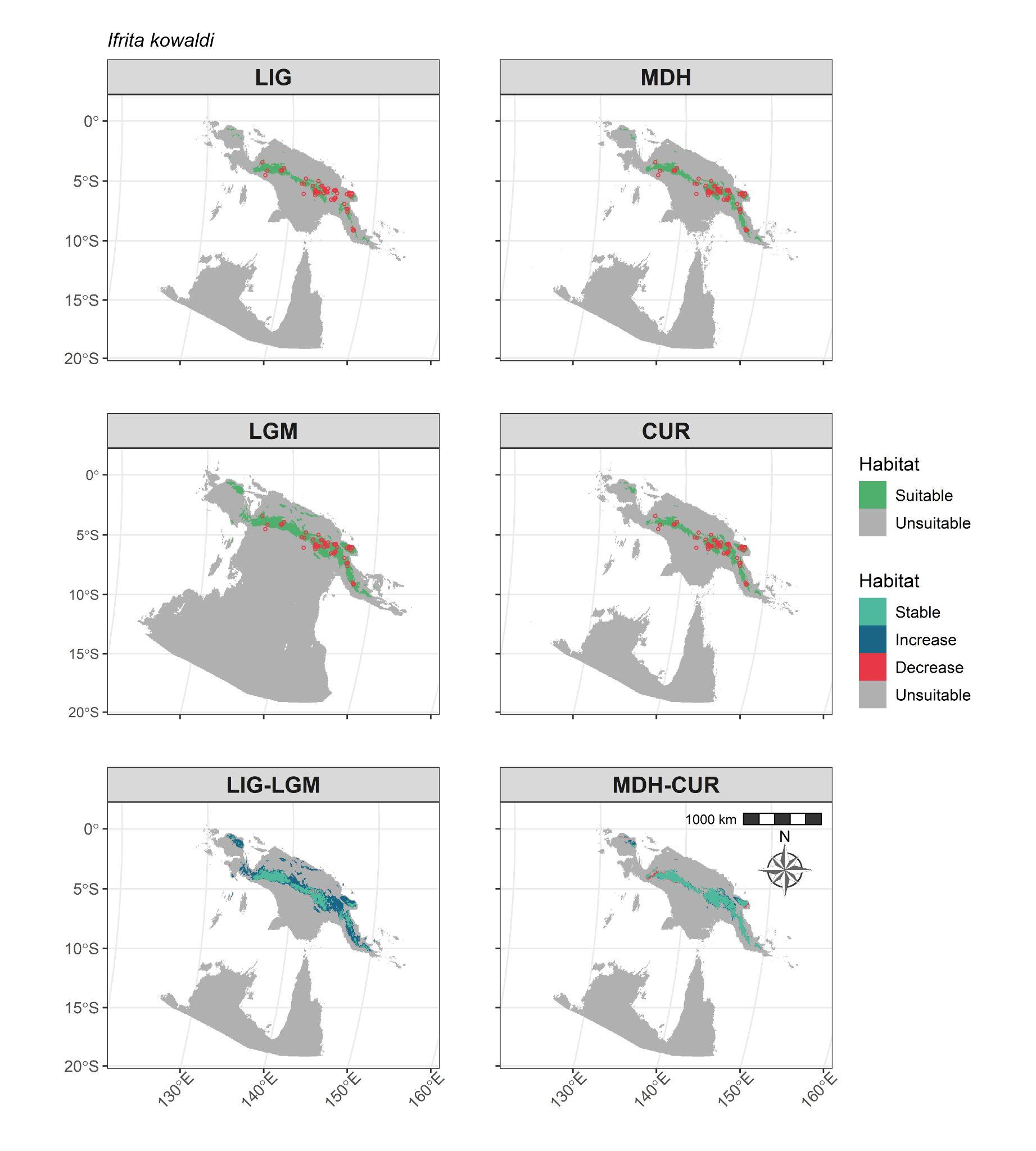

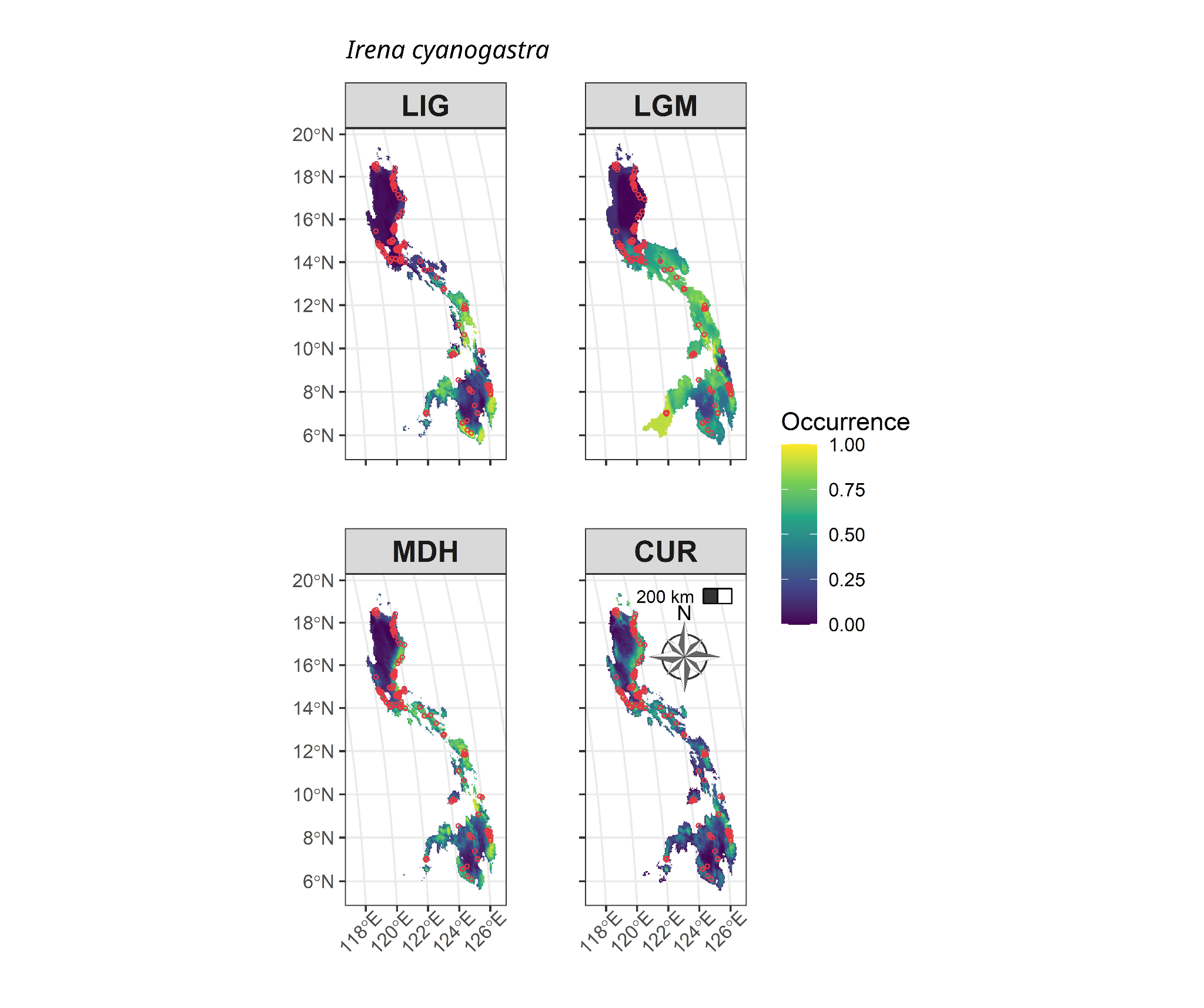

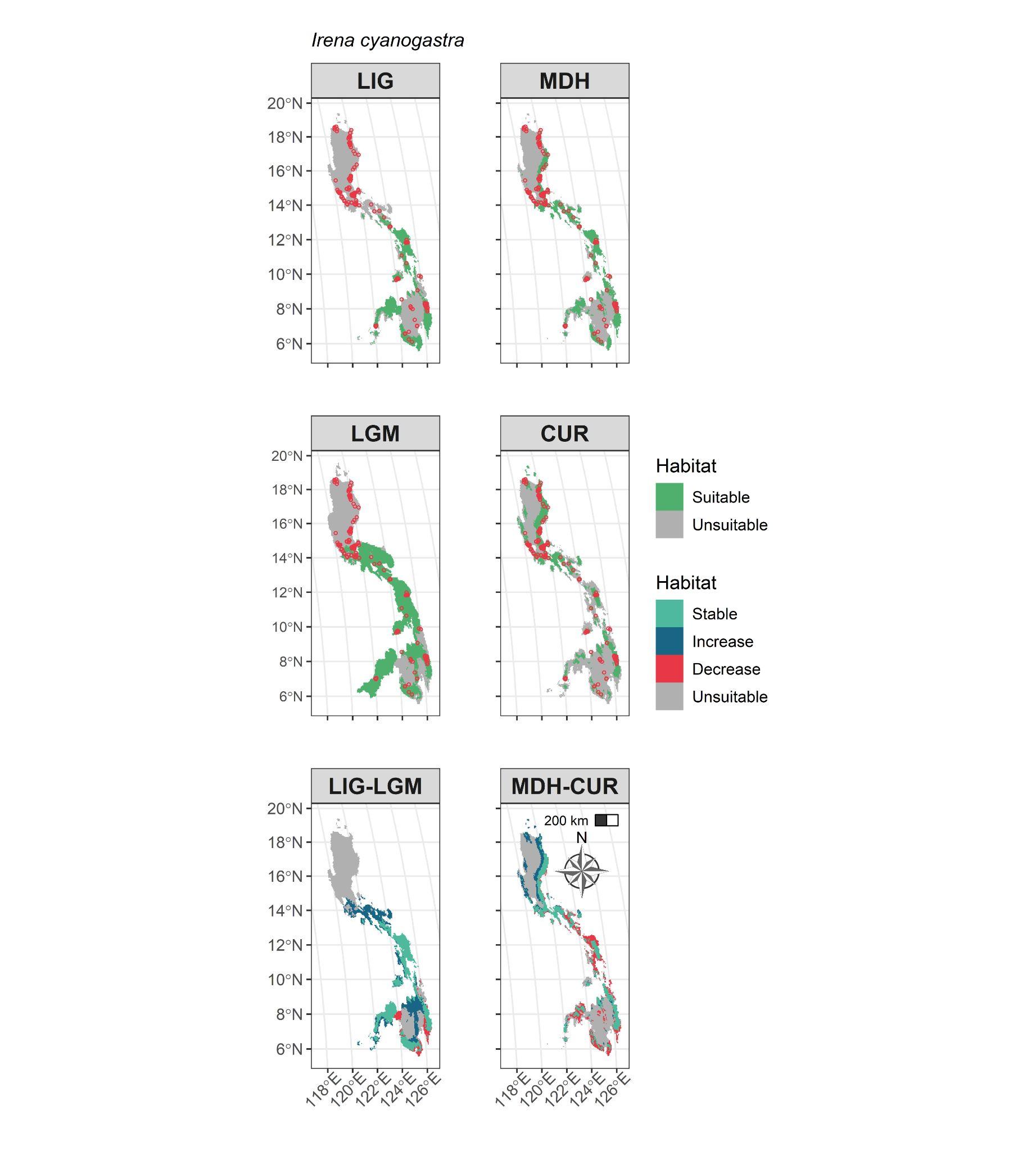

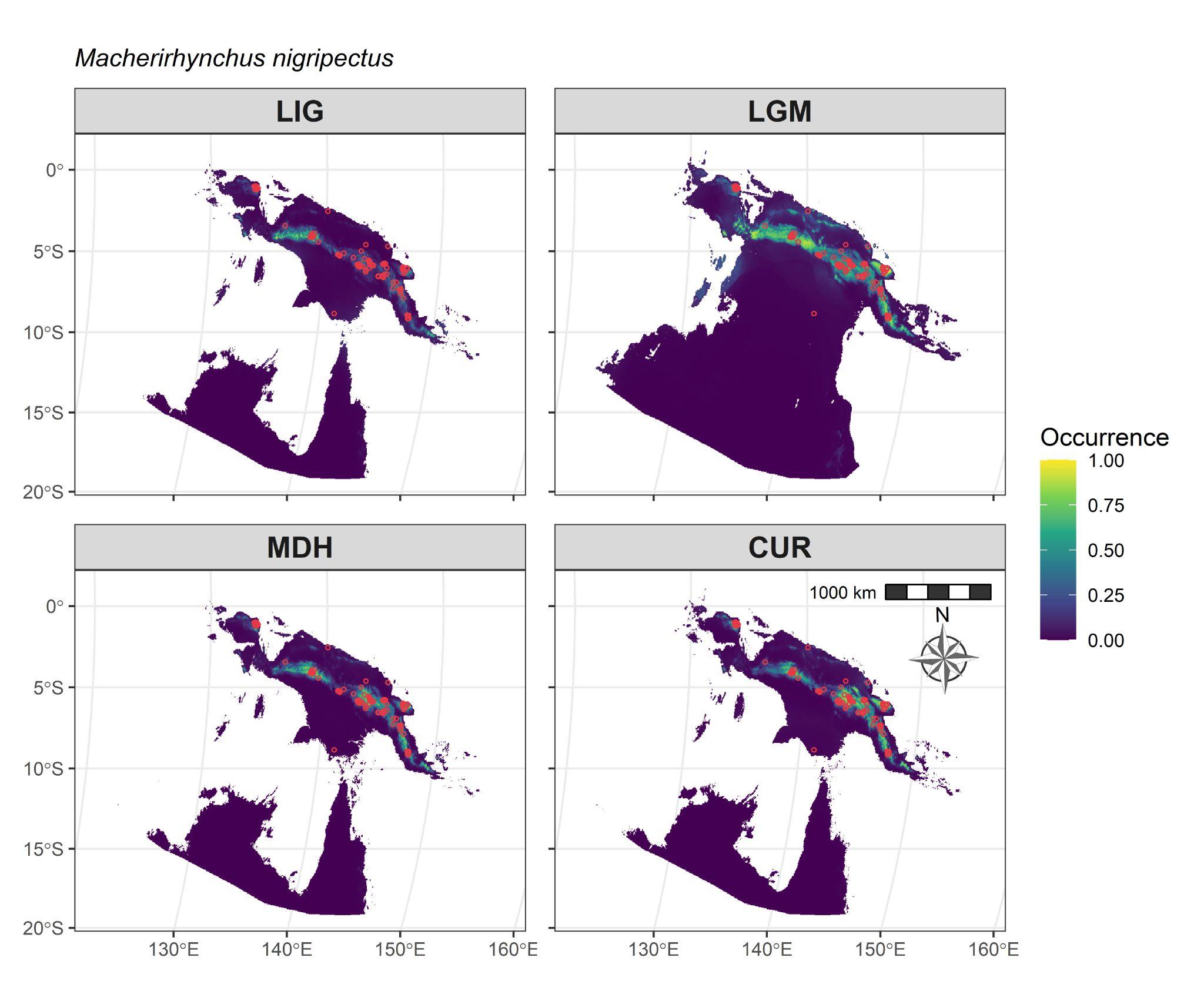

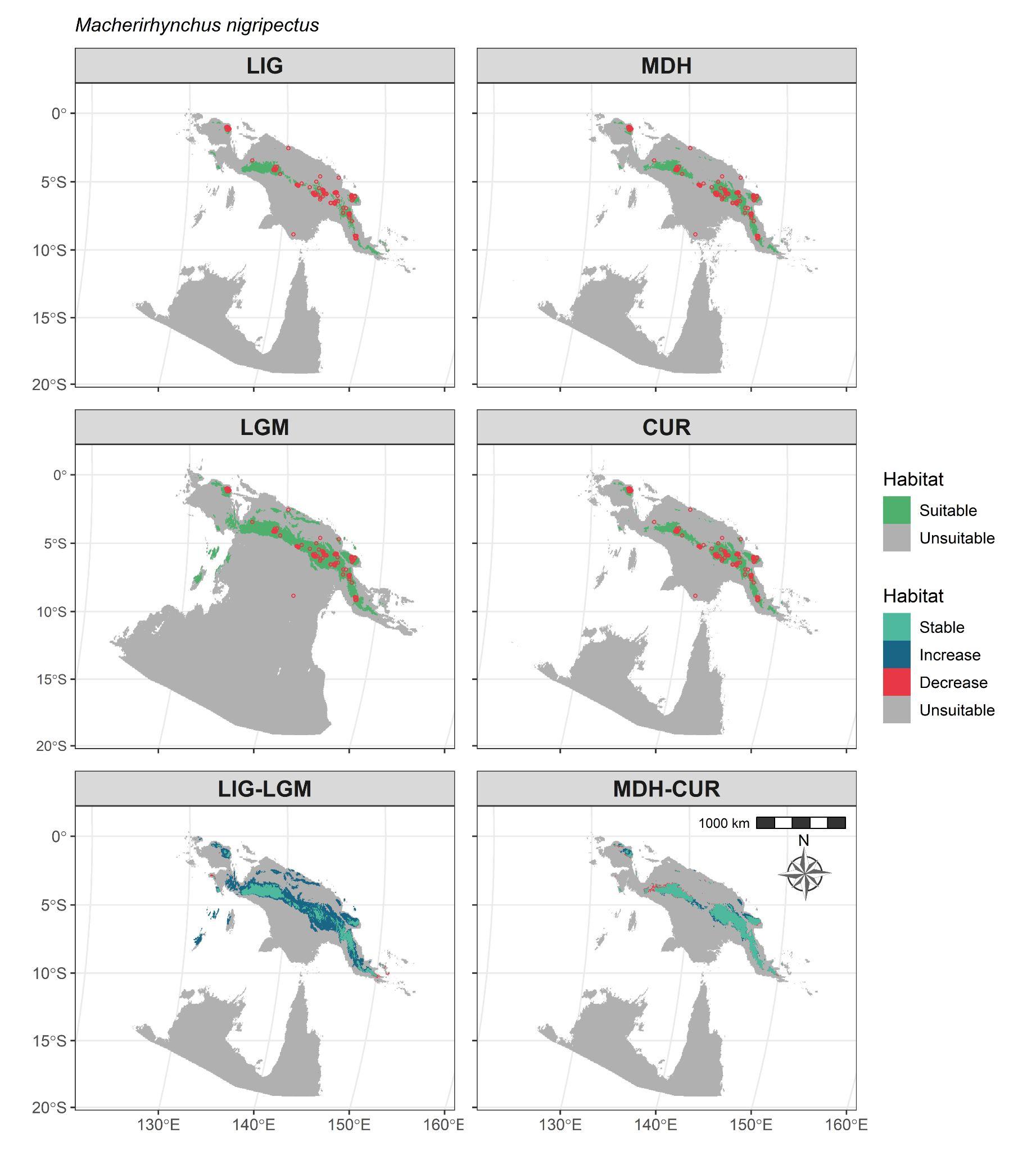

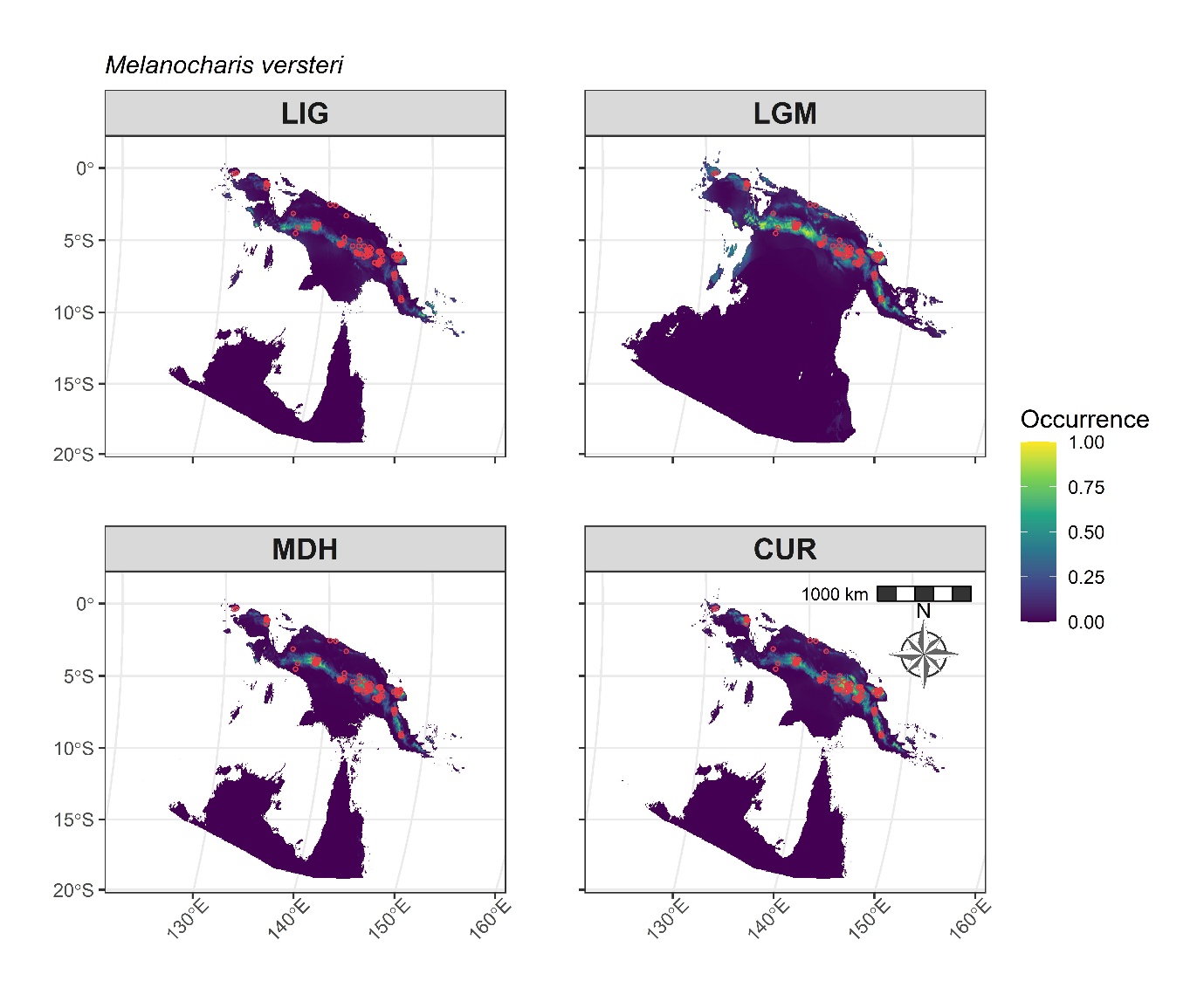

### **Figure S3: PSMC comparisons between methods**

### Comparisons between PSMC plots with different methods of sex chromosome exclusion for five species. Red line = Scaffolds mapping to sex chromosomes removed after synteny analysis. Blue line = SNPs selected only from autosomal regions using the bcftools pipelines.

### *Amazona vittata*

### *Dicaeum eximium*

### *Diphyllodes magnificus*

### *Irena cyanogastra*

### *Rhynochetos jubatus*

### **Figure S4: Boxplot of PSMC comparisons between methods**

### Comparisons of Effective Population Size (Ne) at the LIG and LGM across methods of removing sex chromosome reads. noBam = Scaffolds mapping to the sex chromosomes were removed after synteny analysis. Bam = SNPs selected only from autosomal regions using the bcftools pipelines. Ne values for the five species are overlaid in blue.

### **Figure S5: PSMC Bootstrapping results across the LGP**

### Comparisons of Effective Population Size (Ne) at the Last Interglacial (LIG) and Last Glacial Maximum (LGM) incorporating bootstrapped Ne values. Boxplots display bootstrapped Ne values, and blue points display the non-bootstrapped Ne value. Outlying bootstrapped and non-bootstrapped Ne values are not displayed. “***” indicates p < 0.001, and “*” indicates p < 0.05. Sequentially, the plots display Ne values using the –p “1 + 1 + 1 + 1 + 30 * 2 + 4 + 6 + 10”, –p “2 + 2 + 30 * 2 + 4 + 6 + 10”, and –p “4 + 30 * 2 + 4 + 6 + 10” PSMC settings respectively.

### Figure S6: Bayesian Multi Level Model results

(A) Trace plots from the best Bayesian Multilevel Model performed (model name = brms11), displaying the sampling behaviour for each parameter across four chains. Each panel corresponds to a specific parameter in the model, with individual chains indicated by different shades. The x-axis represents the iteration number, and the y-axis shows the sampled values. These indicate that our model has converged for each chain.

(B) A pair plot displaying the relationships among various parameters from the best Bayesian Multilevel Model performed. Each row and column pair represents an interaction between model parameters Each parameter’s posterior distribution is shown on the diagonal. The off-diagonal cells contain scatter plots showing how pairs of parameters interact. Each point in the scatter plots represents a sample from the posterior distribution. Scatterplots display positive or negative slopes for any parameter pairs indicating the pair distribution influenced by particular predictors.

**SUPPLEMENTARY TABLES**

### Table S1: Details of the bioclimatic variables used for Ecological Niche Modelling (ENM).

Table S2: Details of the total suitable land area estimated for each species in our panel.

Table S3: Details of PSMC analyses done. Effective population size (Ne) values at various time points were approximated from graphs. LIG = Last Interglacial, LGM = Last Glacial Maximum.

Table S4: Concordance of the direction of change in Ne during the LGP as inferred from boxplots.

Table S5: Model coefficients of the best performing Bayesian model.

Table S6: Life history and trait details of the species. Unavailable data are left blank.

Table S7: Data sources for the genomes used in our study.

Table S8: Details of Global Biodiversity Information Facility (GBIF) keys used to access the occurrence data points used in this study.

Table S9: Details of the Ecological Niche Modelling (ENM) analyses done.

Table S10: Details of the Bayesian multilevel models tested. Models are arranged in descending order of model performance.
